# Ancient tropical extinctions contributed to the latitudinal diversity gradient

**DOI:** 10.1101/236646

**Authors:** Andrea S. Meseguer, Fabien L. Condamine

## Abstract

Biodiversity currently peaks at the equator, decreasing toward the poles. Growing fossil evidence suggest that this hump-shaped latitudinal diversity gradient (LDG) has not been persistent through time, with similar species diversity across latitudes flattening out the LDG during past greenhouse periods. This provides a new starting point for LDG research. Most studies assume the processes shaping the LDG have acted constantly through time and seek to understand why diversity accumulated in the Holarctic at lower levels than at the equator, *e.g.* as the result of limited dispersal, or higher turnover in Holarctic regions. However, fossil investigations suggest that we need to explain when and why diversity was lost at high latitudes to generate the LDG. Unfortunately, diversity lost scenarios in the Holarctic have been repeatedly proposed but not yet clearly demonstrated. Here, we use diversification approaches for both phylogenies and fossils to study the LDG of Testudines, Crocodilia and Lepidosauria. We show the LDG of these groups has varied over time, with high latitudes serving as a source of tropical diversity but suffering disproportionate extinction during transitional periods to cold climate. We outline the ‘asymmetric gradient of extinction and dispersal’ (AGED) framework, which contextualizes previous ideas behind the LDG under a time-variable scenario. We suggest the current steep LDG may be explained by the extinction of clades adapted to warmer conditions from the new temperate regions formed in the Neogene, together with the equator-ward dispersal of organisms tracking their own climatic preferences, when tropical biomes became restricted to the equator. Conversely, high rates of speciation and pole-ward dispersals can account for the formation of an ancient flat LDG during the Cretaceous–Paleogene greenhouse period. Our results demonstrate that the inclusion of fossils in macroevolutionary studies allows detecting extinction events less detectable in analyses restricted to present-day data only.

## Introduction

The current increase in species richness from the poles toward the equator, known as the latitudinal diversity gradient (LDG), is one of the most conspicuous patterns in ecology and evolution. This pattern has been described for microbes, insects, vertebrates, and plants, and for marine, freshwater, and terrestrial ecosystems^1–6^.

For decades, it has been thought that the modern-type steep LDG (with higher diversity at the equator) persisted throughout the Phanerozoic (the last 540 million years), even if the gradient was sometimes shallower^7^, based on published fossil record studies^8, 9^. However, the methodological limitations of fossil sampling have called this conclusion into question. Analyses controlling for sampling bias have suggested that, for many groups, the LDG was less marked in the past than it is today, and was flat (*i.e.* with similar species diversity across latitudes) or even developed a paleotemperate peak during some periods (see^10^ for a review). This sampling-corrected flatter LDG in deep time has been demonstrated for non-avian dinosaurs^11^, mammals^12, 13^, tetrapods^14^, insects^15–17^, brachiopods^18–20^, bivalves^21^, coral reefs^22^, crocodiles^23^, turtles^24, 25^, and plants^26–28^. The pattern emerging from fossil studies also suggests that steep LDGs, such as that currently observed, have been restricted to the relatively small number of short coldhouse periods during the history of the Earth: the Ordovician/Silurian, the Carboniferous/Permian, the end of the Jurassic, and the Neogene. Most of the Phanerozoic has instead been characterized by warm greenhouse climates associated with a flatter LDG^1^ (**Fig. 1**).

**Figure 1.**
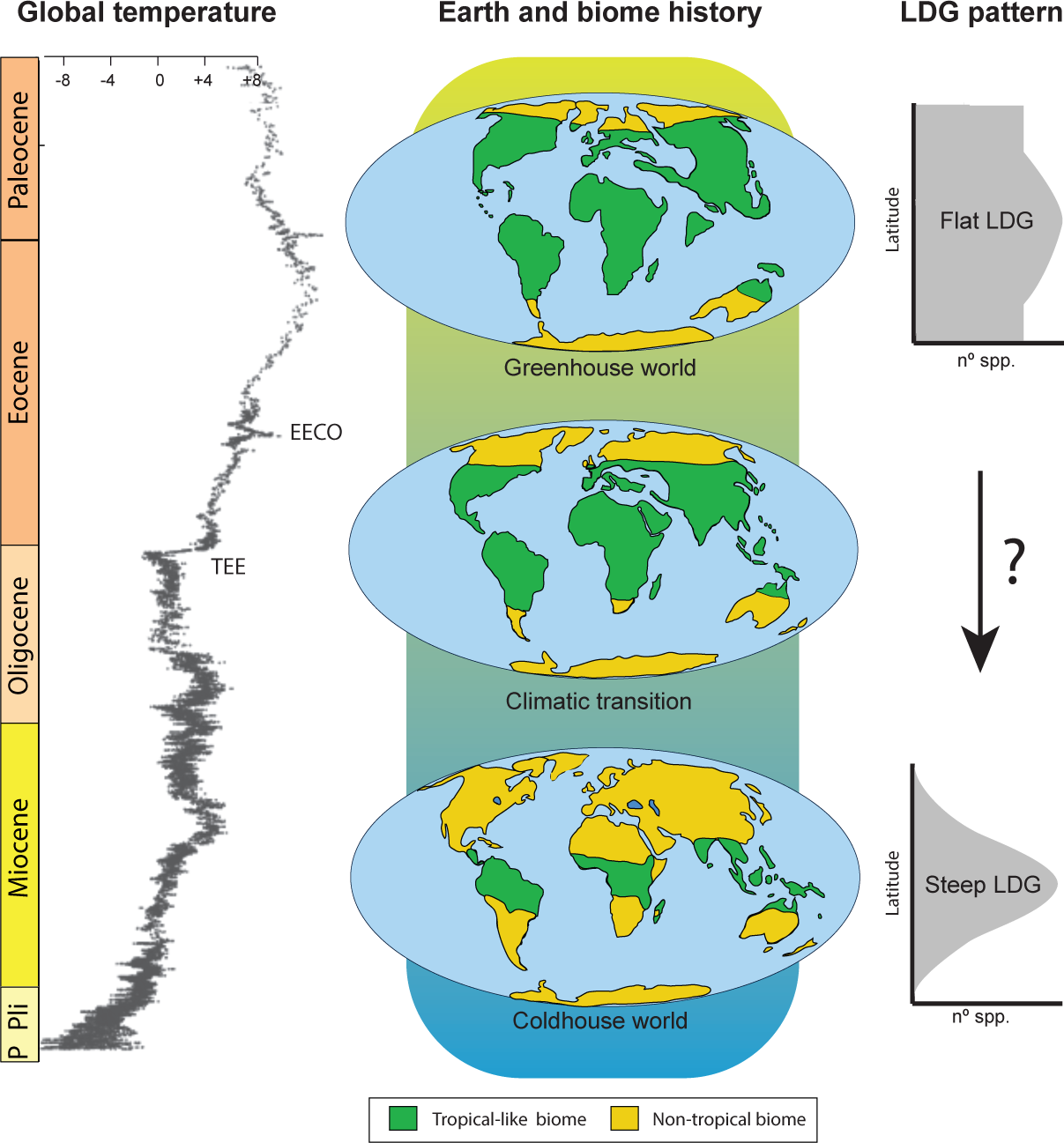
Changes in global temperature and extension of the tropical belt during the Cenozoic, in relation with the shape of the LDG. Early Cenozoic global temperatures were higher than today and paratropical conditions extended over northern and southern latitudes. From the early Eocene climatic optimum (EECO; ca. 53-51 Ma), a global cooling trend intensified on Earth and culminated with the Pleistocene glaciations. Global cooling was punctuated by sharp declines of temperatures and periods of relative warmth. Warm-equable regimes got then restricted to the equator. The LDG evolved following these global changes; during greenhouse periods diversity was uniform across latitudes, such that the LDG flattened, while in cold periods diversity peaked at the equator (a steep LDG)^10^. The question mark denotes the focus of this study, which is to unveil the processes that mediated the transition between a flat and steep LDG. The relative temperature curve of the Cenozoic is adapted from^73^. Maps represent the extension of the tropical belt and Earth tectonic changes as derived from^70, 71^. P=Pleistocene, Pli=Pliocene.

This recent fossil evidence provides a new starting point for LDG research. Up to now, most studies have explained the origin of the LDG as a result of greater tropical diversification and limited dispersal out of the equatorial region^7, 29, 30^, or by high rates of turnover in the Holarctic (*i.e.* similar high speciation (*λ*) and extinction (*μ*) rates; *λ*≈*µ*) (**Table 1**) for amphibians^31, 32^, birds^33–35^, butterflies^36^, conifers^37^, fishes^38^, mammals^30, 35^, and lepidosaurs^39^. Behind these ideas rely the assumption that the equatorial regions are the source of world diversity^40, 41^, and the LDG resulted from lower levels of diversity accumulation in the Holarctic than at the equator through time^7, 29, 33^, hereafter referred to as ‘slow Holarctic diversity accumulation’ hypotheses, which in turns implies that the current steep shape of the LDG, and so the processes driving this pattern, have been persistent through time. However, the recent fossil investigations showing, for many lineages, similar diversity levels across latitudes in the past suggest we do not necessarily need to explain why diversity accumulated at slower rates in the Holarctic through time, but the question being how and when diversity was lost at high latitudes, giving rise to the current shape of the LDG^10^?

**Table 1.**
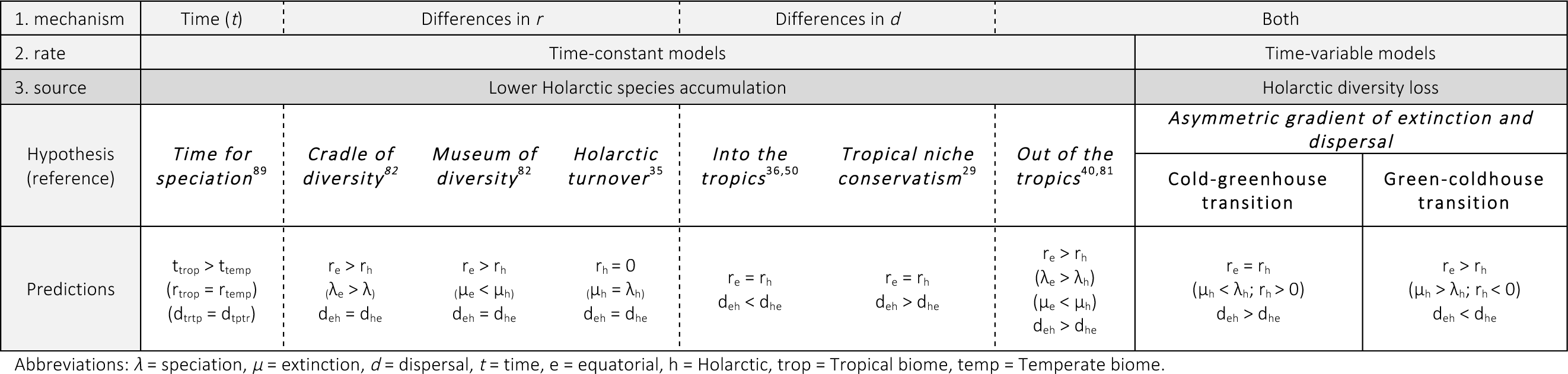
Predictions of the most common LDG hypotheses, including the *Asymmetric gradient of tropical extinction and dispersal* (AGED) proposed here. The main evolutionary hypotheses can be classified according to three criteria: (1) the mechanisms behind regional differences in species richness^7^, including explanations based on evolutionary time, on dispersal *(d)*, and on diversification (the composite value *r* = *λ* − *μ*). Explanations based on evolutionary time assume most groups originated in the tropics and had less time to diversify in the temperate regions^89^. Hypotheses focusing on the role of geographic movements *(d)*, include the “*tropical niche conservatism*” model assuming that most groups originated in the tropics and the LDG results from limited dispersal to the temperate regions, as only few tropical species succeeded to adapt to temperate regimes^41^. The “*Into the tropics*” model assumes instead the LDG results from dispersals towards the equator^36, 50^. Hypotheses that emphasize the LDG is generated by regional differences in net diversification rates, being higher in the tropics^31, 33^ assume the outstanding tropical diversity is the outcome of higher rates of speciation in the tropics than in the extra-tropical regions (*λ*_*t*_ > *λ*_*e*_) under the “*cradle of diversity*”, and/or result from lower rates of extinction ((*μ*_*t*_ > *μ*_*e*_) under the “*museum of diversity*”^82^. The LDG could also results from higher turnover rates (*i.e.* higher *λ* and *μ*) in the Holarctic^35^. Diversification and dispersal hypotheses are not mutually exclusive. In the “*out of the tropics*” model the tropics are regarded as both cradle and museum, with lineages preferentially originating in the tropics and expanding into high latitudes^40, 81^. Hypotheses could be classified according to (2) the rate at which processes acted through time to explain the LDG; most studies assumed evolutionary processes acted constantly. The AGED model, conversely, includes various diversification/dispersal parameters for each temporal interval (the transition from coldhouse to greenhouse, and *viceversa*). (3) Finally, hypotheses can be classified according to the source of tropical diversity: “*Lower Holarctic species accumulation”* hypotheses assume that the equator is the source of world diversity and species accumulated at slower rates on the higher latitudes. Conversely, “*Holarctic diversity loss*” hypothesis assumes the Holarctic was also a source of diversity but this diversity was lost during evolutionary history.

Diversity losses in the Holarctic have been traditionally considered to underlie the LDG^7, 35^. They were initially attributed to Pleistocene glaciations^42^, but this hypothesis can be called into question since the LDG substantially predates the Pleistocene^7^. More ancient extinctions have also been considered^43–49^. For example, Hawkins *et al.*^46^ suggested the avian LDG resulted from the differential extirpation of older warm-adapted clades from the temperate regions newly formed in the Neogene. Pyron^39^ suggested that higher temperate extinction represents a dominant force for the origin and maintenance of LDG. More recently, Pulido-Santacruz & Weir^49^ proposed the terrestrial LDG is largely the effect of a post-Eocene increase in extinction rates at high latitudes resulting from the cooling Cenozoic trend.

Unfortunately, using phylogenies alone, studies on the LDG have not clearly demonstrated diversity losses in the Holarctic but instead high regional turnover (*λ*≈*µ*)^30, 32, 34–37, 50^. Nonetheless, high turnover can only explain a slow accumulation of lineages, with one fauna being replaced by another, but does not explain diversity decline (*i.e.* a reduction in the net number of species). Diversity declines occur when extinction exceeds speciation, resulting in negative net diversification rates (*r= λ −µ*; *r<0*). Accordingly, ‘diversity loss’ hypotheses differ from ‘high turnover’ hypotheses.

The perceived difficulty for inferring negative diversification rates from present-day phylogenetic data^51, 52^ and the assumption that diversity levels were always lower in the Holarctic than at the equator have resulted in ‘diversity loss’ hypotheses being repeatedly proposed but seldom demonstrated. Meanwhile, numerous fossil investigations have detected signatures of extinction and diversity loss in the Northern Hemisphere. For instance, Archibald *et al.*^15, 16^ sampled insect diversity at an Eocene site in Canada, and in present-day temperate Massachusetts (USA) and tropical sites of Costa Rica. Insect diversity was higher at the Eocene paleotropical site than the modern temperate locality, and comparable to the modern-day tropical locality, suggesting that post-Eocene insects have thus suffered greater levels of extinction in the Nearctic regions than around the equator. This pattern is consistent with other studies on various taxonomic groups, including birds^53^, invertebrates^15, 16, 54^, mammals^12, 13, 55^ and plants^56–58^. However, fossil studies are generally restricted to a geographic and temporal scale, which makes difficult to extrapolate local inferences of extinction in the context of the LDG.

Here, we use comparative methods for both phylogenies and fossils to estimate the evolutionary processes behind the LDG of Testudines, Crocodilia and Lepidosauria, and test the hypothesis of Holarctic diversity loss (**Fig. 1**). To this end, we propose to include a temporal component to study the LDG in which prevailing speciation, extinction and dispersal dynamics may change between warm- and cold-time intervals. The modern-day Crocodilia and Lepidosauria comprise mostly tropical-adapted species with a classic LDG pattern as shown by diversity peaks at equatorial latitudes^39, 44^. We investigate the origin of the LDG in subtropical taxa as well, by extending the study to Testudines, which display a hump-shaped gradient of diversity centred on subtropical latitudes (10°S–30°N)^59^. By contrast, the paleolatitudinal distribution of turtles was concentrated in the Holarctic (30–60°N) during the Cretaceous^24, 25^. All these lineages are ancient and likely experienced climatic transitions during the early Cenozoic^23, 25, 39, 44, 59^. They display contrasting patterns of species richness: turtles and crocodiles are species-poor (330 and 25 species, respectively), while lepidosaurs include a large number of species (9500+ species), and all have a rich fossil record extending back to the Triassic (Early Cretaceous for crocodiles), providing information about the variation of latitudinal species richness accumulation during their evolutionary history.

## Results

### Phylogeny-based diversification analyses: are diversification rates higher at the equator?

According to current distribution data, the species richness of turtles, lepidosaurs and crocodiles peaks near the equator, with 84% of all extant species living in the tropics, only 15% living in temperate regions and 1% spanning both biomes. We classified each species reported in the phylogeny (Supplementary Tables 1-3) as living close to the equator (the modern-day tropical biome) or the Holarctic and Southern Hemisphere (the modern-day temperate biome). For turtles, there were 239 tropical species, 84 temperate and 6 spanning both biomes (7 were marine species). For lepidosaurs, there were 7955 tropical species, 1337 temperate and 124 spanning both biomes. The species-poor crocodile clade had only 23 tropical and two temperate species.

We analysed differences in diversification rates between the Holarctic and equatorial regions, with the binary-state speciation and extinction model (BiSSE^60, 61^, see *Methods*). We did not use the geographic-state speciation and extinction model^62^, which is appropriate for dealing with widespread species, because most of the species in our datasets were endemic to the Holarctic or equatorial regions, and, for a character state to be considered in SSE models, it must account for at least 10% of the total diversity^63^. We did not apply the BiSSE model to crocodiles, because simulations have shown that trees containing fewer than 300 species may have to weak a phylogenetic signal to generate sufficient statistical power^63^.

We first used the time-constant BiSSE model, which is generally used in studies of the LDG. For turtles, net diversification rates were higher in the Holarctic than at the equator (**Table 2**, Supplementary Fig. 1a), but this difference was not significant, and rates of dispersal ‘*into the equator’* were ten times higher than those ‘*out of the equator’*. For lepidosaurs, a similar dispersal pattern was recovered, but net diversification rates were significantly higher in the equator (Supplementary Fig. 1b). We additionally introduced two shift times, at 51 and 23 million years ago (Ma), to detect differences in diversification dynamics between greenhouse, transitional, and coldhouse periods. This model indicated that the net diversification of turtles was similar in the Holarctic and at the equator, whereas it was lower in the Holarctic for lepidosaurs until the coldhouse period, when Holarctic diversification increased (**Table 2**, Supplementary Fig. 2). Dispersal was considered to be symmetric between regions (*into the equator = out of the equator*) during greenhouse periods, and asymmetric (*into the equator > out of the equator*) during the climatic transition and coldhouse period. The same patterns were obtained for analyses with the same model but with different combinations of shift times (51/66 Ma and 23/34 Ma; Supplementary Fig. 3).

**Table 2.** Results of the diversification and biogeographic analyses performed in this study for the transition from greenhouse to coldhouse climates. Abbreviations: *λ* = speciation, *µ* = extinction, *d* = dispersal, *Rμ* = range extirpations, e = equatorial, h = Holarctic.

|  | Data source | Model | Turtles | Squamates | Crocodiles |
| --- | --- | --- | --- | --- | --- |
|  |  |  | Green- to coldhouse transition | Green- to coldhouse transition | Green- to coldhouse transition |
| Diversification analyses | Fossil | PyRate | $\lambda_e > \lambda_h$<br>$\mu_e < \mu_h$<br>$r_e > r_h$ | $\lambda_e = \lambda_h$<br>$\mu_e > \mu_h$<br>$r_e < r_h$ | $\lambda_e > \lambda_h$<br>$\mu_e < \mu_h$<br>$r_e > r_h$ |
| | Present | BiSSE (time-variable) | $\lambda_e = \lambda_h$<br>$\mu_e = \mu_h$<br>$d_{he} > d_{eh}$ | $\lambda_e > \lambda_h$<br>$\mu_e < \mu_h$<br>$d_{he} > d_{eh}$ | - |
| | Present | BiSSE (constant) | $\lambda_e < \lambda_h$<br>$\mu_e = \mu_h$<br>$d_{he} > d_{eh}$ | $\lambda_e < \lambda_h$<br>$\mu_e < \mu_h$<br>$d_{he} > d_{eh}$ | - |
| Biogeographic analyses | Present | DEC | $R\mu_e < R\mu_h$ ($R\mu_e < R\mu_h$)*<br>$d_{he} < d_{eh}$ ($d_{he} < d_{eh}$)* | $R\mu_e > R\mu_h$ ($R\mu_e = R\mu_h$)*<br>$d_{he} < d_{eh}$ ($d_{he} = d_{eh}$)* | $R\mu_e = R\mu_h$ ($R\mu_e = R\mu_h$)*<br>$d_{he} = d_{eh}$ ($d_{he} = d_{eh}$)* |
| | Present + fossil | DEC fossil | $R\mu_e < R\mu_h$ ($R\mu_e < R\mu_h$)*<br>$d_{he} > d_{eh}$ ($d_{he} > d_{eh}$)* | $R\mu_e > R\mu_h$ ($R\mu_e < R\mu_h$)*<br>$d_{he} < d_{eh}$ ($d_{he} > d_{eh}$)* | $R\mu_e < R\mu_h$ ($R\mu_e < R\mu_h$)*<br>$d_{he} > d_{eh}$ ($d_{he} > d_{eh}$)* |

\* Prevalent dynamics when the number of events is calculated relative to the number of taxa currently distributed in each region (see methods)

### Fossil-based diversification analyses: evidence for ancient tropical extinctions?

We also analyzed differences in diversification rates between the Holarctic and equatorial regions based exclusively on fossil data. The turtle fossil dataset comprised 4084 occurrences for 420 genera (65 extant and 355 extinct; Supplementary Table 4). The lepidosaur fossil dataset comprised 4798 occurrences for 638 genera (120 extant and 518 extinct; Supplementary Table 5). The crocodile fossil dataset comprised 1596 occurrences for 121 genera (9 extant and 112 extinct; Supplementary Table 6). We first inferred global diversification dynamics by analyzing the fossil datasets as a whole with a Bayesian approach to inferring the temporal dynamics of origination and extinction rates based on fossil occurrences (PyRate^64^, see *Methods*). For turtles, origination rates peaked during the Jurassic, subsequently decreasing until the present day. Extinction rates were generally low and constant during the Mesozoic, but increased during the coldhouse periods of the Jurassic and Paleogene, resulting in negative net diversification during the Paleogene (**Fig. 2, Table 2**, Supplementary Figs. 4, 5).

**Figure 2.**
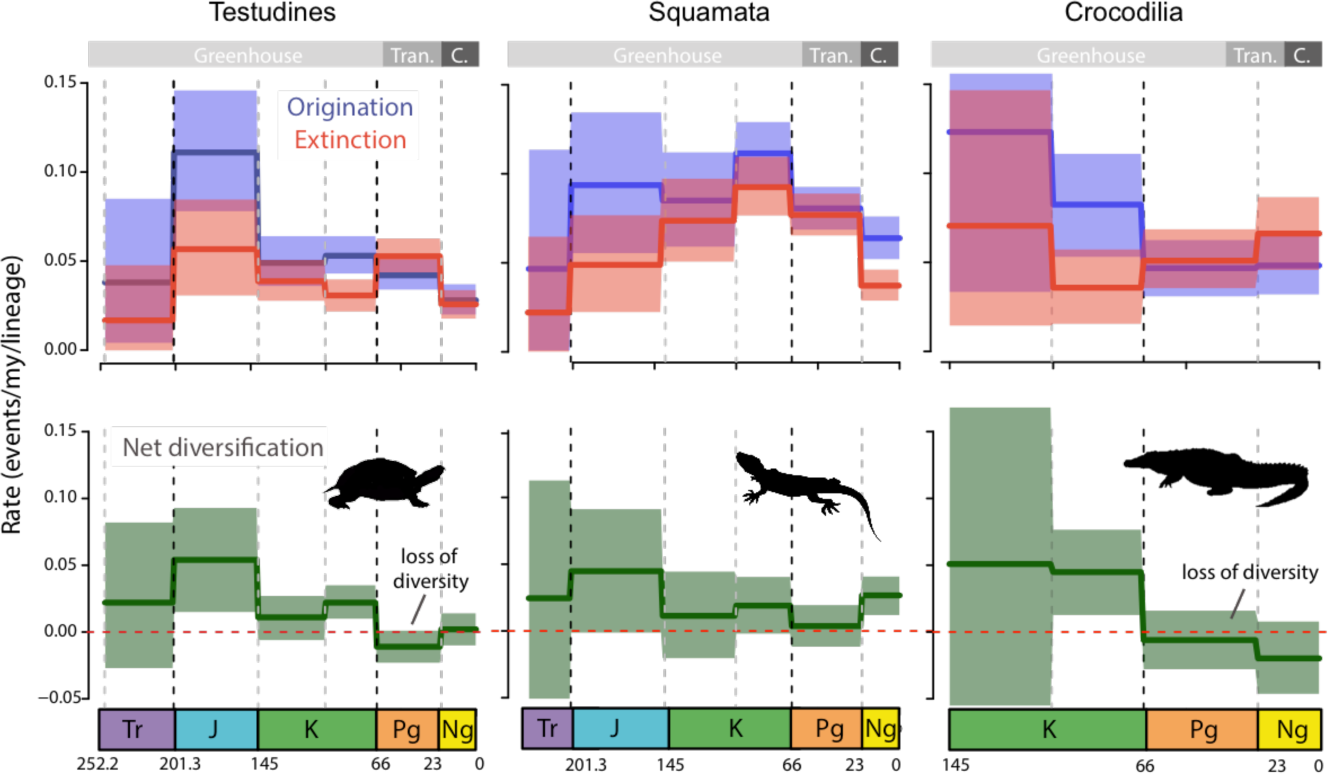
Global pattern of turtles, squamates and crocodiles diversification through time based on the fossil record, and analysed with a Bayesian model. Origination (blue) and extinction (red) rates were estimated using time bins as defined by epochs of the geological timescale (on the top, main climatic periods are shown as follows: Greenhouse, Tran. = climatic transition, and C. = coldhouse). Solid lines indicate mean posterior rates, whereas the shaded areas show 95% credibility intervals. Net diversification rates (green) are the difference between origination and extinction. The vertical lines indicate the boundaries between geological periods. Tr=Triassic; J=Jurassic; K=Cretaceous; Pg=Paleogene, and Ng=Neogene.

For lepidosaurs, origination rates peaked in the Jurassic and Late Cretaceous, whereas extinction increased steadily until the Late Cretaceous. In the Paleogene, net diversification approached zero, suggesting a high turnover (**Fig. 2**, Supplementary Figs. 6, 7). Crocodile origination peaked in the Early Cretaceous, subsequently decreasing toward the present day, and extinction rates were generally low and constant. We also identified diversity losses in the Paleogene extending to the present, suggesting that crocodiles are still in a phase of declining diversity (**Fig. 2**, Supplementary Figs. 8, 9).

We performed additional analyses with different subsets of the three fossil datasets, to separate the origination and extinction signals between geographic regions (equator or Holarctic) and ecological conditions (temperate or tropical, see *Methods*). These analyses showed that the diversity losses experienced by turtles and crocodiles during the Paleogene were mostly attributable to species living in the Holarctic and under tropical conditions (**Figs. 3, 4, Table 2**). The global diversity loss inferred for crocodiles during the Neogene was attributed to taxa living in both the Holarctic and equatorial regions (adapted to temperate and tropical conditions respectively), providing further support for the hypothesis that this whole group is in decline. For all groups, temperate taxa have been estimated to have high rates of diversification during the Oligocene, but lower rates during the Neogene. For the equatorial datasets, extinction and origination rates decreased over time, resulting in constant net diversification rates (except for lepidosaurs, which displayed a decrease in diversification during the Paleogene, followed by an increase during the Neogene).

**Figure 3.**
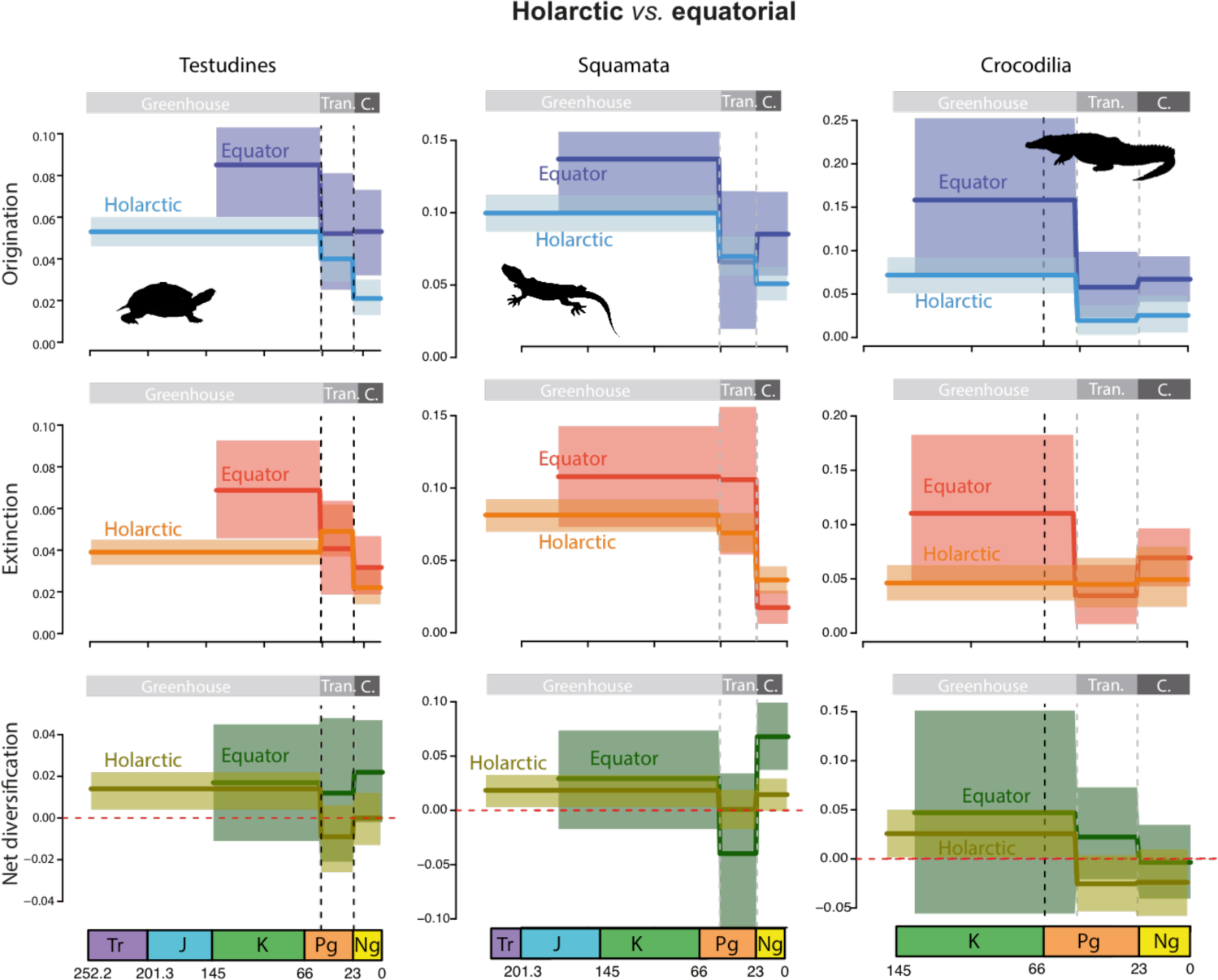
Global pattern of turtle, squamate and crocodile diversification between Holarctic and equatorial regions, based on the fossil record. Diversification dynamics are compared between fossils distributed in Holarctic and equatorial regions. Origination (blue) and extinction (red) rates were estimated using time bins as defined by the main climatic intervals since the Mesozoic (on the top, climatic periods are shown as follows: Greenhouse, Tran. = climatic transition, and C. = coldhouse). Solid lines indicate mean posterior rates, whereas the shaded areas show 95% credibility intervals. Net diversification rates (green) are the difference between origination and extinction. The vertical lines indicate the boundaries between climatic intervals. Tr=Triassic; J=Jurassic; K=Cretaceous; Pg=Paleogene, and Ng=Neogene.

**Figure 4.**
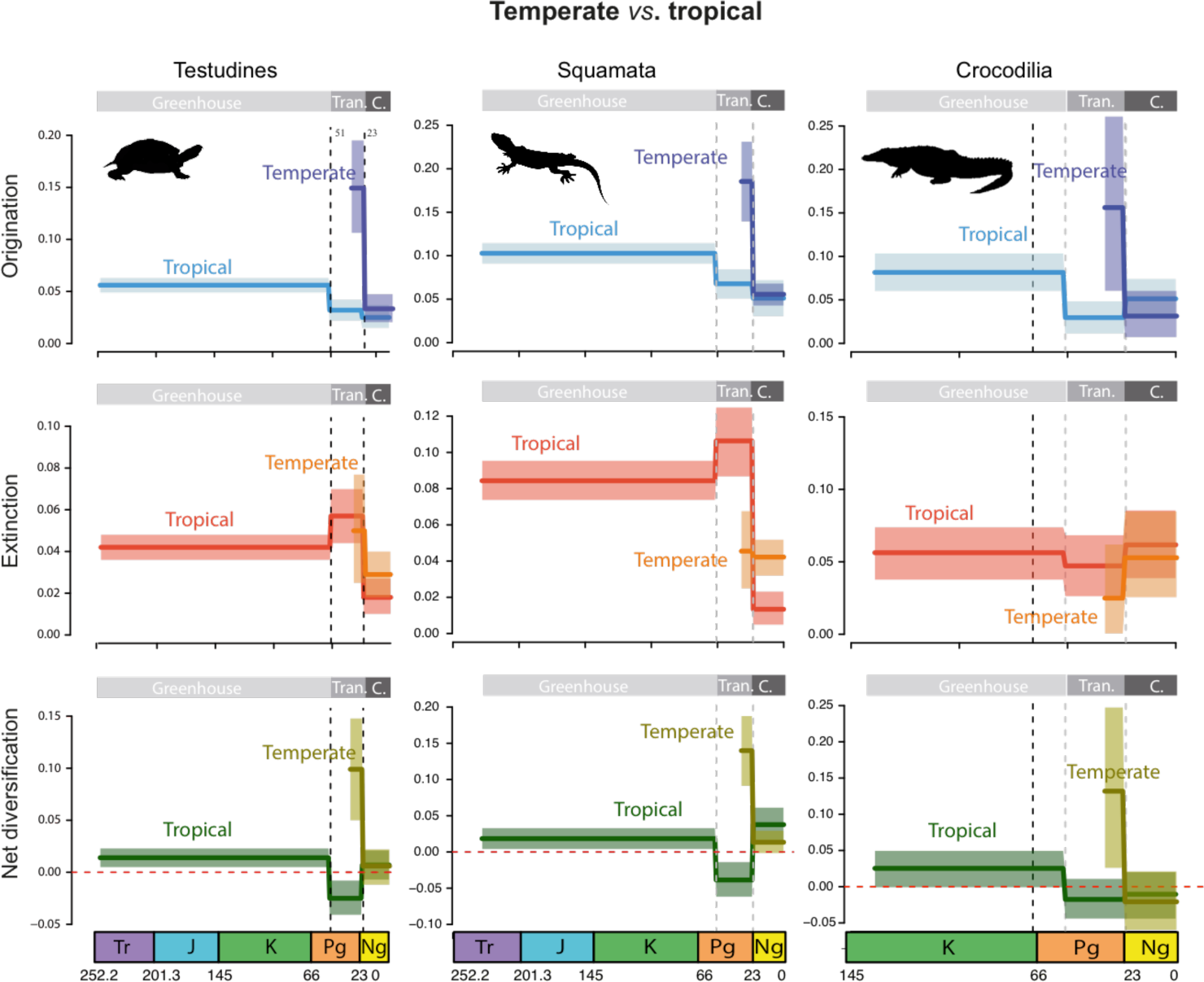
Global pattern of turtle, squamate and crocodile diversification across temperate and tropical climates, based on the fossil record. Diversification dynamics are compared between fossils inhabiting under temperate and tropical macroclimates. Origination (blue) and extinction (red) rates were estimated using time bins as defined by the main climatic intervals since the Mesozoic (on the top, climatic periods are shown as follows: Greenhouse, Trans. = climatic transition, and C. = coldhouse). Solid lines indicate mean posterior rates, whereas shaded areas show 95% credibility intervals. Net diversification rates (green) are the difference between origination and extinction. Vertical lines show boundaries between climatic intervals. Tr=Triassic; J=Jurassic; K=Cretaceous; Pg=Paleogene, Ng=Neogene.

### Estimations of ancestral origins: did groups preferentially originate close to the equator?

We performed biogeographic analyses with the dispersal-extinction-cladogenesis (DEC) model^65^ and dated phylogenies (see *Methods*). We first analyzed the data in an unconstrained DEC analysis in which all ranges covering three areas could be in an ancestral state. We inferred an equatorial distribution for the deepest nodes for the turtles and lepidosaurs, whence these lineages colonized the other regions (**Fig. 5a, Table 2**, Supplementary Fig. 10). Crocodile ancestors were found to have been widespread during the Cretaceous, with an early vicariant speciation event separating *Alligator* in the Holarctic from the other genera of Alligatoridae in equatorial regions (Supplementary Fig. 11). Our biogeographic estimates based exclusively on extant data conflict with the fossil record^23, 24, 66^. We overcame this bias by introducing information about the distribution of fossils into DEC, in the form of hard (HFC) and soft (SFC) geographic fossil constraints at specific nodes (see *Methods*; Supplementary Tables 7–9). The inclusion of fossil information yielded very different biogeographic histories for the three groups (**Table 2**; turtles: **Fig. 5b**, Supplementary Fig. 12; lepidosaurs: Supplementary Figs. 13, 14; and crocodiles: Supplementary Figs. 15, 16). Under the SFC model, turtles were found to have originated in the Northern Hemisphere (under the HFC model they were spread over both regions), whence lineages migrated toward the equator and southern regions (**Fig. 5b**, Supplementary Fig. 12). Most dispersal therefore occurred ‘*into the equator*’ (Supplementary Fig. 17, Supplementary Table 10). We also detected a larger number of geographic extinctions when fossil ranges were considered, predominantly for turtle lineages in the Holarctic (53 and 11 lineages disappeared from this region under the HFC and SFC models, respectively) and in southern temperate regions (9 in the HFC model; Supplementary Fig. 17, Supplementary Table 11). The same trend was observed when the number of extinction/dispersal events was controlled for the number of lineages currently distributed in each region (**Fig. 6**). The uncertainty associated to this estimation does not affect the overall result (Supplementary Table 12 and Appendix 1).

**Figure 5.**
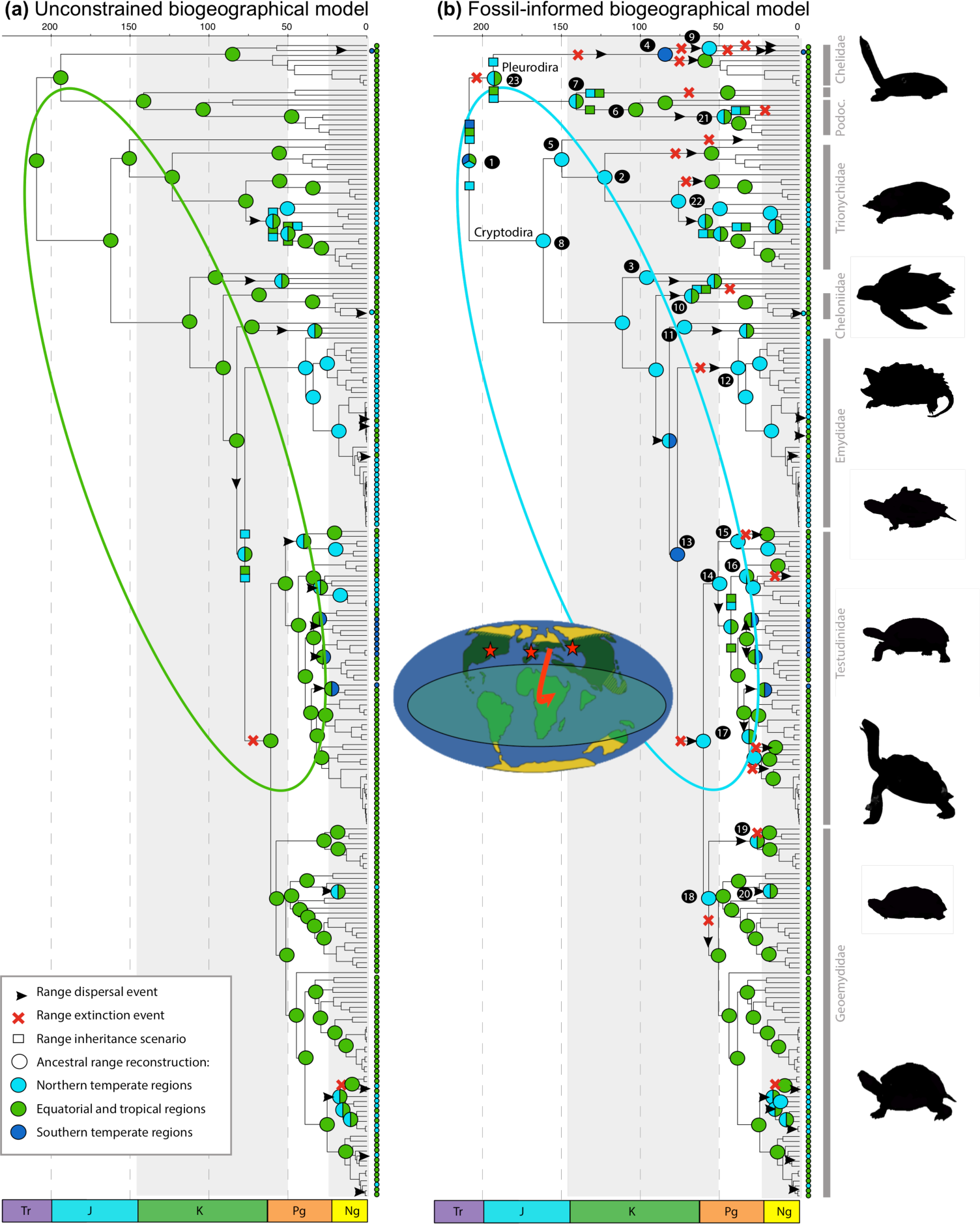
Biogeographic reconstruction of Testudines showing the effects of the incorporation of fossil information into biogeographic inference. **a**, Biogeographic reconstruction inferred with DEC based on the distribution of extant taxa. **b**, Biogeographic reconstruction under the fossil-informed HFC (hard fossil constraint) model. Coloured circles at tips and nodes represent current and ancestral ranges, respectively, while squares represent range inheritance scenarios. Colours correspond with the discrete areas in the legend. Black circles indicate fossil range constraints included in the analysis, with numbers corresponding with taxa in Table Supplementary S7. The reconstruction under the soft fossil constraint (SFC, see text) model is presented in Supplementary Fig. 12. Tr=Triassic; J=Jurassic; K=Cretaceous; Pg=Paleogene, and Ng=Neogene.

**Figure 6.**
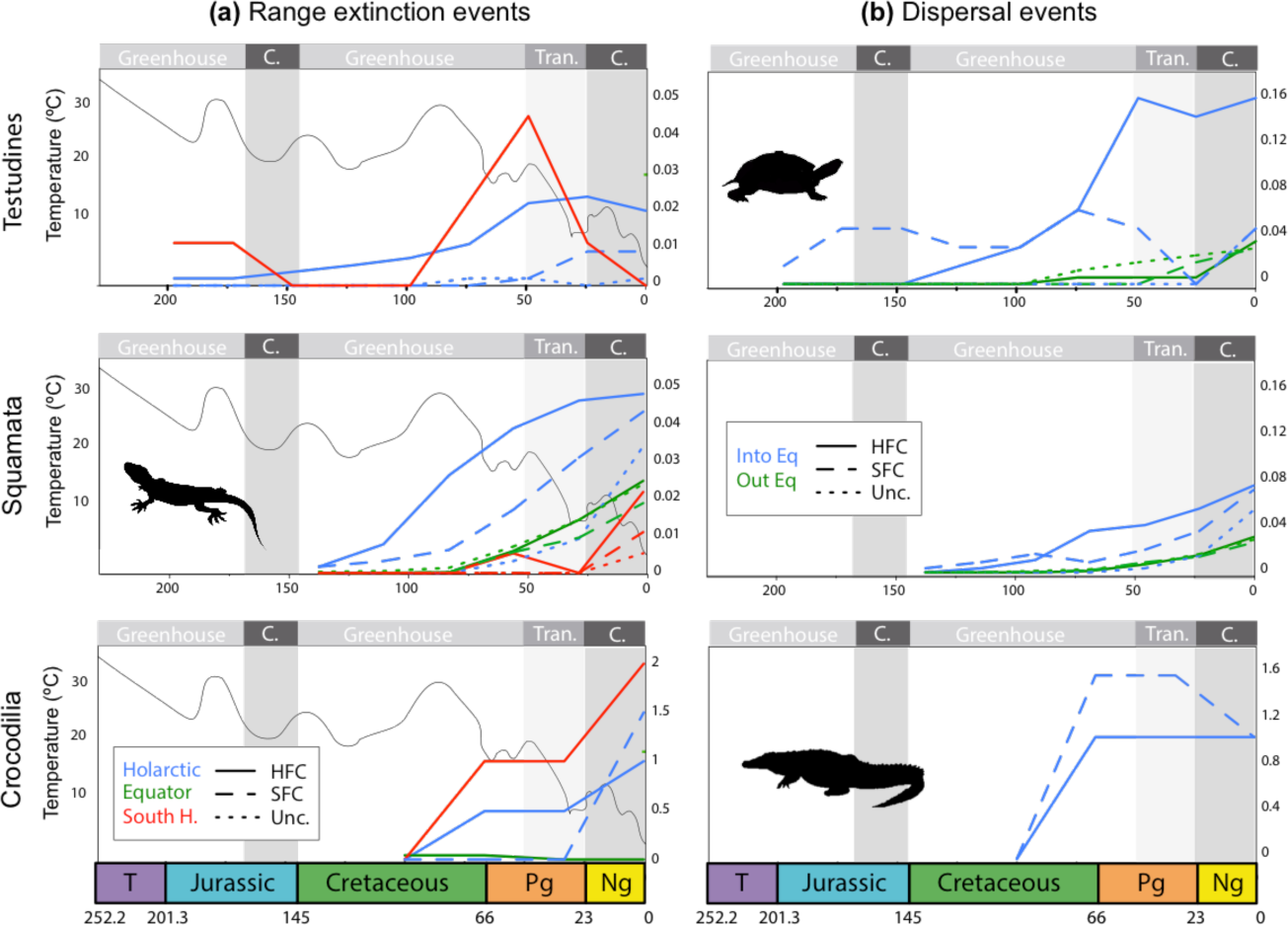
Estimated number of range-extinction and dispersal events through time. Analyses were performed for Testudines, Squamata and Crocodiles under the unconstrained model (Unc.), based on present evidence only, and the fossil-based hard (HFC) and soft fossil constraint (SFC) biogeographic models. **a**, Inferred number of range extinction events through time and across regions relative to the number of lineages currently distributed in each region. The black line represents the global mean temperature curve as modified from^73^. **b**, Inferred number of dispersal events from the Holarctic into the equator (IntoEq) and out of the equatorial zone (OutEq), relative to the current number of lineages distributed in the Holarctic and equatorial zones, respectively. Tr, Triassic; J, Jurassic; K, Cretaceous; Pg, Paleogene; and Ng, Neogene, Trans. = climatic transition, and C. = coldhouse.

The most supported biogeographic scenarios in both SFC and HFC analyses also suggest that lepidosaur ancestors were widespread (Supplementary Figs. 13, 14; uncertainty presented in Supplementary Tables 13 and Appendix 1). During the greenhouse period, dispersal ‘*into the equator*’ occurred at the same rate (or at a higher rate in the HFC model) than dispersal ‘*out of the equator*’, and dispersal ‘*out of the equator*’ prevailed thereafter (Supplementary Fig. 17, Supplementary Table 10). Estimated range extinction rates were high in this group under the unconstrained model, with 30 lineages extirpated from the Holarctic, two from southern temperate regions and 152 from the equator (Supplementary Fig. 17, Supplementary Table 11).

Under fossil-informed models, the number of Holarctic extinctions was higher (109 and 66 lineages in the HFC and SFC models, respectively), whereas the number of lineages extirpated from the equator was similar (144 and 109 in the HFC and SFC models, respectively; Supplementary Fig. 17). When the number of events was controlled for the actual number of lineages distributed in each region, the number of Holarctic extinctions and dispersals ‘*into the equator*’ increased dramatically, exceeding equatorial dispersal/extinctions (**Fig. 6**). For crocodiles, analyses including fossil ranges showed that all basal nodes were distributed in the Holarctic (Supplementary Figs. 15, 16), and range extinctions were detected: four lineages disappeared from the Holarctic, three from southern temperate regions, and two from the equator (HFC model; Supplementary Fig. 17, Supplementary Tables 11, 15). Only two lineages disappeared from the Holarctic in the SFC model. The same trends were observed after controlling the number of events for the current number of lineages in each region (**Fig. 6**). The uncertainty associated to this estimation does not affect the overall result (Supplementary Table 14 and Appendix 1).

## Discussion

### The role of time-varying evolutionary processes in the generation of latitudinal diversity gradients

Fossil investigations have shown that, at certain times during the Phanerozoic, the LDG has weakened, flattened, or developed a paleotemperate peak, with diversity at high latitudes being greater than currently for many groups^10, 13^. Hypotheses relating to ‘slow Holarctic diversity accumulation’, such as limited dispersal to the Holarctic^29^, high Holarctic turnover^35, 39, 49^, or high rates of equatorial diversification^30–33, 67^, cannot themselves account for the formation of a flatten LDG, or the transition from higher to lower diversity in the Holarctic observed in many groups. Furthermore, although the processes shaping biodiversity vary over time and space, this has been largely overlooked in the context of the LDG, which has been generally explained in terms of time-constant processes.

Over Earth’s history, the geographic extent of the tropical biome around the equator has fluctuated, with periods of pole-ward expansion during which warm paratropical conditions appeared at high latitudes, followed by periods of equator-ward contractions^68–72^ (**Fig. 1**). For instance, the last 100 million years have witnessed the contraction of tropical conditions toward the equator, due to the global cooling since the latest Cretaceous–early Cenozoic (the most recent greenhouse period), culminating in the Pleistocene glaciations^73^. To account for these environmental changes and their effect on the global distribution of biodiversity, we propose to include a temporal component to study the LDG. Doing this, our time-variable models identified gains and losses of tropical diversity at high latitudes, with prevailing speciation, extinction and dispersal dynamics changing between warm and cold intervals. In particular, fossil-based analyses show that diversification rates in the Holarctic and equatorial regions were similar during the greenhouse period of the Cretaceous-early Cenozoic for all groups studied here (overlapping credibility intervals; **Fig. 3**; Supplementary Figs. 4–9), consistent with the idea of the existence of a flattened LDG during this phase^10, 13^. Thereafter, diversification rates of turtles and crocodiles decreased in all regions in the transition to colder climates, but the slowing of diversification was much stronger in the Holarctic than at the equator, with extinction exceeding speciation in this region (*i.e.* Holarctic diversity loss; **Fig. 3**). In addition, by using phylogenetic-based biogeographic models informed by fossils, we inferred that all groups had a widespread ancestral distribution that subsequently contracted toward the equator during Cenozoic cooling. This result is in agreement with previous fossil investigations for turtles^24, 25, 66^ and crocodiles^23, 44^. Range contraction in our study results from higher levels of range extirpations (*i.e.* when a lineage losses part of its range) at higher latitudes and ‘*into the equator*’ dispersals^36, 50^ (**Figs. 5, 6**). All together, these results suggest that Holarctic diversity loss (*r<0*), Holarctic range extinctions, and ‘*into the equator*’ dispersals during the Paleogene impoverished the region and could also explain the origin of a steep LDG for these groups.

Results exclusively based on extant species (time-constant and time-variable BiSSE analyses) differed from the analyses including fossils, suggesting a different scenario: higher levels of Holarctic diversification for turtles during Cenozoic cooling. Biogeographic analyses based exclusively on extant species estimate an equatorial origin for turtles and recent invasion of high-latitude regions, resulting in less time for lineages to diversify (**Figs. 5, 6**), in agreement with the ‘*tropical niche conservatism*’ hypothesis^29^ and recent investigations^74, 75^. For crocodiles they support the diversification hypothesis, with higher origination rates close to the equator and no effect on dispersal (**Fig. 6, Table 2**, Supplementary Fig. 11).

Diversification patterns inferred based on fossils for lepidosaurs differ from turtles and crocodiles. Although we found similar diversification rates in the Holarctic and equator during the greenhouse period for this group and widespread ancestral distributions, diversity losses occurred in the equator and a high turnover at higher latitudes during cooling climate (**Fig. 3**; Supplementary Figs 13, 14). Diversity dynamics for species distributed at the equator, however, may not be entirely reliable, due to the poverty of the equatorial dataset in terms of the number of fossil lineages and the small number of records per lineage (Supplementary Table 15). Uncertainties therefore remain on these estimates, which have large credibility intervals probably due to geographic and/or preservation biases in the fossil record^76^. Turnover rates were high in the Holarctic during the transitional period to cold, indicating that species did disappear from high latitudes, but that a lepidosaur community got replaced by another. This result suggests the number of lepidosaur species may always have been unbalanced between regions; the high Holarctic turnover thus contributing to the maintenance of this pattern, together with the inferred temporal increases in diversification at the equator (**Fig. 3**), as previously hypothesized^39^. In addition, in absolute terms, we found that more lepidosaurs species migrated ‘*out of’* than ‘*into the equator’* (Supplementary Fig. 17), but the number of species in the equatorial region today is four times the number of lineages elsewhere. After controlling for the imbalance in species sampling in our tree, we found that a higher proportion of lepidosaur species actually lost their ancestral Holarctic distribution and emigrated ‘*into the equator*’^39^ than the other way around (**Fig. 6**). Although the number of fossil constraints in the biogeographic analyses of lepidosaurs was relatively low given the size of the tree and in comparison with the other groups (30 Holarctic and equatorial fossils for 4161 nodes), these constraints significantly increased the absolute number of Holarctic range extinctions (from 30 to 109) and ‘*into the equator*’ dispersals (from 40 to 124) relative to estimates without such constraints (Supplementary Tables 10, 11). Meanwhile, the inclusion of fossil data did not alter the number of events estimated for equatorial taxa. This finding suggests that a deeper understanding of lepidosaur fossil taxonomy might facilitate the assignment of fossils on the tree, and the detection of additional high-latitude range extinctions not detected here.

In summary, the general pattern is that ‘Holarctic diversity loss’ scenarios were supported for crocodiles and turtles using fossil and fossil-informed phylogenetic investigations (**Table 2**), whether if we rely only on analyses based on extant species this evolutionary scenario was poorly supported. Support for diversity loss is mixed in lepidosaurs: on the one hand the detected Holarctic range contractions are in agreement with this hypothesis, and on the other hand the evidence for high Holarctic turnover is more in line with previous ‘slower Holarctic diversity accumulation’ hypotheses.

### The role of climate dynamics in shaping latitudinal diversity gradients

Recent fossil investigations suggest that changes of the LDG shape have been associated with major climatic oscillations^10, 13^. In agreement with this idea, we detect an impoverishment of the Holarctic during the last greenhouse to coldhouse transition defined here between 51 and 23 Ma, after the early Eocene Climatic Optimum (EECO; see *Methods*). In addition, the ancestors of turtles, lepidosaurs and crocodiles were adapted to tropical conditions during the Late Cretaceous^44, 77, 78^. Our diversification results based on fossils when the dataset is divided according to their ecological preferences further indicate that extinction events were not random, instead preferentially affecting taxa living in tropical-like climates at high latitudes^48^ (**Fig. 4**). In agreement with the niche conservatism hypothesis^29, 41, 79^, in which macroclimatic conditions play a major role in the structuring of global diversity^80^, our results suggest that many species adapted to warm conditions living in the Holarctic were unable to adapt to the new temperate regimes of the Neogene, and either went extinct or escaped extinction by contracting their ranges in a southerly direction (**Fig. 6**).

It could be hypothesized that the expansion of tropical conditions to higher latitudes during greenhouse periods might have induced species diversification in the new paratropical areas and facilitated movements within the broad ‘tropical belt’, such that tropical equatorial clades were able to disperse ‘*out of the equator’* and diversify into high-latitude warm regions^40, 81^. An equable Cretaceous-early Cenozoic greenhouse climate thus triggered the formation of a flat LDG (**Fig. 7a**). By contrast, the contraction of the tropical biome following climate cooling induced periods of declining diversity at high latitudes (where climate change was more intensively felt; **Fig. 3**), and mediated biotic movements ‘*into the equator’* (**Figs. 6, 7b**). Extinction rates were high for tropical-adapted lineages at high latitudes, but lower for low-latitude tropical lineages (**Fig. 4**). Climate change would thus have driven the development of an asymmetric gradient of extinction and dispersal (AGED) within the tropical biome, and could have mediated the formation of a steep LDG (**Fig. 7c**).

**Figure 7.**
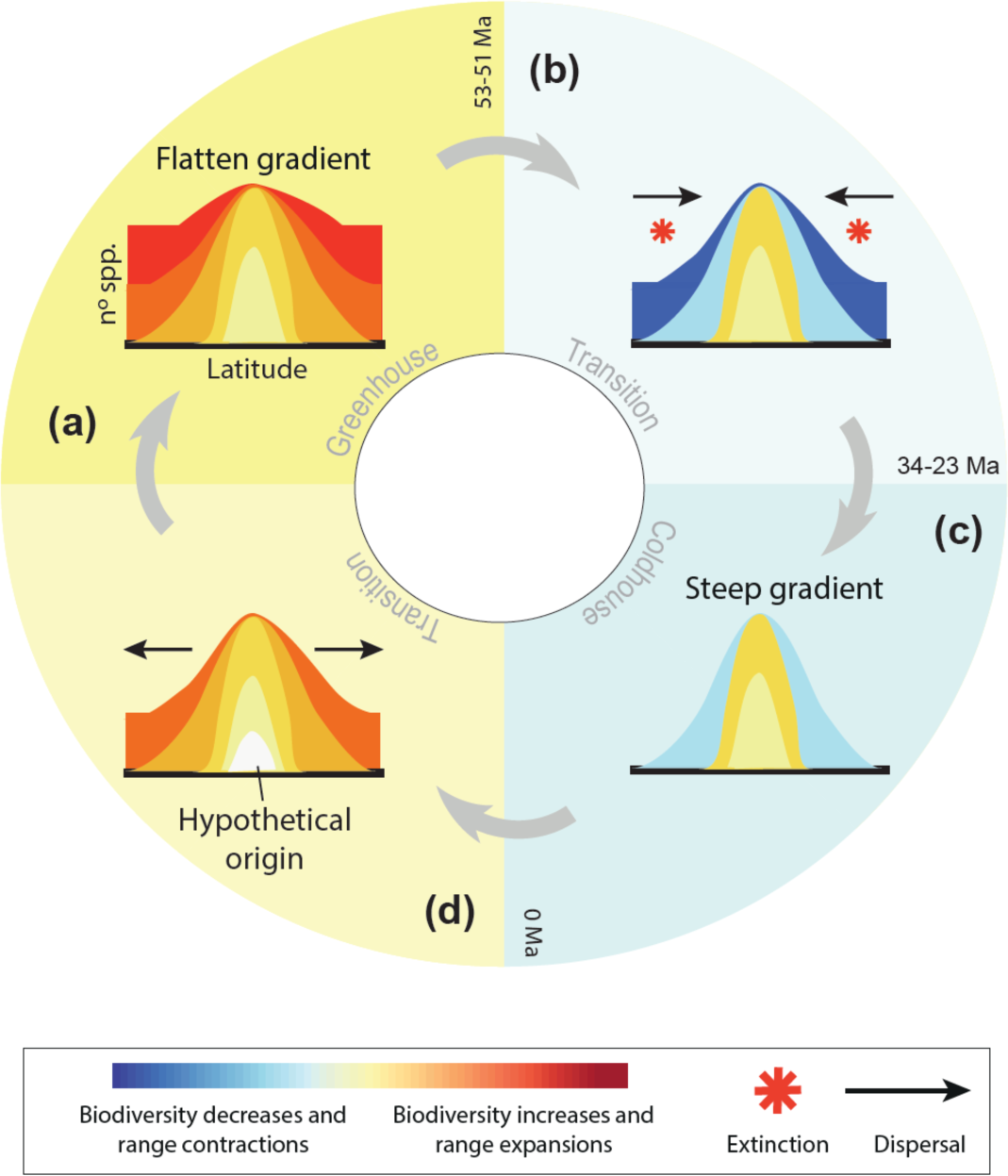
Prevalent evolutionary processes behind the latitudinal diversity gradient under the AGED time-variable framework. The graphic shows the hypothetic change in evolutionary dynamics between Holarctic and equatorial regions and across main climatic intervals: a, the greenhouse Late Cretaceous-early Cenozoic period. b, the late Eocene-Oligocene climatic transition. c, the current Neogene coldhouse interval. d, the past and future transition to greenhouse climates. For each period, inset Figs. represent the distribution of species richness across latitudes (the LDG shape) and the hypothetical change in global evolutionary dynamics under the AGED hypothesis.

We propose the AGED hypothesis to reconcile previous contending ideas on the origin of the LDG by placing them in a temporal scenario (**Table 1, Fig. 7**). For instance, there is controversial support around the tropics being ‘*cradle*’ or ‘*museum of diversity*’^82^, and dispersal prevailing ‘*out of*’^40, 81^ or ‘*into the tropics*’^36, 39, 50^. The interpretation of our results alternatively invokes the ‘*museum of diversity*’ regarding the equatorial tropics as refuge during coldhouse transitions, but also the ‘*cradle of diversity*’ during greenhouse periods. Similarly, our hypothesis invokes ‘*out of the equator*’ dispersals during greenhouse transitions and ‘*into the equator*’ dispersals during coldhouse transitions. Hence, the AGED hypothesis is not entirely a new idea, but mostly the contextualization of previous ideas on the origin of the LDG in the frame of a time-variable scenario.

The AGED hypothesis might be applicable for groups adapted to tropical conditions. However, lineages possessing or having evolved the appropriate adaptations to cope with climate change^83, 84^ might have diversified in the new temperate areas. We actually found that the diversification rates of turtles, crocodiles and lepidosaurs living in temperate climatic conditions were significantly higher than those of tropical-adapted taxa living in Holarctic and equatorial regions after the transition to temperate climates in the late Eocene (**Fig. 4, Table 2**). The new temperate habitats could have constituted an opportunity for diversification because they increased geographic ranges and ecological niches^31^, and may have eventually driven an inverse LDG for some groups^37, 84^. Several radiations following the appearance of the temperate biome have been identified in other groups of organisms, such as plants^83, 85^, mammals^86, 87^ or insects^88^. After this period, speciation decreased dramatically in the temperate lineages of our focal groups, possibly due to the effect of the Pleistocene glaciations, and we estimated similar diversification rates between tropical and temperate lineages (**Fig. 4**). In summary, our study suggests that differences in species richness between geographic regions (*i.e.* the Holarctic vs. the equator) may be explained by differences in diversification and dispersal rates. Differences in species richness between ecological types (*i.e.* tropical- vs. temperate-adapted taxa) may be explained by the longer time available for tropical-adapted clades to diversify in tropical areas^89^ rather than higher rate of speciation under warm tropical environments^90^.

Nonetheless, we cannot exclude that diversity losses at high latitudes occurred since the Cretaceous, as suggested by our fossil-based diversification analyses with time intervals defined by the main geological periods (Supplementary Figs. 4, 6, 8) and by our fossil-based biogeographic analyses suggesting the prevalence of ‘*into the equator’* dispersals since the Cretaceous (**Fig. 6**, Supplementary Table 11). These findings could suggest that other processes different to climate change mediated the extinction and range contraction of Holarctic lineages, or alternatively, that a transition phase to cold started before the relatively short interval considered here (between 51 and 23 Ma). We defined this climatic interval based on paleontological evidence showing that paratropical conditions and the associated warm-adapted taxa disappeared from high latitudes between the mid-late Eocene and the Neogene^68, 70, 71^. However, some studies consider the EECO only represented a transient temperature peak within an otherwise cooling trend that started in the Cretaceous^73, 91^. This trend was intensified by the Cretaceous-Paleogene (K-Pg) mass extinction, and the drop in temperatures caused by the impact-associated winter^92, 93^. In our study, lineage extinctions, range extinctions and southward dispersals increased between the K-Pg and Neogene (**Figs. 3, 6**), suggesting an additive effect of K-Pg and Neogene cooling on depopulation of the Holarctic.

### Reconciling fossil and phylogenetic evidence

Our results unequivocally demonstrate that the inclusion of fossils in macroevolutionary studies greatly improves our ability to detect signals of ancient high-latitude extinctions and range extirpations (**Figs. 3–6**), otherwise hardly detectable with analyses based exclusively on extant species. This conflict between extant and fossil evidence may extend beyond our study, pervading the LDG literature. High extinction rates have occasionally been inferred in tropical lineages^83, 94–96^, with hypotheses related to extinction being focused on temperate groups and recent time scales, such as the effects of recent Pleistocene glaciations^30, 35, 39, 42^. In cases where extinction is inferred, speciation rates were also elevated in high-latitude groups (high turnover)^30, 35–37, 39, 50^, while diversity losses (*r<0*) have to our knowledge never been inferred in phylogenetic studies of the LDG (with the exception of a recent study^49^ using time-constant BiSSE models, but see below). On the other hand, description of ancient tropical extinction at high latitudes is supported by fossil studies on various taxonomic groups^15, 16, 28, 53–56^.

The last decade has seen ample efforts to reconcile fossil and phylogenetic evidence. Birth-death diversification models have been developed to detect negative diversification rates from reconstructed phylogenies or total-evidence trees^97–99^. Still, their use in the literature is limited and these models are difficult to implement in a trait evolution context, such as in the study of the LDG (but see^100^). LDG studies are often based on state-dependent speciation and extinction models^32, 36, 38, 39, 49, 50^ that are designed to test differential diversification and asymmetric transition scenarios, such as that suggested here, but LDG studies often assume that diversification parameters remain constant over time. If the evolutionary processes shaping the LDG have varied across latitudes and time, then time-constant models are not appropriate for testing more complex scenarios underlying the LDG. Moreover, the potential of time-constant models for detecting negative diversification rates is questionable, since inferring negative diversification for the entire history of lineages conflicts with the fact that these groups are still extant. Testing the hypothesis of diversity declines thus requires the implementation of time-variable models^100^. When applied to the study of diversity patterns, these models have revealed extinction signatures in ancestral tropical plant clades^101^. The incorporation of time-shifts into our BiSSE analyses improves but not completely reconciles the fossil evidence with extant diversity. Identifying the causes of this problem and finding solutions are beyond the scope of this study, but this artefact highlights the importance of fossils in macroevolutionary inferences^102^. Fossil records remain incomplete, but they nevertheless provide the only direct evidence of the diversity that existed in the past, and thus are fundamental in the study of extinction scenarios, as the one postulated here. By contrast to molecular phylogenies, the incompleteness of the fossil record has a less problematic effect on the estimation of speciation and extinction rates, because removing a random set of taxa does not affect the observed occurrences of other lineages^64^. Indeed, simulations have shown that PyRate correctly estimates the dynamics of speciation and extinction rates under low levels of preservation or severely incomplete taxon sampling

## Conclusion

The processes shaping the LDG remain among the most hotly debated topics in ecology and evolutionary biology. Our analyses indicated that these processes have changed over time. The current LDG of turtles and crocodilians can be, partially at least, explained by ancient high-latitude tropical diversity loss and range contractions as a consequence of the retraction of the tropical biome due to climate cooling. Meanwhile, equivalent diversification rates across latitudes during greenhouse periods could explain the formation of a flatten LDG. Overall, this suggests that changes in global diversification and dispersal dynamics imposed by large-scale climatic transitions could represent a mechanism that shapes the LDG. This ‘asymmetric gradient of extinction and dispersal’ hypothesis might account for the LDG of tropical-adapted groups that were once diverse at high latitudes, but might not be fully applicable to all organisms currently displaying a LDG, as shown here for lepidosaurs.

## Methods

### Time-calibrated phylogenies and the fossil record

. We study the LDG of three vertebrate groups: turtles (order Testudines), crocodiles (order Crocodilia), and scaled lizards (order Lepidosauria). A time-calibrated phylogeny for each group was obtained from published data. For turtles, we used the phylogeny of Jaffe et al.^103^, including 233 species. We preferred this phylogeny over other more recent and slightly better sampled trees^104^ because the divergence time estimates of Jaffe et al. are more consistent with recent estimates based on genomic datasets^74, 105^. For lepidosaurs, we retrieved the most comprehensive dated tree available, including 4161 species^39^, and a complete phylogeny was obtained for crocodiles^106^.

Fossil occurrences were downloaded from the Paleobiology Database (https://paleobiodb.org/#/, last accessed October 25^th^ 2017). We reduced potential biases in the taxonomic assignation of turtle, crocodile and lepidosaur fossils, by compiling occurrence data at the genus level. The fossil datasets were cleaned by checking for synonymies between taxa and for assignment to a particular genus or family on the basis of published results (Supplementary Table 4–6).

### Estimation of origination and extinction rates with phylogenies

We investigated possible differences between Holarctic and equatorial regions, by combining the turtle and lepidosaur phylogenies with distributional data (Supplementary Tables 1, 2) to fit trait-dependent diversification models in BiSSE^60^. We accounted for incomplete taxon sampling in the form of trait-specific global sampling fraction of extant species^107^.

We ensured comparability with previous LDG studies, by initially using a constant-rate trait-dependent diversification model. The constant-rate BiSSE model has six parameters: two speciation rates (without range shift, or in situ speciation), one associated with the Holarctic (hereafter ‘H’, λ_H_) and the other with other equatorial and subtropical regions (hereafter ‘equator’ or ‘E’, λ_E_), two extinction rates associated with the Holarctic (μ_H_) and the equator (μ_E_), and two transition rates (dispersal or range shift), one for the Holarctic to equator direction (q_H-E_), and the other for the equator to Holarctic direction (q_E-H_).

We then assessed the effect of species distribution on diversification, allowing for rate changes at specific time points. This approach is associated with a lower bias than the use of constant rates. We used the time-dependent BiSSE (BiSSE.td) model, in which speciation, extinction, and dispersal rates are allowed to vary between regions and to change after the shift times. We introduced two shift times to model different diversification dynamics between greenhouse, transitional, and coldhouse periods. We assumed that a global warm tropical-like climate dominated the world from the origin of the clades until 51 Ma (corresponding to the temperature peak in the Cenozoic). Thereafter, the climate progressively cooled until 23 Ma (the transitional period), when the climate definitively shifted to a temperate-like biome in the Holarctic^70, 71, 73^. The shift times at 51 Ma and at 23 Ma are initial values that are re-estimated by the model during the likelihood calculation. The climatic transition in the Cenozoic may have different temporal boundaries, with potential effects on the results. We thus applied the same model but with different combinations of shift times (we tested 51/66 Ma and 23/34 Ma for the upper and lower bounds of the climatic transition).

Analyses were performed with the R package *diversitree* 0.9-7^61^, using the *make.bisse* function to construct likelihood functions for each model from the data, and the functions constrain and *find.mle* to apply different diversification scenarios. Finally, we used a Markov Chain Monte Carlo (MCMC) approach to investigate the credibility intervals of the parameter estimates. Following previous recommendations^61^, we used an exponential prior 1/(2r) and initiated the chain with the parameters obtained by maximum likelihood methods. We ran 10,000 MCMC steps, with a burn-in of 10%.

### Estimation of origination and extinction rates with fossils

We also used fossil data to estimate diversification rates over time. We analysed the three fossil records, using a Bayesian model for simultaneous inference of the temporal dynamics of origination and extinction, and of preservation rates^64^. This approach, implemented in PyRate^108^, uses fossil occurrences that can be assigned to a taxon, in this case fossil genera. The preservation process is used to infer the individual origination and extinction times of each taxon from all fossil occurrences and an estimated preservation rate; it is expressed as expected occurrences per taxon per million years.

We followed a birth-death shift approach^109^, which focuses on the variation of origination and extinction at a global scale and over large temporal ranges. We used a homogeneous Poisson process of preservation (-mHPP option). We also accounted for the variation of preservation rates across taxa, using a Gamma model with gamma-distributed rate heterogeneity (-mG option). We used four rate categories to discretize the gamma distribution, to allow for a greater variability of preservation rates across taxa.

Given the large number of occurrences analysed and the vast timescale considered, we dissected the birth–death process into time intervals, and estimated origination and extinction rates within these intervals. In one set of analyses, we defined the time intervals using the geological epochs of the stratigraphic timescale^110^ (Supplementary Figs. 4, 6, 8). In another set of analyses, we defined the intervals according to the major climatic periods characterizing the Cenozoic (Supplementary Figs. 5, 7, 9): the greenhouse world (Cretaceous), the climatic transition (Paleogene), and the coldhouse world (Neogene until the present). We adopted this solution as an alternative to the algorithms implemented in the original PyRate software for joint estimation of the number of rate shifts and the times at which origination and extinction shift^64^. The estimation of origination and extinction rates within fixed time intervals improved the mixing of the MCMC and made it possible to obtain an overview of the general trends in rate variation over a long timescale ^109^. Both the preservation and birth–death processes were modelled in continuous time but without being based on boundary crossings. Thus, the origination and extinction rates were measured as the expected number of origination and extinction events per lineage per million years. One potential problem when fixing the number of rate shifts *a priori* is over-parameterization. We overcame this problem by assuming that the rates of origination and extinction belonged to two families of parameters following a common prior distribution, with parameters estimated from the data with hyper-priors^111^.

We ran PyRate for 10 million MCMC generations on each of the 10 randomly replicated datasets. We monitored chain mixing and effective sample sizes by examining the log files in Tracer 1.6^112^. After excluding the first 20% of the samples as a burn-in, we combined the posterior estimates of the origination and extinction rates across all replicates to generate plots of the change in rate over time. The rates of two adjacent intervals were considered significantly different if the mean of one lay outside the 95% credibility interval of the other, and vice versa. We looked at the marginal posterior distributions of origination and extinction rates through the evolutionary history of the three groups and assessed the effect of different environments.

In the context of the LDG, we performed additional analyses with different subsets of fossils, to separate the speciation and extinction signals of different geographic regions (equator or Holarctic) and ecological conditions (temperate or tropical). For example, for turtles, we split the global fossil dataset into four subsets: one for the fossil genera occurring at the equator (429 occurrences), one for the fossils occurring in the Holarctic (3568 occurrences), one for the fossil genera considered to be adapted to temperate conditions (993 occurrences), and one for the fossils considered to be adapted to tropical conditions (2996 occurrences). We excluded the few fossil occurrences for the southern regions of the South Hemisphere (about 180) only in subset analyses, as they were poorly represented in our dataset. Note that a given fossil can be present in both the ‘Holarctic’ and ‘tropical’ datasets. We encoded tropical/temperate preferences by considering macro-conditions in the Holarctic to be paratropical until the end of the Eocene, as previously reported ^70, 71^ (and references therein). We also assumed that taxa inhabiting the warm Holarctic were adapted to tropical-like conditions (i.e. a high global temperature, indicating probable adaptation to tropical climates). This is, of course, an oversimplification that may introduce bias into the analysis, but general patterns may nevertheless emerge from such analyses^80^. For turtles, crocodiles and lepidosaurs this assumption is supported by Cenozoic climatic niche modelling^77, 78^, stable isotope analyses and other climate proxies^44, 113^. After the late Eocene, we categorized each species as living in the temperate biome or the tropical biome, according to the threshold latitudes defining the tropics (23.4°N and 23.4°S) suggested in a previous study^30^. This delineation is also consistent overall with the Köppen climate classification. With these datasets, we reproduced the same PyRate analyses as for the whole dataset (see above). In general, the fossil datasets included mostly Holarctic fossils, with a smaller number of occurrences for the equator. Caution is therefore required when drawing conclusions from the equatorial datasets.

### Inferring ancestral geographic distribution with phylogenies and fossils

We performed biogeographic analyses with the parametric likelihood method DEC^65^ using the fast C++ version^114^ (https://github.com/rhr/lagrange-cpp). Turtle, lepidosaur, and crocodile species distributions were obtained from online databases (www.iucnredlist.org and www.reptile-database.org). We chose 23.4°N and 23.4°S as the threshold latitudes defining the tropics, and categorized each species as living in the Holarctic, in the southern temperate regions, or in the equatorial tropics and subtropical regions. We considered that all ranges comprising three areas could be considered an ancestral state (*maxareas* =3).

We set up three different DEC analyses. We first ran DEC with no particular constraints, using only the distribution of extant species. We then performed DEC analyses including fossil information in the form of ‘fossil constraints’ at certain nodes, according to the range of distribution of fossil occurrences assigned to a particular taxon during the relevant time frame. For example the crown age of Carettochelyidae (Testudines) dates back to the Late Jurassic (150 Ma, *node 5*, **Fig. 3**; Supplementary Table 7), and we set a constraint on this node reflecting the distribution of all the Late Jurassic fossils attributed to Carettochelyidae. Similarly, for the origin of turtles (210 Ma, *node 1*), distribution constraints represent the range of Late Triassic fossils assigned to turtles. For the crown of Trionychidae, in the Early Cretaceous (123 Ma, *node 2*), the early fossils assigned to the clade were used to constrain the geographic origin of Trionychidae. In total, we implemented 23 fossil constraints for turtles (Supplementary Table 7), 30 fossil constraints for lepidosaurs (Supplementary Table 8), and 8 for crocodiles (Supplementary Table 9).

We included the fossil distribution in two different approaches: *(i)* a soft (SFC), and *(ii)* hard fossil constraints (HFC). For the SFC approach, fossil data were incorporated into the anagenetic component of the likelihood framework. The direct impact of a given fossil is limited to the particular branch to which it has been assigned, although it may indirectly influence other branches. The inclusion of a fossil conditions the estimated geographic-transition probability matrix for that branch by imposing a spatiotemporal constraint on the simulation process. Only the simulations resulting in a geographic range including the area of fossil occurrence contribute to the geographic-range transition probability matrix for the branch concerned; simulations not meeting this constraint are discarded^115^. For SFC, we used the command ‘*fossil*’ in DEC. We consider this to be a ‘soft’ constraint, because other areas different from that in which the fossil was found could be included in the ancestral states. In some cases, in which today’s diversity is not representative of past diversity (*e.g.* due to extreme levels of extinction), the SFC model may still overlook known fossil information. We therefore also implemented an HFC model in which the estimation of ancestral areas was fixed to the location of fossils. This was achieved with existing functions in the C++ version of Lagrange, using the command ‘*fixnode*’. By fixing nodes to the distribution area of fossils, we assume fossil occurrences reflect the distribution of the ancestors, *i.e.* that the fossil record is complete. This is a strong assumption, but it makes it possible to recover all fossil ranges in the ancestral estimations. The real scenario probably lies somewhere between the SFC and HFC inferences.

We then compared the timing and number of range extinction and dispersal events inferred with the three different biogeographic analyses. In DEC, range-subdivision (inheritance) scenarios (vicariance, duplication and peripatric isolation) occur at cladogenetic events, whereas extinction (range contraction) and dispersal (range expansion) are modelled as stochastic processes occurring along the branches of the tree^116^. As the probability of any extinction/dispersal event is constant along the entire length of the branch, we estimate the periods at which range extinction and dispersal occurred by dividing the phylogeny into intervals of 25 million years and calculating the number of branches for which extinction/dispersal was inferred crossing a particular time interval (the same branch could cross two continuous intervals).

## Data accessibility

All the data used in this manuscript are presented in the manuscript and its supplementary material or have been published or archived elsewhere.

## Supplementary material

https://www.biorxiv.org/content/10.1101/236646v3.supplementary-material,

## Acknowledgements

This preprint has been reviewed and recommended by Peer Community In Evolutionary Biology (https://dx.doi.org/10.24072/pci.evolbiol.100068). The authors are very grateful to Joaquín Hortal, Juan Arroyo, Arne Mooers, Joaquín Calatayud, and two anonymous reviewers for comments and suggestions that greatly improved the study. Previous versions of the manuscript benefited from the comments of Gary Mittlebach, Emmanuelle Jousselin and Jonathan Rolland. Financial support was provided by a Marie-Curie FP7–COFUND (AgreenSkills fellowship–26719) grant to A.S.M. and a Marie Curie FP7-IOF (project 627684 BIOMME) grant to F.L.C. This work benefited from an “Investissements d’Avenir” grant managed by the “Agence Nationale de la Recherche” (CEBA, ref. ANR-10-LABX-25-01).

## Appendix 1

https://zenodo.org/record/2572685#.XGvv3idCe8U

