## Supplementary Fig. 1 for "Ancient tropical extinctions contributed to the latitudinal diversity gradient"

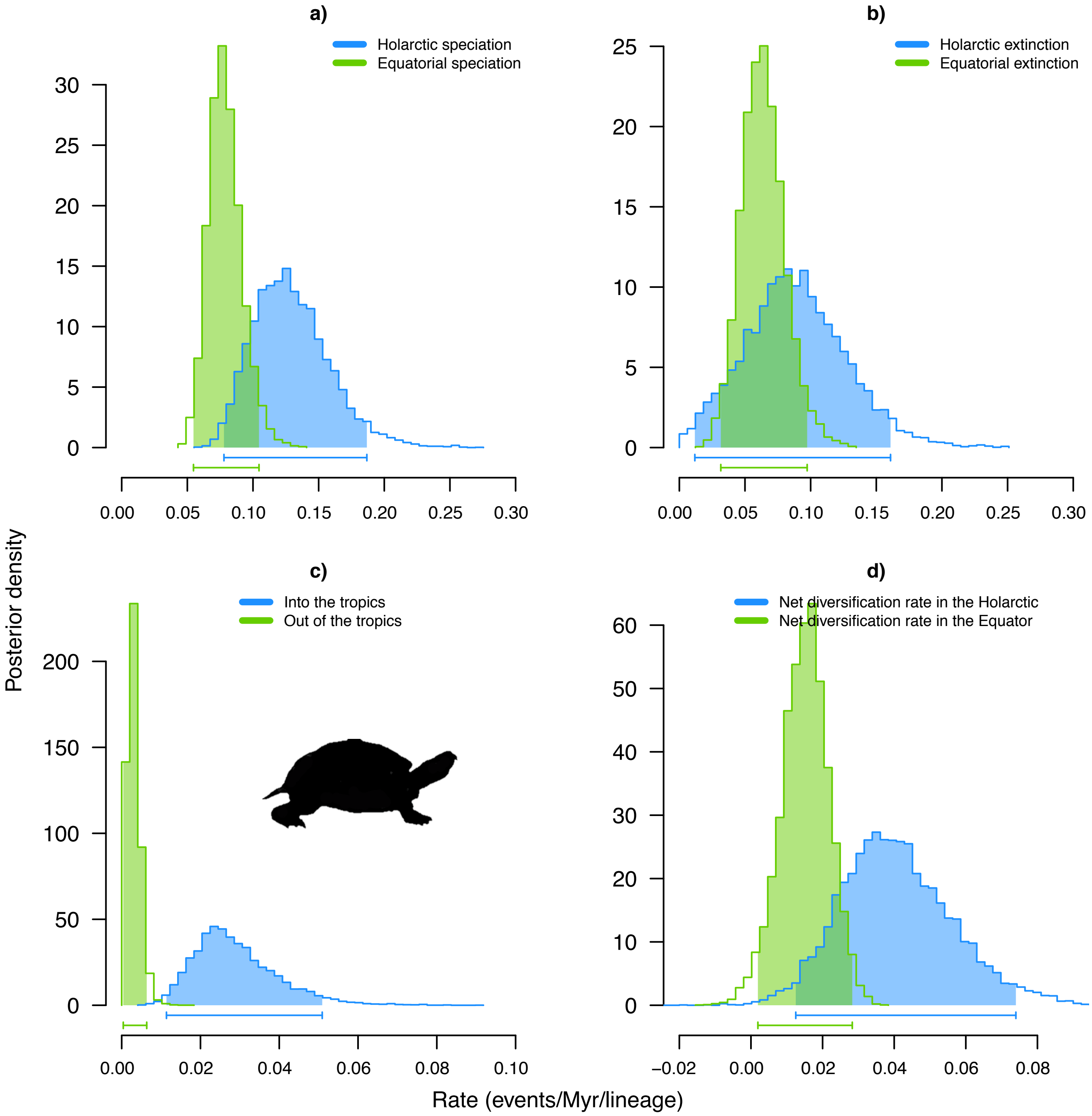


Figure S1a Latitudinal diversification pattern of turtles inferred with BiSSE showing the estimates for speciation (a), extinction (b), transition (c) and net diversification (d) rates for Holarctic and Equatorial species. Bayesian posterior distributions were computed using MCMC analyses with the full model on the MCC tree. Bars below each distribution correspond to the shaded area and represent the 95% credibility interval of each estimated parameter.

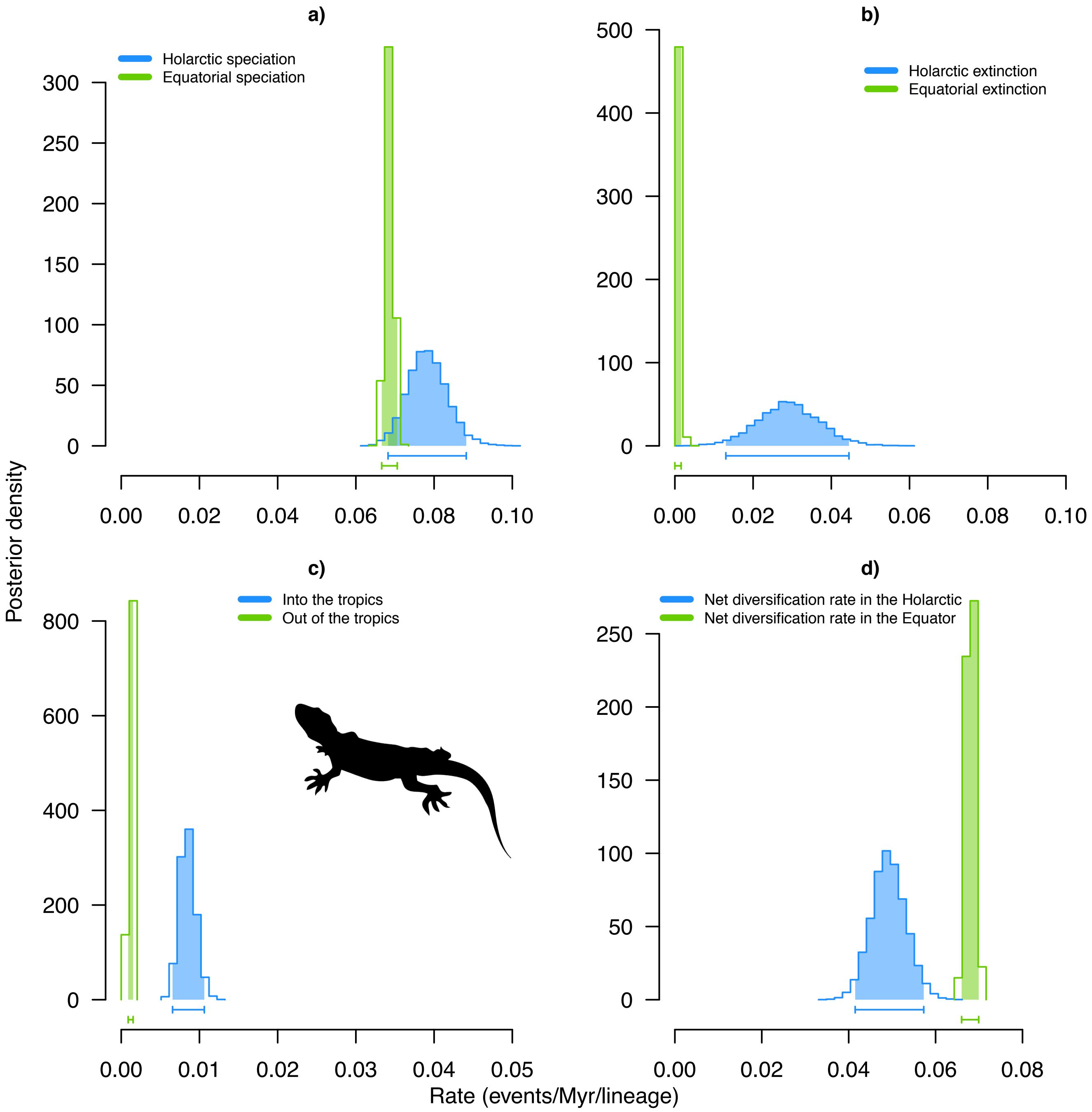


Figure S1b. Latitudinal diversification pattern of squamates inferred with BiSSE showing the estimates for speciation (a), extinction (b), transition (c) and net diversification (d) rates for Holarctic and Equatorial species. Bayesian posterior distributions were computed using MCMC analyses with the full model on the MCC tree. Bars below each distribution correspond to the shaded area and represent the 95% credibility interval of each estimated parameter.
