## Supplementary Fig. 2 for "Ancient tropical extinctions contributed to the latitudinal diversity gradient"

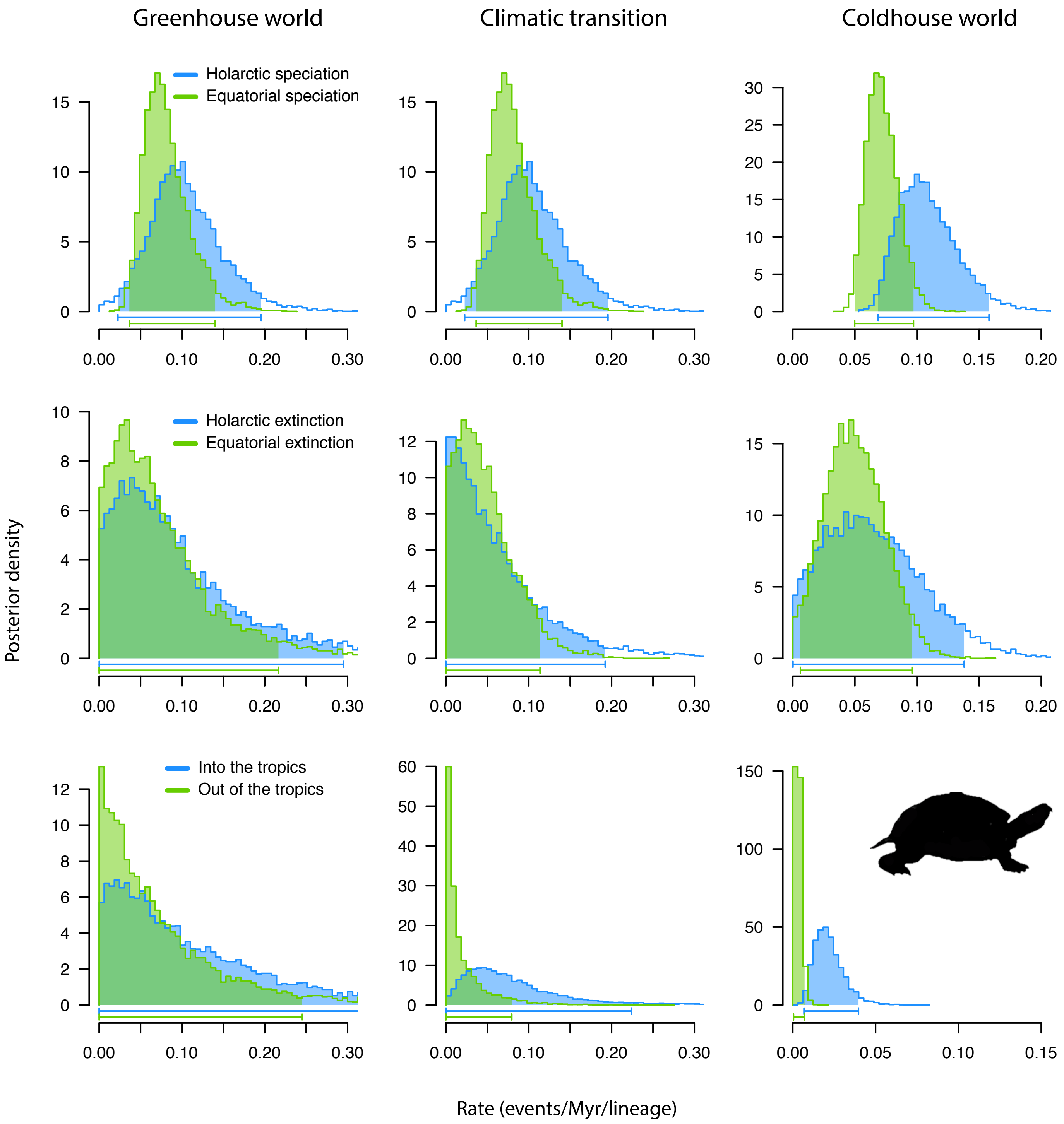


Figure S2a. Latitudinal time-dependent diversification patterns of turtles inferred with BiSSE.td showing the diversification dynamics of turtles during the greenhouse (left panel), the climatic transition (central panel), and the coldhouse (right panel) intervals for Holarctic and Equatorial species. In this analysis, the climatic transition begins 51 Ma and ends 23 Ma. Bayesian posterior distributions were computed using MCMC analyses with the full model on the MCC tree. Bars below each distribution correspond to the shaded area and represent the 95% credibility interval of each estimated parameter.

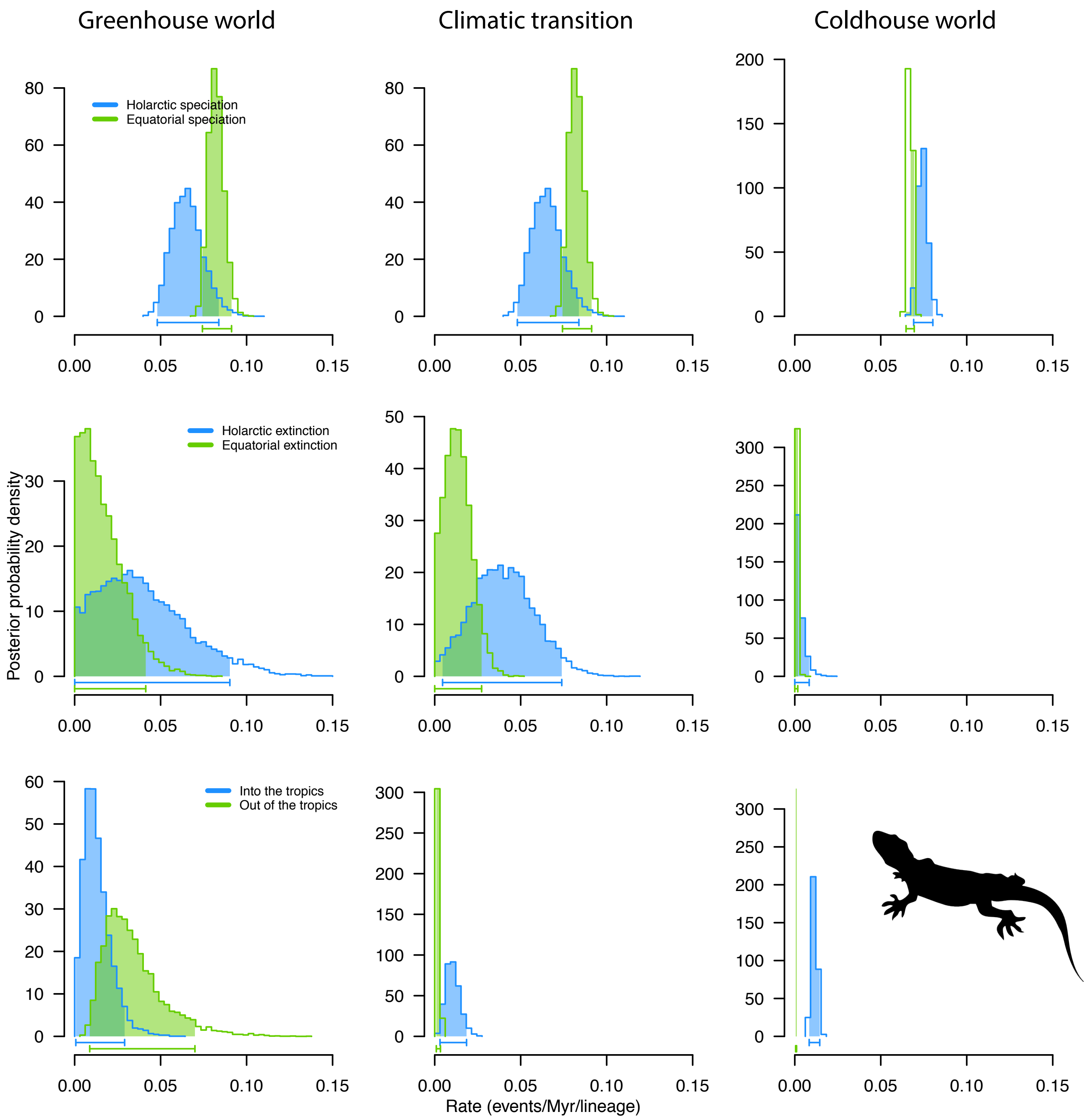


Figure S2b. Latitudinal time-dependent diversification patterns of squamates inferred with BiSSE.td showing the diversification dynamics of turtles during the greenhouse (left panel), the climatic transition (central panel), and the coldhouse (right panel) intervals for Holarctic and Equatorial species. In this analysis, the climatic transition begins 51 Ma and ends 23 Ma. Bayesian posterior distributions were computed using MCMC analyses with the full model on the MCC tree. Bars below each distribution correspond to the shaded area and represent the 95% credibility interval of each estimated parameter.
