## Supplementary Fig. 3 for "Ancient tropical extinctions contributed to the latitudinal diversity gradient"

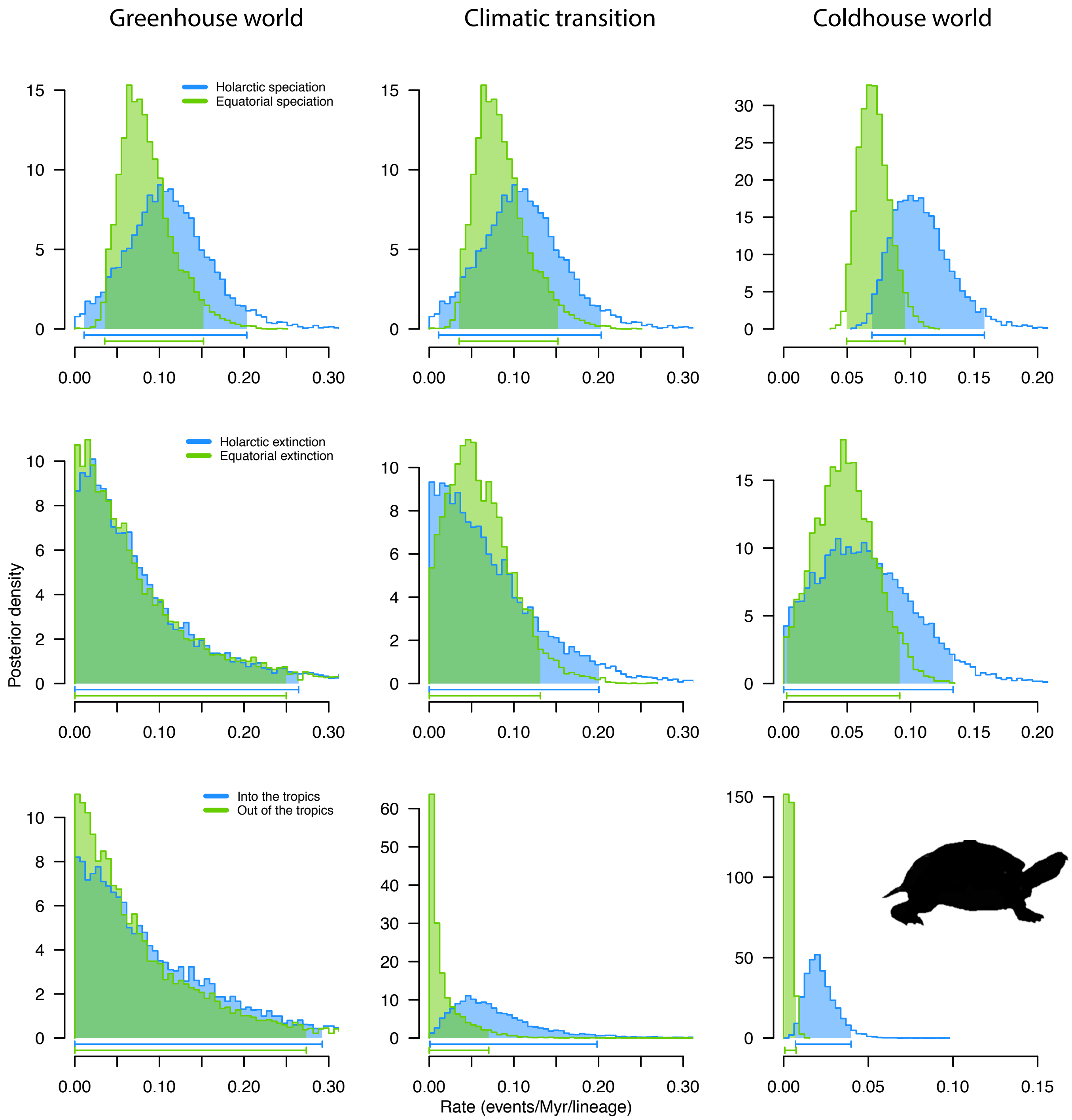


Figure S3a. Latitudinal time-dependent diversification patterns of turtles inferred with BiSSE.td showing the diversification dynamics during the greenhouse (left panel), the climatic transition (central panel), and the coldhouse (right panel) intervals for Holarctic and Equatorial species. In this analysis, the climatic transition begins 66 Ma and ends 23 Ma. Bayesian posterior distributions were computed using MCMC analyses with the full model on the MCC tree. Bars below each distribution correspond to the shaded area and represent the 95% credibility interval of each estimated parameter.

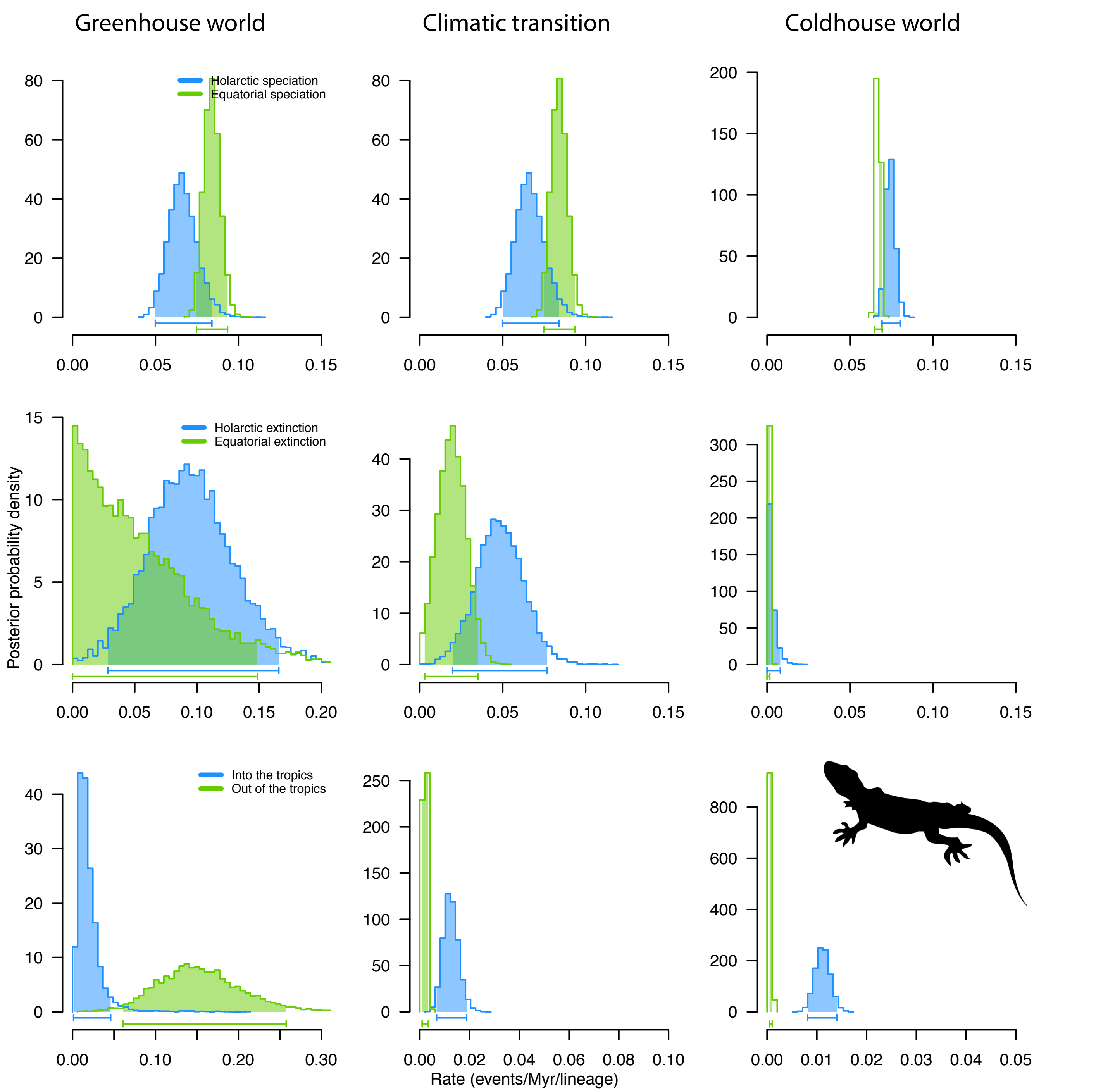


Figure S3b. Latitudinal time-dependent diversification patterns of squamates inferred with BiSSE.td showing the diversification dynamics during the greenhouse (left panel), the climatic transition (central panel), and the coldhouse (right panel) intervals for Holarctic and Equatorial species. In this analysis, the climatic transition begins 66 Ma and ends 23 Ma. Bayesian posterior distributions were computed using MCMC analyses with the full model on the MCC tree. Bars below each distribution correspond to the shaded area and represent the 95% credibility interval of each estimated parameter.
