## Supplementary Fig. 4 for "Ancient tropical extinctions contributed to the latitudinal diversity gradient"

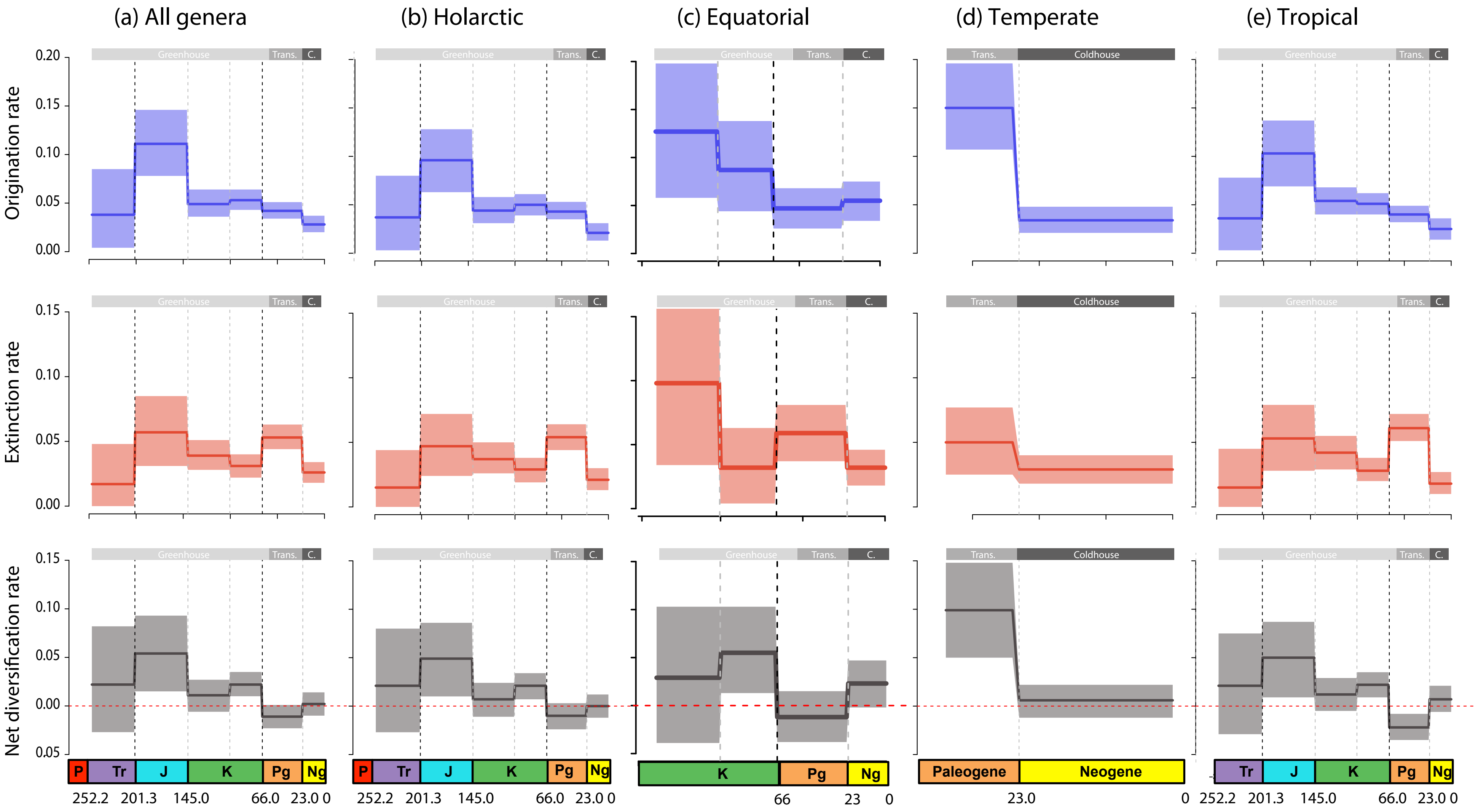


Figure S4. Global pattern of turtle diversification based on the fossil record and estimated using time bins as defined by geological periods. Results are shown independently for the total genus diversity (a), the Holarctic genus diversity (b), the Equatorial genus diversity only (c), the temperate genus diversity (d), and the tropical genus diversity (e). Origination (blue) and extinction (red) rates were estimated using time bins as defined by epochs of the geological timescale (on the top, climatic periods are shown as follows: Greenhouse, Trans. = climatic transition, and C. = coldhouse). Solid lines indicate mean posterior rates, whereas the shaded areas show 95% confidence intervals. Net diversification rates (black) are the difference between origination and extinction. The vertical lines indicate the boundaries between geological boundaries and major mass extinction events. Tr, Triassic; J, Jurassic; K, Cretaceous; Pg, Paleogene; and Ng, Neogene.
