## Supplementary Fig. 9 for "Ancient tropical extinctions contributed to the latitudinal diversity gradient"

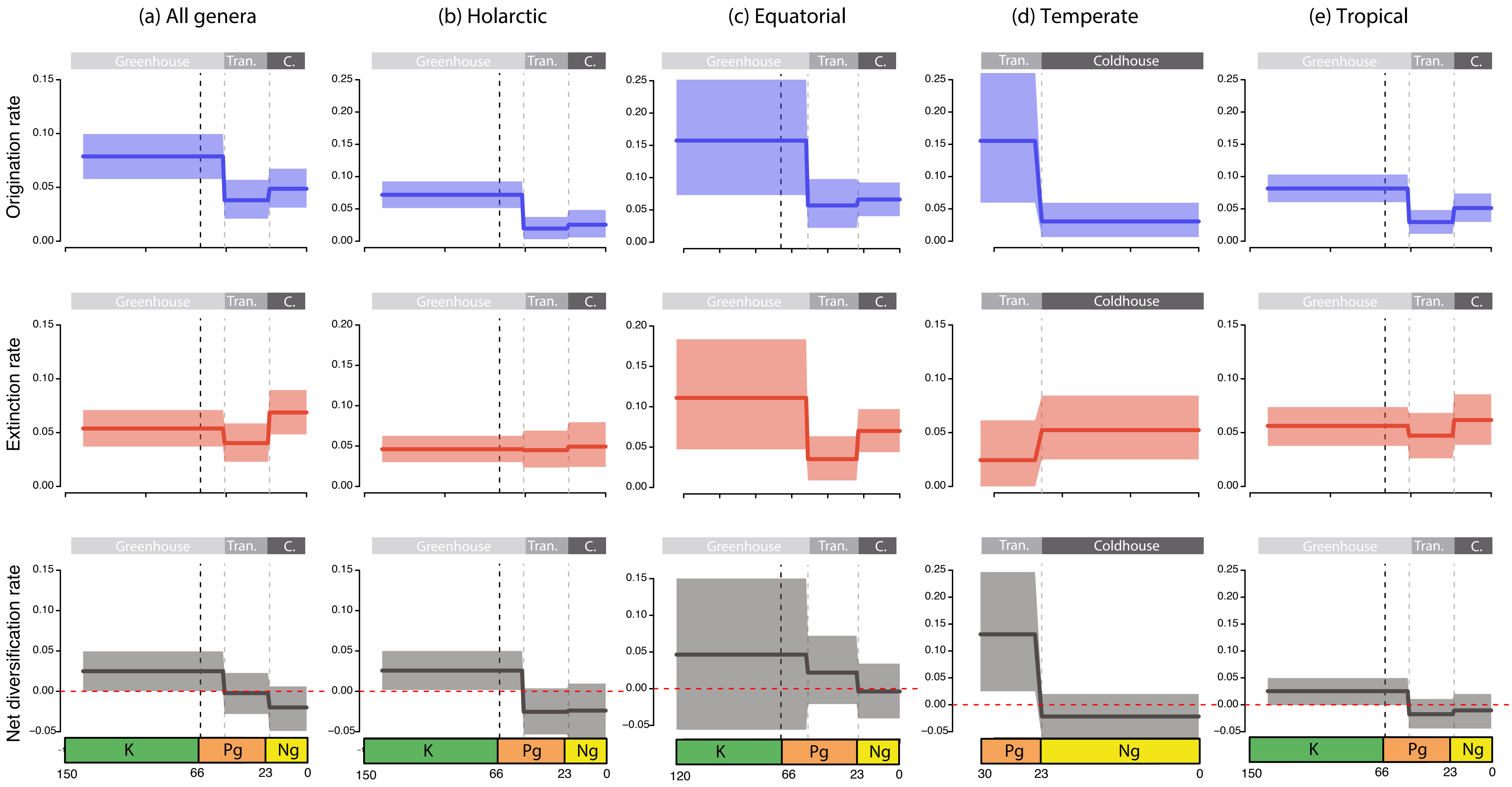


Figure S9. Global pattern of crocodiles diversification based on the fossil record and estimated using time bins as defined by climatic epochs. Results are shown for all the datasets combined (a), the Holarctic genus diversity only (b), the Equatorial genus diversity (c), the temperate genus diversity (d), and the tropical genus diversity (e). Origination (blue) and extinction (red) rates were estimated using time bins as defined by climatic epochs (on the top, climatic periods are shown as follows: Greenhouse, Trans. = climatic transition, and C. = coldhouse). Solid lines indicate mean posterior rates, whereas the shaded areas show 95% confidence intervals. Net diversification rates (black) are the difference between origination and extinction. The vertical lines indicate the boundaries between geological boundaries and major mass extinction events. Tr, Triassic; J, Jurassic; K, Cretaceous; Pg, Paleogene; and Ng, Neogene.
