## Supplementary Fig. 11 for "Ancient tropical extinctions contributed to the latitudinal diversity gradient"

Figure S11. Biogeographic reconstruction of Crocodiles on DEC under the Unconstrained model. Letters in the nodes represent ancestral ranges and range inheritance scenarios; a = Holarctic; b = Equator; c = Southern temperate regions. Range extinction events are represented with red asterisks, range dispersal events with black arrows.

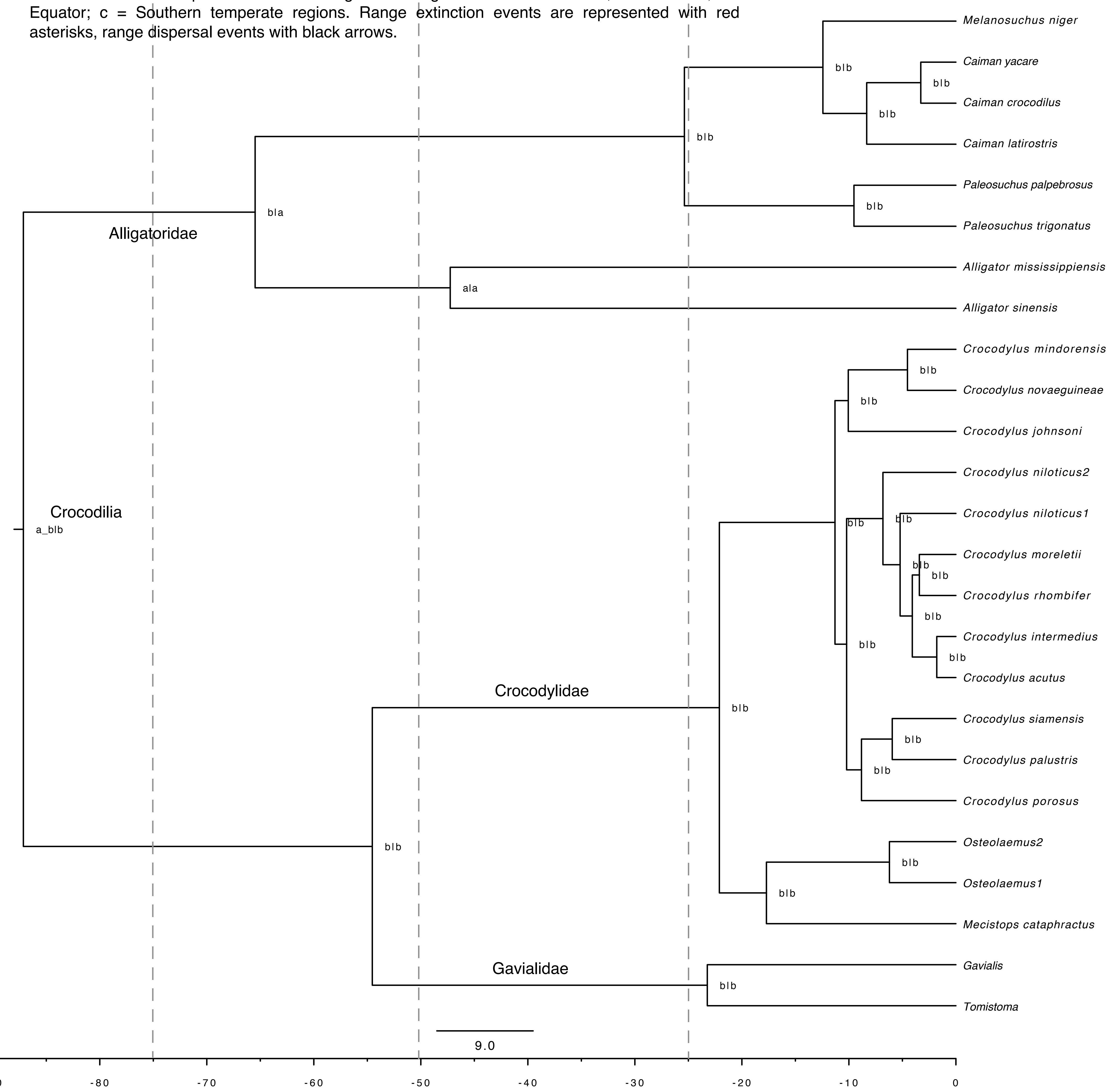
