## Supplementary Fig. 12 for "Ancient tropical extinctions contributed to the latitudinal diversity gradient"

Figure S12. Biogeographical reconstruction of Testudines inferred with DEC including soft fossil constraints (SFC, see main text). Coloured circles at the tips and nodes represent current and ancestral ranges, respectively, and correspond with the discrete areas in the legend.

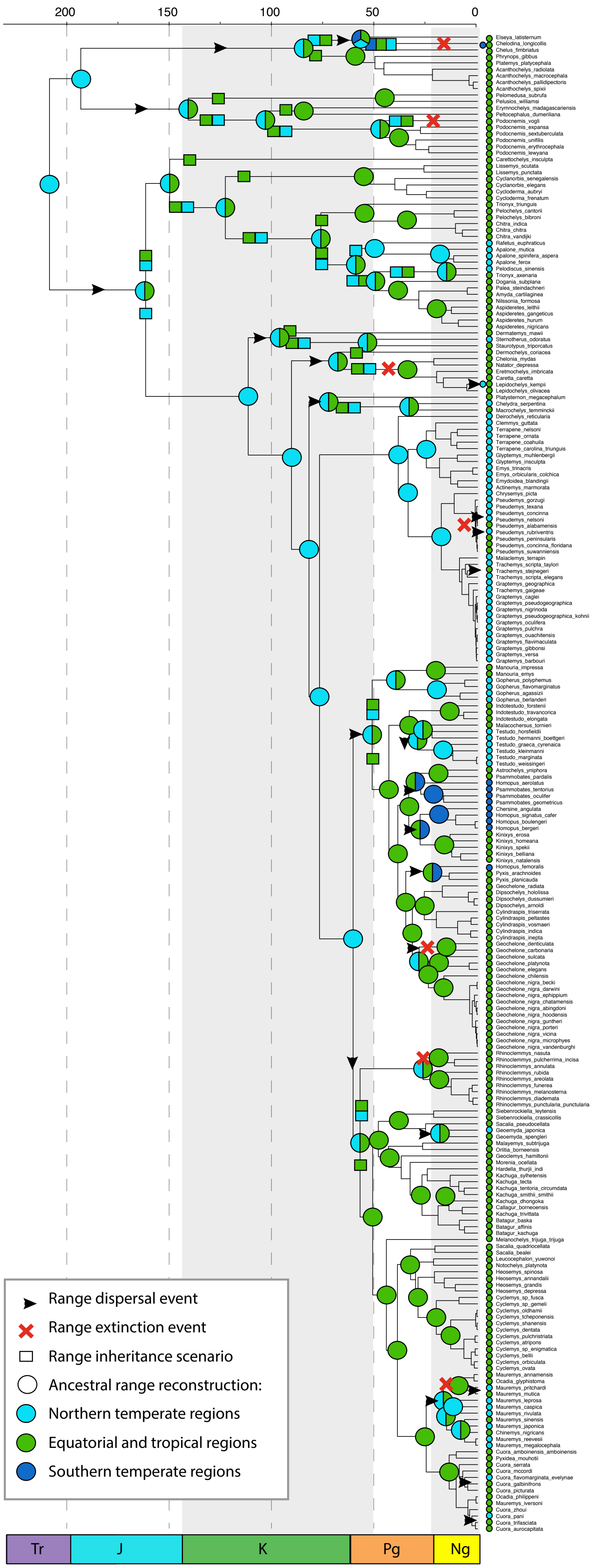
