## Supplementary Fig. 16 for "Ancient tropical extinctions contributed to the latitudinal diversity gradient"

Figure S16. Biogeographic reconstruction of Crocodiles showing the effects of the incorporation of fossil information into biogeographic inference under a hard fossil constraint (HFC, see main text) model. Letters in the nodes represent ancestral ranges and range inheritance scenarios; a = Holarctic; b = Equator; c = Southern temperate regions. Black circles indicate fossil range constraints included in the analysis, with numbers corresponding with taxa in Table S3. Range extinction events are represented with red asterisks, range dispersal events with black arrows.

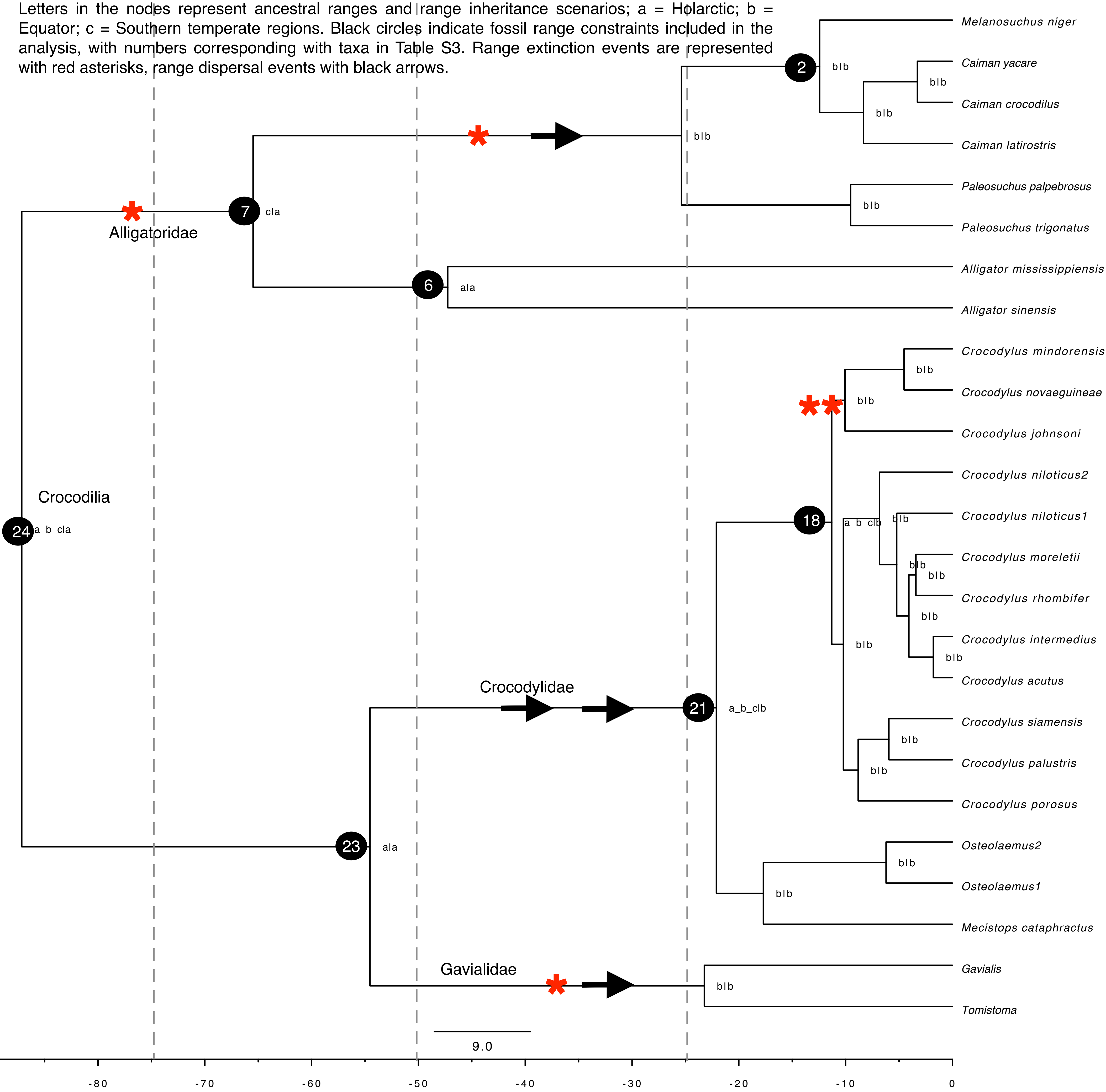
