## Supplementary Fig. 17 for "Ancient tropical extinctions contributed to the latitudinal diversity gradient"

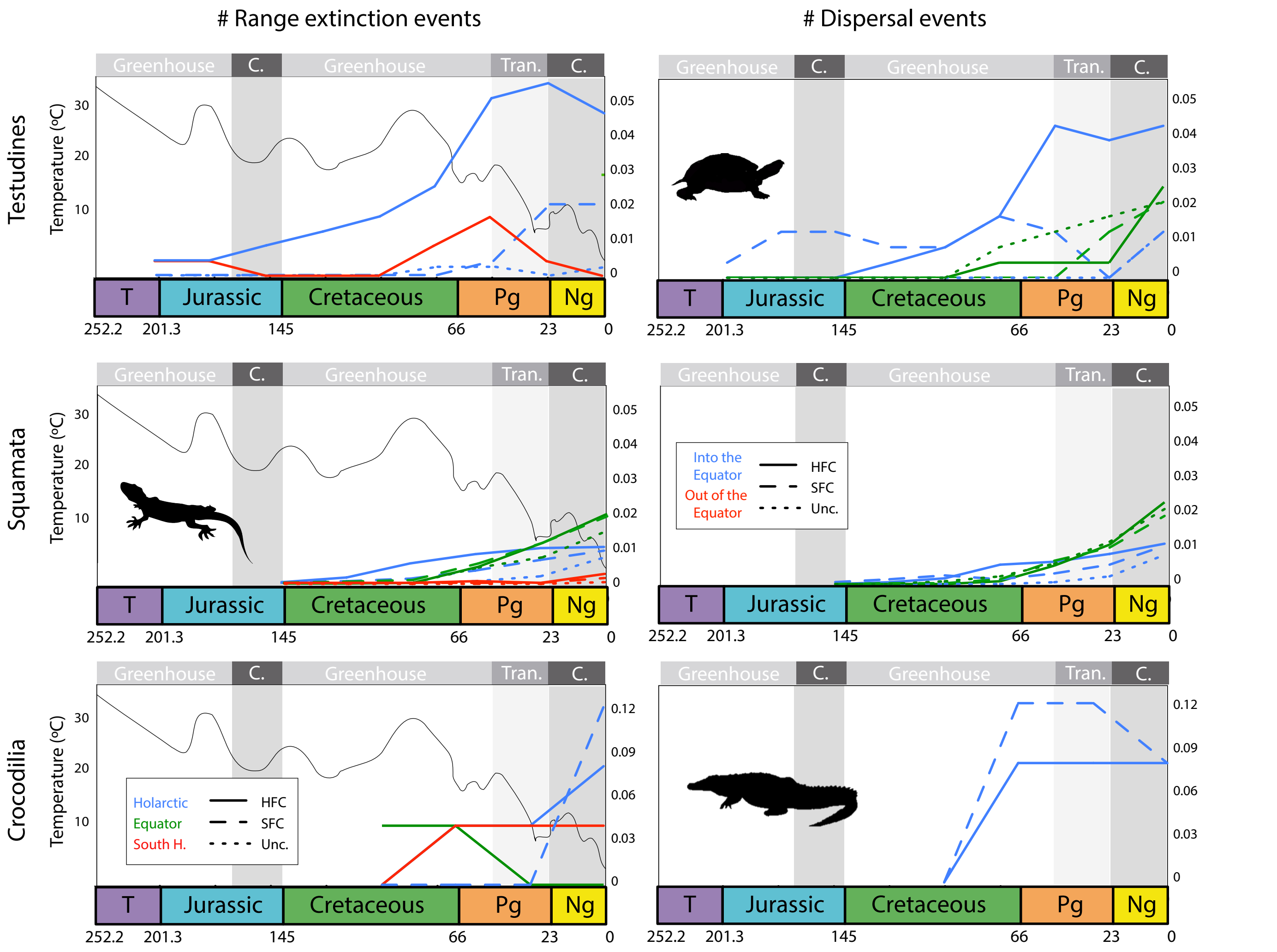


Figure S17. Estimated number of range-extinction and dispersal events across regions and through time for Testudines, Squamata and Crocodiles, relative to the current number of lineages in each group. Results are compared between the unconstrained (Unc.), based on present evidence only, and the fossil-based hard (HFC) and soft fossil constraint (SFC) biogeographic models. The global mean temperature curve is modified from Zachos et al. (2008). Abbreviations: Tr, Triassic; J, Jurassic; K, Cretaceous; Pg, Paleogene; and Ng, Neogene, Trans. = climatic transition, and C. = coldhouse.
