## Supplementary Table 1 for "Ancient tropical extinctions contributed to the latitudinal diversity gradient"

**Table S1.** Geographic distribution of extant turtle species based on the *Reptile Database* and the *IUCN Red List*. For each species, we coded its geographic range as defined by 3 discrete areas. We also provide the coding for BiSSE analysis: whether species were distributed in the Holarctic (0) or Equatorial areas (1). Note that southern temperate areas are not considered for this analysis. *Abbreviations for the 3 areas: North Temp. and South Temp. = Northern (Southern) temperate regions.*

| **Species** | **North Temp.** | **Tropical** | **South Temp.** | **BiSSE** |
| --- | --- | --- | --- | --- |
| *Acanthochelys macrocephala* | 0 | 1 | 0 | 1 |
| *Acanthochelys pallidipectoris* | 0 | 1 | 0 | 1 |
| *Acanthochelys radiolata* | 0 | 1 | 0 | 1 |
| *Acanthochelys spixii* | 0 | 1 | 0 | 1 |
| *Actinemys marmorata* | 1 | 0 | 0 | 0 |
| *Amyda cartilaginea* | 0 | 1 | 0 | 1 |
| *Apalone ferox* | 1 | 0 | 0 | 0 |
| *Apalone mutica* | 1 | 0 | 0 | 0 |
| *Apalone spinifera aspera* | 1 | 0 | 0 | 0 |
| *Aspideretes gangeticus* | 0 | 1 | 0 | 1 |
| *Aspideretes hurum* | 0 | 1 | 0 | 1 |
| *Aspideretes leithii* | 0 | 1 | 0 | 1 |
| *Aspideretes nigricans* | 0 | 1 | 0 | 1 |
| *Astrochelys yniphora* | 0 | 1 | 0 | 1 |
| *Batagur affinis* | 0 | 1 | 0 | 1 |
| *Batagur baska* | 0 | 1 | 0 | 1 |
| *Batagur kachuga* | 0 | 1 | 0 | 1 |
| *Callagur borneoensis* | 0 | 1 | 0 | 1 |
| *Caretta caretta* | 0 | 1 | 0 | 1 |
| *Carettochelys insculpta* | 0 | 1 | 0 | 1 |
| *Chelodina longicollis* | 0 | 1 | 1 | 1 |
| *Chelonia mydas* | 0 | 1 | 0 | 1 |
| *Chelus fimbriatus* | 0 | 1 | 0 | 1 |
| *Chelydra serpentina* | 1 | 0 | 0 | 0 |
| *Chersine angulata* | 0 | 0 | 1 | 0 |
| *Chinemys nigricans* | 0 | 1 | 0 | 1 |
| *Chitra chitra* | 0 | 1 | 0 | 1 |
| *Chitra indica* | 0 | 1 | 0 | 1 |
| *Chitra vandijki* | 0 | 1 | 0 | 1 |
| *Chrysemys picta* | 1 | 0 | 0 | 0 |
| *Clemmys guttata* | 1 | 0 | 0 | 0 |
| *Cuora amboinensis amboinensis* | 0 | 1 | 0 | 1 |
| *Cuora aurocapitata* | 0 | 1 | 0 | 1 |
| *Cuora flavomarginata evelynae* | 1 | 0 | 0 | 0 |
| *Cuora galbinifrons* | 0 | 1 | 0 | 1 |
| *Cuora mccordi* | 0 | 1 | 0 | 1 |
| *Cuora pani* | 1 | 0 | 0 | 0 |
| *Cuora picturata* | 0 | 1 | 0 | 1 |
| *Cuora serrata* | 0 | 1 | 0 | 1 |
| *Cuora trifasciata* | 0 | 1 | 0 | 1 |
| *Cuora zhoui* | 0 | 1 | 0 | 1 |
| *Cyclanorbis elegans* | 0 | 1 | 0 | 1 |
| *Cyclanorbis senegalensis* | 0 | 1 | 0 | 1 |
| *Cyclemys atripons* | 0 | 1 | 0 | 1 |
| *Cyclemys bellii* | 0 | 1 | 0 | 1 |
| *Cyclemys dentata* | 0 | 1 | 0 | 1 |
| *Cyclemys oldhamii* | 0 | 1 | 0 | 1 |
| *Cyclemys orbiculata* | 0 | 1 | 0 | 1 |
| *Cyclemys ovata* | 0 | 1 | 0 | 1 |
| *Cyclemys pulchristriata* | 0 | 1 | 0 | 1 |
| *Cyclemys shanensis* | 0 | 1 | 0 | 1 |
| *Cyclemys sp. enigmatica* | 0 | 1 | 0 | 1 |
| *Cyclemys sp. fusca* | 0 | 1 | 0 | 1 |
| *Cyclemys sp. gemeli* | 0 | 1 | 0 | 1 |
| *Cyclemys tcheponensis* | 0 | 1 | 0 | 1 |
| *Cycloderma aubryi* | 0 | 1 | 0 | 1 |
| *Cycloderma frenatum* | 0 | 1 | 0 | 1 |
| *Cylindraspis indica* | 0 | 1 | 0 | 1 |
| *Cylindraspis inepta* | 0 | 1 | 0 | 1 |
| *Cylindraspis peltastes* | 0 | 1 | 0 | 1 |
| *Cylindraspis triserrata* | 0 | 1 | 0 | 1 |
| *Cylindraspis vosmaeri* | 0 | 1 | 0 | 1 |
| *Deirochelys reticularia* | 1 | 0 | 0 | 0 |
| *Dermatemys mawii* | 0 | 1 | 0 | 1 |
| *Dermochelys coriacea* | 0 | 1 | 0 | 1 |
| *Dipsochelys arnoldi* | 0 | 1 | 0 | 1 |
| *Dipsochelys dussumieri* | 0 | 1 | 0 | 1 |
| *Dipsochelys hololissa* | 0 | 1 | 0 | 1 |
| *Dogania subplana* | 0 | 1 | 0 | 1 |
| *Elseya latisternum* | 0 | 1 | 0 | 1 |
| *Emydoidea blandingii* | 1 | 0 | 0 | 0 |
| *Emys orbicularis colchica* | 1 | 0 | 0 | 0 |
| *Emys trinacris* | 1 | 0 | 0 | 0 |
| *Eretmochelys imbricata* | 0 | 1 | 0 | 1 |
| *Erymnochelys madagascariensis* | 0 | 1 | 0 | 1 |
| *Geochelone carbonaria* | 0 | 1 | 0 | 1 |
| *Geochelone chilensis* | 0 | 1 | 0 | 1 |
| *Geochelone denticulata* | 0 | 1 | 0 | 1 |
| *Geochelone elegans* | 0 | 1 | 0 | 1 |
| *Geochelone nigra abingdoni* | 0 | 1 | 0 | 1 |
| *Geochelone nigra becki* | 0 | 1 | 0 | 1 |
| *Geochelone nigra chatamensis* | 0 | 1 | 0 | 1 |
| *Geochelone nigra darwini* | 0 | 1 | 0 | 1 |
| *Geochelone nigra ephippium* | 0 | 1 | 0 | 1 |
| *Geochelone nigra guntheri* | 0 | 1 | 0 | 1 |
| *Geochelone nigra hoodensis* | 0 | 1 | 0 | 1 |
| *Geochelone nigra microphyes* | 0 | 1 | 0 | 1 |
| *Geochelone nigra porteri* | 0 | 1 | 0 | 1 |
| *Geochelone nigra vandenburghi* | 0 | 1 | 0 | 1 |
| *Geochelone nigra vicina* | 0 | 1 | 0 | 1 |
| *Geochelone platynota* | 0 | 1 | 0 | 1 |
| *Geochelone radiata* | 0 | 1 | 0 | 1 |
| *Geochelone sulcata* | 0 | 1 | 0 | 1 |
| *Geoclemys hamiltonii* | 0 | 1 | 0 | 1 |
| *Geoemyda japonica* | 1 | 0 | 0 | 0 |
| *Geoemyda spengleri* | 0 | 1 | 0 | 1 |
| *Glyptemys insculpta* | 1 | 0 | 0 | 0 |
| *Glyptemys muhlenbergii* | 1 | 0 | 0 | 0 |
| *Gopherus agassizii* | 1 | 0 | 0 | 0 |
| *Gopherus berlanderi* | 1 | 0 | 0 | 0 |
| *Gopherus flavomarginatus* | 1 | 0 | 0 | 0 |
| *Gopherus polyphemus* | 1 | 0 | 0 | 0 |
| *Graptemys barbouri* | 1 | 0 | 0 | 0 |
| *Graptemys caglei* | 1 | 0 | 0 | 0 |
| *Graptemys flavimaculata* | 1 | 0 | 0 | 0 |
| *Graptemys geographica* | 1 | 0 | 0 | 0 |
| *Graptemys gibbonsi* | 1 | 0 | 0 | 0 |
| *Graptemys nigrinoda* | 1 | 0 | 0 | 0 |
| *Graptemys oculifera* | 1 | 0 | 0 | 0 |
| *Graptemys ouachitensis* | 1 | 0 | 0 | 0 |
| *Graptemys pseudogeographica* | 1 | 0 | 0 | 0 |
| *Graptemys pseudogeographica kohnii* | 1 | 0 | 0 | 0 |
| *Graptemys pulchra* | 1 | 0 | 0 | 0 |
| *Graptemys versa* | 1 | 0 | 0 | 0 |
| *Hardella thurjii indi* | 0 | 1 | 0 | 1 |
| *Heosemys annandalii* | 0 | 1 | 0 | 1 |
| *Heosemys depressa* | 0 | 1 | 0 | 1 |
| *Heosemys grandis* | 0 | 1 | 0 | 1 |
| *Heosemys spinosa* | 0 | 1 | 0 | 1 |
| *Homopus aerolatus* | 0 | 0 | 1 | 0 |
| *Homopus bergeri* | 0 | 0 | 1 | 0 |
| *Homopus boulengeri* | 0 | 0 | 1 | 0 |
| *Homopus femoralis* | 0 | 0 | 1 | 0 |
| *Homopus signatus cafer* | 0 | 0 | 1 | 0 |
| *Indotestudo elongata* | 0 | 1 | 0 | 1 |
| *Indotestudo forstenii* | 0 | 1 | 0 | 1 |
| *Indotestudo travancorica* | 0 | 1 | 0 | 1 |
| *Kachuga dhongoka* | 0 | 1 | 0 | 1 |
| *Kachuga smithii smithii* | 0 | 1 | 0 | 1 |
| *Kachuga sylhetensis* | 0 | 1 | 0 | 1 |
| *Kachuga tecta* | 0 | 1 | 0 | 1 |
| *Kachuga tentoria circumdata* | 0 | 1 | 0 | 1 |
| *Kachuga trivittata* | 0 | 1 | 0 | 1 |
| *Kinixys belliana* | 0 | 1 | 0 | 1 |
| *Kinixys erosa* | 0 | 1 | 0 | 1 |
| *Kinixys homeana* | 0 | 1 | 0 | 1 |
| *Kinixys natalensis* | 0 | 1 | 0 | 1 |
| *Kinixys spekii* | 0 | 1 | 0 | 1 |
| *Lepidochelys kempii* | 1 | 1 | 0 | 1 |
| *Lepidochelys olivacea* | 0 | 1 | 0 | 1 |
| *Leucocephalon yuwonoi* | 0 | 1 | 0 | 1 |
| *Lissemys punctata* | 0 | 1 | 0 | 1 |
| *Lissemys scutata* | 0 | 1 | 0 | 1 |
| *Macrochelys temminckii* | 0 | 1 | 0 | 1 |
| *Malaclemys terrapin* | 1 | 0 | 0 | 0 |
| *Malacochersus tornieri* | 0 | 1 | 0 | 1 |
| *Malayemys subtrijuga* | 0 | 1 | 0 | 1 |
| *Manouria emys* | 0 | 1 | 0 | 1 |
| *Manouria impressa* | 0 | 1 | 0 | 1 |
| *Mauremys annamensis* | 0 | 1 | 0 | 1 |
| *Mauremys caspica* | 1 | 0 | 0 | 0 |
| *Mauremys iversoni* | 0 | 1 | 0 | 1 |
| *Mauremys japonica* | 1 | 0 | 0 | 0 |
| *Mauremys leprosa* | 1 | 0 | 0 | 0 |
| *Mauremys megalocephala* | 1 | 0 | 0 | 0 |
| *Mauremys mutica* | 0 | 1 | 0 | 1 |
| *Mauremys pritchardi* | 1 | 0 | 0 | 0 |
| *Mauremys reevesii* | 1 | 0 | 0 | 0 |
| *Mauremys rivulata* | 1 | 0 | 0 | 0 |
| *Mauremys sinensis* | 0 | 1 | 0 | 1 |
| *Melanochelys trijuga trijuga* | 0 | 1 | 0 | 1 |
| *Morenia ocellata* | 0 | 1 | 0 | 1 |
| *Natator depressa* | 0 | 1 | 0 | 1 |
| *Nilssonia formosa* | 0 | 1 | 0 | 1 |
| *Notochelys platynota* | 0 | 1 | 0 | 1 |
| *Ocadia glyphistoma* | 0 | 1 | 0 | 1 |
| *Ocadia philippeni* | 0 | 1 | 0 | 1 |
| *Orlitia borneensis* | 0 | 1 | 0 | 1 |
| *Palea steindachneri* | 0 | 1 | 0 | 1 |
| *Pelochelys bibroni* | 0 | 1 | 0 | 1 |
| *Pelochelys cantorii* | 0 | 1 | 0 | 1 |
| *Pelodiscus sinensis* | 1 | 0 | 0 | 0 |
| *Pelomedusa subrufa* | 0 | 1 | 0 | 1 |
| *Peltocephalus dumeriliana* | 0 | 1 | 0 | 1 |
| *Pelusios williamsi* | 0 | 1 | 0 | 1 |
| *Phrynops gibbus* | 0 | 1 | 0 | 1 |
| *Platemys platycephala* | 0 | 1 | 0 | 1 |
| *Platysternon megacephalum* | 0 | 1 | 0 | 1 |
| *Podocnemis erythrocephala* | 0 | 1 | 0 | 1 |
| *Podocnemis expansa* | 0 | 1 | 0 | 1 |
| *Podocnemis lewyana* | 0 | 1 | 0 | 1 |
| *Podocnemis sextuberculata* | 0 | 1 | 0 | 1 |
| *Podocnemis unifilis* | 0 | 1 | 0 | 1 |
| *Podocnemis vogli* | 0 | 1 | 0 | 1 |
| *Psammobates geometricus* | 0 | 0 | 1 | 0 |
| *Psammobates oculifer* | 0 | 0 | 1 | 0 |
| *Psammobates pardalis* | 0 | 1 | 0 | 1 |
| *Psammobates tentorius* | 0 | 0 | 1 | 0 |
| *Pseudemys alabamensis* | 0 | 1 | 0 | 1 |
| *Pseudemys concinna* | 1 | 0 | 0 | 0 |
| *Pseudemys concinna floridana* | 0 | 1 | 0 | 1 |
| *Pseudemys gorzugi* | 1 | 0 | 0 | 0 |
| *Pseudemys nelsoni* | 0 | 1 | 0 | 1 |
| *Pseudemys peninsularis* | 0 | 1 | 0 | 1 |
| *Pseudemys rubriventris* | 1 | 0 | 0 | 0 |
| *Pseudemys suwanniensis* | 0 | 1 | 0 | 1 |
| *Pseudemys texana* | 1 | 0 | 0 | 0 |
| *Pyxidea mouhotii* | 0 | 1 | 0 | 1 |
| *Pyxis arachnoides* | 0 | 1 | 0 | 1 |
| *Pyxis planicauda* | 0 | 1 | 0 | 1 |
| *Rafetus euphraticus* | 1 | 0 | 0 | 0 |
| *Rhinoclemmys annulata* | 0 | 1 | 0 | 1 |
| *Rhinoclemmys areolata* | 0 | 1 | 0 | 1 |
| *Rhinoclemmys diademata* | 0 | 1 | 0 | 1 |
| *Rhinoclemmys funerea* | 0 | 1 | 0 | 1 |
| *Rhinoclemmys melanosterna* | 0 | 1 | 0 | 1 |
| *Rhinoclemmys nasuta* | 0 | 1 | 0 | 1 |
| *Rhinoclemmys pulcherrima incisa* | 0 | 1 | 0 | 1 |
| *Rhinoclemmys punctularia punctularia* | 0 | 1 | 0 | 1 |
| *Rhinoclemmys rubida* | 0 | 1 | 0 | 1 |
| *Sacalia bealei* | 0 | 1 | 0 | 1 |
| *Sacalia pseudocellata* | 0 | 1 | 0 | 1 |
| *Sacalia quadriocellata* | 0 | 1 | 0 | 1 |
| *Siebenrockiella crassicollis* | 0 | 1 | 0 | 1 |
| *Siebenrockiella leytensis* | 0 | 1 | 0 | 1 |
| *Staurotypus triporcatus* | 0 | 1 | 0 | 1 |
| *Sternotherus odoratus* | 1 | 0 | 0 | 0 |
| *Terrapene carolina triunguis* | 1 | 0 | 0 | 0 |
| *Terrapene coahuila* | 1 | 0 | 0 | 0 |
| *Terrapene nelsoni* | 1 | 0 | 0 | 0 |
| *Terrapene ornata* | 1 | 0 | 0 | 0 |
| *Testudo graeca cyrenaica* | 1 | 0 | 0 | 0 |
| *Testudo hermanni boettgeri* | 1 | 0 | 0 | 0 |
| *Testudo horsfieldii* | 1 | 0 | 0 | 0 |
| *Testudo kleinmanni* | 1 | 0 | 0 | 0 |
| *Testudo marginata* | 1 | 0 | 0 | 0 |
| *Testudo weissingeri* | 1 | 0 | 0 | 0 |
| *Trachemys gaigeae* | 1 | 0 | 0 | 0 |
| *Trachemys scripta elegans* | 1 | 0 | 0 | 0 |
| *Trachemys scripta taylori* | 1 | 0 | 0 | 0 |
| *Trachemys stejnegeri* | 0 | 1 | 0 | 1 |
| *Trionyx axenaria* | 0 | 1 | 0 | 1 |
| *Trionyx triunguis* | 0 | 1 | 0 | 1 |
