## Supplementary Table 2 for "Ancient tropical extinctions contributed to the latitudinal diversity gradient"

**Table S2.** Geographic distribution of extant squamate species based on the *Reptile Database* and the *IUCN Red List*. For each species, we coded its geographic range as defined by 3 discrete areas. We also provide the coding for BiSSE analysis: whether species were distributed in the Holarctic (0) or Equatorial areas (1). Note that southern temperate areas are not considered for this analysis. *Abbreviations for the 3 areas: North Temp. and South Temp. = Northern (Southern) temperate regions.*

| Species | North Temp. | Tropical | South Temp. | BiSSE |
| --- | --- | --- | --- | --- |
| *Ablepharus budaki* | 1 | 0 | 0 | 0 |
| *Ablepharus chernovi* | 1 | 0 | 0 | 0 |
| *Ablepharus kitaibelii* | 1 | 0 | 0 | 0 |
| *Ablepharus pannonicus* | 1 | 0 | 0 | 0 |
| *Abronia anzuetoi* | 0 | 1 | 0 | 1 |
| *Abronia aurita* | 0 | 1 | 0 | 1 |
| *Abronia campbelli* | 0 | 1 | 0 | 1 |
| *Abronia chiszari* | 0 | 1 | 0 | 1 |
| *Abronia fimbriata* | 0 | 1 | 0 | 1 |
| *Abronia frosti* | 0 | 1 | 0 | 1 |
| *Abronia graminea* | 0 | 1 | 0 | 1 |
| *Abronia lythrochila* | 0 | 1 | 0 | 1 |
| *Abronia matudai* | 0 | 1 | 0 | 1 |
| *Abronia mixteca* | 0 | 1 | 0 | 1 |
| *Abronia oaxacae* | 0 | 1 | 0 | 1 |
| *Abronia ornelasi* | 0 | 1 | 0 | 1 |
| *Acalyptophis peronii* | 0 | 1 | 0 | 1 |
| *Acanthocercus atricollis* | 0 | 1 | 0 | 1 |
| *Acanthodactylus aureus* | 1 | 0 | 0 | 0 |
| *Acanthodactylus beershebensis* | 1 | 0 | 0 | 0 |
| *Acanthodactylus blanci* | 1 | 0 | 0 | 0 |
| *Acanthodactylus boskianus* | 1 | 0 | 0 | 0 |
| *Acanthodactylus busacki* | 1 | 0 | 0 | 0 |
| *Acanthodactylus cantoris* | 1 | 0 | 0 | 0 |
| *Acanthodactylus erythrurus* | 1 | 0 | 0 | 0 |
| *Acanthodactylus gongrorhynchatus* | 1 | 0 | 0 | 0 |
| *Acanthodactylus longipes* | 1 | 0 | 0 | 0 |
| *Acanthodactylus maculatus* | 1 | 0 | 0 | 0 |
| *Acanthodactylus masirae* | 1 | 0 | 0 | 0 |
| *Acanthodactylus opheodurus* | 1 | 0 | 0 | 0 |
| *Acanthodactylus orientalis* | 1 | 0 | 0 | 0 |
| *Acanthodactylus pardalis* | 1 | 0 | 0 | 0 |
| *Acanthodactylus schmidti* | 1 | 0 | 0 | 0 |
| *Acanthodactylus schreiberi* | 1 | 0 | 0 | 0 |
| *Acanthodactylus scutellatus* | 1 | 0 | 0 | 0 |
| *Acanthodactylus tristrami* | 1 | 0 | 0 | 0 |
| *Acanthophis antarcticus* | 0 | 1 | 0 | 1 |
| *Acanthophis praelongus* | 0 | 1 | 0 | 1 |
| *Acanthosaura armata* | 0 | 1 | 0 | 1 |
| *Acanthosaura capra* | 0 | 1 | 0 | 1 |
| *Acanthosaura crucigera* | 0 | 1 | 0 | 1 |
| *Acanthosaura lepidogaster* | 0 | 1 | 0 | 1 |
| *Achalinus meiguensis* | 1 | 1 | 0 | 0 |
| *Achalinus rufescens* | 0 | 1 | 0 | 1 |
| *Acontias breviceps* | 0 | 1 | 0 | 1 |
| *Acontias gracilicauda* | 0 | 1 | 0 | 1 |
| *Acontias meleagris* | 0 | 1 | 0 | 1 |
| *Acontias percivali* | 0 | 1 | 0 | 1 |
| *Acontias plumbeus* | 0 | 1 | 0 | 1 |
| *Acontias poecilus* | 0 | 1 | 0 | 1 |
| *Acontophiops lineatus* | 0 | 1 | 0 | 1 |
| *Acrantophis dumerili* | 0 | 1 | 0 | 1 |
| *Acrantophis madagascariensis* | 0 | 1 | 0 | 1 |
| *Acrochordus arafurae* | 0 | 1 | 0 | 1 |
| *Acrochordus granulatus* | 0 | 1 | 0 | 1 |
| *Acrochordus javanicus* | 0 | 1 | 0 | 1 |
| *Acutotyphlops kunuaensis* | 0 | 1 | 0 | 1 |
| *Acutotyphlops subocularis* | 0 | 1 | 0 | 1 |
| *Adelophis foxi* | 0 | 1 | 0 | 1 |
| *Adelphicos quadrivirgatus* | 0 | 1 | 0 | 1 |
| *Adolfus africanus* | 0 | 1 | 0 | 1 |
| *Adolfus alleni* | 0 | 1 | 0 | 1 |
| *Adolfus jacksoni* | 0 | 1 | 0 | 1 |
| *Adolfus vauereselli* | 0 | 1 | 0 | 1 |
| *Aeluroscalabotes felinus* | 0 | 1 | 0 | 1 |
| *Afroablepharus africanus* | 0 | 1 | 0 | 1 |
| *Afroablepharus annobonensis* | 0 | 1 | 0 | 1 |
| *Afroablepharus wahlbergi* | 0 | 1 | 0 | 1 |
| *Afroedura karroica* | 0 | 1 | 0 | 1 |
| *Afroedura pondolia* | 0 | 1 | 0 | 1 |
| *Afrogecko porphyreus* | 0 | 1 | 0 | 1 |
| *Afrogecko swartbergensis* | 0 | 1 | 0 | 1 |
| *Afronatrix anoscopus* | 0 | 1 | 0 | 1 |
| *Agama aculeata* | 0 | 1 | 0 | 1 |
| *Agama agama* | 0 | 1 | 0 | 1 |
| *Agama anchietae* | 0 | 1 | 0 | 1 |
| *Agama armata* | 0 | 1 | 0 | 1 |
| *Agama atra* | 0 | 1 | 0 | 1 |
| *Agama boueti* | 1 | 0 | 0 | 0 |
| *Agama boulengeri* | 1 | 0 | 0 | 0 |
| *Agama castroviejoi* | 1 | 1 | 0 | 0 |
| *Agama caudospinosa* | 0 | 1 | 0 | 1 |
| *Agama doriae* | 0 | 1 | 0 | 1 |
| *Agama finchi* | 0 | 1 | 0 | 1 |
| *Agama gracilimembris* | 0 | 1 | 0 | 1 |
| *Agama hispida* | 0 | 1 | 0 | 1 |
| *Agama impalearis* | 0 | 1 | 0 | 1 |
| *Agama insularis* | 0 | 1 | 0 | 1 |
| *Agama kaimosae* | 0 | 1 | 0 | 1 |
| *Agama lionotus* | 0 | 1 | 0 | 1 |
| *Agama mwanzae* | 0 | 1 | 0 | 1 |
| *Agama paragama* | 0 | 1 | 0 | 1 |
| *Agama planiceps* | 0 | 1 | 0 | 1 |
| *Agama rueppelli* | 0 | 1 | 0 | 1 |
| *Agama sankaranica* | 0 | 1 | 0 | 1 |
| *Agama spinosa* | 1 | 0 | 0 | 0 |
| *Agama weidholzi* | 0 | 1 | 0 | 1 |
| *Agamura persica* | 1 | 0 | 0 | 0 |
| *Agkistrodon bilineatus* | 0 | 1 | 0 | 1 |
| *Agkistrodon contortrix* | 1 | 0 | 0 | 0 |
| *Agkistrodon piscivorus* | 1 | 0 | 0 | 0 |
| *Agkistrodon taylori* | 0 | 1 | 0 | 1 |
| *Ahaetulla fronticincta* | 0 | 1 | 0 | 1 |
| *Ahaetulla nasuta* | 0 | 1 | 0 | 1 |
| *Ahaetulla pulverulenta* | 0 | 1 | 0 | 1 |
| *Ailuronyx seychellensis* | 0 | 1 | 0 | 1 |
| *Ailuronyx tachyscopaeus* | 0 | 1 | 0 | 1 |
| *Ailuronyx trachygaster* | 0 | 1 | 0 | 1 |
| *Aipysurus apraefrontalis* | 0 | 1 | 0 | 1 |
| *Aipysurus duboisii* | 0 | 1 | 0 | 1 |
| *Aipysurus eydouxii* | 0 | 1 | 0 | 1 |
| *Aipysurus fuscus* | 0 | 1 | 0 | 1 |
| *Aipysurus laevis* | 0 | 1 | 0 | 1 |
| *Algyroides fitzingeri* | 1 | 0 | 0 | 0 |
| *Algyroides marchi* | 1 | 0 | 0 | 0 |
| *Algyroides moreoticus* | 1 | 0 | 0 | 0 |
| *Algyroides nigropunctatus* | 1 | 0 | 0 | 0 |
| *Alluaudina bellyi* | 0 | 1 | 0 | 1 |
| *Alopoglossus angulatus* | 0 | 1 | 0 | 1 |
| *Alopoglossus atriventris* | 0 | 1 | 0 | 1 |
| *Alopoglossus copii* | 0 | 1 | 0 | 1 |
| *Alsophis anomalus* | 0 | 1 | 0 | 1 |
| *Alsophis antiguae* | 0 | 1 | 0 | 1 |
| *Alsophis antillensis* | 0 | 1 | 0 | 1 |
| *Alsophis biserialis* | 0 | 1 | 0 | 1 |
| *Alsophis cantherigerus* | 0 | 1 | 0 | 1 |
| *Alsophis elegans* | 0 | 1 | 0 | 1 |
| *Alsophis portoricensis* | 0 | 1 | 0 | 1 |
| *Alsophis rijgersmaei* | 0 | 1 | 0 | 1 |
| *Alsophis rufiventris* | 0 | 1 | 0 | 1 |
| *Alsophis vudii* | 0 | 1 | 0 | 1 |
| *Alsophylax pipiens* | 1 | 0 | 0 | 0 |
| *Amastridium veliferum* | 0 | 1 | 0 | 1 |
| *Amblyodipsas dimidiata* | 0 | 1 | 0 | 1 |
| *Amblyodipsas polylepis* | 0 | 1 | 0 | 1 |
| *Amblyrhynchus cristatus* | 0 | 1 | 0 | 1 |
| *Ameiva ameiva* | 0 | 1 | 0 | 1 |
| *Ameiva auberi* | 0 | 1 | 0 | 1 |
| *Ameiva bifrontata* | 0 | 1 | 0 | 1 |
| *Ameiva chrysolaema* | 0 | 1 | 0 | 1 |
| *Ameiva corax* | 0 | 1 | 0 | 1 |
| *Ameiva dorsalis* | 0 | 1 | 0 | 1 |
| *Ameiva erythrocephala* | 0 | 1 | 0 | 1 |
| *Ameiva exsul* | 0 | 1 | 0 | 1 |
| *Ameiva festiva* | 0 | 1 | 0 | 1 |
| *Ameiva fuscata* | 0 | 1 | 0 | 1 |
| *Ameiva griswoldi* | 0 | 1 | 0 | 1 |
| *Ameiva leberi* | 0 | 1 | 0 | 1 |
| *Ameiva lineolata* | 0 | 1 | 0 | 1 |
| *Ameiva maynardi* | 0 | 1 | 0 | 1 |
| *Ameiva plei* | 0 | 1 | 0 | 1 |
| *Ameiva pluvianotata* | 0 | 1 | 0 | 1 |
| *Ameiva polops* | 0 | 1 | 0 | 1 |
| *Ameiva quadrilineata* | 0 | 1 | 0 | 1 |
| *Ameiva taeniura* | 0 | 1 | 0 | 1 |
| *Ameiva undulata* | 0 | 1 | 0 | 1 |
| *Ameiva wetmorei* | 0 | 1 | 0 | 1 |
| *Amphibolurus muricatus* | 0 | 1 | 0 | 1 |
| *Amphibolurus nobbi* | 0 | 1 | 0 | 1 |
| *Amphibolurus norrisi* | 0 | 1 | 0 | 1 |
| *Amphiesma craspedogaster* | 0 | 1 | 0 | 1 |
| *Amphiesma sauteri* | 0 | 1 | 0 | 1 |
| *Amphiesma stolatum* | 0 | 1 | 0 | 1 |
| *Amphiglossus anosyensis* | 0 | 1 | 0 | 1 |
| *Amphiglossus astrolabi* | 0 | 1 | 0 | 1 |
| *Amphiglossus frontoparietalis* | 0 | 1 | 0 | 1 |
| *Amphiglossus macrocercus* | 0 | 1 | 0 | 1 |
| *Amphiglossus mandokava* | 0 | 1 | 0 | 1 |
| *Amphiglossus melanurus* | 0 | 1 | 0 | 1 |
| *Amphiglossus nanus* | 0 | 1 | 0 | 1 |
| *Amphiglossus ornaticeps* | 0 | 1 | 0 | 1 |
| *Amphiglossus polleni* | 0 | 1 | 0 | 1 |
| *Amphiglossus punctatus* | 0 | 1 | 0 | 1 |
| *Amphiglossus reticulatus* | 0 | 1 | 0 | 1 |
| *Amphiglossus splendidus* | 0 | 1 | 0 | 1 |
| *Amphiglossus tanysoma* | 0 | 1 | 0 | 1 |
| *Amphiglossus tsaratananensis* | 0 | 1 | 0 | 1 |
| *Amphisbaena alba* | 0 | 1 | 0 | 1 |
| *Amphisbaena anaemariae* | 0 | 1 | 0 | 1 |
| *Amphisbaena angustifrons* | 0 | 1 | 0 | 1 |
| *Amphisbaena bakeri* | 0 | 1 | 0 | 1 |
| *Amphisbaena barbouri* | 0 | 1 | 0 | 1 |
| *Amphisbaena bolivica* | 0 | 1 | 0 | 1 |
| *Amphisbaena caeca* | 0 | 1 | 0 | 1 |
| *Amphisbaena camura* | 0 | 1 | 0 | 1 |
| *Amphisbaena carlgansi* | 0 | 1 | 0 | 1 |
| *Amphisbaena cubana* | 0 | 1 | 0 | 1 |
| *Amphisbaena cunhai* | 0 | 1 | 0 | 1 |
| *Amphisbaena darwini* | 0 | 1 | 0 | 1 |
| *Amphisbaena fenestrata* | 0 | 1 | 0 | 1 |
| *Amphisbaena fuliginosa* | 0 | 1 | 0 | 1 |
| *Amphisbaena hastata* | 0 | 1 | 0 | 1 |
| *Amphisbaena hyporissor* | 0 | 1 | 0 | 1 |
| *Amphisbaena ignatiana* | 0 | 1 | 0 | 1 |
| *Amphisbaena innocens* | 0 | 1 | 0 | 1 |
| *Amphisbaena leali* | 0 | 1 | 0 | 1 |
| *Amphisbaena leeseri* | 0 | 1 | 0 | 1 |
| *Amphisbaena manni* | 0 | 1 | 0 | 1 |
| *Amphisbaena mertensii* | 0 | 1 | 0 | 1 |
| *Amphisbaena munoai* | 0 | 1 | 0 | 1 |
| *Amphisbaena schmidti* | 0 | 1 | 0 | 1 |
| *Amphisbaena silvestrii* | 0 | 1 | 0 | 1 |
| *Amphisbaena vermicularis* | 0 | 1 | 0 | 1 |
| *Amphisbaena xera* | 0 | 1 | 0 | 1 |
| *Amplorhinus multimaculatus* | 0 | 1 | 0 | 1 |
| *Anatololacerta anatolica* | 1 | 0 | 0 | 0 |
| *Anatololacerta danfordi* | 1 | 0 | 0 | 0 |
| *Anatololacerta oertzeni* | 1 | 0 | 0 | 0 |
| *Androngo trivittatus* | 0 | 1 | 0 | 1 |
| *Anelytropsis papillosus* | 0 | 1 | 0 | 1 |
| *Anguis fragilis* | 1 | 0 | 0 | 0 |
| *Anilius scytale* | 0 | 1 | 0 | 1 |
| *Anisolepis longicauda* | 0 | 1 | 0 | 1 |
| *Anniella geronimensis* | 1 | 0 | 0 | 0 |
| *Anniella pulchra* | 1 | 0 | 0 | 0 |
| *Anolis acutus* | 0 | 1 | 0 | 1 |
| *Anolis aeneus* | 0 | 1 | 0 | 1 |
| *Anolis aequatorialis* | 0 | 1 | 0 | 1 |
| *Anolis agassizi* | 0 | 1 | 0 | 1 |
| *Anolis ahli* | 0 | 1 | 0 | 1 |
| *Anolis alayoni* | 0 | 1 | 0 | 1 |
| *Anolis alfaroi* | 0 | 1 | 0 | 1 |
| *Anolis aliniger* | 0 | 1 | 0 | 1 |
| *Anolis allisoni* | 0 | 1 | 0 | 1 |
| *Anolis allogus* | 0 | 1 | 0 | 1 |
| *Anolis altae* | 0 | 1 | 0 | 1 |
| *Anolis alumina* | 0 | 1 | 0 | 1 |
| *Anolis alutaceus* | 0 | 1 | 0 | 1 |
| *Anolis anatoloros* | 0 | 1 | 0 | 1 |
| *Anolis angusticeps* | 0 | 1 | 0 | 1 |
| *Anolis annectens* | 0 | 1 | 0 | 1 |
| *Anolis aquaticus* | 0 | 1 | 0 | 1 |
| *Anolis argenteolus* | 0 | 1 | 0 | 1 |
| *Anolis argillaceus* | 0 | 1 | 0 | 1 |
| *Anolis armouri* | 0 | 1 | 0 | 1 |
| *Anolis auratus* | 0 | 1 | 0 | 1 |
| *Anolis bahorucoensis* | 0 | 1 | 0 | 1 |
| *Anolis baleatus* | 0 | 1 | 0 | 1 |
| *Anolis baracoae* | 0 | 1 | 0 | 1 |
| *Anolis barahonae* | 0 | 1 | 0 | 1 |
| *Anolis barbatus* | 0 | 1 | 0 | 1 |
| *Anolis barbouri* | 0 | 1 | 0 | 1 |
| *Anolis bartschi* | 0 | 1 | 0 | 1 |
| *Anolis bicaorum* | 0 | 1 | 0 | 1 |
| *Anolis bimaculatus* | 0 | 1 | 0 | 1 |
| *Anolis biporcatus* | 0 | 1 | 0 | 1 |
| *Anolis bitectus* | 0 | 1 | 0 | 1 |
| *Anolis boettgeri* | 0 | 1 | 0 | 1 |
| *Anolis bombiceps* | 0 | 1 | 0 | 1 |
| *Anolis bonairensis* | 0 | 1 | 0 | 1 |
| *Anolis bremeri* | 0 | 1 | 0 | 1 |
| *Anolis brevirostris* | 0 | 1 | 0 | 1 |
| *Anolis brunneus* | 0 | 1 | 0 | 1 |
| *Anolis calimae* | 0 | 1 | 0 | 1 |
| *Anolis capito* | 0 | 1 | 0 | 1 |
| *Anolis carolinensis* | 1 | 1 | 0 | 0 |
| *Anolis carpenteri* | 0 | 1 | 0 | 1 |
| *Anolis casildae* | 0 | 1 | 0 | 1 |
| *Anolis caudalis* | 0 | 1 | 0 | 1 |
| *Anolis centralis* | 0 | 1 | 0 | 1 |
| *Anolis chamaeleonides* | 0 | 1 | 0 | 1 |
| *Anolis chloris* | 0 | 1 | 0 | 1 |
| *Anolis chlorocyanus* | 0 | 1 | 0 | 1 |
| *Anolis chocorum* | 0 | 1 | 0 | 1 |
| *Anolis christophei* | 0 | 1 | 0 | 1 |
| *Anolis chrysolepis* | 0 | 1 | 0 | 1 |
| *Anolis clivicola* | 0 | 1 | 0 | 1 |
| *Anolis coelestinus* | 0 | 1 | 0 | 1 |
| *Anolis confusus* | 0 | 1 | 0 | 1 |
| *Anolis conspersus* | 0 | 1 | 0 | 1 |
| *Anolis cooki* | 0 | 1 | 0 | 1 |
| *Anolis crassulus* | 0 | 1 | 0 | 1 |
| *Anolis cristatellus* | 0 | 1 | 0 | 1 |
| *Anolis cupeyalensis* | 0 | 1 | 0 | 1 |
| *Anolis cupreus* | 0 | 1 | 0 | 1 |
| *Anolis cuvieri* | 0 | 1 | 0 | 1 |
| *Anolis cyanopleurus* | 0 | 1 | 0 | 1 |
| *Anolis cybotes* | 0 | 1 | 0 | 1 |
| *Anolis danieli* | 0 | 1 | 0 | 1 |
| *Anolis darlingtoni* | 0 | 1 | 0 | 1 |
| *Anolis desechensis* | 0 | 1 | 0 | 1 |
| *Anolis distichus* | 0 | 1 | 0 | 1 |
| *Anolis dolichocephalus* | 0 | 1 | 0 | 1 |
| *Anolis equestris* | 0 | 1 | 0 | 1 |
| *Anolis ernestwilliamsi* | 0 | 1 | 0 | 1 |
| *Anolis etheridgei* | 0 | 1 | 0 | 1 |
| *Anolis eugenegrahami* | 0 | 1 | 0 | 1 |
| *Anolis euskalerriari* | 0 | 1 | 0 | 1 |
| *Anolis evermanni* | 0 | 1 | 0 | 1 |
| *Anolis extremus* | 0 | 1 | 0 | 1 |
| *Anolis ferreus* | 0 | 1 | 0 | 1 |
| *Anolis festae* | 0 | 1 | 0 | 1 |
| *Anolis fitchi* | 0 | 1 | 0 | 1 |
| *Anolis fowleri* | 0 | 1 | 0 | 1 |
| *Anolis fraseri* | 0 | 1 | 0 | 1 |
| *Anolis frenatus* | 0 | 1 | 0 | 1 |
| *Anolis fuscoauratus* | 0 | 1 | 0 | 1 |
| *Anolis garmani* | 0 | 1 | 0 | 1 |
| *Anolis garridoi* | 0 | 1 | 0 | 1 |
| *Anolis gemmosus* | 0 | 1 | 0 | 1 |
| *Anolis gingivinus* | 0 | 1 | 0 | 1 |
| *Anolis grahami* | 0 | 1 | 0 | 1 |
| *Anolis griseus* | 0 | 1 | 0 | 1 |
| *Anolis guafe* | 0 | 1 | 0 | 1 |
| *Anolis guamuhaya* | 0 | 1 | 0 | 1 |
| *Anolis guazuma* | 0 | 1 | 0 | 1 |
| *Anolis gundlachi* | 0 | 1 | 0 | 1 |
| *Anolis haetianus* | 0 | 1 | 0 | 1 |
| *Anolis hendersoni* | 0 | 1 | 0 | 1 |
| *Anolis heterodermus* | 0 | 1 | 0 | 1 |
| *Anolis homolechis* | 0 | 1 | 0 | 1 |
| *Anolis huilae* | 0 | 1 | 0 | 1 |
| *Anolis humilis* | 0 | 1 | 0 | 1 |
| *Anolis imias* | 0 | 1 | 0 | 1 |
| *Anolis inderenae* | 0 | 1 | 0 | 1 |
| *Anolis inexpectatus* | 0 | 1 | 0 | 1 |
| *Anolis insignis* | 0 | 1 | 0 | 1 |
| *Anolis insolitus* | 0 | 1 | 0 | 1 |
| *Anolis intermedius* | 0 | 1 | 0 | 1 |
| *Anolis isolepis* | 0 | 1 | 0 | 1 |
| *Anolis isthmicus* | 0 | 1 | 0 | 1 |
| *Anolis jacare* | 0 | 1 | 0 | 1 |
| *Anolis jubar* | 0 | 1 | 0 | 1 |
| *Anolis kemptoni* | 0 | 1 | 0 | 1 |
| *Anolis krugi* | 0 | 1 | 0 | 1 |
| *Anolis laeviventris* | 0 | 1 | 0 | 1 |
| *Anolis leachii* | 0 | 1 | 0 | 1 |
| *Anolis lemurinus* | 0 | 1 | 0 | 1 |
| *Anolis limifrons* | 0 | 1 | 0 | 1 |
| *Anolis lineatopus* | 0 | 1 | 0 | 1 |
| *Anolis lineatus* | 0 | 1 | 0 | 1 |
| *Anolis lionotus* | 0 | 1 | 0 | 1 |
| *Anolis lividus* | 0 | 1 | 0 | 1 |
| *Anolis longiceps* | 0 | 1 | 0 | 1 |
| *Anolis longitibialis* | 0 | 1 | 0 | 1 |
| *Anolis loveridgei* | 0 | 1 | 0 | 1 |
| *Anolis loysiana* | 0 | 1 | 0 | 1 |
| *Anolis luciae* | 0 | 1 | 0 | 1 |
| *Anolis lucius* | 0 | 1 | 0 | 1 |
| *Anolis luteogularis* | 0 | 1 | 0 | 1 |
| *Anolis macilentus* | 0 | 1 | 0 | 1 |
| *Anolis maculigula* | 0 | 1 | 0 | 1 |
| *Anolis marcanoi* | 0 | 1 | 0 | 1 |
| *Anolis marmoratus* | 0 | 1 | 0 | 1 |
| *Anolis marron* | 0 | 1 | 0 | 1 |
| *Anolis maynardi* | 0 | 1 | 0 | 1 |
| *Anolis meridionalis* | 0 | 1 | 0 | 1 |
| *Anolis mestrei* | 0 | 1 | 0 | 1 |
| *Anolis microtus* | 0 | 1 | 0 | 1 |
| *Anolis monensis* | 0 | 1 | 0 | 1 |
| *Anolis monticola* | 0 | 1 | 0 | 1 |
| *Anolis neblininus* | 0 | 1 | 0 | 1 |
| *Anolis nicefori* | 0 | 1 | 0 | 1 |
| *Anolis nitens* | 0 | 1 | 0 | 1 |
| *Anolis noblei* | 0 | 1 | 0 | 1 |
| *Anolis nubilis* | 0 | 1 | 0 | 1 |
| *Anolis occultus* | 0 | 1 | 0 | 1 |
| *Anolis ocelloscapularis* | 0 | 1 | 0 | 1 |
| *Anolis oculatus* | 0 | 1 | 0 | 1 |
| *Anolis olssoni* | 0 | 1 | 0 | 1 |
| *Anolis onca* | 0 | 1 | 0 | 1 |
| *Anolis opalinus* | 0 | 1 | 0 | 1 |
| *Anolis ophiolepis* | 0 | 1 | 0 | 1 |
| *Anolis oporinus* | 0 | 1 | 0 | 1 |
| *Anolis ortonii* | 0 | 1 | 0 | 1 |
| *Anolis oxylophus* | 0 | 1 | 0 | 1 |
| *Anolis pachypus* | 0 | 1 | 0 | 1 |
| *Anolis paternus* | 0 | 1 | 0 | 1 |
| *Anolis peraccae* | 0 | 1 | 0 | 1 |
| *Anolis placidus* | 0 | 1 | 0 | 1 |
| *Anolis poecilopus* | 0 | 1 | 0 | 1 |
| *Anolis pogus* | 0 | 1 | 0 | 1 |
| *Anolis polylepis* | 0 | 1 | 0 | 1 |
| *Anolis polyrhachis* | 0 | 1 | 0 | 1 |
| *Anolis poncensis* | 0 | 1 | 0 | 1 |
| *Anolis porcatus* | 0 | 1 | 0 | 1 |
| *Anolis porcus* | 0 | 1 | 0 | 1 |
| *Anolis princeps* | 0 | 1 | 0 | 1 |
| *Anolis pulchellus* | 0 | 1 | 0 | 1 |
| *Anolis pumilus* | 0 | 1 | 0 | 1 |
| *Anolis punctatus* | 0 | 1 | 0 | 1 |
| *Anolis purpurgularis* | 0 | 1 | 0 | 1 |
| *Anolis quadriocellifer* | 0 | 1 | 0 | 1 |
| *Anolis quercorum* | 0 | 1 | 0 | 1 |
| *Anolis reconditus* | 0 | 1 | 0 | 1 |
| *Anolis rejectus* | 0 | 1 | 0 | 1 |
| *Anolis richardii* | 0 | 1 | 0 | 1 |
| *Anolis ricordi* | 0 | 1 | 0 | 1 |
| *Anolis roquet* | 0 | 1 | 0 | 1 |
| *Anolis rubribarbaris* | 0 | 1 | 0 | 1 |
| *Anolis sabanus* | 0 | 1 | 0 | 1 |
| *Anolis sagrei* | 0 | 1 | 0 | 1 |
| *Anolis scriptus* | 0 | 1 | 0 | 1 |
| *Anolis semilineatus* | 0 | 1 | 0 | 1 |
| *Anolis sericeus* | 0 | 1 | 0 | 1 |
| *Anolis sheplani* | 0 | 1 | 0 | 1 |
| *Anolis shrevei* | 0 | 1 | 0 | 1 |
| *Anolis singularis* | 0 | 1 | 0 | 1 |
| *Anolis smallwoodi* | 0 | 1 | 0 | 1 |
| *Anolis smaragdinus* | 0 | 1 | 0 | 1 |
| *Anolis sminthus* | 0 | 1 | 0 | 1 |
| *Anolis strahmi* | 0 | 1 | 0 | 1 |
| *Anolis stratulus* | 0 | 1 | 0 | 1 |
| *Anolis terraealtae* | 0 | 1 | 0 | 1 |
| *Anolis tigrinus* | 0 | 1 | 0 | 1 |
| *Anolis trachyderma* | 0 | 1 | 0 | 1 |
| *Anolis transversalis* | 0 | 1 | 0 | 1 |
| *Anolis trinitatis* | 0 | 1 | 0 | 1 |
| *Anolis tropidogaster* | 0 | 1 | 0 | 1 |
| *Anolis tropidonotus* | 0 | 1 | 0 | 1 |
| *Anolis uniformis* | 0 | 1 | 0 | 1 |
| *Anolis utilensis* | 0 | 1 | 0 | 1 |
| *Anolis valencienni* | 0 | 1 | 0 | 1 |
| *Anolis vanidicus* | 0 | 1 | 0 | 1 |
| *Anolis vanzolinii* | 0 | 1 | 0 | 1 |
| *Anolis ventrimaculatus* | 0 | 1 | 0 | 1 |
| *Anolis vermiculatus* | 0 | 1 | 0 | 1 |
| *Anolis wattsi* | 0 | 1 | 0 | 1 |
| *Anolis websteri* | 0 | 1 | 0 | 1 |
| *Anolis whitemani* | 0 | 1 | 0 | 1 |
| *Anolis woodi* | 0 | 1 | 0 | 1 |
| *Anolis zeus* | 0 | 1 | 0 | 1 |
| *Anomalopus leuckartii* | 0 | 1 | 0 | 1 |
| *Anomalopus mackayi* | 0 | 1 | 0 | 1 |
| *Anomalopus swansoni* | 0 | 1 | 0 | 1 |
| *Anomalopus verreauxi* | 0 | 1 | 0 | 1 |
| *Anomochilus leonardi* | 0 | 1 | 0 | 1 |
| *Anops kingii* | 0 | 1 | 0 | 1 |
| *Anotosaura collaris* | 0 | 1 | 0 | 1 |
| *Antaresia childreni* | 0 | 1 | 0 | 1 |
| *Antaresia maculosa* | 0 | 1 | 0 | 1 |
| *Antaresia perthensis* | 0 | 1 | 0 | 1 |
| *Antaresia stimsoni* | 0 | 1 | 0 | 1 |
| *Antillophis andreae* | 0 | 1 | 0 | 1 |
| *Antillophis parvifrons* | 0 | 1 | 0 | 1 |
| *Aparallactus capensis* | 0 | 1 | 0 | 1 |
| *Aparallactus guentheri* | 0 | 1 | 0 | 1 |
| *Aparallactus modestus* | 0 | 1 | 0 | 1 |
| *Aparallactus werneri* | 0 | 1 | 0 | 1 |
| *Apathya cappadocica* | 1 | 0 | 0 | 0 |
| *Aphaniotis fusca* | 0 | 1 | 0 | 1 |
| *Aplopeltura boa* | 0 | 1 | 0 | 1 |
| *Apodora papuana* | 0 | 1 | 0 | 1 |
| *Apostolepis albicollaris* | 0 | 1 | 0 | 1 |
| *Apostolepis assimilis* | 0 | 1 | 0 | 1 |
| *Apostolepis cearensis* | 0 | 1 | 0 | 1 |
| *Apostolepis dimidiata* | 0 | 1 | 0 | 1 |
| *Apostolepis flavotorquata* | 0 | 1 | 0 | 1 |
| *Apostolepis sanctaeritae* | 0 | 1 | 0 | 1 |
| *Aprasia aurita* | 0 | 1 | 0 | 1 |
| *Aprasia fusca* | 0 | 1 | 0 | 1 |
| *Aprasia inaurita* | 0 | 1 | 0 | 1 |
| *Aprasia parapulchella* | 0 | 1 | 0 | 1 |
| *Aprasia picturata* | 0 | 1 | 0 | 1 |
| *Aprasia pseudopulchella* | 0 | 1 | 0 | 1 |
| *Aprasia pulchella* | 0 | 1 | 0 | 1 |
| *Aprasia repens* | 0 | 1 | 0 | 1 |
| *Aprasia smithi* | 0 | 1 | 0 | 1 |
| *Aprasia striolata* | 0 | 1 | 0 | 1 |
| *Apterygodon vittatus* | 0 | 1 | 0 | 1 |
| *Archaeolacerta bedriagae* | 1 | 0 | 0 | 0 |
| *Aristelliger georgeensis* | 0 | 1 | 0 | 1 |
| *Aristelliger lar* | 0 | 1 | 0 | 1 |
| *Aristelliger praesignis* | 0 | 1 | 0 | 1 |
| *Arizona elegans* | 1 | 0 | 0 | 0 |
| *Arrhyton callilaemum* | 0 | 1 | 0 | 1 |
| *Arrhyton dolichura* | 0 | 1 | 0 | 1 |
| *Arrhyton exiguum* | 0 | 1 | 0 | 1 |
| *Arrhyton funereum* | 0 | 1 | 0 | 1 |
| *Arrhyton landoi* | 0 | 1 | 0 | 1 |
| *Arrhyton polylepis* | 0 | 1 | 0 | 1 |
| *Arrhyton procerum* | 0 | 1 | 0 | 1 |
| *Arrhyton supernum* | 0 | 1 | 0 | 1 |
| *Arrhyton taeniatum* | 0 | 1 | 0 | 1 |
| *Arrhyton tanyplectum* | 0 | 1 | 0 | 1 |
| *Arrhyton vittatum* | 0 | 1 | 0 | 1 |
| *Arthrosaura kockii* | 0 | 1 | 0 | 1 |
| *Arthrosaura reticulata* | 0 | 1 | 0 | 1 |
| *Asaccus platyrhynchus* | 1 | 0 | 0 | 0 |
| *Aspidelaps scutatus* | 0 | 1 | 0 | 1 |
| *Aspidites melanocephalus* | 0 | 1 | 0 | 1 |
| *Aspidites ramsayi* | 0 | 1 | 0 | 1 |
| *Aspidomorphus lineaticollis* | 0 | 1 | 0 | 1 |
| *Aspidomorphus muelleri* | 0 | 1 | 0 | 1 |
| *Aspidomorphus schlegeli* | 0 | 1 | 0 | 1 |
| *Aspidoscelis burti* | 1 | 0 | 0 | 0 |
| *Aspidoscelis ceralbensis* | 1 | 0 | 0 | 0 |
| *Aspidoscelis communis* | 0 | 1 | 0 | 1 |
| *Aspidoscelis costatus* | 0 | 1 | 0 | 1 |
| *Aspidoscelis deppei* | 0 | 1 | 0 | 1 |
| *Aspidoscelis gularis* | 1 | 0 | 0 | 0 |
| *Aspidoscelis guttatus* | 0 | 1 | 0 | 1 |
| *Aspidoscelis hyperythrus* | 1 | 0 | 0 | 0 |
| *Aspidoscelis inornatus* | 1 | 0 | 0 | 0 |
| *Aspidoscelis laredoensis* | 1 | 0 | 0 | 0 |
| *Aspidoscelis lineattissimus* | 0 | 1 | 0 | 1 |
| *Aspidoscelis marmoratus* | 1 | 0 | 0 | 0 |
| *Aspidoscelis sexlineatus* | 1 | 0 | 0 | 0 |
| *Aspidoscelis tigris* | 1 | 0 | 0 | 0 |
| *Aspidoscelis velox* | 1 | 0 | 0 | 0 |
| *Aspidura drummondhayi* | 0 | 1 | 0 | 1 |
| *Aspidura guentheri* | 0 | 1 | 0 | 1 |
| *Aspidura trachyprocta* | 0 | 1 | 0 | 1 |
| *Asthenodipsas vertebralis* | 0 | 1 | 0 | 1 |
| *Astrotia stokesii* | 0 | 1 | 0 | 1 |
| *Asymblepharus alaicus* | 1 | 0 | 0 | 0 |
| *Asymblepharus sikimmensis* | 0 | 1 | 0 | 1 |
| *Ateuchosaurus pellopleurus* | 0 | 1 | 0 | 1 |
| *Atheris barbouri* | 0 | 1 | 0 | 1 |
| *Atheris ceratophora* | 0 | 1 | 0 | 1 |
| *Atheris chlorechis* | 0 | 1 | 0 | 1 |
| *Atheris desaixi* | 0 | 1 | 0 | 1 |
| *Atheris hispida* | 0 | 1 | 0 | 1 |
| *Atheris nitschei* | 0 | 1 | 0 | 1 |
| *Atheris squamigera* | 0 | 1 | 0 | 1 |
| *Atlantolacerta andreanskyi* | 1 | 0 | 0 | 0 |
| *Atractaspis bibronii* | 0 | 1 | 0 | 1 |
| *Atractaspis boulengeri* | 0 | 1 | 0 | 1 |
| *Atractaspis corpulenta* | 0 | 1 | 0 | 1 |
| *Atractaspis irregularis* | 0 | 1 | 0 | 1 |
| *Atractaspis microlepidota* | 0 | 1 | 0 | 1 |
| *Atractaspis micropholis* | 0 | 1 | 0 | 1 |
| *Atractus albuquerquei* | 0 | 1 | 0 | 1 |
| *Atractus badius* | 0 | 1 | 0 | 1 |
| *Atractus elaps* | 0 | 1 | 0 | 1 |
| *Atractus flammigerus* | 0 | 1 | 0 | 1 |
| *Atractus reticulatus* | 0 | 1 | 0 | 1 |
| *Atractus schach* | 0 | 1 | 0 | 1 |
| *Atractus trihedrurus* | 0 | 1 | 0 | 1 |
| *Atractus wagleri* | 0 | 1 | 0 | 1 |
| *Atractus zebrinus* | 0 | 1 | 0 | 1 |
| *Atractus zidoki* | 0 | 1 | 0 | 1 |
| *Atretium schistosum* | 0 | 1 | 0 | 1 |
| *Atretium yunnanensis* | 0 | 1 | 0 | 1 |
| *Atropoides nummifer* | 0 | 1 | 0 | 1 |
| *Atropoides occiduus* | 0 | 1 | 0 | 1 |
| *Atropoides olmec* | 0 | 1 | 0 | 1 |
| *Atropoides picadoi* | 0 | 1 | 0 | 1 |
| *Aulura anomala* | 0 | 1 | 0 | 1 |
| *Australolacerta australis* | 0 | 1 | 0 | 1 |
| *Austrelaps labialis* | 0 | 1 | 0 | 1 |
| *Austrelaps superbus* | 0 | 1 | 0 | 1 |
| *Austrotyphlops ammodytes* | 0 | 1 | 0 | 1 |
| *Austrotyphlops australis* | 0 | 1 | 0 | 1 |
| *Austrotyphlops bituberculatus* | 0 | 1 | 0 | 1 |
| *Austrotyphlops diversus* | 0 | 1 | 0 | 1 |
| *Austrotyphlops endoterus* | 0 | 1 | 0 | 1 |
| *Austrotyphlops ganei* | 0 | 1 | 0 | 1 |
| *Austrotyphlops grypus* | 0 | 1 | 0 | 1 |
| *Austrotyphlops guentheri* | 0 | 1 | 0 | 1 |
| *Austrotyphlops hamatus* | 0 | 1 | 0 | 1 |
| *Austrotyphlops howi* | 0 | 1 | 0 | 1 |
| *Austrotyphlops kimberleyensis* | 0 | 1 | 0 | 1 |
| *Austrotyphlops leptosomus* | 0 | 1 | 0 | 1 |
| *Austrotyphlops ligatus* | 0 | 1 | 0 | 1 |
| *Austrotyphlops longissimus* | 0 | 1 | 0 | 1 |
| *Austrotyphlops pilbarensis* | 0 | 1 | 0 | 1 |
| *Austrotyphlops pinguis* | 0 | 1 | 0 | 1 |
| *Austrotyphlops splendidus* | 0 | 1 | 0 | 1 |
| *Austrotyphlops troglodytes* | 0 | 1 | 0 | 1 |
| *Austrotyphlops unguirostris* | 0 | 1 | 0 | 1 |
| *Austrotyphlops waitii* | 0 | 1 | 0 | 1 |
| *Azemiops feae* | 0 | 1 | 0 | 1 |
| *Bachia barbouri* | 0 | 1 | 0 | 1 |
| *Bachia bicolor* | 0 | 1 | 0 | 1 |
| *Bachia bresslaui* | 0 | 1 | 0 | 1 |
| *Bachia dorbignyi* | 0 | 1 | 0 | 1 |
| *Bachia flavescens* | 0 | 1 | 0 | 1 |
| *Bachia heteropa* | 0 | 1 | 0 | 1 |
| *Bachia huallagana* | 0 | 1 | 0 | 1 |
| *Bachia intermedia* | 0 | 1 | 0 | 1 |
| *Bachia panoplia* | 0 | 1 | 0 | 1 |
| *Bachia peruana* | 0 | 1 | 0 | 1 |
| *Bachia scolecoides* | 0 | 1 | 0 | 1 |
| *Bachia trisanale* | 0 | 1 | 0 | 1 |
| *Balanophis ceylonensis* | 0 | 1 | 0 | 1 |
| *Barisia herrerae* | 0 | 1 | 0 | 0 |
| *Barisia imbricata* | 1 | 1 | 0 | 0 |
| *Barisia levicollis* | 1 | 0 | 0 | 0 |
| *Barisia rudicollis* | 0 | 1 | 0 | 0 |
| *Bartleia jigurru* | 0 | 1 | 0 | 1 |
| *Basiliscus basiliscus* | 0 | 1 | 0 | 1 |
| *Basiliscus galeritus* | 0 | 1 | 0 | 1 |
| *Basiliscus plumifrons* | 0 | 1 | 0 | 1 |
| *Basiliscus vittatus* | 0 | 1 | 0 | 1 |
| *Bassiana duperreyi* | 0 | 1 | 0 | 1 |
| *Bassiana trilineata* | 0 | 1 | 0 | 1 |
| *Bavayia cyclura* | 0 | 1 | 0 | 1 |
| *Bavayia geitaina* | 0 | 1 | 0 | 1 |
| *Bavayia goroensis* | 0 | 1 | 0 | 1 |
| *Bavayia madjo* | 0 | 1 | 0 | 1 |
| *Bavayia montana* | 0 | 1 | 0 | 1 |
| *Bavayia ornata* | 0 | 1 | 0 | 1 |
| *Bavayia pulchella* | 0 | 1 | 0 | 1 |
| *Bavayia sauvagii* | 0 | 1 | 0 | 1 |
| *Bellatorias frerei* | 0 | 1 | 0 | 1 |
| *Bellatorias major* | 0 | 1 | 0 | 1 |
| *Bibilava epistibes* | 0 | 1 | 0 | 1 |
| *Bibilava infrasignatus* | 0 | 1 | 0 | 1 |
| *Bibilava lateralis* | 0 | 1 | 0 | 1 |
| *Bibilava martae* | 0 | 1 | 0 | 1 |
| *Bibilava stumpffi* | 0 | 1 | 0 | 1 |
| *Bipes biporus* | 1 | 1 | 0 | 0 |
| *Bipes canaliculatus* | 0 | 1 | 0 | 1 |
| *Bipes tridactylus* | 0 | 1 | 0 | 1 |
| *Bitia hydroides* | 0 | 1 | 0 | 1 |
| *Bitis arietans* | 0 | 1 | 0 | 1 |
| *Bitis atropos* | 0 | 1 | 0 | 1 |
| *Bitis caudalis* | 0 | 1 | 0 | 1 |
| *Bitis cornuta* | 0 | 1 | 0 | 1 |
| *Bitis gabonica* | 0 | 1 | 0 | 1 |
| *Bitis nasicornis* | 0 | 1 | 0 | 1 |
| *Bitis peringueyi* | 0 | 1 | 0 | 1 |
| *Bitis rubida* | 0 | 1 | 0 | 1 |
| *Bitis worthingtoni* | 0 | 1 | 0 | 1 |
| *Bitis xeropaga* | 0 | 1 | 0 | 1 |
| *Blaesodactylus antongilensis* | 0 | 1 | 0 | 1 |
| *Blaesodactylus boivini* | 0 | 1 | 0 | 1 |
| *Blaesodactylus sakalava* | 0 | 1 | 0 | 1 |
| *Blanus cinereus* | 1 | 0 | 0 | 0 |
| *Blanus mettetali* | 1 | 0 | 0 | 0 |
| *Blanus strauchi* | 1 | 0 | 0 | 0 |
| *Blanus tingitanus* | 1 | 0 | 0 | 0 |
| *Boa constrictor* | 0 | 1 | 0 | 1 |
| *Bogertia lutzae* | 0 | 1 | 0 | 1 |
| *Bogertophis rosaliae* | 1 | 0 | 0 | 0 |
| *Bogertophis subocularis* | 1 | 0 | 0 | 0 |
| *Boiga barnesii* | 0 | 1 | 0 | 1 |
| *Boiga beddomei* | 0 | 1 | 0 | 1 |
| *Boiga ceylonensis* | 0 | 1 | 0 | 1 |
| *Boiga cynodon* | 0 | 1 | 0 | 1 |
| *Boiga dendrophila* | 0 | 1 | 0 | 1 |
| *Boiga forsteni* | 0 | 1 | 0 | 1 |
| *Boiga irregularis* | 0 | 1 | 0 | 1 |
| *Boiga kraepelini* | 0 | 1 | 0 | 1 |
| *Boiga multomaculata* | 0 | 1 | 0 | 1 |
| *Boiga pulverulenta* | 0 | 1 | 0 | 1 |
| *Boiga trigonata* | 0 | 1 | 0 | 1 |
| *Boiruna maculata* | 0 | 1 | 0 | 1 |
| *Bothriechis aurifer* | 0 | 1 | 0 | 1 |
| *Bothriechis bicolor* | 0 | 1 | 0 | 1 |
| *Bothriechis lateralis* | 0 | 1 | 0 | 1 |
| *Bothriechis marchi* | 0 | 1 | 0 | 1 |
| *Bothriechis nigroviridis* | 0 | 1 | 0 | 1 |
| *Bothriechis rowleyi* | 0 | 1 | 0 | 1 |
| *Bothriechis schlegelii* | 0 | 1 | 0 | 1 |
| *Bothriechis thalassinus* | 0 | 1 | 0 | 1 |
| *Bothriopsis bilineata* | 0 | 1 | 0 | 1 |
| *Bothriopsis chloromelas* | 0 | 1 | 0 | 1 |
| *Bothriopsis pulchra* | 0 | 1 | 0 | 1 |
| *Bothriopsis taeniata* | 0 | 1 | 0 | 1 |
| *Bothrochilus boa* | 0 | 1 | 0 | 1 |
| *Bothrocophias campbelli* | 0 | 1 | 0 | 1 |
| *Bothrocophias hyoprora* | 0 | 1 | 0 | 1 |
| *Bothrocophias microphthalmus* | 0 | 1 | 0 | 1 |
| *Bothrolycus ater* | 0 | 1 | 0 | 1 |
| *Bothrophthalmus brunneus* | 0 | 1 | 0 | 1 |
| *Bothrophthalmus lineatus* | 0 | 1 | 0 | 1 |
| *Bothrops alcatraz* | 0 | 1 | 0 | 1 |
| *Bothrops alternatus* | 0 | 1 | 0 | 1 |
| *Bothrops ammodytoides* | 0 | 0 | 1 | 0 |
| *Bothrops asper* | 0 | 1 | 0 | 1 |
| *Bothrops atrox* | 0 | 1 | 0 | 1 |
| *Bothrops brazili* | 0 | 1 | 0 | 1 |
| *Bothrops caribbaeus* | 0 | 1 | 0 | 1 |
| *Bothrops colombiensis* | 0 | 1 | 0 | 1 |
| *Bothrops cotiara* | 0 | 1 | 0 | 1 |
| *Bothrops diporus* | 0 | 1 | 0 | 1 |
| *Bothrops erythromelas* | 0 | 1 | 0 | 1 |
| *Bothrops fonsecai* | 0 | 1 | 0 | 1 |
| *Bothrops insularis* | 0 | 1 | 0 | 1 |
| *Bothrops itapetiningae* | 0 | 1 | 0 | 1 |
| *Bothrops jararaca* | 0 | 1 | 0 | 1 |
| *Bothrops jararacussu* | 0 | 1 | 0 | 1 |
| *Bothrops lanceolatus* | 0 | 1 | 0 | 1 |
| *Bothrops leucurus* | 0 | 1 | 0 | 1 |
| *Bothrops marajoensis* | 0 | 1 | 0 | 1 |
| *Bothrops moojeni* | 0 | 1 | 0 | 1 |
| *Bothrops neuwiedi* | 0 | 1 | 0 | 1 |
| *Bothrops pictus* | 0 | 1 | 0 | 1 |
| *Bothrops punctata* | 0 | 1 | 0 | 1 |
| *Brachylophus fasciatus* | 0 | 1 | 0 | 1 |
| *Brachylophus vitiensis* | 0 | 1 | 0 | 1 |
| *Brachymeles apus* | 0 | 1 | 0 | 1 |
| *Brachymeles bicolor* | 0 | 1 | 0 | 1 |
| *Brachymeles bonitae* | 0 | 1 | 0 | 1 |
| *Brachymeles boulengeri* | 0 | 1 | 0 | 1 |
| *Brachymeles cebuensis* | 0 | 1 | 0 | 1 |
| *Brachymeles elerae* | 0 | 1 | 0 | 1 |
| *Brachymeles gracilis* | 0 | 1 | 0 | 1 |
| *Brachymeles minimus* | 0 | 1 | 0 | 1 |
| *Brachymeles pathfinderi* | 0 | 1 | 0 | 1 |
| *Brachymeles samarensis* | 0 | 1 | 0 | 1 |
| *Brachymeles schadenbergi* | 0 | 1 | 0 | 1 |
| *Brachymeles talinis* | 0 | 1 | 0 | 1 |
| *Brachymeles tridactylus* | 0 | 1 | 0 | 1 |
| *Brachyophidium rhodogaster* | 0 | 1 | 0 | 1 |
| *Bradypodion atromontanum* | 0 | 1 | 0 | 1 |
| *Bradypodion caffrum* | 0 | 1 | 0 | 1 |
| *Bradypodion damaranum* | 0 | 1 | 0 | 1 |
| *Bradypodion dracomontanum* | 0 | 1 | 0 | 1 |
| *Bradypodion gutturale* | 0 | 1 | 0 | 1 |
| *Bradypodion karrooicum* | 0 | 1 | 0 | 1 |
| *Bradypodion melanocephalum* | 0 | 1 | 0 | 1 |
| *Bradypodion nemorale* | 0 | 1 | 0 | 1 |
| *Bradypodion occidentale* | 0 | 1 | 0 | 1 |
| *Bradypodion pumilum* | 0 | 1 | 0 | 1 |
| *Bradypodion setaroi* | 0 | 1 | 0 | 1 |
| *Bradypodion taeniabronchum* | 0 | 1 | 0 | 1 |
| *Bradypodion thamnobates* | 0 | 1 | 0 | 1 |
| *Bradypodion transvaalense* | 0 | 1 | 0 | 1 |
| *Bradypodion ventrale* | 0 | 1 | 0 | 1 |
| *Broghammerus reticulatus* | 0 | 1 | 0 | 1 |
| *Broghammerus timoriensis* | 0 | 1 | 0 | 1 |
| *Bronchocela cristatella* | 0 | 1 | 0 | 1 |
| *Bronia brasiliana* | 0 | 1 | 0 | 1 |
| *Bronia kraoh* | 0 | 1 | 0 | 1 |
| *Bronia saxosa* | 0 | 1 | 0 | 1 |
| *Brookesia ambreensis* | 0 | 1 | 0 | 1 |
| *Brookesia antakarana* | 0 | 1 | 0 | 1 |
| *Brookesia betschi* | 0 | 1 | 0 | 1 |
| *Brookesia bonsi* | 0 | 1 | 0 | 1 |
| *Brookesia brygooi* | 0 | 1 | 0 | 1 |
| *Brookesia decaryi* | 0 | 1 | 0 | 1 |
| *Brookesia dentata* | 0 | 1 | 0 | 1 |
| *Brookesia ebenaui* | 0 | 1 | 0 | 1 |
| *Brookesia exarmata* | 0 | 1 | 0 | 1 |
| *Brookesia griveaudi* | 0 | 1 | 0 | 1 |
| *Brookesia karchei* | 0 | 1 | 0 | 1 |
| *Brookesia lineata* | 0 | 1 | 0 | 1 |
| *Brookesia lolontany* | 0 | 1 | 0 | 1 |
| *Brookesia minima* | 0 | 1 | 0 | 1 |
| *Brookesia nasus* | 0 | 1 | 0 | 1 |
| *Brookesia perarmata* | 0 | 1 | 0 | 1 |
| *Brookesia peyrierasi* | 0 | 1 | 0 | 1 |
| *Brookesia stumpffi* | 0 | 1 | 0 | 1 |
| *Brookesia superciliaris* | 0 | 1 | 0 | 1 |
| *Brookesia therezieni* | 0 | 1 | 0 | 1 |
| *Brookesia thieli* | 0 | 1 | 0 | 1 |
| *Brookesia tuberculata* | 0 | 1 | 0 | 1 |
| *Brookesia vadoni* | 0 | 1 | 0 | 1 |
| *Brookesia valerieae* | 0 | 1 | 0 | 1 |
| *Bufoniceps laungwalaensis* | 0 | 1 | 0 | 1 |
| *Buhoma depressiceps* | 0 | 1 | 0 | 1 |
| *Buhoma procterae* | 0 | 1 | 0 | 1 |
| *Bungarus bungaroides* | 0 | 1 | 0 | 1 |
| *Bungarus caeruleus* | 0 | 1 | 0 | 1 |
| *Bungarus candidus* | 0 | 1 | 0 | 1 |
| *Bungarus ceylonicus* | 0 | 1 | 0 | 1 |
| *Bungarus fasciatus* | 0 | 1 | 0 | 1 |
| *Bungarus flaviceps* | 0 | 1 | 0 | 1 |
| *Bungarus multicinctus* | 0 | 1 | 0 | 1 |
| *Bungarus niger* | 0 | 1 | 0 | 1 |
| *Bungarus sindanus* | 0 | 1 | 0 | 1 |
| *Bunopus crassicauda* | 1 | 0 | 0 | 0 |
| *Bunopus tuberculatus* | 1 | 0 | 0 | 0 |
| *Cacophis squamulosus* | 0 | 1 | 0 | 1 |
| *Cadea blanoides* | 0 | 1 | 0 | 1 |
| *Caimanops amphiboluroides* | 0 | 1 | 0 | 1 |
| *Calabaria reinhardtii* | 0 | 1 | 0 | 1 |
| *Calamaria pavimentata* | 0 | 1 | 0 | 1 |
| *Calamaria yunnanensis* | 0 | 1 | 0 | 1 |
| *Calamodontophis paucidens* | 0 | 1 | 0 | 1 |
| *Caledoniscincus aquilonius* | 0 | 1 | 0 | 1 |
| *Caledoniscincus atropunctatus* | 0 | 1 | 0 | 1 |
| *Caledoniscincus auratus* | 0 | 1 | 0 | 1 |
| *Caledoniscincus austrocaledonicus* | 0 | 1 | 0 | 1 |
| *Caledoniscincus chazeaui* | 0 | 1 | 0 | 1 |
| *Caledoniscincus festivus* | 0 | 1 | 0 | 1 |
| *Caledoniscincus haplorhinus* | 0 | 1 | 0 | 1 |
| *Caledoniscincus orestes* | 0 | 1 | 0 | 1 |
| *Caledoniscincus renevieri* | 0 | 1 | 0 | 1 |
| *Caledoniscincus terma* | 0 | 1 | 0 | 1 |
| *Calliophis bivirgata* | 0 | 1 | 0 | 1 |
| *Calliophis melanurus* | 0 | 1 | 0 | 1 |
| *Callisaurus draconoides* | 1 | 0 | 0 | 0 |
| *Callopistes flavipunctatus* | 0 | 1 | 0 | 1 |
| *Callopistes maculatus* | 0 | 0 | 1 | 0 |
| *Calloselasma rhodostoma* | 0 | 1 | 0 | 1 |
| *Calodactylodes aureus* | 0 | 1 | 0 | 1 |
| *Calodactylodes illingworthorum* | 0 | 1 | 0 | 1 |
| *Calotes calotes* | 0 | 1 | 0 | 1 |
| *Calotes ceylonensis* | 0 | 1 | 0 | 1 |
| *Calotes chincollium* | 0 | 1 | 0 | 1 |
| *Calotes emma* | 0 | 1 | 0 | 1 |
| *Calotes htunwini* | 0 | 1 | 0 | 1 |
| *Calotes irawadi* | 0 | 1 | 0 | 1 |
| *Calotes liocephalus* | 0 | 1 | 0 | 1 |
| *Calotes liolepis* | 0 | 1 | 0 | 1 |
| *Calotes mystaceus* | 0 | 1 | 0 | 1 |
| *Calotes nigrilabris* | 0 | 1 | 0 | 1 |
| *Calotes versicolor* | 1 | 0 | 0 | 0 |
| *Calumma boettgeri* | 0 | 1 | 0 | 1 |
| *Calumma brevicorne* | 0 | 1 | 0 | 1 |
| *Calumma capuroni* | 0 | 1 | 0 | 1 |
| *Calumma crypticum* | 0 | 1 | 0 | 1 |
| *Calumma cucullatum* | 0 | 1 | 0 | 1 |
| *Calumma fallax* | 0 | 1 | 0 | 1 |
| *Calumma furcifer* | 0 | 1 | 0 | 1 |
| *Calumma gallus* | 0 | 1 | 0 | 1 |
| *Calumma gastrotaenia* | 0 | 1 | 0 | 1 |
| *Calumma globifer* | 0 | 1 | 0 | 1 |
| *Calumma guibei* | 0 | 1 | 0 | 1 |
| *Calumma hilleniusi* | 0 | 1 | 0 | 1 |
| *Calumma malthe* | 0 | 1 | 0 | 1 |
| *Calumma nasutum* | 0 | 1 | 0 | 1 |
| *Calumma oshaughnessyi* | 0 | 1 | 0 | 1 |
| *Calumma parsonii* | 0 | 1 | 0 | 1 |
| *Calumma tsaratananense* | 0 | 1 | 0 | 1 |
| *Calyptommatus confusionibus* | 0 | 1 | 0 | 1 |
| *Calyptommatus leiolepis* | 0 | 1 | 0 | 1 |
| *Calyptommatus nicterus* | 0 | 1 | 0 | 1 |
| *Calyptommatus sinebrachiatus* | 0 | 1 | 0 | 1 |
| *Calyptotis lepidorostrum* | 0 | 1 | 0 | 1 |
| *Calyptotis ruficauda* | 0 | 1 | 0 | 1 |
| *Calyptotis scutirostrum* | 0 | 1 | 0 | 1 |
| *Candoia aspera* | 0 | 1 | 0 | 1 |
| *Candoia bibroni* | 0 | 1 | 0 | 1 |
| *Candoia carinata* | 0 | 1 | 0 | 1 |
| *Cantoria violacea* | 0 | 1 | 0 | 1 |
| *Carinatogecko heteropholis* | 1 | 0 | 0 | 0 |
| *Carlia amax* | 0 | 1 | 0 | 1 |
| *Carlia bicarinata* | 0 | 1 | 0 | 1 |
| *Carlia coensis* | 0 | 1 | 0 | 1 |
| *Carlia dogare* | 0 | 1 | 0 | 1 |
| *Carlia fusca* | 0 | 1 | 0 | 1 |
| *Carlia gracilis* | 0 | 1 | 0 | 1 |
| *Carlia jarnoldae* | 0 | 1 | 0 | 1 |
| *Carlia johnstonei* | 0 | 1 | 0 | 1 |
| *Carlia longipes* | 0 | 1 | 0 | 1 |
| *Carlia munda* | 0 | 1 | 0 | 1 |
| *Carlia mundivensis* | 0 | 1 | 0 | 1 |
| *Carlia mysi* | 0 | 1 | 0 | 1 |
| *Carlia pectoralis* | 0 | 1 | 0 | 1 |
| *Carlia rhomboidalis* | 0 | 1 | 0 | 1 |
| *Carlia rostralis* | 0 | 1 | 0 | 1 |
| *Carlia rubrigularis* | 0 | 1 | 0 | 1 |
| *Carlia rufilatus* | 0 | 1 | 0 | 1 |
| *Carlia schmeltzii* | 0 | 1 | 0 | 1 |
| *Carlia scirtetis* | 0 | 1 | 0 | 1 |
| *Carlia storri* | 0 | 1 | 0 | 1 |
| *Carlia tetradactyla* | 0 | 1 | 0 | 1 |
| *Carlia triacantha* | 0 | 1 | 0 | 1 |
| *Carlia vivax* | 0 | 1 | 0 | 1 |
| *Carphodactylus laevis* | 0 | 1 | 0 | 1 |
| *Carphophis amoenus* | 1 | 0 | 0 | 0 |
| *Casarea dussumieri* | 0 | 1 | 0 | 1 |
| *Causus defilippii* | 0 | 1 | 0 | 1 |
| *Causus resimus* | 0 | 1 | 0 | 1 |
| *Causus rhombeatus* | 0 | 1 | 0 | 1 |
| *Cautula zia* | 0 | 1 | 0 | 1 |
| *Celatiscincus euryotis* | 0 | 1 | 0 | 1 |
| *Celatiscincus similis* | 0 | 1 | 0 | 1 |
| *Celestus agasepsoides* | 0 | 1 | 0 | 1 |
| *Celestus enneagrammus* | 0 | 1 | 0 | 1 |
| *Celestus haetianus* | 0 | 1 | 0 | 1 |
| *Cemophora coccinea* | 1 | 0 | 0 | 0 |
| *Cerastes cerastes* | 1 | 0 | 0 | 0 |
| *Cerastes gasperettii* | 1 | 0 | 0 | 0 |
| *Cerastes vipera* | 1 | 0 | 0 | 0 |
| *Ceratophora aspera* | 0 | 1 | 0 | 1 |
| *Ceratophora erdeleni* | 0 | 1 | 0 | 1 |
| *Ceratophora karu* | 0 | 1 | 0 | 1 |
| *Ceratophora stoddartii* | 0 | 1 | 0 | 1 |
| *Cerberus australis* | 0 | 1 | 0 | 1 |
| *Cerberus microlepis* | 0 | 1 | 0 | 1 |
| *Cerberus rynchops* | 0 | 1 | 0 | 1 |
| *Cercaspis carinatus* | 0 | 1 | 0 | 1 |
| *Cercolophia cuiabana* | 0 | 1 | 0 | 1 |
| *Cercolophia roberti* | 0 | 1 | 0 | 1 |
| *Cercosaura argulus* | 0 | 1 | 0 | 1 |
| *Cercosaura eigenmanni* | 0 | 1 | 0 | 1 |
| *Cercosaura ocellata* | 0 | 1 | 0 | 1 |
| *Cercosaura oshaughnessyi* | 0 | 1 | 0 | 1 |
| *Cercosaura quadrilineata* | 0 | 1 | 0 | 1 |
| *Cercosaura schreibersii* | 0 | 1 | 0 | 1 |
| *Cerrophidion barbouri* | 0 | 1 | 0 | 1 |
| *Cerrophidion godmani* | 0 | 1 | 0 | 1 |
| *Cerrophidion petlalcalensis* | 0 | 1 | 0 | 1 |
| *Cerrophidion tzotzilorum* | 0 | 1 | 0 | 1 |
| *Chalarodon madagascariensis* | 0 | 1 | 0 | 1 |
| *Chalcides bedriagai* | 1 | 0 | 0 | 0 |
| *Chalcides boulengeri* | 1 | 0 | 0 | 0 |
| *Chalcides chalcides* | 1 | 0 | 0 | 0 |
| *Chalcides coeruleopunctatus* | 1 | 0 | 0 | 0 |
| *Chalcides colosii* | 1 | 0 | 0 | 0 |
| *Chalcides guentheri* | 1 | 0 | 0 | 0 |
| *Chalcides lanzai* | 1 | 0 | 0 | 0 |
| *Chalcides manueli* | 1 | 0 | 0 | 0 |
| *Chalcides mauritanicus* | 1 | 0 | 0 | 0 |
| *Chalcides minutus* | 1 | 0 | 0 | 0 |
| *Chalcides mionecton* | 1 | 0 | 0 | 0 |
| *Chalcides montanus* | 1 | 0 | 0 | 0 |
| *Chalcides ocellatus* | 1 | 0 | 0 | 0 |
| *Chalcides parallelus* | 1 | 0 | 0 | 0 |
| *Chalcides polylepis* | 1 | 0 | 0 | 0 |
| *Chalcides pseudostriatus* | 1 | 0 | 0 | 0 |
| *Chalcides sepsoides* | 1 | 0 | 0 | 0 |
| *Chalcides sexlineatus* | 1 | 0 | 0 | 0 |
| *Chalcides sphenopsiformis* | 1 | 0 | 0 | 0 |
| *Chalcides striatus* | 1 | 0 | 0 | 0 |
| *Chalcides viridanus* | 1 | 0 | 0 | 0 |
| *Chamaeleo affinis* | 0 | 1 | 0 | 1 |
| *Chamaeleo africanus* | 0 | 1 | 0 | 1 |
| *Chamaeleo arabicus* | 0 | 1 | 0 | 1 |
| *Chamaeleo balebicornutus* | 0 | 1 | 0 | 1 |
| *Chamaeleo bitaeniatus* | 0 | 1 | 0 | 1 |
| *Chamaeleo calcaricarens* | 0 | 1 | 0 | 1 |
| *Chamaeleo calyptratus* | 0 | 1 | 0 | 1 |
| *Chamaeleo chamaeleon* | 1 | 0 | 0 | 0 |
| *Chamaeleo cristatus* | 0 | 1 | 0 | 1 |
| *Chamaeleo deremensis* | 0 | 1 | 0 | 1 |
| *Chamaeleo dilepis* | 0 | 1 | 0 | 1 |
| *Chamaeleo ellioti* | 0 | 1 | 0 | 1 |
| *Chamaeleo feae* | 0 | 1 | 0 | 1 |
| *Chamaeleo fuelleborni* | 0 | 1 | 0 | 1 |
| *Chamaeleo goetzei* | 0 | 1 | 0 | 1 |
| *Chamaeleo gracilis* | 0 | 1 | 0 | 1 |
| *Chamaeleo harennae* | 0 | 1 | 0 | 1 |
| *Chamaeleo hoehnelii* | 0 | 1 | 0 | 1 |
| *Chamaeleo jacksonii* | 0 | 1 | 0 | 1 |
| *Chamaeleo johnstoni* | 0 | 1 | 0 | 1 |
| *Chamaeleo laevigatus* | 0 | 1 | 0 | 1 |
| *Chamaeleo melleri* | 0 | 1 | 0 | 1 |
| *Chamaeleo monachus* | 0 | 1 | 0 | 1 |
| *Chamaeleo montium* | 0 | 1 | 0 | 1 |
| *Chamaeleo namaquensis* | 0 | 1 | 0 | 1 |
| *Chamaeleo narraioca* | 0 | 1 | 0 | 1 |
| *Chamaeleo necasi* | 0 | 1 | 0 | 1 |
| *Chamaeleo oweni* | 0 | 1 | 0 | 1 |
| *Chamaeleo pfefferi* | 0 | 1 | 0 | 1 |
| *Chamaeleo quadricornis* | 0 | 1 | 0 | 1 |
| *Chamaeleo quilensis* | 0 | 1 | 0 | 1 |
| *Chamaeleo roperi* | 0 | 1 | 0 | 1 |
| *Chamaeleo rudis* | 0 | 1 | 0 | 1 |
| *Chamaeleo schubotzi* | 0 | 1 | 0 | 1 |
| *Chamaeleo senegalensis* | 0 | 1 | 0 | 1 |
| *Chamaeleo sternfeldi* | 0 | 1 | 0 | 1 |
| *Chamaeleo tempeli* | 0 | 1 | 0 | 1 |
| *Chamaeleo werneri* | 0 | 1 | 0 | 1 |
| *Chamaeleo wiedersheimi* | 0 | 1 | 0 | 1 |
| *Chamaeleo zeylanicus* | 0 | 1 | 0 | 1 |
| *Chamaesaura aenea* | 0 | 1 | 0 | 1 |
| *Chamaesaura anguina* | 0 | 1 | 0 | 1 |
| *Charina bottae* | 1 | 0 | 0 | 0 |
| *Chelosania brunnea* | 0 | 1 | 0 | 1 |
| *Chilomeniscus stramineus* | 1 | 0 | 0 | 0 |
| *Chionactis occipitalis* | 1 | 0 | 0 | 0 |
| *Chioninia delalandii* | 0 | 1 | 0 | 1 |
| *Chioninia fogoensis* | 0 | 1 | 0 | 1 |
| *Chioninia spinalis* | 0 | 1 | 0 | 1 |
| *Chioninia stangeri* | 0 | 1 | 0 | 1 |
| *Chioninia vaillantii* | 0 | 1 | 0 | 1 |
| *Chirindia swynnertoni* | 0 | 1 | 0 | 1 |
| *Chironius bicarinatus* | 0 | 1 | 0 | 1 |
| *Chironius carinatus* | 0 | 1 | 0 | 1 |
| *Chironius exoletus* | 0 | 1 | 0 | 1 |
| *Chironius flavolineatus* | 0 | 1 | 0 | 1 |
| *Chironius fuscus* | 0 | 1 | 0 | 1 |
| *Chironius grandisquamis* | 0 | 1 | 0 | 1 |
| *Chironius laevicollis* | 0 | 1 | 0 | 1 |
| *Chironius laurenti* | 0 | 1 | 0 | 1 |
| *Chironius monticola* | 0 | 1 | 0 | 1 |
| *Chironius multiventris* | 0 | 1 | 0 | 1 |
| *Chironius quadricarinatus* | 0 | 1 | 0 | 1 |
| *Chironius scurrulus* | 0 | 1 | 0 | 1 |
| *Chlamydosaurus kingii* | 0 | 1 | 0 | 1 |
| *Chondrodactylus angulifer* | 0 | 1 | 0 | 1 |
| *Chondrodactylus bibronii* | 0 | 1 | 0 | 1 |
| *Chondrodactylus fitzsimonsi* | 0 | 1 | 0 | 1 |
| *Chondrodactylus turneri* | 0 | 1 | 0 | 1 |
| *Christinus marmoratus* | 0 | 1 | 0 | 1 |
| *Chrysopelea ornata* | 0 | 1 | 0 | 1 |
| *Chrysopelea paradisi* | 0 | 1 | 0 | 1 |
| *Chrysopelea taprobanica* | 0 | 1 | 0 | 1 |
| *Clelia bicolor* | 0 | 1 | 0 | 1 |
| *Clelia clelia* | 0 | 1 | 0 | 1 |
| *Clelia rustica* | 0 | 1 | 0 | 1 |
| *Clonophis kirtlandii* | 1 | 0 | 0 | 0 |
| *Cnemaspis africana* | 0 | 1 | 0 | 1 |
| *Cnemaspis dickersoni* | 0 | 1 | 0 | 1 |
| *Cnemaspis kandiana* | 0 | 1 | 0 | 1 |
| *Cnemaspis kendallii* | 0 | 1 | 0 | 1 |
| *Cnemaspis limi* | 0 | 1 | 0 | 1 |
| *Cnemaspis podihuna* | 0 | 1 | 0 | 1 |
| *Cnemaspis tropidogaster* | 0 | 1 | 0 | 1 |
| *Cnemaspis uzungwae* | 0 | 1 | 0 | 1 |
| *Cnemidophorus arenivagus* | 0 | 1 | 0 | 1 |
| *Cnemidophorus gramivagus* | 0 | 1 | 0 | 1 |
| *Cnemidophorus lacertoides* | 0 | 1 | 0 | 1 |
| *Cnemidophorus lemniscatus* | 0 | 1 | 0 | 1 |
| *Cnemidophorus longicaudus* | 0 | 0 | 1 | 0 |
| *Cnemidophorus ocellifer* | 0 | 1 | 0 | 1 |
| *Cnemidophorus vanzoi* | 0 | 1 | 0 | 1 |
| *Coelognathus erythrurus* | 0 | 1 | 0 | 1 |
| *Coelognathus flavolineatus* | 0 | 1 | 0 | 1 |
| *Coelognathus helena* | 0 | 1 | 0 | 1 |
| *Coelognathus radiata* | 0 | 1 | 0 | 1 |
| *Coelognathus subradiata* | 0 | 1 | 0 | 1 |
| *Coeranoscincus frontalis* | 0 | 1 | 0 | 1 |
| *Coeranoscincus reticulatus* | 0 | 1 | 0 | 1 |
| *Coggeria naufragus* | 0 | 1 | 0 | 1 |
| *Coleodactylus amazonicus* | 0 | 1 | 0 | 1 |
| *Coleodactylus brachystoma* | 0 | 1 | 0 | 1 |
| *Coleodactylus meridionalis* | 0 | 1 | 0 | 1 |
| *Coleodactylus natalensis* | 0 | 1 | 0 | 1 |
| *Coleodactylus septentrionalis* | 0 | 1 | 0 | 1 |
| *Coleonyx brevis* | 1 | 0 | 0 | 0 |
| *Coleonyx elegans* | 0 | 1 | 0 | 1 |
| *Coleonyx mitratus* | 0 | 1 | 0 | 1 |
| *Coleonyx variegatus* | 1 | 0 | 0 | 0 |
| *Colobodactylus dalcyanus* | 0 | 1 | 0 | 1 |
| *Colobodactylus taunayi* | 0 | 1 | 0 | 1 |
| *Colobosaura mentalis* | 0 | 1 | 0 | 1 |
| *Colobosaura modesta* | 0 | 1 | 0 | 1 |
| *Colobosauroides cearensis* | 0 | 1 | 0 | 1 |
| *Coloptychon rhombifer* | 0 | 1 | 0 | 1 |
| *Colopus kochii* | 0 | 1 | 0 | 1 |
| *Colopus wahlbergii* | 0 | 1 | 0 | 1 |
| *Coluber constrictor* | 1 | 0 | 0 | 0 |
| *Coluber dorri* | 0 | 1 | 0 | 1 |
| *Coluber zebrinus* | 0 | 1 | 0 | 1 |
| *Compsophis albiventris* | 0 | 1 | 0 | 1 |
| *Compsophis boulengeri* | 0 | 1 | 0 | 1 |
| *Compsophis infralineatus* | 0 | 1 | 0 | 1 |
| *Compsophis laphystius* | 0 | 1 | 0 | 1 |
| *Coniophanes fissidens* | 0 | 1 | 0 | 1 |
| *Conolophus pallidus* | 0 | 1 | 0 | 1 |
| *Conolophus subcristatus* | 0 | 1 | 0 | 1 |
| *Conophis lineatus* | 0 | 1 | 0 | 1 |
| *Conopsis biserialis* | 0 | 1 | 0 | 1 |
| *Conopsis nasus* | 0 | 1 | 0 | 1 |
| *Contia tenuis* | 1 | 0 | 0 | 0 |
| *Cophosaurus texanus* | 1 | 0 | 0 | 0 |
| *Cophotis ceylanica* | 0 | 1 | 0 | 1 |
| *Cophotis dumbara* | 0 | 1 | 0 | 1 |
| *Corallus annulatus* | 0 | 1 | 0 | 1 |
| *Corallus caninus* | 0 | 1 | 0 | 1 |
| *Corallus hortulanus* | 0 | 1 | 0 | 1 |
| *Cordylosaurus subtessellatus* | 0 | 1 | 0 | 1 |
| *Cordylus aridus* | 0 | 1 | 0 | 1 |
| *Cordylus beraduccii* | 0 | 1 | 0 | 1 |
| *Cordylus campbelli* | 0 | 1 | 0 | 1 |
| *Cordylus capensis* | 0 | 1 | 0 | 1 |
| *Cordylus cataphractus* | 0 | 1 | 0 | 1 |
| *Cordylus coeruleopunctatus* | 0 | 1 | 0 | 1 |
| *Cordylus cordylus* | 0 | 1 | 0 | 1 |
| *Cordylus giganteus* | 0 | 1 | 0 | 1 |
| *Cordylus imkeae* | 0 | 1 | 0 | 1 |
| *Cordylus jonesii* | 0 | 1 | 0 | 1 |
| *Cordylus jordani* | 0 | 1 | 0 | 1 |
| *Cordylus langi* | 0 | 1 | 0 | 1 |
| *Cordylus lawrenci* | 0 | 1 | 0 | 1 |
| *Cordylus macropholis* | 0 | 1 | 0 | 1 |
| *Cordylus mclachlani* | 0 | 1 | 0 | 1 |
| *Cordylus meculae* | 0 | 1 | 0 | 1 |
| *Cordylus melanotus* | 0 | 1 | 0 | 1 |
| *Cordylus microlepidotus* | 0 | 1 | 0 | 1 |
| *Cordylus minor* | 0 | 1 | 0 | 1 |
| *Cordylus namaquensis* | 0 | 1 | 0 | 1 |
| *Cordylus nebulosus* | 0 | 1 | 0 | 1 |
| *Cordylus niger* | 0 | 1 | 0 | 1 |
| *Cordylus oelofseni* | 0 | 1 | 0 | 1 |
| *Cordylus peersi* | 0 | 1 | 0 | 1 |
| *Cordylus polyzonus* | 0 | 1 | 0 | 1 |
| *Cordylus pustulatus* | 0 | 1 | 0 | 1 |
| *Cordylus rhodesianus* | 0 | 1 | 0 | 1 |
| *Cordylus spinosus* | 0 | 1 | 0 | 1 |
| *Cordylus tasmani* | 0 | 1 | 0 | 1 |
| *Cordylus tropidosternum* | 0 | 1 | 0 | 1 |
| *Cordylus ukingensis* | 0 | 1 | 0 | 1 |
| *Cordylus vittifer* | 0 | 1 | 0 | 1 |
| *Cordylus warreni* | 0 | 1 | 0 | 1 |
| *Coronella austriaca* | 1 | 0 | 0 | 0 |
| *Coronella girondica* | 1 | 0 | 0 | 0 |
| *Corucia zebrata* | 0 | 1 | 0 | 1 |
| *Coryphophylax subcristatus* | 0 | 1 | 0 | 1 |
| *Corytophanes cristatus* | 0 | 1 | 0 | 1 |
| *Corytophanes percarinatus* | 0 | 1 | 0 | 1 |
| *Crenadactylus ocellatus* | 0 | 1 | 0 | 1 |
| *Cricosaura typica* | 0 | 1 | 0 | 1 |
| *Crisantophis nevermanni* | 0 | 1 | 0 | 1 |
| *Crocodilurus amazonicus* | 0 | 1 | 0 | 1 |
| *Crossobamon orientalis* | 0 | 1 | 0 | 1 |
| *Crotalus adamanteus* | 1 | 0 | 0 | 0 |
| *Crotalus aquilus* | 0 | 1 | 0 | 1 |
| *Crotalus atrox* | 1 | 0 | 0 | 0 |
| *Crotalus basiliscus* | 0 | 1 | 0 | 1 |
| *Crotalus catalinensis* | 1 | 0 | 0 | 0 |
| *Crotalus cerastes* | 1 | 0 | 0 | 0 |
| *Crotalus durissus* | 0 | 1 | 0 | 1 |
| *Crotalus enyo* | 0 | 1 | 0 | 1 |
| *Crotalus horridus* | 1 | 0 | 0 | 0 |
| *Crotalus intermedius* | 0 | 1 | 0 | 1 |
| *Crotalus lepidus* | 1 | 0 | 0 | 0 |
| *Crotalus mitchellii* | 1 | 0 | 0 | 0 |
| *Crotalus molossus* | 1 | 1 | 0 | 0 |
| *Crotalus oreganus* | 1 | 0 | 0 | 0 |
| *Crotalus polystictus* | 0 | 1 | 0 | 1 |
| *Crotalus pricei* | 1 | 0 | 0 | 0 |
| *Crotalus pusillus* | 0 | 1 | 0 | 1 |
| *Crotalus ravus* | 0 | 1 | 0 | 1 |
| *Crotalus ruber* | 1 | 0 | 0 | 0 |
| *Crotalus scutulatus* | 1 | 0 | 0 | 0 |
| *Crotalus simus* | 0 | 1 | 0 | 1 |
| *Crotalus tancitarensis* | 0 | 1 | 0 | 1 |
| *Crotalus tigris* | 1 | 0 | 0 | 0 |
| *Crotalus tortugensis* | 1 | 0 | 0 | 0 |
| *Crotalus totonacus* | 0 | 1 | 0 | 1 |
| *Crotalus transversus* | 0 | 1 | 0 | 1 |
| *Crotalus triseriatus* | 0 | 1 | 0 | 1 |
| *Crotalus viridis* | 1 | 0 | 0 | 0 |
| *Crotalus willardi* | 1 | 0 | 0 | 0 |
| *Crotaphopeltis tornieri* | 0 | 1 | 0 | 1 |
| *Crotaphytus antiquus* | 1 | 0 | 0 | 0 |
| *Crotaphytus bicinctores* | 1 | 0 | 0 | 0 |
| *Crotaphytus collaris* | 1 | 0 | 0 | 0 |
| *Crotaphytus grismeri* | 1 | 0 | 0 | 0 |
| *Crotaphytus insularis* | 1 | 0 | 0 | 0 |
| *Crotaphytus nebrius* | 1 | 0 | 0 | 0 |
| *Crotaphytus reticulatus* | 1 | 0 | 0 | 0 |
| *Crotaphytus vestigium* | 1 | 0 | 0 | 0 |
| *Cryophis hallbergi* | 0 | 1 | 0 | 1 |
| *Cryptactites peringueyi* | 0 | 1 | 0 | 1 |
| *Cryptelytrops albolabris* | 0 | 1 | 0 | 1 |
| *Cryptelytrops andersonii* | 0 | 1 | 0 | 1 |
| *Cryptelytrops cantori* | 0 | 1 | 0 | 1 |
| *Cryptelytrops erythrurus* | 0 | 1 | 0 | 1 |
| *Cryptelytrops fasciatus* | 0 | 1 | 0 | 1 |
| *Cryptelytrops insularis* | 0 | 1 | 0 | 1 |
| *Cryptelytrops kanburiensis* | 0 | 1 | 0 | 1 |
| *Cryptelytrops macrops* | 0 | 1 | 0 | 1 |
| *Cryptelytrops purpureomaculatus* | 0 | 1 | 0 | 1 |
| *Cryptelytrops septentrionalis* | 0 | 1 | 0 | 1 |
| *Cryptelytrops venustus* | 0 | 1 | 0 | 1 |
| *Cryptoblepharus boutonii* | 0 | 1 | 0 | 1 |
| *Cryptoblepharus nigropunctatus* | 0 | 1 | 0 | 1 |
| *Cryptoblepharus novocaledonicus* | 0 | 1 | 0 | 1 |
| *Ctenoblepharys adspersa* | 0 | 1 | 0 | 1 |
| *Ctenophorus adelaidensis* | 0 | 1 | 0 | 1 |
| *Ctenophorus caudicinctus* | 0 | 1 | 0 | 1 |
| *Ctenophorus clayi* | 0 | 1 | 0 | 1 |
| *Ctenophorus cristatus* | 0 | 1 | 0 | 1 |
| *Ctenophorus decresii* | 0 | 1 | 0 | 1 |
| *Ctenophorus femoralis* | 0 | 1 | 0 | 1 |
| *Ctenophorus fionni* | 0 | 1 | 0 | 1 |
| *Ctenophorus fordi* | 0 | 1 | 0 | 1 |
| *Ctenophorus gibba* | 0 | 1 | 0 | 1 |
| *Ctenophorus isolepis* | 0 | 1 | 0 | 1 |
| *Ctenophorus maculatus* | 0 | 1 | 0 | 1 |
| *Ctenophorus maculosus* | 0 | 1 | 0 | 1 |
| *Ctenophorus mckenziei* | 0 | 1 | 0 | 1 |
| *Ctenophorus nuchalis* | 0 | 1 | 0 | 1 |
| *Ctenophorus ornatus* | 0 | 1 | 0 | 1 |
| *Ctenophorus pictus* | 0 | 1 | 0 | 1 |
| *Ctenophorus reticulatus* | 0 | 1 | 0 | 1 |
| *Ctenophorus rufescens* | 0 | 1 | 0 | 1 |
| *Ctenophorus salinarum* | 0 | 1 | 0 | 1 |
| *Ctenophorus scutulatus* | 0 | 1 | 0 | 1 |
| *Ctenophorus tjantjalka* | 0 | 1 | 0 | 1 |
| *Ctenophorus vadnappa* | 0 | 1 | 0 | 1 |
| *Ctenosaura acanthura* | 0 | 1 | 0 | 1 |
| *Ctenosaura bakeri* | 0 | 1 | 0 | 1 |
| *Ctenosaura flavidorsalis* | 0 | 1 | 0 | 1 |
| *Ctenosaura hemilopha* | 0 | 1 | 0 | 1 |
| *Ctenosaura melanosterna* | 0 | 1 | 0 | 1 |
| *Ctenosaura oaxacana* | 0 | 1 | 0 | 1 |
| *Ctenosaura oedirhina* | 0 | 1 | 0 | 1 |
| *Ctenosaura palearis* | 0 | 1 | 0 | 1 |
| *Ctenosaura pectinata* | 0 | 1 | 0 | 1 |
| *Ctenosaura quinquecarinata* | 0 | 1 | 0 | 1 |
| *Ctenosaura similis* | 0 | 1 | 0 | 1 |
| *Ctenotus angusticeps* | 0 | 1 | 0 | 1 |
| *Ctenotus astarte* | 0 | 1 | 0 | 1 |
| *Ctenotus atlas* | 0 | 1 | 0 | 1 |
| *Ctenotus australis* | 0 | 1 | 0 | 1 |
| *Ctenotus brooksi* | 0 | 1 | 0 | 1 |
| *Ctenotus calurus* | 0 | 1 | 0 | 1 |
| *Ctenotus essingtonii* | 0 | 1 | 0 | 1 |
| *Ctenotus fallens* | 0 | 1 | 0 | 1 |
| *Ctenotus gagudju* | 0 | 1 | 0 | 1 |
| *Ctenotus grandis* | 0 | 1 | 0 | 1 |
| *Ctenotus greeri* | 0 | 1 | 0 | 1 |
| *Ctenotus hanloni* | 0 | 1 | 0 | 1 |
| *Ctenotus hebetior* | 0 | 1 | 0 | 1 |
| *Ctenotus hilli* | 0 | 1 | 0 | 1 |
| *Ctenotus inornatus* | 0 | 1 | 0 | 1 |
| *Ctenotus labillardieri* | 0 | 1 | 0 | 1 |
| *Ctenotus leae* | 0 | 1 | 0 | 1 |
| *Ctenotus leonhardii* | 0 | 1 | 0 | 1 |
| *Ctenotus maryani* | 0 | 1 | 0 | 1 |
| *Ctenotus mimetes* | 0 | 1 | 0 | 1 |
| *Ctenotus nasutus* | 0 | 1 | 0 | 1 |
| *Ctenotus olympicus* | 0 | 1 | 0 | 1 |
| *Ctenotus pantherinus* | 0 | 1 | 0 | 1 |
| *Ctenotus piankai* | 0 | 1 | 0 | 1 |
| *Ctenotus pulchellus* | 0 | 1 | 0 | 1 |
| *Ctenotus quattuordecimlineatus* | 0 | 1 | 0 | 1 |
| *Ctenotus rawlinsoni* | 0 | 1 | 0 | 1 |
| *Ctenotus regius* | 0 | 1 | 0 | 1 |
| *Ctenotus robustus* | 0 | 1 | 0 | 1 |
| *Ctenotus rubicundus* | 0 | 1 | 0 | 1 |
| *Ctenotus rutilans* | 0 | 1 | 0 | 1 |
| *Ctenotus saxatilis* | 0 | 1 | 0 | 1 |
| *Ctenotus schomburgkii* | 0 | 1 | 0 | 1 |
| *Ctenotus septenarius* | 0 | 1 | 0 | 1 |
| *Ctenotus serventyi* | 0 | 1 | 0 | 1 |
| *Ctenotus spaldingi* | 0 | 1 | 0 | 1 |
| *Ctenotus strauchii* | 0 | 1 | 0 | 1 |
| *Ctenotus taeniolatus* | 0 | 1 | 0 | 1 |
| *Ctenotus tanamiensis* | 0 | 1 | 0 | 1 |
| *Ctenotus uber* | 0 | 1 | 0 | 1 |
| *Ctenotus youngsoni* | 0 | 1 | 0 | 1 |
| *Cyclodina aenea* | 0 | 0 | 1 | 0 |
| *Cyclodina alani* | 0 | 0 | 1 | 0 |
| *Cyclodina hardyi* | 0 | 0 | 1 | 0 |
| *Cyclodina levidensa* | 0 | 0 | 1 | 0 |
| *Cyclodina lichenigera* | 0 | 1 | 0 | 1 |
| *Cyclodina macgregori* | 0 | 0 | 1 | 0 |
| *Cyclodina oliveri* | 0 | 0 | 1 | 0 |
| *Cyclodina ornata* | 0 | 0 | 1 | 0 |
| *Cyclodina townsi* | 0 | 0 | 1 | 0 |
| *Cyclodina whitakeri* | 0 | 0 | 1 | 0 |
| *Cyclodomorphus branchialis* | 0 | 1 | 0 | 1 |
| *Cyclodomorphus casuarinae* | 0 | 1 | 0 | 1 |
| *Cyclodomorphus michaeli* | 0 | 1 | 0 | 1 |
| *Cyclophiops major* | 0 | 1 | 0 | 1 |
| *Cyclura carinata* | 0 | 1 | 0 | 1 |
| *Cyclura collei* | 0 | 1 | 0 | 1 |
| *Cyclura cornuta* | 0 | 1 | 0 | 1 |
| *Cyclura cychlura* | 0 | 1 | 0 | 1 |
| *Cyclura nubila* | 0 | 1 | 0 | 1 |
| *Cyclura pinguis* | 0 | 1 | 0 | 1 |
| *Cyclura ricordi* | 0 | 1 | 0 | 1 |
| *Cyclura rileyi* | 0 | 1 | 0 | 1 |
| *Cylindrophis maculatus* | 0 | 1 | 0 | 1 |
| *Cylindrophis ruffus* | 0 | 1 | 0 | 1 |
| *Cynisca leucura* | 0 | 1 | 0 | 1 |
| *Cyrtodactylus agusanensis* | 0 | 1 | 0 | 1 |
| *Cyrtodactylus angularis* | 0 | 1 | 0 | 1 |
| *Cyrtodactylus annulatus* | 0 | 1 | 0 | 1 |
| *Cyrtodactylus ayeyarwadyensis* | 0 | 1 | 0 | 1 |
| *Cyrtodactylus consobrinus* | 0 | 1 | 0 | 1 |
| *Cyrtodactylus epiroticus* | 0 | 1 | 0 | 1 |
| *Cyrtodactylus intermedius* | 0 | 1 | 0 | 1 |
| *Cyrtodactylus irregularis* | 0 | 1 | 0 | 1 |
| *Cyrtodactylus jarujini* | 0 | 1 | 0 | 1 |
| *Cyrtodactylus klugei* | 0 | 1 | 0 | 1 |
| *Cyrtodactylus loriae* | 0 | 1 | 0 | 1 |
| *Cyrtodactylus louisiadensis* | 0 | 1 | 0 | 1 |
| *Cyrtodactylus marmoratus* | 0 | 1 | 0 | 1 |
| *Cyrtodactylus novaeguineae* | 0 | 1 | 0 | 1 |
| *Cyrtodactylus oldhami* | 0 | 1 | 0 | 1 |
| *Cyrtodactylus philippinicus* | 0 | 1 | 0 | 1 |
| *Cyrtodactylus pulchellus* | 0 | 1 | 0 | 1 |
| *Cyrtodactylus robustus* | 0 | 1 | 0 | 1 |
| *Cyrtodactylus sermowaiensis* | 0 | 1 | 0 | 1 |
| *Cyrtodactylus tripartitus* | 0 | 1 | 0 | 1 |
| *Cyrtodactylus tuberculatus* | 0 | 1 | 0 | 1 |
| *Cyrtopodion agamuroides* | 1 | 0 | 0 | 0 |
| *Cyrtopodion caspium* | 1 | 0 | 0 | 0 |
| *Cyrtopodion gastrophole* | 1 | 0 | 0 | 0 |
| *Cyrtopodion heterocercum* | 1 | 0 | 0 | 0 |
| *Cyrtopodion kotschyi* | 1 | 0 | 0 | 0 |
| *Cyrtopodion longipes* | 1 | 0 | 0 | 0 |
| *Cyrtopodion russowii* | 1 | 0 | 0 | 0 |
| *Cyrtopodion sagittiferum* | 1 | 0 | 0 | 0 |
| *Cyrtopodion scabrum* | 1 | 0 | 0 | 0 |
| *Cyrtopodion sistanensis* | 1 | 0 | 0 | 0 |
| *Cyrtopodion spinicaudum* | 1 | 0 | 0 | 0 |
| *Daboia russelii* | 0 | 1 | 0 | 1 |
| *Dalmatolacerta oxycephala* | 1 | 0 | 0 | 0 |
| *Darevskia alpina* | 1 | 0 | 0 | 0 |
| *Darevskia armeniaca* | 1 | 0 | 0 | 0 |
| *Darevskia bendimahiensis* | 1 | 0 | 0 | 0 |
| *Darevskia brauneri* | 1 | 0 | 0 | 0 |
| *Darevskia caucasica* | 1 | 0 | 0 | 0 |
| *Darevskia chlorogaster* | 1 | 0 | 0 | 0 |
| *Darevskia clarkorum* | 1 | 0 | 0 | 0 |
| *Darevskia daghestanica* | 1 | 0 | 0 | 0 |
| *Darevskia derjugini* | 1 | 0 | 0 | 0 |
| *Darevskia lindholmi* | 1 | 0 | 0 | 0 |
| *Darevskia mixta* | 1 | 0 | 0 | 0 |
| *Darevskia parvula* | 1 | 0 | 0 | 0 |
| *Darevskia portschinskii* | 1 | 0 | 0 | 0 |
| *Darevskia praticola* | 1 | 0 | 0 | 0 |
| *Darevskia raddei* | 1 | 0 | 0 | 0 |
| *Darevskia rostombekovi* | 1 | 0 | 0 | 0 |
| *Darevskia rudis* | 1 | 0 | 0 | 0 |
| *Darevskia sapphirina* | 1 | 0 | 0 | 0 |
| *Darevskia saxicola* | 1 | 0 | 0 | 0 |
| *Darevskia uzzelli* | 1 | 0 | 0 | 0 |
| *Darevskia valentini* | 1 | 0 | 0 | 0 |
| *Darlingtonia haetiana* | 0 | 1 | 0 | 1 |
| *Dasia grisea* | 0 | 1 | 0 | 1 |
| *Dasia olivacea* | 0 | 1 | 0 | 1 |
| *Dasypeltis atra* | 0 | 1 | 0 | 1 |
| *Dasypeltis confusa* | 0 | 1 | 0 | 1 |
| *Dasypeltis fasciata* | 0 | 1 | 0 | 1 |
| *Dasypeltis gansi* | 0 | 1 | 0 | 1 |
| *Dasypeltis sahelensis* | 0 | 1 | 0 | 1 |
| *Dasypeltis scabra* | 1 | 0 | 0 | 0 |
| *Davewakeum miriamae* | 0 | 1 | 0 | 1 |
| *Deinagkistrodon acutus* | 0 | 1 | 0 | 1 |
| *Delma australis* | 0 | 1 | 0 | 1 |
| *Delma borea* | 0 | 1 | 0 | 1 |
| *Delma butleri* | 0 | 1 | 0 | 1 |
| *Delma concinna* | 0 | 1 | 0 | 1 |
| *Delma fraseri* | 0 | 1 | 0 | 1 |
| *Delma grayii* | 0 | 1 | 0 | 1 |
| *Delma impar* | 0 | 1 | 0 | 1 |
| *Delma inornata* | 0 | 1 | 0 | 1 |
| *Delma labialis* | 0 | 1 | 0 | 1 |
| *Delma mitella* | 0 | 1 | 0 | 1 |
| *Delma molleri* | 0 | 1 | 0 | 1 |
| *Delma nasuta* | 0 | 1 | 0 | 1 |
| *Delma pax* | 0 | 1 | 0 | 1 |
| *Delma tincta* | 0 | 1 | 0 | 1 |
| *Delma torquata* | 0 | 1 | 0 | 1 |
| *Demansia papuensis* | 0 | 1 | 0 | 1 |
| *Demansia psammophis* | 0 | 1 | 0 | 1 |
| *Demansia vestigiata* | 0 | 1 | 0 | 1 |
| *Dendrelaphis bifrenalis* | 0 | 1 | 0 | 1 |
| *Dendrelaphis caudolineatus* | 0 | 1 | 0 | 1 |
| *Dendrelaphis caudolineolatus* | 0 | 1 | 0 | 1 |
| *Dendrelaphis schokari* | 0 | 1 | 0 | 1 |
| *Dendrelaphis tristis* | 0 | 1 | 0 | 1 |
| *Dendroaspis angusticeps* | 0 | 1 | 0 | 1 |
| *Dendroaspis polylepis* | 0 | 1 | 0 | 1 |
| *Dendrophidion dendrophis* | 0 | 1 | 0 | 1 |
| *Dendrophidion percarinatus* | 0 | 1 | 0 | 1 |
| *Denisonia devisi* | 0 | 1 | 0 | 1 |
| *Diadophis punctatus* | 1 | 0 | 0 | 0 |
| *Dibamus bourreti* | 0 | 1 | 0 | 1 |
| *Dibamus celebensis* | 0 | 1 | 0 | 1 |
| *Dibamus greeri* | 0 | 1 | 0 | 1 |
| *Dibamus montanus* | 0 | 1 | 0 | 1 |
| *Dibamus novaeguineae* | 0 | 1 | 0 | 1 |
| *Dibamus seramensis* | 0 | 1 | 0 | 1 |
| *Dibamus tiomanensis* | 0 | 1 | 0 | 1 |
| *Dicrodon guttulatum* | 0 | 1 | 0 | 1 |
| *Dierogekko inexpectatus* | 0 | 1 | 0 | 1 |
| *Dierogekko insularis* | 0 | 1 | 0 | 1 |
| *Dierogekko kaalaensis* | 0 | 1 | 0 | 1 |
| *Dierogekko koniambo* | 0 | 1 | 0 | 1 |
| *Dierogekko nehoueensis* | 0 | 1 | 0 | 1 |
| *Dierogekko poumensis* | 0 | 1 | 0 | 1 |
| *Dierogekko thomaswhitei* | 0 | 1 | 0 | 1 |
| *Dierogekko validiclavis* | 0 | 1 | 0 | 1 |
| *Dinarolacerta montenegrina* | 1 | 0 | 0 | 0 |
| *Dinarolacerta mosorensis* | 1 | 0 | 0 | 0 |
| *Dinodon rufozonatum* | 1 | 0 | 0 | 0 |
| *Dinodon semicarinatum* | 0 | 1 | 0 | 1 |
| *Diplodactylus capensis* | 0 | 1 | 0 | 1 |
| *Diplodactylus conspicillatus* | 0 | 1 | 0 | 1 |
| *Diplodactylus fulleri* | 0 | 1 | 0 | 1 |
| *Diplodactylus galeatus* | 0 | 1 | 0 | 1 |
| *Diplodactylus granariensis* | 0 | 1 | 0 | 1 |
| *Diplodactylus klugei* | 0 | 1 | 0 | 1 |
| *Diplodactylus mitchelli* | 0 | 1 | 0 | 1 |
| *Diplodactylus ornatus* | 0 | 1 | 0 | 1 |
| *Diplodactylus polyophthalmus* | 0 | 1 | 0 | 1 |
| *Diplodactylus pulcher* | 0 | 1 | 0 | 1 |
| *Diplodactylus savagei* | 0 | 1 | 0 | 1 |
| *Diplodactylus tessellatus* | 0 | 1 | 0 | 1 |
| *Diplodactylus vittatus* | 0 | 1 | 0 | 1 |
| *Diploglossus bilobatus* | 0 | 1 | 0 | 1 |
| *Diploglossus pleii* | 0 | 1 | 0 | 1 |
| *Diplolaemus darwinii* | 0 | 0 | 1 | 0 |
| *Diplometopon zarudnyi* | 1 | 0 | 0 | 0 |
| *Diporiphora albilabris* | 0 | 1 | 0 | 1 |
| *Diporiphora arnhemica* | 0 | 1 | 0 | 1 |
| *Diporiphora australis* | 0 | 1 | 0 | 1 |
| *Diporiphora bennettii* | 0 | 1 | 0 | 1 |
| *Diporiphora bilineata* | 0 | 1 | 0 | 1 |
| *Diporiphora lalliae* | 0 | 1 | 0 | 1 |
| *Diporiphora linga* | 0 | 1 | 0 | 1 |
| *Diporiphora magna* | 0 | 1 | 0 | 1 |
| *Diporiphora pindan* | 0 | 1 | 0 | 1 |
| *Diporiphora reginae* | 0 | 1 | 0 | 1 |
| *Diporiphora superba* | 0 | 1 | 0 | 1 |
| *Diporiphora valens* | 0 | 1 | 0 | 1 |
| *Diporiphora winneckei* | 0 | 1 | 0 | 1 |
| *Dipsadoboa unicolor* | 0 | 1 | 0 | 1 |
| *Dipsas albifrons* | 0 | 1 | 0 | 1 |
| *Dipsas articulata* | 0 | 1 | 0 | 1 |
| *Dipsas catesbyi* | 0 | 1 | 0 | 1 |
| *Dipsas indica* | 0 | 1 | 0 | 1 |
| *Dipsas neivai* | 0 | 1 | 0 | 1 |
| *Dipsas pratti* | 0 | 1 | 0 | 1 |
| *Dipsas variegata* | 0 | 1 | 0 | 1 |
| *Dipsina multimaculata* | 0 | 1 | 0 | 1 |
| *Dipsosaurus dorsalis* | 1 | 0 | 0 | 0 |
| *Dispholidus typus* | 0 | 1 | 0 | 1 |
| *Disteira kingii* | 0 | 1 | 0 | 1 |
| *Disteira major* | 0 | 1 | 0 | 1 |
| *Ditypophis vivax* | 0 | 1 | 0 | 1 |
| *Dixonius melanostictus* | 0 | 1 | 0 | 1 |
| *Dixonius siamensis* | 0 | 1 | 0 | 1 |
| *Dixonius vietnamensis* | 0 | 1 | 0 | 1 |
| *Dolichophis caspius* | 1 | 0 | 0 | 0 |
| *Dolichophis jugularis* | 1 | 0 | 0 | 0 |
| *Dolichophis schmidti* | 1 | 0 | 0 | 0 |
| *Dracaena guianensis* | 0 | 1 | 0 | 1 |
| *Draco beccarii* | 0 | 1 | 0 | 1 |
| *Draco biaro* | 0 | 1 | 0 | 1 |
| *Draco bimaculatus* | 0 | 1 | 0 | 1 |
| *Draco blanfordii* | 0 | 1 | 0 | 1 |
| *Draco boschmai* | 0 | 1 | 0 | 1 |
| *Draco bourouniensis* | 0 | 1 | 0 | 1 |
| *Draco caerulhians* | 0 | 1 | 0 | 1 |
| *Draco cornutus* | 0 | 1 | 0 | 1 |
| *Draco cristatellus* | 0 | 1 | 0 | 1 |
| *Draco cyanopterus* | 0 | 1 | 0 | 1 |
| *Draco dussumieri* | 0 | 1 | 0 | 1 |
| *Draco fimbriatus* | 0 | 1 | 0 | 1 |
| *Draco guentheri* | 0 | 1 | 0 | 1 |
| *Draco haematopogon* | 0 | 1 | 0 | 1 |
| *Draco indochinensis* | 0 | 1 | 0 | 1 |
| *Draco lineatus* | 0 | 1 | 0 | 1 |
| *Draco maculatus* | 0 | 1 | 0 | 1 |
| *Draco maximus* | 0 | 1 | 0 | 1 |
| *Draco melanopogon* | 0 | 1 | 0 | 1 |
| *Draco mindanensis* | 0 | 1 | 0 | 1 |
| *Draco obscurus* | 0 | 1 | 0 | 1 |
| *Draco ornatus* | 0 | 1 | 0 | 1 |
| *Draco palawanensis* | 0 | 1 | 0 | 1 |
| *Draco quadrasi* | 0 | 1 | 0 | 1 |
| *Draco quinquefasciatus* | 0 | 1 | 0 | 1 |
| *Draco reticulatus* | 0 | 1 | 0 | 1 |
| *Draco rhytisma* | 0 | 1 | 0 | 1 |
| *Draco spilonotus* | 0 | 1 | 0 | 1 |
| *Draco spilopterus* | 0 | 1 | 0 | 1 |
| *Draco taeniopterus* | 0 | 1 | 0 | 1 |
| *Draco timorensis* | 0 | 1 | 0 | 1 |
| *Draco volans* | 0 | 1 | 0 | 1 |
| *Drepanoides anomalus* | 0 | 1 | 0 | 1 |
| *Dromicodryas bernieri* | 0 | 1 | 0 | 1 |
| *Dromicodryas quadrilineatus* | 0 | 1 | 0 | 1 |
| *Drymarchon corais* | 0 | 1 | 0 | 1 |
| *Drymobius rhombifer* | 0 | 1 | 0 | 1 |
| *Drymoluber brazili* | 0 | 1 | 0 | 1 |
| *Drymoluber dichrous* | 0 | 1 | 0 | 1 |
| *Dryocalamus nympha* | 0 | 1 | 0 | 1 |
| *Drysdalia coronoides* | 0 | 1 | 0 | 1 |
| *Drysdalia mastersii* | 0 | 1 | 0 | 1 |
| *Duberria lutrix* | 0 | 1 | 0 | 1 |
| *Duberria variegata* | 0 | 1 | 0 | 1 |
| *Ebenavia inunguis* | 0 | 1 | 0 | 1 |
| *Echinanthera melanostigma* | 0 | 1 | 0 | 1 |
| *Echinanthera undulata* | 0 | 1 | 0 | 1 |
| *Echiopsis atriceps* | 0 | 1 | 0 | 1 |
| *Echiopsis curta* | 0 | 1 | 0 | 1 |
| *Echis carinatus* | 1 | 0 | 0 | 0 |
| *Echis coloratus* | 1 | 0 | 0 | 0 |
| *Echis jogeri* | 1 | 0 | 0 | 0 |
| *Echis leucogaster* | 1 | 0 | 0 | 0 |
| *Echis ocellatus* | 0 | 1 | 0 | 1 |
| *Echis omanensis* | 1 | 0 | 0 | 0 |
| *Echis pyramidum* | 1 | 0 | 0 | 0 |
| *Ecpleopus gaudichaudii* | 0 | 1 | 0 | 1 |
| *Egernia depressa* | 0 | 1 | 0 | 1 |
| *Egernia guthega* | 0 | 1 | 0 | 1 |
| *Egernia hosmeri* | 0 | 1 | 0 | 1 |
| *Egernia inornata* | 0 | 1 | 0 | 1 |
| *Egernia kingii* | 0 | 1 | 0 | 1 |
| *Egernia kintorei* | 0 | 1 | 0 | 1 |
| *Egernia luctuosa* | 0 | 1 | 0 | 1 |
| *Egernia margaretae* | 0 | 1 | 0 | 1 |
| *Egernia modesta* | 0 | 1 | 0 | 1 |
| *Egernia montana* | 0 | 1 | 0 | 1 |
| *Egernia multiscutata* | 0 | 1 | 0 | 1 |
| *Egernia napoleonis* | 0 | 1 | 0 | 1 |
| *Egernia pulchra* | 0 | 1 | 0 | 1 |
| *Egernia richardi* | 0 | 1 | 0 | 1 |
| *Egernia saxatilis* | 0 | 1 | 0 | 1 |
| *Egernia stokesii* | 0 | 1 | 0 | 1 |
| *Egernia striata* | 0 | 1 | 0 | 1 |
| *Egernia whitii* | 0 | 1 | 0 | 1 |
| *Eirenis aurolineatus* | 1 | 0 | 0 | 0 |
| *Eirenis barani* | 1 | 0 | 0 | 0 |
| *Eirenis collaris* | 1 | 0 | 0 | 0 |
| *Eirenis coronelloides* | 1 | 0 | 0 | 0 |
| *Eirenis decemlineatus* | 1 | 0 | 0 | 0 |
| *Eirenis eiselti* | 1 | 0 | 0 | 0 |
| *Eirenis levantinus* | 1 | 0 | 0 | 0 |
| *Eirenis lineomaculatus* | 1 | 0 | 0 | 0 |
| *Eirenis medus* | 1 | 0 | 0 | 0 |
| *Eirenis modestus* | 1 | 0 | 0 | 0 |
| *Eirenis punctatolineatus* | 1 | 0 | 0 | 0 |
| *Eirenis rothii* | 1 | 0 | 0 | 0 |
| *Eirenis thospitis* | 1 | 0 | 0 | 0 |
| *Elaphe bimaculata* | 1 | 0 | 0 | 0 |
| *Elaphe carinata* | 1 | 0 | 0 | 0 |
| *Elaphe climacophora* | 1 | 0 | 0 | 0 |
| *Elaphe davidi* | 1 | 0 | 0 | 0 |
| *Elaphe dione* | 1 | 0 | 0 | 0 |
| *Elaphe quadrivirgata* | 1 | 0 | 0 | 0 |
| *Elaphe quatuorlineata* | 1 | 0 | 0 | 0 |
| *Elaphe rufodorsata* | 1 | 0 | 0 | 0 |
| *Elaphe sauromates* | 1 | 0 | 0 | 0 |
| *Elaphe schrenckii* | 1 | 0 | 0 | 0 |
| *Elapognathus coronata* | 0 | 1 | 0 | 1 |
| *Elapomorphus quinquelineatus* | 0 | 1 | 0 | 1 |
| *Elapsoidea nigra* | 0 | 1 | 0 | 1 |
| *Elapsoidea semiannulata* | 0 | 1 | 0 | 1 |
| *Elapsoidea sundevallii* | 0 | 1 | 0 | 1 |
| *Elasmodactylus tetensis* | 0 | 1 | 0 | 1 |
| *Elasmodactylus tuberculosus* | 0 | 1 | 0 | 1 |
| *Elgaria coerulea* | 1 | 0 | 0 | 0 |
| *Elgaria kingii* | 1 | 0 | 0 | 0 |
| *Elgaria multicarinata* | 1 | 0 | 0 | 0 |
| *Elgaria panamintina* | 1 | 0 | 0 | 0 |
| *Elgaria paucicarinata* | 1 | 0 | 0 | 0 |
| *Emoia atrocostata* | 0 | 1 | 0 | 1 |
| *Emoia caeruleocauda* | 0 | 1 | 0 | 1 |
| *Emoia concolor* | 0 | 1 | 0 | 1 |
| *Emoia cyanogaster* | 0 | 1 | 0 | 1 |
| *Emoia cyanura* | 0 | 1 | 0 | 1 |
| *Emoia impar* | 0 | 1 | 0 | 1 |
| *Emoia isolata* | 0 | 1 | 0 | 1 |
| *Emoia jakati* | 0 | 1 | 0 | 1 |
| *Emoia loyaltiensis* | 0 | 1 | 0 | 1 |
| *Emoia physicae* | 0 | 1 | 0 | 1 |
| *Emoia pseudocyanura* | 0 | 1 | 0 | 1 |
| *Emoia schmidti* | 0 | 1 | 0 | 1 |
| *Emoia tongana* | 0 | 1 | 0 | 1 |
| *Emydocephalus annulatus* | 0 | 1 | 0 | 1 |
| *Enhydrina schistosa* | 0 | 1 | 0 | 1 |
| *Enhydris bocourti* | 0 | 1 | 0 | 1 |
| *Enhydris chinensis* | 0 | 1 | 0 | 1 |
| *Enhydris enhydris* | 0 | 1 | 0 | 1 |
| *Enhydris innominata* | 0 | 1 | 0 | 1 |
| *Enhydris jagorii* | 0 | 1 | 0 | 1 |
| *Enhydris longicauda* | 0 | 1 | 0 | 1 |
| *Enhydris matannensis* | 0 | 1 | 0 | 1 |
| *Enhydris plumbea* | 0 | 1 | 0 | 1 |
| *Enhydris polylepis* | 0 | 1 | 0 | 1 |
| *Enhydris punctata* | 0 | 1 | 0 | 1 |
| *Enyalioides heterolepis* | 0 | 1 | 0 | 1 |
| *Enyalioides laticeps* | 0 | 1 | 0 | 1 |
| *Enyalioides microlepis* | 0 | 1 | 0 | 1 |
| *Enyalioides oshaughnessyi* | 0 | 1 | 0 | 1 |
| *Enyalioides palpebralis* | 0 | 1 | 0 | 1 |
| *Enyalioides praestabilis* | 0 | 1 | 0 | 1 |
| *Enyalius bilineatus* | 0 | 1 | 0 | 1 |
| *Enyalius leechii* | 0 | 1 | 0 | 1 |
| *Ephalophis greyae* | 0 | 1 | 0 | 1 |
| *Epicrates angulifer* | 0 | 1 | 0 | 1 |
| *Epicrates cenchria* | 0 | 1 | 0 | 1 |
| *Epicrates chrysogaster* | 0 | 1 | 0 | 1 |
| *Epicrates exsul* | 0 | 1 | 0 | 1 |
| *Epicrates fordi* | 0 | 1 | 0 | 1 |
| *Epicrates inornatus* | 0 | 1 | 0 | 1 |
| *Epicrates monensis* | 0 | 1 | 0 | 1 |
| *Epicrates striatus* | 0 | 1 | 0 | 1 |
| *Epicrates subflavus* | 0 | 1 | 0 | 1 |
| *Eremias argus* | 1 | 0 | 0 | 0 |
| *Eremias arguta* | 1 | 0 | 0 | 0 |
| *Eremias brenchleyi* | 1 | 0 | 0 | 0 |
| *Eremias grammica* | 1 | 0 | 0 | 0 |
| *Eremias montanus* | 1 | 0 | 0 | 0 |
| *Eremias multiocellata* | 1 | 0 | 0 | 0 |
| *Eremias nigrolateralis* | 1 | 0 | 0 | 0 |
| *Eremias persica* | 1 | 0 | 0 | 0 |
| *Eremias pleskei* | 1 | 0 | 0 | 0 |
| *Eremias przewalskii* | 1 | 0 | 0 | 0 |
| *Eremias velox* | 1 | 0 | 0 | 0 |
| *Eremias vermiculata* | 1 | 0 | 0 | 0 |
| *Eremiascincus fasciolatus* | 0 | 1 | 0 | 1 |
| *Eremiascincus richardsonii* | 0 | 1 | 0 | 1 |
| *Eristicophis macmahoni* | 1 | 0 | 0 | 0 |
| *Erpeton tentaculatum* | 0 | 1 | 0 | 1 |
| *Erythrolamprus aesculapii* | 0 | 1 | 0 | 1 |
| *Erythrolamprus mimus* | 0 | 1 | 0 | 1 |
| *Eryx elegans* | 1 | 0 | 0 | 0 |
| *Eryx jaculus* | 1 | 0 | 0 | 0 |
| *Eryx jayakari* | 1 | 0 | 0 | 0 |
| *Eryx johnii* | 1 | 0 | 0 | 0 |
| *Eryx miliaris* | 1 | 0 | 0 | 0 |
| *Eryx tataricus* | 1 | 0 | 0 | 0 |
| *Eublepharis macularius* | 1 | 0 | 0 | 0 |
| *Eublepharis turcmenicus* | 1 | 0 | 0 | 0 |
| *Eugongylus albofasciolatus* | 0 | 1 | 0 | 1 |
| *Eugongylus rufescens* | 0 | 1 | 0 | 1 |
| *Eulamprus amplus* | 0 | 1 | 0 | 1 |
| *Eulamprus brachyosoma* | 0 | 1 | 0 | 1 |
| *Eulamprus frerei* | 0 | 1 | 0 | 1 |
| *Eulamprus heatwolei* | 0 | 1 | 0 | 1 |
| *Eulamprus kosciuskoi* | 0 | 1 | 0 | 1 |
| *Eulamprus leuraensis* | 0 | 1 | 0 | 1 |
| *Eulamprus luteilateralis* | 0 | 1 | 0 | 1 |
| *Eulamprus martini* | 0 | 1 | 0 | 1 |
| *Eulamprus murrayi* | 0 | 1 | 0 | 1 |
| *Eulamprus quoyii* | 0 | 1 | 0 | 1 |
| *Eulamprus sokosoma* | 0 | 1 | 0 | 1 |
| *Eulamprus tenuis* | 0 | 1 | 0 | 1 |
| *Eulamprus tigrinus* | 0 | 1 | 0 | 1 |
| *Eulamprus tryoni* | 0 | 1 | 0 | 1 |
| *Eulamprus tympanum* | 0 | 1 | 0 | 1 |
| *Euleptes europaea* | 1 | 0 | 0 | 0 |
| *Eumeces algeriensis* | 1 | 0 | 0 | 0 |
| *Eumeces schneideri* | 1 | 0 | 0 | 0 |
| *Eumecia anchietae* | 0 | 1 | 0 | 1 |
| *Eunectes murinus* | 0 | 1 | 0 | 1 |
| *Eunectes notaeus* | 0 | 1 | 0 | 1 |
| *Euprepiophis conspicillata* | 1 | 0 | 0 | 0 |
| *Euprepiophis mandarina* | 0 | 1 | 0 | 1 |
| *Eurolophosaurus amathites* | 0 | 1 | 0 | 1 |
| *Eurolophosaurus divaricatus* | 0 | 1 | 0 | 1 |
| *Eurolophosaurus nanuzae* | 0 | 1 | 0 | 1 |
| *Eurydactylodes agricolae* | 0 | 1 | 0 | 1 |
| *Eurydactylodes occidentalis* | 0 | 1 | 0 | 1 |
| *Eurydactylodes symmetricus* | 0 | 1 | 0 | 1 |
| *Eurydactylodes vieillardi* | 0 | 1 | 0 | 1 |
| *Eurylepis taeniolatus* | 1 | 0 | 0 | 0 |
| *Eutropis beddomii* | 0 | 1 | 0 | 1 |
| *Eutropis bibronii* | 0 | 1 | 0 | 1 |
| *Eutropis clivicola* | 0 | 1 | 0 | 1 |
| *Eutropis cumingi* | 0 | 1 | 0 | 1 |
| *Eutropis longicaudata* | 0 | 1 | 0 | 1 |
| *Eutropis macularia* | 0 | 1 | 0 | 1 |
| *Eutropis multicarinata* | 0 | 1 | 0 | 1 |
| *Eutropis multifasciata* | 0 | 1 | 0 | 1 |
| *Eutropis nagarjuni* | 0 | 1 | 0 | 1 |
| *Eutropis rudis* | 0 | 1 | 0 | 1 |
| *Eutropis trivittata* | 0 | 1 | 0 | 1 |
| *Exiliboa placata* | 0 | 1 | 0 | 1 |
| *Farancia abacura* | 1 | 0 | 0 | 0 |
| *Farancia erytrogramma* | 1 | 0 | 0 | 0 |
| *Feylinia currori* | 0 | 1 | 0 | 1 |
| *Feylinia grandisquamis* | 0 | 1 | 0 | 1 |
| *Feylinia polylepis* | 0 | 1 | 0 | 1 |
| *Ficimia streckeri* | 1 | 0 | 0 | 0 |
| *Fordonia leucobalia* | 0 | 1 | 0 | 1 |
| *Furcifer angeli* | 0 | 1 | 0 | 1 |
| *Furcifer antimena* | 0 | 1 | 0 | 1 |
| *Furcifer balteatus* | 0 | 1 | 0 | 1 |
| *Furcifer belalandaensis* | 0 | 1 | 0 | 1 |
| *Furcifer bifidus* | 0 | 1 | 0 | 1 |
| *Furcifer campani* | 0 | 1 | 0 | 1 |
| *Furcifer cephalolepis* | 0 | 1 | 0 | 1 |
| *Furcifer labordi* | 0 | 1 | 0 | 1 |
| *Furcifer lateralis* | 0 | 1 | 0 | 1 |
| *Furcifer minor* | 0 | 1 | 0 | 1 |
| *Furcifer oustaleti* | 0 | 1 | 0 | 1 |
| *Furcifer pardalis* | 0 | 1 | 0 | 1 |
| *Furcifer petteri* | 0 | 1 | 0 | 1 |
| *Furcifer polleni* | 0 | 1 | 0 | 1 |
| *Furcifer verrucosus* | 0 | 1 | 0 | 1 |
| *Furcifer willsii* | 0 | 1 | 0 | 1 |
| *Furina diadema* | 0 | 1 | 0 | 1 |
| *Furina ornata* | 0 | 1 | 0 | 1 |
| *Gallotia atlantica* | 1 | 0 | 0 | 0 |
| *Gallotia caesaris* | 1 | 0 | 0 | 0 |
| *Gallotia galloti* | 1 | 0 | 0 | 0 |
| *Gallotia gomerana* | 1 | 0 | 0 | 0 |
| *Gallotia intermedia* | 1 | 0 | 0 | 0 |
| *Gallotia simonyi* | 1 | 0 | 0 | 0 |
| *Gallotia stehlini* | 1 | 0 | 0 | 0 |
| *Gambelia copeii* | 1 | 0 | 0 | 0 |
| *Gambelia sila* | 1 | 0 | 0 | 0 |
| *Gambelia wislizenii* | 1 | 0 | 0 | 0 |
| *Garthius chaseni* | 0 | 1 | 0 | 1 |
| *Gastropholis prasina* | 0 | 1 | 0 | 1 |
| *Gastropholis vittata* | 0 | 1 | 0 | 1 |
| *Geckoella triedrus* | 0 | 1 | 0 | 1 |
| *Geckolepis maculata* | 0 | 1 | 0 | 1 |
| *Geckolepis typica* | 0 | 1 | 0 | 1 |
| *Geckonia chazaliae* | 1 | 0 | 0 | 0 |
| *Gehyra australis* | 0 | 1 | 0 | 1 |
| *Gehyra baliola* | 0 | 1 | 0 | 1 |
| *Gehyra barea* | 0 | 1 | 0 | 1 |
| *Gehyra borroloola* | 0 | 1 | 0 | 1 |
| *Gehyra brevipalmata* | 0 | 1 | 0 | 1 |
| *Gehyra catenata* | 0 | 1 | 0 | 1 |
| *Gehyra dubia* | 0 | 1 | 0 | 1 |
| *Gehyra fehlmanni* | 0 | 1 | 0 | 1 |
| *Gehyra koira* | 0 | 1 | 0 | 1 |
| *Gehyra lacerata* | 0 | 1 | 0 | 1 |
| *Gehyra marginata* | 0 | 1 | 0 | 1 |
| *Gehyra membranacruralis* | 0 | 1 | 0 | 1 |
| *Gehyra minuta* | 0 | 1 | 0 | 1 |
| *Gehyra montium* | 0 | 1 | 0 | 1 |
| *Gehyra mutilata* | 0 | 1 | 0 | 1 |
| *Gehyra nana* | 0 | 1 | 0 | 1 |
| *Gehyra occidentalis* | 0 | 1 | 0 | 1 |
| *Gehyra oceanica* | 0 | 1 | 0 | 1 |
| *Gehyra pamela* | 0 | 1 | 0 | 1 |
| *Gehyra pilbara* | 0 | 1 | 0 | 1 |
| *Gehyra punctata* | 0 | 1 | 0 | 1 |
| *Gehyra purpurascens* | 0 | 1 | 0 | 1 |
| *Gehyra robusta* | 0 | 1 | 0 | 1 |
| *Gehyra variegata* | 0 | 1 | 0 | 1 |
| *Gehyra xenopus* | 0 | 1 | 0 | 1 |
| *Gekko athymus* | 0 | 1 | 0 | 1 |
| *Gekko auriverrucosus* | 1 | 0 | 0 | 0 |
| *Gekko badenii* | 0 | 1 | 0 | 1 |
| *Gekko chinensis* | 0 | 1 | 0 | 1 |
| *Gekko crombota* | 0 | 1 | 0 | 1 |
| *Gekko gecko* | 0 | 1 | 0 | 1 |
| *Gekko grossmanni* | 0 | 1 | 0 | 1 |
| *Gekko hokouensis* | 0 | 1 | 0 | 1 |
| *Gekko japonicus* | 1 | 0 | 0 | 0 |
| *Gekko mindorensis* | 0 | 1 | 0 | 1 |
| *Gekko monarchus* | 0 | 1 | 0 | 1 |
| *Gekko petricolus* | 0 | 1 | 0 | 1 |
| *Gekko porosus* | 0 | 1 | 0 | 1 |
| *Gekko romblon* | 0 | 1 | 0 | 1 |
| *Gekko smithii* | 0 | 1 | 0 | 1 |
| *Gekko swinhonis* | 1 | 0 | 0 | 0 |
| *Gekko vittatus* | 0 | 1 | 0 | 1 |
| *Geocalamus acutus* | 0 | 1 | 0 | 1 |
| *Geophis carinosus* | 0 | 1 | 0 | 1 |
| *Geophis godmani* | 0 | 1 | 0 | 1 |
| *Gerarda prevostiana* | 0 | 1 | 0 | 1 |
| *Gerrhonotus infernalis* | 1 | 0 | 0 | 0 |
| *Gerrhonotus liocephalus* | 0 | 1 | 0 | 1 |
| *Gerrhonotus parvus* | 1 | 0 | 0 | 0 |
| *Gerrhosaurus flavigularis* | 0 | 1 | 0 | 1 |
| *Gerrhosaurus major* | 0 | 1 | 0 | 1 |
| *Gerrhosaurus multilineatus* | 0 | 1 | 0 | 1 |
| *Gerrhosaurus nigrolineatus* | 0 | 1 | 0 | 1 |
| *Gerrhosaurus skoogi* | 0 | 1 | 0 | 1 |
| *Gerrhosaurus typicus* | 0 | 1 | 0 | 1 |
| *Gerrhosaurus validus* | 0 | 1 | 0 | 1 |
| *Glaphyromorphus cracens* | 0 | 1 | 0 | 1 |
| *Glaphyromorphus darwiniensis* | 0 | 1 | 0 | 1 |
| *Glaphyromorphus douglasi* | 0 | 1 | 0 | 1 |
| *Glaphyromorphus fuscicaudis* | 0 | 1 | 0 | 1 |
| *Glaphyromorphus gracilipes* | 0 | 1 | 0 | 1 |
| *Glaphyromorphus isolepis* | 0 | 1 | 0 | 1 |
| *Glaphyromorphus mjobergi* | 0 | 1 | 0 | 1 |
| *Glaphyromorphus pardalis* | 0 | 1 | 0 | 1 |
| *Glaphyromorphus pumilus* | 0 | 1 | 0 | 1 |
| *Glaphyromorphus punctulatus* | 0 | 1 | 0 | 1 |
| *Gloydius blomhoffii* | 1 | 0 | 0 | 0 |
| *Gloydius brevicaudus* | 1 | 0 | 0 | 0 |
| *Gloydius halys* | 1 | 0 | 0 | 0 |
| *Gloydius intermedius* | 1 | 0 | 0 | 0 |
| *Gloydius saxatilis* | 1 | 0 | 0 | 0 |
| *Gloydius shedaoensis* | 1 | 0 | 0 | 0 |
| *Gloydius strauchi* | 1 | 0 | 0 | 0 |
| *Gloydius tsushimaensis* | 1 | 0 | 0 | 0 |
| *Gloydius ussuriensis* | 1 | 0 | 0 | 0 |
| *Gnypetoscincus queenslandiae* | 0 | 1 | 0 | 1 |
| *Goggia lineata* | 0 | 1 | 0 | 1 |
| *Gomesophis brasiliensis* | 0 | 1 | 0 | 1 |
| *Gonatodes albogularis* | 0 | 1 | 0 | 1 |
| *Gonatodes alexandermendesi* | 0 | 1 | 0 | 1 |
| *Gonatodes annularis* | 0 | 1 | 0 | 1 |
| *Gonatodes antillensis* | 0 | 1 | 0 | 1 |
| *Gonatodes caudiscutatus* | 0 | 1 | 0 | 1 |
| *Gonatodes ceciliae* | 0 | 1 | 0 | 1 |
| *Gonatodes concinnatus* | 0 | 1 | 0 | 1 |
| *Gonatodes daudini* | 0 | 1 | 0 | 1 |
| *Gonatodes eladioi* | 0 | 1 | 0 | 1 |
| *Gonatodes falconensis* | 0 | 1 | 0 | 1 |
| *Gonatodes hasemani* | 0 | 1 | 0 | 1 |
| *Gonatodes humeralis* | 0 | 1 | 0 | 1 |
| *Gonatodes infernalis* | 0 | 1 | 0 | 1 |
| *Gonatodes ocellatus* | 0 | 1 | 0 | 1 |
| *Gonatodes petersi* | 0 | 1 | 0 | 1 |
| *Gonatodes purpurogularis* | 0 | 1 | 0 | 1 |
| *Gonatodes seigliei* | 0 | 1 | 0 | 1 |
| *Gonatodes superciliaris* | 0 | 1 | 0 | 1 |
| *Gonatodes taniae* | 0 | 1 | 0 | 1 |
| *Gonatodes vittatus* | 0 | 1 | 0 | 1 |
| *Gongylomorphus bojerii* | 0 | 1 | 0 | 1 |
| *Gongylophis colubrinus* | 0 | 1 | 0 | 1 |
| *Gongylophis conicus* | 0 | 1 | 0 | 1 |
| *Gonionotophis brussauxi* | 0 | 1 | 0 | 1 |
| *Goniurosaurus araneus* | 0 | 1 | 0 | 1 |
| *Goniurosaurus catbaensis* | 0 | 1 | 0 | 1 |
| *Goniurosaurus kuroiwae* | 0 | 1 | 0 | 1 |
| *Goniurosaurus lichtenfelderi* | 0 | 1 | 0 | 1 |
| *Goniurosaurus luii* | 0 | 1 | 0 | 1 |
| *Gonocephalus chamaeleontinus* | 0 | 1 | 0 | 1 |
| *Gonocephalus grandis* | 0 | 1 | 0 | 1 |
| *Gonocephalus kuhlii* | 0 | 1 | 0 | 1 |
| *Gonocephalus robinsonii* | 0 | 1 | 0 | 1 |
| *Gonyosoma jansenii* | 0 | 1 | 0 | 1 |
| *Gonyosoma oxycephalum* | 0 | 1 | 0 | 1 |
| *Graciliscincus shonae* | 0 | 1 | 0 | 1 |
| *Grayia ornata* | 0 | 1 | 0 | 1 |
| *Grayia smithii* | 0 | 1 | 0 | 1 |
| *Grayia tholloni* | 0 | 1 | 0 | 1 |
| *Gyalopion canum* | 1 | 0 | 0 | 0 |
| *Gymnophthalmus cryptus* | 0 | 1 | 0 | 1 |
| *Gymnophthalmus leucomystax* | 0 | 1 | 0 | 1 |
| *Gymnophthalmus pleei* | 0 | 1 | 0 | 1 |
| *Gymnophthalmus speciosus* | 0 | 1 | 0 | 1 |
| *Gymnophthalmus underwoodi* | 0 | 1 | 0 | 1 |
| *Gymnophthalmus vanzoi* | 0 | 1 | 0 | 1 |
| *Haemodracon riebeckii* | 0 | 1 | 0 | 1 |
| *Hakaria simonyi* | 0 | 1 | 0 | 1 |
| *Haplocercus ceylonensis* | 0 | 1 | 0 | 1 |
| *Hapsidophrys lineatus* | 0 | 1 | 0 | 1 |
| *Hapsidophrys principis* | 0 | 1 | 0 | 1 |
| *Hapsidophrys smaragdina* | 0 | 1 | 0 | 1 |
| *Helicops angulatus* | 0 | 1 | 0 | 1 |
| *Helicops carinicaudus* | 0 | 1 | 0 | 1 |
| *Helicops gomesi* | 0 | 1 | 0 | 1 |
| *Helicops hagmanni* | 0 | 1 | 0 | 1 |
| *Helicops infrataeniatus* | 0 | 1 | 0 | 1 |
| *Heliobolus lugubris* | 0 | 1 | 0 | 1 |
| *Heliobolus spekii* | 0 | 1 | 0 | 1 |
| *Hellenolacerta graeca* | 1 | 0 | 0 | 0 |
| *Heloderma horridum* | 0 | 1 | 0 | 1 |
| *Heloderma suspectum* | 1 | 0 | 0 | 0 |
| *Hemachatus haemachatus* | 0 | 1 | 0 | 1 |
| *Hemerophis socotrae* | 0 | 1 | 0 | 1 |
| *Hemiaspis damelii* | 0 | 1 | 0 | 1 |
| *Hemiaspis signata* | 0 | 1 | 0 | 1 |
| *Hemibungarus calligaster* | 0 | 1 | 0 | 1 |
| *Hemidactylus aaronbaueri* | 0 | 1 | 0 | 1 |
| *Hemidactylus agrius* | 0 | 1 | 0 | 1 |
| *Hemidactylus albofasciatus* | 0 | 1 | 0 | 1 |
| *Hemidactylus angulatus* | 0 | 1 | 0 | 1 |
| *Hemidactylus bouvieri* | 0 | 1 | 0 | 1 |
| *Hemidactylus bowringii* | 0 | 1 | 0 | 1 |
| *Hemidactylus brasilianus* | 0 | 1 | 0 | 1 |
| *Hemidactylus brookii* | 0 | 1 | 0 | 1 |
| *Hemidactylus citernii* | 0 | 1 | 0 | 1 |
| *Hemidactylus depressus* | 0 | 1 | 0 | 1 |
| *Hemidactylus dracaenacolus* | 0 | 1 | 0 | 1 |
| *Hemidactylus fasciatus* | 0 | 1 | 0 | 1 |
| *Hemidactylus flaviviridis* | 1 | 0 | 0 | 0 |
| *Hemidactylus forbesii* | 0 | 1 | 0 | 1 |
| *Hemidactylus foudaii* | 1 | 0 | 0 | 0 |
| *Hemidactylus frenatus* | 0 | 1 | 0 | 1 |
| *Hemidactylus garnotii* | 0 | 1 | 0 | 1 |
| *Hemidactylus giganteus* | 0 | 1 | 0 | 1 |
| *Hemidactylus gracilis* | 0 | 1 | 0 | 1 |
| *Hemidactylus granti* | 0 | 1 | 0 | 1 |
| *Hemidactylus greefii* | 0 | 1 | 0 | 1 |
| *Hemidactylus haitianus* | 0 | 1 | 0 | 1 |
| *Hemidactylus homoeolepis* | 0 | 1 | 0 | 1 |
| *Hemidactylus karenorum* | 0 | 1 | 0 | 1 |
| *Hemidactylus lemurinus* | 0 | 1 | 0 | 1 |
| *Hemidactylus leschenaultii* | 0 | 1 | 0 | 1 |
| *Hemidactylus longicephalus* | 0 | 1 | 0 | 1 |
| *Hemidactylus mabouia* | 0 | 1 | 0 | 1 |
| *Hemidactylus macropholis* | 0 | 1 | 0 | 1 |
| *Hemidactylus maculatus* | 0 | 1 | 0 | 1 |
| *Hemidactylus mercatorius* | 0 | 1 | 0 | 1 |
| *Hemidactylus mindiae* | 1 | 0 | 0 | 0 |
| *Hemidactylus modestus* | 0 | 1 | 0 | 1 |
| *Hemidactylus oxyrhinus* | 0 | 1 | 0 | 1 |
| *Hemidactylus palaichthus* | 0 | 1 | 0 | 1 |
| *Hemidactylus persicus* | 1 | 0 | 0 | 0 |
| *Hemidactylus platycephalus* | 0 | 1 | 0 | 1 |
| *Hemidactylus platyurus* | 0 | 1 | 0 | 1 |
| *Hemidactylus prashadi* | 0 | 1 | 0 | 1 |
| *Hemidactylus pumilio* | 0 | 1 | 0 | 1 |
| *Hemidactylus reticulatus* | 0 | 1 | 0 | 1 |
| *Hemidactylus robustus* | 1 | 0 | 0 | 0 |
| *Hemidactylus sataraensis* | 0 | 1 | 0 | 1 |
| *Hemidactylus triedrus* | 0 | 1 | 0 | 1 |
| *Hemidactylus turcicus* | 1 | 0 | 0 | 0 |
| *Hemidactylus yerburyi* | 0 | 1 | 0 | 1 |
| *Hemiergis decresiensis* | 0 | 1 | 0 | 1 |
| *Hemiergis initialis* | 0 | 1 | 0 | 1 |
| *Hemiergis millewae* | 0 | 1 | 0 | 1 |
| *Hemiergis peronii* | 0 | 1 | 0 | 1 |
| *Hemiergis quadrilineatum* | 0 | 1 | 0 | 1 |
| *Hemiphyllodactylus aurantiacus* | 0 | 1 | 0 | 1 |
| *Hemiphyllodactylus typus* | 0 | 1 | 0 | 1 |
| *Hemiphyllodactylus yunnanensis* | 0 | 1 | 0 | 1 |
| *Hemirhagerrhis hildebrandtii* | 0 | 1 | 0 | 1 |
| *Hemirhagerrhis kelleri* | 0 | 1 | 0 | 1 |
| *Hemirhagerrhis viperina* | 0 | 1 | 0 | 1 |
| *Hemitheconyx caudicinctus* | 0 | 1 | 0 | 1 |
| *Hemitheconyx taylori* | 0 | 1 | 0 | 1 |
| *Hemorrhois algirus* | 1 | 0 | 0 | 0 |
| *Hemorrhois hippocrepis* | 1 | 0 | 0 | 0 |
| *Hemorrhois nummifer* | 1 | 0 | 0 | 0 |
| *Hemorrhois ravergieri* | 1 | 0 | 0 | 0 |
| *Heterodactylus imbricatus* | 0 | 1 | 0 | 1 |
| *Heterodon nasicus* | 1 | 0 | 0 | 0 |
| *Heterodon platirhinos* | 1 | 0 | 0 | 0 |
| *Heterodon simus* | 1 | 0 | 0 | 0 |
| *Heteroliodon occipitalis* | 0 | 1 | 0 | 1 |
| *Heteronotia binoei* | 0 | 1 | 0 | 1 |
| *Heteronotia planiceps* | 0 | 1 | 0 | 1 |
| *Heteronotia spelea* | 0 | 1 | 0 | 1 |
| *Hierophis gemonensis* | 1 | 0 | 0 | 0 |
| *Hierophis spinalis* | 1 | 0 | 0 | 0 |
| *Hierophis viridiflavus* | 1 | 0 | 0 | 0 |
| *Himalayophis tibetanus* | 1 | 0 | 0 | 0 |
| *Holaspis guentheri* | 0 | 1 | 0 | 1 |
| *Holaspis laevis* | 0 | 1 | 0 | 1 |
| *Holbrookia lacerata* | 1 | 0 | 0 | 0 |
| *Holbrookia maculata* | 1 | 0 | 0 | 0 |
| *Holbrookia propinqua* | 1 | 0 | 0 | 0 |
| *Holodactylus africanus* | 0 | 1 | 0 | 1 |
| *Homalopsis buccata* | 0 | 1 | 0 | 1 |
| *Homonota andicola* | 0 | 0 | 1 | 0 |
| *Homonota borellii* | 0 | 0 | 1 | 0 |
| *Homonota darwinii* | 0 | 0 | 1 | 0 |
| *Homonota fasciata* | 0 | 0 | 1 | 0 |
| *Homonota gaudichaudii* | 0 | 0 | 1 | 0 |
| *Homonota underwoodi* | 0 | 0 | 1 | 0 |
| *Homopholis fasciata* | 0 | 1 | 0 | 1 |
| *Homopholis mulleri* | 0 | 1 | 0 | 1 |
| *Homopholis walbergii* | 0 | 1 | 0 | 1 |
| *Homoroselaps lacteus* | 0 | 1 | 0 | 1 |
| *Hoplocephalus bitorquatus* | 0 | 1 | 0 | 1 |
| *Hoplocercus spinosus* | 0 | 1 | 0 | 1 |
| *Hoplodactylus chrysosireticus* | 0 | 0 | 1 | 0 |
| *Hoplodactylus cryptozoicus* | 0 | 0 | 1 | 0 |
| *Hoplodactylus duvaucelii* | 0 | 0 | 1 | 0 |
| *Hoplodactylus granulatus* | 0 | 0 | 1 | 0 |
| *Hoplodactylus kahutarae* | 0 | 0 | 1 | 0 |
| *Hoplodactylus maculatus* | 0 | 0 | 1 | 0 |
| *Hoplodactylus nebulosus* | 0 | 0 | 1 | 0 |
| *Hoplodactylus pacificus* | 0 | 0 | 1 | 0 |
| *Hoplodactylus rakiurae* | 0 | 0 | 1 | 0 |
| *Hoplodactylus stephensi* | 0 | 0 | 1 | 0 |
| *Hormonotus modestus* | 0 | 1 | 0 | 1 |
| *Hydrelaps darwiniensis* | 0 | 1 | 0 | 1 |
| *Hydrodynastes bicinctus* | 0 | 1 | 0 | 1 |
| *Hydrodynastes gigas* | 0 | 1 | 0 | 1 |
| *Hydromorphus concolor* | 0 | 1 | 0 | 1 |
| *Hydrophis atriceps* | 0 | 1 | 0 | 1 |
| *Hydrophis brooki* | 0 | 1 | 0 | 1 |
| *Hydrophis cyanocinctus* | 0 | 1 | 0 | 1 |
| *Hydrophis czeblukovi* | 0 | 1 | 0 | 1 |
| *Hydrophis elegans* | 0 | 1 | 0 | 1 |
| *Hydrophis lapemoides* | 0 | 1 | 0 | 1 |
| *Hydrophis macdowelli* | 0 | 1 | 0 | 1 |
| *Hydrophis melanocephalus* | 0 | 1 | 0 | 1 |
| *Hydrophis ornatus* | 0 | 1 | 0 | 1 |
| *Hydrophis pacificus* | 0 | 1 | 0 | 1 |
| *Hydrophis parviceps* | 0 | 1 | 0 | 1 |
| *Hydrophis semperi* | 0 | 1 | 0 | 1 |
| *Hydrophis spiralis* | 0 | 1 | 0 | 1 |
| *Hydrops triangularis* | 0 | 1 | 0 | 1 |
| *Hydrosaurus amboinensis* | 0 | 1 | 0 | 1 |
| *Hypnale hypnale* | 0 | 1 | 0 | 1 |
| *Hypnale nepa* | 0 | 1 | 0 | 1 |
| *Hypnale zara* | 0 | 1 | 0 | 1 |
| *Hypsiglena affinis* | 0 | 1 | 0 | 1 |
| *Hypsiglena chlorophaea* | 1 | 0 | 0 | 0 |
| *Hypsiglena jani* | 1 | 0 | 0 | 0 |
| *Hypsiglena ochrorhyncha* | 1 | 0 | 0 | 0 |
| *Hypsiglena slevini* | 0 | 1 | 0 | 1 |
| *Hypsiglena torquata* | 1 | 0 | 0 | 0 |
| *Hypsilurus boydii* | 0 | 1 | 0 | 1 |
| *Hypsilurus bruijnii* | 0 | 1 | 0 | 1 |
| *Hypsilurus dilophus* | 0 | 1 | 0 | 1 |
| *Hypsilurus modestus* | 0 | 1 | 0 | 1 |
| *Hypsilurus nigrigularis* | 0 | 1 | 0 | 1 |
| *Hypsilurus papuensis* | 0 | 1 | 0 | 1 |
| *Hypsilurus spinipes* | 0 | 1 | 0 | 1 |
| *Hypsirhynchus ferox* | 0 | 1 | 0 | 1 |
| *Ialtris dorsalis* | 0 | 1 | 0 | 1 |
| *Iberolacerta aranica* | 1 | 0 | 0 | 0 |
| *Iberolacerta aurelioi* | 1 | 0 | 0 | 0 |
| *Iberolacerta bonnali* | 1 | 0 | 0 | 0 |
| *Iberolacerta cyreni* | 1 | 0 | 0 | 0 |
| *Iberolacerta galani* | 1 | 0 | 0 | 0 |
| *Iberolacerta horvathi* | 1 | 0 | 0 | 0 |
| *Iberolacerta monticola* | 1 | 0 | 0 | 0 |
| *Ichnotropis capensis* | 0 | 1 | 0 | 1 |
| *Ichnotropis squamulosa* | 0 | 1 | 0 | 1 |
| *Iguana delicatissima* | 0 | 1 | 0 | 1 |
| *Iguana iguana* | 0 | 1 | 0 | 1 |
| *Imantodes cenchoa* | 0 | 1 | 0 | 1 |
| *Imantodes gemmistratus* | 0 | 1 | 0 | 1 |
| *Imantodes inornatus* | 0 | 1 | 0 | 1 |
| *Imantodes lentiferus* | 0 | 1 | 0 | 1 |
| *Iphisa elegans* | 0 | 1 | 0 | 1 |
| *Iranolacerta brandtii* | 1 | 0 | 0 | 0 |
| *Iranolacerta zagrosica* | 1 | 0 | 0 | 0 |
| *Isopachys anguinoides* | 0 | 1 | 0 | 1 |
| *Ithycyphus miniatus* | 0 | 1 | 0 | 1 |
| *Ithycyphus oursi* | 0 | 1 | 0 | 1 |
| *Janetaescincus braueri* | 0 | 1 | 0 | 1 |
| *Janetaescincus veseyfitzgeraldi* | 0 | 1 | 0 | 1 |
| *Japalura flaviceps* | 1 | 0 | 0 | 0 |
| *Japalura polygonata* | 0 | 1 | 0 | 1 |
| *Japalura splendida* | 1 | 0 | 0 | 0 |
| *Japalura tricarinata* | 0 | 1 | 0 | 1 |
| *Japalura variegata* | 0 | 1 | 0 | 1 |
| *Kanakysaurus viviparus* | 0 | 1 | 0 | 1 |
| *Kentropyx altamazonica* | 0 | 1 | 0 | 1 |
| *Kentropyx calcarata* | 0 | 1 | 0 | 1 |
| *Kentropyx paulensis* | 0 | 1 | 0 | 1 |
| *Kentropyx pelviceps* | 0 | 1 | 0 | 1 |
| *Kentropyx striata* | 0 | 1 | 0 | 1 |
| *Kentropyx vanzoi* | 0 | 1 | 0 | 1 |
| *Kentropyx viridistriga* | 0 | 1 | 0 | 1 |
| *Kinyongia adolfifriderici* | 0 | 1 | 0 | 1 |
| *Kinyongia boehmei* | 0 | 1 | 0 | 1 |
| *Kinyongia carpenteri* | 0 | 1 | 0 | 1 |
| *Kinyongia excubitor* | 0 | 1 | 0 | 1 |
| *Kinyongia fischeri* | 0 | 1 | 0 | 1 |
| *Kinyongia matschiei* | 0 | 1 | 0 | 1 |
| *Kinyongia multituberculata* | 0 | 1 | 0 | 1 |
| *Kinyongia oxyrhina* | 0 | 1 | 0 | 1 |
| *Kinyongia tavetana* | 0 | 1 | 0 | 1 |
| *Kinyongia tenue* | 0 | 1 | 0 | 1 |
| *Kinyongia uthmoelleri* | 0 | 1 | 0 | 1 |
| *Kinyongia vosseleri* | 0 | 1 | 0 | 1 |
| *Kinyongia xenorhina* | 0 | 1 | 0 | 1 |
| *Lacerta agilis* | 1 | 0 | 0 | 0 |
| *Lacerta bilineata* | 1 | 0 | 0 | 0 |
| *Lacerta media* | 1 | 0 | 0 | 0 |
| *Lacerta pamphylica* | 1 | 0 | 0 | 0 |
| *Lacerta schreiberi* | 1 | 0 | 0 | 0 |
| *Lacerta strigata* | 1 | 0 | 0 | 0 |
| *Lacerta trilineata* | 1 | 0 | 0 | 0 |
| *Lacerta viridis* | 1 | 0 | 0 | 0 |
| *Lacertaspis chriswildi* | 0 | 1 | 0 | 1 |
| *Lacertaspis gemmiventris* | 0 | 1 | 0 | 1 |
| *Lacertaspis lepesmei* | 0 | 1 | 0 | 1 |
| *Lacertaspis reichenowi* | 0 | 1 | 0 | 1 |
| *Lacertaspis rohdei* | 0 | 1 | 0 | 1 |
| *Lacertoides pardalis* | 0 | 1 | 0 | 1 |
| *Lachesis muta* | 0 | 1 | 0 | 1 |
| *Lachesis stenophrys* | 0 | 1 | 0 | 1 |
| *Laemanctus longipes* | 0 | 1 | 0 | 1 |
| *Lamprolepis smaragdina* | 0 | 1 | 0 | 1 |
| *Lampropeltis alterna* | 1 | 0 | 0 | 0 |
| *Lampropeltis californiae* | 1 | 0 | 0 | 0 |
| *Lampropeltis calligaster* | 1 | 0 | 0 | 0 |
| *Lampropeltis elapsoides* | 1 | 0 | 0 | 0 |
| *Lampropeltis extenuata* | 1 | 0 | 0 | 0 |
| *Lampropeltis getula* | 1 | 0 | 0 | 0 |
| *Lampropeltis holbrooki* | 1 | 0 | 0 | 0 |
| *Lampropeltis mexicana* | 1 | 0 | 0 | 0 |
| *Lampropeltis nigra* | 1 | 0 | 0 | 0 |
| *Lampropeltis pyromelana* | 1 | 0 | 0 | 0 |
| *Lampropeltis ruthveni* | 0 | 1 | 0 | 1 |
| *Lampropeltis splendida* | 1 | 0 | 0 | 0 |
| *Lampropeltis triangulum* | 1 | 1 | 0 | 0 |
| *Lampropeltis webbi* | 0 | 1 | 0 | 1 |
| *Lampropeltis zonata* | 1 | 0 | 0 | 0 |
| *Lamprophis aurora* | 0 | 1 | 0 | 1 |
| *Lamprophis fiskii* | 0 | 1 | 0 | 1 |
| *Lamprophis fuliginosus* | 0 | 1 | 0 | 1 |
| *Lamprophis fuscus* | 0 | 1 | 0 | 1 |
| *Lamprophis guttatus* | 0 | 1 | 0 | 1 |
| *Lamprophis inornatus* | 0 | 1 | 0 | 1 |
| *Lamprophis lineatus* | 0 | 1 | 0 | 1 |
| *Lamprophis olivaceus* | 0 | 1 | 0 | 1 |
| *Lamprophis swazicus* | 0 | 1 | 0 | 1 |
| *Lamprophis virgatus* | 0 | 1 | 0 | 1 |
| *Lampropholis coggeri* | 0 | 1 | 0 | 1 |
| *Lampropholis delicata* | 0 | 1 | 0 | 1 |
| *Lampropholis guichenoti* | 0 | 1 | 0 | 1 |
| *Lampropholis robertsi* | 0 | 1 | 0 | 1 |
| *Langaha madagascariensis* | 0 | 1 | 0 | 1 |
| *Lankascincus fallax* | 0 | 1 | 0 | 1 |
| *Lanthanotus borneensis* | 0 | 1 | 0 | 1 |
| *Lapemis curtus* | 0 | 1 | 0 | 1 |
| *Larutia seribuatensis* | 0 | 1 | 0 | 1 |
| *Latastia longicaudata* | 0 | 1 | 0 | 1 |
| *Laticauda colubrina* | 0 | 1 | 0 | 1 |
| *Laticauda guineai* | 0 | 1 | 0 | 1 |
| *Laticauda laticaudata* | 0 | 1 | 0 | 1 |
| *Laticauda saintgironsi* | 0 | 1 | 0 | 1 |
| *Laudakia caucasia* | 1 | 0 | 0 | 0 |
| *Laudakia erythrogastra* | 1 | 0 | 0 | 0 |
| *Laudakia himalayana* | 1 | 0 | 0 | 0 |
| *Laudakia lehmanni* | 1 | 0 | 0 | 0 |
| *Laudakia microlepis* | 1 | 0 | 0 | 0 |
| *Laudakia nupta* | 1 | 0 | 0 | 0 |
| *Laudakia sacra* | 1 | 0 | 0 | 0 |
| *Laudakia stellio* | 1 | 0 | 0 | 0 |
| *Laudakia stoliczkana* | 1 | 0 | 0 | 0 |
| *Laudakia tuberculata* | 1 | 0 | 0 | 0 |
| *Leiocephalus barahonensis* | 0 | 1 | 0 | 1 |
| *Leiocephalus carinatus* | 0 | 1 | 0 | 1 |
| *Leiocephalus personatus* | 0 | 1 | 0 | 1 |
| *Leiocephalus psammodromus* | 0 | 1 | 0 | 1 |
| *Leiocephalus raviceps* | 0 | 1 | 0 | 1 |
| *Leiocephalus schreibersii* | 0 | 1 | 0 | 1 |
| *Leioheterodon geayi* | 0 | 1 | 0 | 1 |
| *Leioheterodon madagascariensis* | 0 | 1 | 0 | 1 |
| *Leioheterodon modestus* | 0 | 1 | 0 | 1 |
| *Leiolepis belliana* | 0 | 1 | 0 | 1 |
| *Leiolepis guentherpetersi* | 0 | 1 | 0 | 1 |
| *Leiolepis guttata* | 0 | 1 | 0 | 1 |
| *Leiolepis reevesii* | 0 | 1 | 0 | 1 |
| *Leiolopisma mauritiana* | 0 | 1 | 0 | 1 |
| *Leiolopisma telfairii* | 0 | 1 | 0 | 1 |
| *Leiopython albertisii* | 0 | 1 | 0 | 1 |
| *Leiosaurus bellii* | 0 | 0 | 1 | 0 |
| *Leiosaurus catamarcensis* | 0 | 0 | 1 | 0 |
| *Leiosaurus paronae* | 0 | 1 | 0 | 1 |
| *Lepidoblepharis festae* | 0 | 1 | 0 | 1 |
| *Lepidoblepharis xanthostigma* | 0 | 1 | 0 | 1 |
| *Lepidodactylus lugubris* | 0 | 1 | 0 | 1 |
| *Lepidodactylus moestus* | 0 | 1 | 0 | 1 |
| *Lepidodactylus novaeguineae* | 0 | 1 | 0 | 1 |
| *Lepidodactylus orientalis* | 0 | 1 | 0 | 1 |
| *Lepidophyma cuicateca* | 0 | 1 | 0 | 1 |
| *Lepidophyma dontomasi* | 0 | 1 | 0 | 1 |
| *Lepidophyma flavimaculatum* | 0 | 1 | 0 | 1 |
| *Lepidophyma gaigeae* | 0 | 1 | 0 | 1 |
| *Lepidophyma lineri* | 0 | 1 | 0 | 1 |
| *Lepidophyma lipetzi* | 0 | 1 | 0 | 1 |
| *Lepidophyma lowei* | 0 | 1 | 0 | 1 |
| *Lepidophyma mayae* | 0 | 1 | 0 | 1 |
| *Lepidophyma micropholis* | 0 | 1 | 0 | 1 |
| *Lepidophyma occulor* | 0 | 1 | 0 | 1 |
| *Lepidophyma pajapanensis* | 0 | 1 | 0 | 1 |
| *Lepidophyma radula* | 0 | 1 | 0 | 1 |
| *Lepidophyma reticulatum* | 0 | 1 | 0 | 1 |
| *Lepidophyma smithii* | 0 | 1 | 0 | 1 |
| *Lepidophyma sylvaticum* | 0 | 1 | 0 | 1 |
| *Lepidophyma tuxtlae* | 0 | 1 | 0 | 1 |
| *Lepidothyris fernandi* | 0 | 1 | 0 | 1 |
| *Leposoma annectans* | 0 | 1 | 0 | 1 |
| *Leposoma baturitensis* | 0 | 1 | 0 | 1 |
| *Leposoma guianense* | 0 | 1 | 0 | 1 |
| *Leposoma nanodactylus* | 0 | 1 | 0 | 1 |
| *Leposoma osvaldoi* | 0 | 1 | 0 | 1 |
| *Leposoma parietale* | 0 | 1 | 0 | 1 |
| *Leposoma percarinatum* | 0 | 1 | 0 | 1 |
| *Leposoma puk* | 0 | 1 | 0 | 1 |
| *Leposoma scincoides* | 0 | 1 | 0 | 1 |
| *Leposoma southi* | 0 | 1 | 0 | 1 |
| *Leposternon infraorbitale* | 0 | 1 | 0 | 1 |
| *Leposternon microcephalum* | 0 | 1 | 0 | 1 |
| *Leposternon polystegum* | 0 | 1 | 0 | 1 |
| *Leptodeira annulata* | 0 | 1 | 0 | 1 |
| *Leptodeira bakeri* | 0 | 1 | 0 | 1 |
| *Leptodeira frenata* | 0 | 1 | 0 | 1 |
| *Leptodeira maculata* | 0 | 1 | 0 | 1 |
| *Leptodeira nigrofasciata* | 0 | 1 | 0 | 1 |
| *Leptodeira punctata* | 0 | 1 | 0 | 1 |
| *Leptodeira rubricata* | 0 | 1 | 0 | 1 |
| *Leptodeira septentrionalis* | 0 | 1 | 0 | 1 |
| *Leptodeira splendida* | 0 | 1 | 0 | 1 |
| *Leptophis ahaetulla* | 0 | 1 | 0 | 1 |
| *Leptosiaphos amieti* | 0 | 1 | 0 | 1 |
| *Leptosiaphos graueri* | 0 | 1 | 0 | 1 |
| *Leptosiaphos hackarsi* | 0 | 1 | 0 | 1 |
| *Leptosiaphos kilimensis* | 0 | 1 | 0 | 1 |
| *Leptosiaphos vigintiserierum* | 0 | 1 | 0 | 1 |
| *Leptotyphlops adleri* | 0 | 1 | 0 | 1 |
| *Leptotyphlops albifrons* | 0 | 1 | 0 | 1 |
| *Leptotyphlops algeriensis* | 0 | 0 | 1 | 0 |
| *Leptotyphlops asbolepis* | 0 | 1 | 0 | 1 |
| *Leptotyphlops bicolor* | 0 | 1 | 0 | 1 |
| *Leptotyphlops blanfordi* | 1 | 0 | 0 | 0 |
| *Leptotyphlops boueti* | 0 | 1 | 0 | 1 |
| *Leptotyphlops breuili* | 0 | 1 | 0 | 1 |
| *Leptotyphlops carlae* | 0 | 1 | 0 | 1 |
| *Leptotyphlops columbi* | 0 | 1 | 0 | 1 |
| *Leptotyphlops conjunctus* | 0 | 1 | 0 | 1 |
| *Leptotyphlops distanti* | 0 | 1 | 0 | 1 |
| *Leptotyphlops dulcis* | 0 | 0 | 1 | 0 |
| *Leptotyphlops goudotii* | 0 | 1 | 0 | 1 |
| *Leptotyphlops humilis* | 0 | 0 | 1 | 0 |
| *Leptotyphlops leptipilepta* | 0 | 1 | 0 | 1 |
| *Leptotyphlops longicaudus* | 0 | 1 | 0 | 1 |
| *Leptotyphlops macrolepis* | 0 | 1 | 0 | 1 |
| *Leptotyphlops macrorhynchus* | 1 | 1 | 0 | 0 |
| *Leptotyphlops nigricans* | 0 | 1 | 0 | 1 |
| *Leptotyphlops nigroterminus* | 0 | 1 | 0 | 1 |
| *Leptotyphlops occidentalis* | 0 | 1 | 0 | 1 |
| *Leptotyphlops pyrites* | 0 | 1 | 0 | 1 |
| *Leptotyphlops rouxestevae* | 0 | 1 | 0 | 1 |
| *Leptotyphlops scutifrons* | 0 | 1 | 0 | 1 |
| *Leptotyphlops septemstriatus* | 0 | 1 | 0 | 1 |
| *Leptotyphlops sylvicolus* | 0 | 1 | 0 | 1 |
| *Lerista aericeps* | 0 | 1 | 0 | 1 |
| *Lerista allochira* | 0 | 1 | 0 | 1 |
| *Lerista ameles* | 0 | 1 | 0 | 1 |
| *Lerista apoda* | 0 | 1 | 0 | 1 |
| *Lerista arenicola* | 0 | 1 | 0 | 1 |
| *Lerista axillaris* | 0 | 1 | 0 | 1 |
| *Lerista baynesi* | 0 | 1 | 0 | 1 |
| *Lerista bipes* | 0 | 1 | 0 | 1 |
| *Lerista borealis* | 0 | 1 | 0 | 1 |
| *Lerista bougainvillii* | 0 | 1 | 0 | 1 |
| *Lerista carpentariae* | 0 | 1 | 0 | 1 |
| *Lerista chordae* | 0 | 1 | 0 | 1 |
| *Lerista christinae* | 0 | 1 | 0 | 1 |
| *Lerista cinerea* | 0 | 1 | 0 | 1 |
| *Lerista connivens* | 0 | 1 | 0 | 1 |
| *Lerista desertorum* | 0 | 1 | 0 | 1 |
| *Lerista distinguenda* | 0 | 1 | 0 | 1 |
| *Lerista dorsalis* | 0 | 1 | 0 | 1 |
| *Lerista edwardsae* | 0 | 1 | 0 | 1 |
| *Lerista elegans* | 0 | 1 | 0 | 1 |
| *Lerista elongata* | 0 | 1 | 0 | 1 |
| *Lerista emmotti* | 0 | 1 | 0 | 1 |
| *Lerista eupoda* | 0 | 1 | 0 | 1 |
| *Lerista flammicauda* | 0 | 1 | 0 | 1 |
| *Lerista fragilis* | 0 | 1 | 0 | 1 |
| *Lerista frosti* | 0 | 1 | 0 | 1 |
| *Lerista gascoynensis* | 0 | 1 | 0 | 1 |
| *Lerista gerrardii* | 0 | 1 | 0 | 1 |
| *Lerista greeri* | 0 | 1 | 0 | 1 |
| *Lerista griffini* | 0 | 1 | 0 | 1 |
| *Lerista haroldi* | 0 | 1 | 0 | 1 |
| *Lerista humphriesi* | 0 | 1 | 0 | 1 |
| *Lerista ingrami* | 0 | 1 | 0 | 1 |
| *Lerista ips* | 0 | 1 | 0 | 1 |
| *Lerista kalumburu* | 0 | 1 | 0 | 1 |
| *Lerista karlschmidti* | 0 | 1 | 0 | 1 |
| *Lerista kendricki* | 0 | 1 | 0 | 1 |
| *Lerista kennedyensis* | 0 | 1 | 0 | 1 |
| *Lerista labialis* | 0 | 1 | 0 | 1 |
| *Lerista lineata* | 0 | 1 | 0 | 1 |
| *Lerista lineopunctulata* | 0 | 1 | 0 | 1 |
| *Lerista macropisthopus* | 0 | 1 | 0 | 1 |
| *Lerista microtis* | 0 | 1 | 0 | 1 |
| *Lerista muelleri* | 0 | 1 | 0 | 1 |
| *Lerista neander* | 0 | 1 | 0 | 1 |
| *Lerista nichollsi* | 0 | 1 | 0 | 1 |
| *Lerista onsloviana* | 0 | 1 | 0 | 1 |
| *Lerista orientalis* | 0 | 1 | 0 | 1 |
| *Lerista petersoni* | 0 | 1 | 0 | 1 |
| *Lerista picturata* | 0 | 1 | 0 | 1 |
| *Lerista planiventralis* | 0 | 1 | 0 | 1 |
| *Lerista praepedita* | 0 | 1 | 0 | 1 |
| *Lerista punctatovittata* | 0 | 1 | 0 | 1 |
| *Lerista puncticauda* | 0 | 1 | 0 | 1 |
| *Lerista robusta* | 0 | 1 | 0 | 1 |
| *Lerista simillima* | 0 | 1 | 0 | 1 |
| *Lerista speciosa* | 0 | 1 | 0 | 1 |
| *Lerista stictopleura* | 0 | 1 | 0 | 1 |
| *Lerista stylis* | 0 | 1 | 0 | 1 |
| *Lerista taeniata* | 0 | 1 | 0 | 1 |
| *Lerista terdigitata* | 0 | 1 | 0 | 1 |
| *Lerista tridactyla* | 0 | 1 | 0 | 1 |
| *Lerista uniduo* | 0 | 1 | 0 | 1 |
| *Lerista varia* | 0 | 1 | 0 | 1 |
| *Lerista vermicularis* | 0 | 1 | 0 | 1 |
| *Lerista viduata* | 0 | 1 | 0 | 1 |
| *Lerista walkeri* | 0 | 1 | 0 | 1 |
| *Lerista wilkinsi* | 0 | 1 | 0 | 1 |
| *Lerista xanthura* | 0 | 1 | 0 | 1 |
| *Lerista yuna* | 0 | 1 | 0 | 1 |
| *Lerista zietzi* | 0 | 1 | 0 | 1 |
| *Lerista zonulata* | 0 | 1 | 0 | 1 |
| *Letheobia obtusa* | 0 | 1 | 0 | 1 |
| *Lialis burtonis* | 0 | 1 | 0 | 1 |
| *Lialis jicari* | 0 | 1 | 0 | 1 |
| *Liasis fuscus* | 0 | 1 | 0 | 1 |
| *Liasis mackloti* | 0 | 1 | 0 | 1 |
| *Liasis olivaceus* | 0 | 1 | 0 | 1 |
| *Lichanura trivirgata* | 0 | 0 | 1 | 0 |
| *Liolaemus abaucan* | 0 | 0 | 1 | 0 |
| *Liolaemus albiceps* | 0 | 0 | 1 | 0 |
| *Liolaemus andinus* | 0 | 0 | 1 | 0 |
| *Liolaemus archeforus* | 0 | 0 | 1 | 0 |
| *Liolaemus atacamensis* | 0 | 0 | 1 | 0 |
| *Liolaemus audituvelatus* | 0 | 0 | 1 | 0 |
| *Liolaemus austromendocinus* | 0 | 0 | 1 | 0 |
| *Liolaemus azarai* | 0 | 0 | 1 | 0 |
| *Liolaemus baguali* | 0 | 0 | 1 | 0 |
| *Liolaemus bellii* | 0 | 0 | 1 | 0 |
| *Liolaemus bibronii* | 0 | 0 | 1 | 0 |
| *Liolaemus bitaeniatus* | 0 | 0 | 1 | 0 |
| *Liolaemus boulengeri* | 0 | 0 | 1 | 0 |
| *Liolaemus buergeri* | 0 | 0 | 1 | 0 |
| *Liolaemus calchaqui* | 0 | 0 | 1 | 0 |
| *Liolaemus canqueli* | 0 | 0 | 1 | 0 |
| *Liolaemus capillitas* | 0 | 0 | 1 | 0 |
| *Liolaemus ceii* | 0 | 0 | 1 | 0 |
| *Liolaemus chacoensis* | 0 | 0 | 1 | 0 |
| *Liolaemus chaltin* | 0 | 0 | 1 | 0 |
| *Liolaemus chehuachekenk* | 0 | 0 | 1 | 0 |
| *Liolaemus chiliensis* | 0 | 0 | 1 | 0 |
| *Liolaemus coeruleus* | 0 | 0 | 1 | 0 |
| *Liolaemus crepuscularis* | 0 | 0 | 1 | 0 |
| *Liolaemus cuyanus* | 0 | 0 | 1 | 0 |
| *Liolaemus cyanogaster* | 0 | 0 | 1 | 0 |
| *Liolaemus darwinii* | 0 | 0 | 1 | 0 |
| *Liolaemus dicktracy* | 0 | 0 | 1 | 0 |
| *Liolaemus donosobarrosi* | 0 | 0 | 1 | 0 |
| *Liolaemus dorbignyi* | 0 | 0 | 1 | 0 |
| *Liolaemus elongatus* | 0 | 0 | 1 | 0 |
| *Liolaemus escarchadosi* | 0 | 0 | 1 | 0 |
| *Liolaemus espinozai* | 0 | 0 | 1 | 0 |
| *Liolaemus fabiani* | 0 | 0 | 1 | 0 |
| *Liolaemus famatinae* | 0 | 0 | 1 | 0 |
| *Liolaemus fitzingerii* | 0 | 0 | 1 | 0 |
| *Liolaemus fuscus* | 0 | 0 | 1 | 0 |
| *Liolaemus gallardoi* | 0 | 0 | 1 | 0 |
| *Liolaemus gracilis* | 0 | 0 | 1 | 0 |
| *Liolaemus gravenhorstii* | 0 | 0 | 1 | 0 |
| *Liolaemus grosseorum* | 0 | 0 | 1 | 0 |
| *Liolaemus hatcheri* | 0 | 0 | 1 | 0 |
| *Liolaemus heliodermis* | 0 | 0 | 1 | 0 |
| *Liolaemus hermannunezi* | 0 | 0 | 1 | 0 |
| *Liolaemus hernani* | 0 | 0 | 1 | 0 |
| *Liolaemus huacahuasicus* | 0 | 0 | 1 | 0 |
| *Liolaemus inacayali* | 0 | 0 | 1 | 0 |
| *Liolaemus irregularis* | 0 | 0 | 1 | 0 |
| *Liolaemus kingii* | 0 | 0 | 1 | 0 |
| *Liolaemus kolengh* | 0 | 0 | 1 | 0 |
| *Liolaemus koslowskyi* | 0 | 0 | 1 | 0 |
| *Liolaemus kriegi* | 0 | 0 | 1 | 0 |
| *Liolaemus laurenti* | 0 | 0 | 1 | 0 |
| *Liolaemus lavillai* | 0 | 0 | 1 | 0 |
| *Liolaemus lemniscatus* | 0 | 0 | 1 | 0 |
| *Liolaemus leopardinus* | 0 | 0 | 1 | 0 |
| *Liolaemus lineomaculatus* | 0 | 0 | 1 | 0 |
| *Liolaemus lutzae* | 0 | 1 | 0 | 1 |
| *Liolaemus magellanicus* | 0 | 0 | 1 | 0 |
| *Liolaemus melanops* | 0 | 0 | 1 | 0 |
| *Liolaemus molinai* | 0 | 0 | 1 | 0 |
| *Liolaemus monticola* | 0 | 0 | 1 | 0 |
| *Liolaemus morenoi* | 0 | 0 | 1 | 0 |
| *Liolaemus multicolor* | 0 | 0 | 1 | 0 |
| *Liolaemus multimaculatus* | 0 | 0 | 1 | 0 |
| *Liolaemus nigromaculatus* | 0 | 0 | 1 | 0 |
| *Liolaemus nigroviridis* | 0 | 0 | 1 | 0 |
| *Liolaemus nitidus* | 0 | 0 | 1 | 0 |
| *Liolaemus occipitalis* | 0 | 0 | 1 | 0 |
| *Liolaemus olongasta* | 0 | 0 | 1 | 0 |
| *Liolaemus orientalis* | 0 | 0 | 1 | 0 |
| *Liolaemus ornatus* | 0 | 0 | 1 | 0 |
| *Liolaemus pagaburoi* | 0 | 0 | 1 | 0 |
| *Liolaemus paulinae* | 0 | 0 | 1 | 0 |
| *Liolaemus petrophilus* | 0 | 0 | 1 | 0 |
| *Liolaemus pictus* | 0 | 0 | 1 | 0 |
| *Liolaemus platei* | 0 | 0 | 1 | 0 |
| *Liolaemus pseudoanomalus* | 0 | 0 | 1 | 0 |
| *Liolaemus pseudolemniscatus* | 0 | 0 | 1 | 0 |
| *Liolaemus puna* | 0 | 0 | 1 | 0 |
| *Liolaemus quilmes* | 0 | 0 | 1 | 0 |
| *Liolaemus ramirezae* | 0 | 0 | 1 | 0 |
| *Liolaemus reichei* | 0 | 0 | 1 | 0 |
| *Liolaemus riojanus* | 0 | 0 | 1 | 0 |
| *Liolaemus robertmertensi* | 0 | 0 | 1 | 0 |
| *Liolaemus rothi* | 0 | 0 | 1 | 0 |
| *Liolaemus ruibali* | 0 | 0 | 1 | 0 |
| *Liolaemus salinicola* | 0 | 0 | 1 | 0 |
| *Liolaemus sarmientoi* | 0 | 0 | 1 | 0 |
| *Liolaemus saxatilis* | 0 | 0 | 1 | 0 |
| *Liolaemus scapularis* | 0 | 0 | 1 | 0 |
| *Liolaemus schroederi* | 0 | 0 | 1 | 0 |
| *Liolaemus scolaroi* | 0 | 0 | 1 | 0 |
| *Liolaemus silvanae* | 0 | 0 | 1 | 0 |
| *Liolaemus somuncurae* | 0 | 0 | 1 | 0 |
| *Liolaemus tari* | 0 | 0 | 1 | 0 |
| *Liolaemus telsen* | 0 | 0 | 1 | 0 |
| *Liolaemus tenuis* | 0 | 0 | 1 | 0 |
| *Liolaemus thermarum* | 0 | 0 | 1 | 0 |
| *Liolaemus tristis* | 0 | 0 | 1 | 0 |
| *Liolaemus umbrifer* | 0 | 0 | 1 | 0 |
| *Liolaemus uptoni* | 0 | 0 | 1 | 0 |
| *Liolaemus uspallatensis* | 0 | 0 | 1 | 0 |
| *Liolaemus vallecurensis* | 0 | 0 | 1 | 0 |
| *Liolaemus walkeri* | 0 | 0 | 1 | 0 |
| *Liolaemus wiegmannii* | 0 | 0 | 1 | 0 |
| *Liolaemus xanthoviridis* | 0 | 0 | 1 | 0 |
| *Liolaemus yanalcu* | 0 | 0 | 1 | 0 |
| *Liolaemus zapallarensis* | 0 | 0 | 1 | 0 |
| *Liolaemus zullyi* | 0 | 0 | 1 | 0 |
| *Liophidium chabaudi* | 0 | 1 | 0 | 1 |
| *Liophidium mayottensis* | 0 | 1 | 0 | 1 |
| *Liophidium rhodogaster* | 0 | 1 | 0 | 1 |
| *Liophidium therezieni* | 0 | 1 | 0 | 1 |
| *Liophidium torquatum* | 0 | 1 | 0 | 1 |
| *Liophidium vaillanti* | 0 | 1 | 0 | 1 |
| *Liophis almadensis* | 0 | 1 | 0 | 1 |
| *Liophis amarali* | 0 | 1 | 0 | 1 |
| *Liophis anomalus* | 0 | 1 | 0 | 1 |
| *Liophis atraventer* | 0 | 1 | 0 | 1 |
| *Liophis breviceps* | 0 | 1 | 0 | 1 |
| *Liophis ceii* | 0 | 0 | 1 | 0 |
| *Liophis elegantissimus* | 0 | 1 | 0 | 1 |
| *Liophis epinephelus* | 0 | 1 | 0 | 1 |
| *Liophis flavifrenatus* | 0 | 1 | 0 | 1 |
| *Liophis jaegeri* | 0 | 1 | 0 | 1 |
| *Liophis juliae* | 0 | 1 | 0 | 1 |
| *Liophis lineatus* | 0 | 1 | 0 | 1 |
| *Liophis meridionalis* | 0 | 1 | 0 | 1 |
| *Liophis miliaris* | 0 | 1 | 0 | 1 |
| *Liophis paucidens* | 0 | 1 | 0 | 1 |
| *Liophis poecilogyrus* | 0 | 1 | 0 | 1 |
| *Liophis reginae* | 0 | 1 | 0 | 1 |
| *Liophis typhlus* | 0 | 1 | 0 | 1 |
| *Liopholidophis dimorphus* | 0 | 1 | 0 | 1 |
| *Liopholidophis dolicocercus* | 0 | 1 | 0 | 1 |
| *Liopholidophis sexlineatus* | 0 | 1 | 0 | 1 |
| *Lioscincus maruia* | 0 | 1 | 0 | 1 |
| *Lioscincus nigrofasciolatum* | 0 | 1 | 0 | 1 |
| *Lioscincus novaecaledoniae* | 0 | 1 | 0 | 1 |
| *Lioscincus steindachneri* | 0 | 1 | 0 | 1 |
| *Lioscincus tillieri* | 0 | 1 | 0 | 1 |
| *Lioscincus vivae* | 0 | 1 | 0 | 1 |
| *Liotyphlops albirostris* | 0 | 1 | 0 | 1 |
| *Lipinia noctua* | 0 | 1 | 0 | 1 |
| *Lipinia pulchella* | 0 | 1 | 0 | 1 |
| *Lipinia vittigera* | 0 | 1 | 0 | 1 |
| *Lophognathus gilberti* | 0 | 1 | 0 | 1 |
| *Lophognathus longirostris* | 0 | 1 | 0 | 1 |
| *Lophognathus temporalis* | 0 | 1 | 0 | 1 |
| *Loxocemus bicolor* | 0 | 1 | 0 | 1 |
| *Lucasium alboguttatum* | 0 | 1 | 0 | 1 |
| *Lucasium byrnei* | 0 | 1 | 0 | 1 |
| *Lucasium damaeum* | 0 | 1 | 0 | 1 |
| *Lucasium immaculatum* | 0 | 1 | 0 | 1 |
| *Lucasium maini* | 0 | 1 | 0 | 1 |
| *Lucasium squarrosum* | 0 | 1 | 0 | 1 |
| *Lucasium steindachneri* | 0 | 1 | 0 | 1 |
| *Lucasium stenodactylum* | 0 | 1 | 0 | 1 |
| *Lucasium wombeyi* | 0 | 1 | 0 | 1 |
| *Luperosaurus cumingii* | 0 | 1 | 0 | 1 |
| *Luperosaurus iskandari* | 0 | 1 | 0 | 1 |
| *Luperosaurus joloensis* | 0 | 1 | 0 | 1 |
| *Luperosaurus macgregori* | 0 | 1 | 0 | 1 |
| *Lycodon aulicus* | 0 | 1 | 0 | 1 |
| *Lycodon capucinus* | 0 | 1 | 0 | 1 |
| *Lycodon fasciatus* | 0 | 1 | 0 | 1 |
| *Lycodon laoensis* | 0 | 1 | 0 | 1 |
| *Lycodon osmanhilli* | 0 | 1 | 0 | 1 |
| *Lycodon paucifasciatus* | 0 | 1 | 0 | 1 |
| *Lycodon ruhstrati* | 0 | 1 | 0 | 1 |
| *Lycodon zawi* | 0 | 1 | 0 | 1 |
| *Lycodonomorphus laevissimus* | 0 | 1 | 0 | 1 |
| *Lycodonomorphus rufulus* | 0 | 1 | 0 | 1 |
| *Lycodonomorphus whytii* | 0 | 1 | 0 | 1 |
| *Lycodryas sanctijohannis* | 0 | 1 | 0 | 1 |
| *Lycognathophis seychellensis* | 0 | 1 | 0 | 1 |
| *Lycophidion capense* | 0 | 1 | 0 | 1 |
| *Lycophidion laterale* | 0 | 1 | 0 | 1 |
| *Lycophidion nigromaculatum* | 0 | 1 | 0 | 1 |
| *Lycophidion ornatum* | 0 | 1 | 0 | 1 |
| *Lygisaurus abscondita* | 0 | 1 | 0 | 1 |
| *Lygisaurus aeratus* | 0 | 1 | 0 | 1 |
| *Lygisaurus foliorum* | 0 | 1 | 0 | 1 |
| *Lygisaurus laevis* | 0 | 1 | 0 | 1 |
| *Lygisaurus macfarlani* | 0 | 1 | 0 | 1 |
| *Lygisaurus malleolus* | 0 | 1 | 0 | 1 |
| *Lygisaurus novaeguineae* | 0 | 1 | 0 | 1 |
| *Lygisaurus parrhasius* | 0 | 1 | 0 | 1 |
| *Lygisaurus sesbrauna* | 0 | 1 | 0 | 1 |
| *Lygisaurus tanneri* | 0 | 1 | 0 | 1 |
| *Lygodactylus angularis* | 0 | 1 | 0 | 1 |
| *Lygodactylus arnoulti* | 0 | 1 | 0 | 1 |
| *Lygodactylus blancae* | 0 | 1 | 0 | 1 |
| *Lygodactylus bradfieldi* | 0 | 1 | 0 | 1 |
| *Lygodactylus capensis* | 0 | 1 | 0 | 1 |
| *Lygodactylus chobiensis* | 0 | 1 | 0 | 1 |
| *Lygodactylus conraui* | 0 | 1 | 0 | 1 |
| *Lygodactylus expectatus* | 0 | 1 | 0 | 1 |
| *Lygodactylus gravis* | 0 | 1 | 0 | 1 |
| *Lygodactylus guibei* | 0 | 1 | 0 | 1 |
| *Lygodactylus gutturalis* | 0 | 1 | 0 | 1 |
| *Lygodactylus heterurus* | 0 | 1 | 0 | 1 |
| *Lygodactylus keniensis* | 0 | 1 | 0 | 1 |
| *Lygodactylus kimhowelli* | 0 | 1 | 0 | 1 |
| *Lygodactylus klugei* | 0 | 1 | 0 | 1 |
| *Lygodactylus lawrencei* | 0 | 1 | 0 | 1 |
| *Lygodactylus luteopicturatus* | 0 | 1 | 0 | 1 |
| *Lygodactylus madagascariensis* | 0 | 1 | 0 | 1 |
| *Lygodactylus miops* | 0 | 1 | 0 | 1 |
| *Lygodactylus mirabilis* | 0 | 1 | 0 | 1 |
| *Lygodactylus montanus* | 0 | 1 | 0 | 1 |
| *Lygodactylus pauliani* | 0 | 1 | 0 | 1 |
| *Lygodactylus picturatus* | 0 | 1 | 0 | 1 |
| *Lygodactylus pictus* | 0 | 1 | 0 | 1 |
| *Lygodactylus rarus* | 0 | 1 | 0 | 1 |
| *Lygodactylus stevensoni* | 0 | 1 | 0 | 1 |
| *Lygodactylus thomensis* | 0 | 1 | 0 | 1 |
| *Lygodactylus tolampyae* | 0 | 1 | 0 | 1 |
| *Lygodactylus tuberosus* | 0 | 1 | 0 | 1 |
| *Lygodactylus verticillatus* | 0 | 1 | 0 | 1 |
| *Lygodactylus williamsi* | 0 | 1 | 0 | 1 |
| *Lygosoma afrum* | 0 | 1 | 0 | 1 |
| *Lygosoma albopunctata* | 0 | 1 | 0 | 1 |
| *Lygosoma bowringii* | 0 | 1 | 0 | 1 |
| *Lygosoma koratense* | 0 | 1 | 0 | 1 |
| *Lygosoma lineolatum* | 0 | 1 | 0 | 1 |
| *Lygosoma punctata* | 0 | 1 | 0 | 1 |
| *Lygosoma quadrupes* | 0 | 1 | 0 | 1 |
| *Lyriocephalus scutatus* | 0 | 1 | 0 | 1 |
| *Lystrophis dorbignyi* | 0 | 1 | 0 | 1 |
| *Lystrophis histricus* | 0 | 1 | 0 | 1 |
| *Lystrophis matogrossensis* | 0 | 1 | 0 | 1 |
| *Lystrophis nattereri* | 0 | 1 | 0 | 1 |
| *Lystrophis pulcher* | 0 | 1 | 0 | 1 |
| *Lystrophis semicinctus* | 0 | 1 | 0 | 1 |
| *Lytorhynchus diadema* | 1 | 0 | 0 | 0 |
| *Mabuya agilis* | 0 | 1 | 0 | 1 |
| *Mabuya agmosticha* | 0 | 1 | 0 | 1 |
| *Mabuya altamazonica* | 0 | 1 | 0 | 1 |
| *Mabuya bistriata* | 0 | 1 | 0 | 1 |
| *Mabuya caissara* | 0 | 1 | 0 | 1 |
| *Mabuya carvalhoi* | 0 | 1 | 0 | 1 |
| *Mabuya cochabambae* | 0 | 1 | 0 | 1 |
| *Mabuya croizati* | 0 | 1 | 0 | 1 |
| *Mabuya dorsivittata* | 0 | 1 | 0 | 1 |
| *Mabuya falconensis* | 0 | 1 | 0 | 1 |
| *Mabuya frenata* | 0 | 1 | 0 | 1 |
| *Mabuya guaporicola* | 0 | 1 | 0 | 1 |
| *Mabuya heathi* | 0 | 1 | 0 | 1 |
| *Mabuya mabouya* | 0 | 1 | 0 | 1 |
| *Mabuya macrophthalma* | 0 | 1 | 0 | 1 |
| *Mabuya macrorhyncha* | 0 | 1 | 0 | 1 |
| *Mabuya meridensis* | 0 | 1 | 0 | 1 |
| *Mabuya nigropalmata* | 0 | 1 | 0 | 1 |
| *Mabuya nigropunctata* | 0 | 1 | 0 | 1 |
| *Mabuya sloanii* | 0 | 1 | 0 | 1 |
| *Mabuya unimarginata* | 0 | 1 | 0 | 1 |
| *Macrelaps microlepidotus* | 0 | 1 | 0 | 1 |
| *Macropisthodon rudis* | 0 | 1 | 0 | 1 |
| *Macroprotodon abubakeri* | 1 | 0 | 0 | 0 |
| *Macroprotodon cucullatus* | 1 | 0 | 0 | 0 |
| *Macroscincus coctei* | 0 | 1 | 0 | 1 |
| *Macrovipera deserti* | 1 | 0 | 0 | 0 |
| *Macrovipera lebetina* | 1 | 0 | 0 | 0 |
| *Macrovipera mauritanica* | 1 | 0 | 0 | 0 |
| *Macrovipera schweizeri* | 1 | 0 | 0 | 0 |
| *Maculophis bella* | 0 | 1 | 0 | 1 |
| *Madagascarophis colubrinus* | 0 | 1 | 0 | 1 |
| *Madagascarophis meridionalis* | 0 | 1 | 0 | 1 |
| *Madascincus igneocaudatus* | 0 | 1 | 0 | 1 |
| *Madascincus intermedius* | 0 | 1 | 0 | 1 |
| *Madascincus melanopleura* | 0 | 1 | 0 | 1 |
| *Madascincus mouroundavae* | 0 | 1 | 0 | 1 |
| *Madascincus stumpffi* | 0 | 1 | 0 | 1 |
| *Malpolon moilensis* | 1 | 0 | 0 | 0 |
| *Malpolon monspessulanus* | 1 | 0 | 0 | 0 |
| *Manolepis putnami* | 0 | 1 | 0 | 1 |
| *Mantheyus phuwuanensis* | 0 | 1 | 0 | 1 |
| *Marmorosphax montana* | 0 | 1 | 0 | 1 |
| *Marmorosphax tricolor* | 0 | 1 | 0 | 1 |
| *Masticophis flagellum* | 1 | 0 | 0 | 0 |
| *Masticophis taeniatus* | 1 | 0 | 0 | 0 |
| *Mastigodryas bifossatus* | 0 | 1 | 0 | 1 |
| *Mastigodryas boddaerti* | 0 | 1 | 0 | 1 |
| *Mastigodryas melanolomus* | 0 | 1 | 0 | 1 |
| *Matoatoa brevipes* | 0 | 1 | 0 | 1 |
| *Mehelya capensis* | 0 | 1 | 0 | 1 |
| *Mehelya nyassae* | 0 | 1 | 0 | 1 |
| *Mehelya poensis* | 0 | 1 | 0 | 1 |
| *Mehelya stenophthalmus* | 0 | 1 | 0 | 1 |
| *Melanophidium punctatum* | 0 | 1 | 0 | 1 |
| *Melanoseps ater* | 0 | 1 | 0 | 1 |
| *Melanoseps loveridgei* | 0 | 1 | 0 | 1 |
| *Melanoseps occidentalis* | 0 | 1 | 0 | 1 |
| *Menetia alanae* | 0 | 1 | 0 | 1 |
| *Menetia greyii* | 0 | 1 | 0 | 1 |
| *Menetia timlowi* | 0 | 1 | 0 | 1 |
| *Meroles anchietae* | 0 | 1 | 0 | 1 |
| *Meroles ctenodactylus* | 0 | 1 | 0 | 1 |
| *Meroles cuneirostris* | 0 | 1 | 0 | 1 |
| *Meroles knoxii* | 0 | 1 | 0 | 1 |
| *Meroles micropholidotus* | 0 | 1 | 0 | 1 |
| *Meroles reticulatus* | 0 | 1 | 0 | 1 |
| *Meroles suborbitalis* | 0 | 1 | 0 | 1 |
| *Mesalina adramitana* | 1 | 0 | 0 | 0 |
| *Mesalina bahaeldini* | 1 | 0 | 0 | 0 |
| *Mesalina balfouri* | 0 | 1 | 0 | 1 |
| *Mesalina brevirostris* | 1 | 0 | 0 | 0 |
| *Mesalina guttulata* | 1 | 0 | 0 | 0 |
| *Mesalina olivieri* | 1 | 0 | 0 | 0 |
| *Mesalina rubropunctata* | 1 | 0 | 0 | 0 |
| *Mesalina simoni* | 1 | 0 | 0 | 0 |
| *Mesaspis gadovii* | 0 | 1 | 0 | 1 |
| *Mesaspis moreletii* | 0 | 1 | 0 | 1 |
| *Mesoscincus managuae* | 0 | 1 | 0 | 1 |
| *Mesoscincus schwartzei* | 0 | 1 | 0 | 1 |
| *Micrablepharus atticolus* | 0 | 1 | 0 | 1 |
| *Micrablepharus maximiliani* | 0 | 1 | 0 | 1 |
| *Micrelaps bicoloratus* | 0 | 1 | 0 | 1 |
| *Microacontias lineatus* | 0 | 1 | 0 | 1 |
| *Microacontias litoralis* | 0 | 1 | 0 | 1 |
| *Microlophus albemarlensis* | 0 | 1 | 0 | 1 |
| *Microlophus atacamensis* | 0 | 0 | 1 | 0 |
| *Microlophus bivittatus* | 0 | 1 | 0 | 1 |
| *Microlophus delanonis* | 0 | 1 | 0 | 1 |
| *Microlophus duncanensis* | 0 | 1 | 0 | 1 |
| *Microlophus grayii* | 0 | 1 | 0 | 1 |
| *Microlophus habelii* | 0 | 1 | 0 | 1 |
| *Microlophus heterolepis* | 0 | 1 | 0 | 1 |
| *Microlophus koepckeorum* | 0 | 1 | 0 | 1 |
| *Microlophus occipitalis* | 0 | 1 | 0 | 1 |
| *Microlophus pacificus* | 0 | 1 | 0 | 1 |
| *Microlophus peruvianus* | 0 | 1 | 0 | 1 |
| *Microlophus quadrivittatus* | 0 | 1 | 0 | 1 |
| *Microlophus stolzmanni* | 0 | 1 | 0 | 1 |
| *Microlophus theresiae* | 0 | 1 | 0 | 1 |
| *Microlophus theresioides* | 0 | 0 | 1 | 0 |
| *Microlophus thoracicus* | 0 | 1 | 0 | 1 |
| *Microlophus tigris* | 0 | 1 | 0 | 1 |
| *Microlophus yanezi* | 0 | 0 | 1 | 0 |
| *Micropechis ikaheka* | 0 | 1 | 0 | 1 |
| *Micropisthodon ochraceus* | 0 | 1 | 0 | 1 |
| *Micruroides euryxanthus* | 1 | 0 | 0 | 0 |
| *Micrurus albicinctus* | 0 | 1 | 0 | 1 |
| *Micrurus altirostris* | 0 | 1 | 0 | 1 |
| *Micrurus baliocoryphus* | 0 | 1 | 0 | 1 |
| *Micrurus brasiliensis* | 0 | 1 | 0 | 1 |
| *Micrurus corallinus* | 0 | 1 | 0 | 1 |
| *Micrurus decoratus* | 0 | 1 | 0 | 1 |
| *Micrurus diastema* | 0 | 1 | 0 | 1 |
| *Micrurus dissoleucus* | 0 | 1 | 0 | 1 |
| *Micrurus frontalis* | 0 | 1 | 0 | 1 |
| *Micrurus fulvius* | 1 | 0 | 0 | 0 |
| *Micrurus hemprichii* | 0 | 1 | 0 | 1 |
| *Micrurus ibiboboca* | 0 | 1 | 0 | 1 |
| *Micrurus lemniscatus* | 0 | 1 | 0 | 1 |
| *Micrurus mipartitus* | 0 | 1 | 0 | 1 |
| *Micrurus narduccii* | 0 | 1 | 0 | 1 |
| *Micrurus psyches* | 0 | 1 | 0 | 1 |
| *Micrurus pyrrhocryptus* | 0 | 1 | 0 | 1 |
| *Micrurus spixii* | 0 | 1 | 0 | 1 |
| *Micrurus surinamensis* | 0 | 1 | 0 | 1 |
| *Mimophis mahfalensis* | 0 | 1 | 0 | 1 |
| *Mochlus brevicaudis* | 0 | 1 | 0 | 1 |
| *Mochlus sundevalli* | 0 | 1 | 0 | 1 |
| *Moloch horridus* | 0 | 1 | 0 | 1 |
| *Monopeltis capensis* | 0 | 1 | 0 | 1 |
| *Morelia amethistina* | 0 | 1 | 0 | 1 |
| *Morelia boeleni* | 0 | 1 | 0 | 1 |
| *Morelia bredli* | 0 | 1 | 0 | 1 |
| *Morelia carinata* | 0 | 1 | 0 | 1 |
| *Morelia oenpelliensis* | 0 | 1 | 0 | 1 |
| *Morelia spilota* | 0 | 1 | 0 | 1 |
| *Morelia viridis* | 0 | 1 | 0 | 1 |
| *Morethia adelaidensis* | 0 | 1 | 0 | 1 |
| *Morethia butleri* | 0 | 1 | 0 | 1 |
| *Morethia ruficauda* | 0 | 1 | 0 | 1 |
| *Morunasaurus annularis* | 0 | 1 | 0 | 1 |
| *Myron richardsonii* | 0 | 1 | 0 | 1 |
| *Nactus acutus* | 0 | 1 | 0 | 1 |
| *Nactus cheverti* | 0 | 1 | 0 | 1 |
| *Nactus eboracensis* | 0 | 1 | 0 | 1 |
| *Nactus galgajuga* | 0 | 1 | 0 | 1 |
| *Nactus multicarinatus* | 0 | 1 | 0 | 1 |
| *Nactus pelagicus* | 0 | 1 | 0 | 1 |
| *Nactus vankampeni* | 0 | 1 | 0 | 1 |
| *Nadzikambia mlanjense* | 0 | 1 | 0 | 1 |
| *Naja annulata* | 0 | 1 | 0 | 1 |
| *Naja annulifera* | 0 | 1 | 0 | 1 |
| *Naja ashei* | 0 | 1 | 0 | 1 |
| *Naja atra* | 0 | 1 | 0 | 1 |
| *Naja haje* | 0 | 1 | 0 | 1 |
| *Naja kaouthia* | 0 | 1 | 0 | 1 |
| *Naja katiensis* | 0 | 1 | 0 | 1 |
| *Naja mandalayensis* | 0 | 1 | 0 | 1 |
| *Naja melanoleuca* | 0 | 1 | 0 | 1 |
| *Naja mossambica* | 0 | 1 | 0 | 1 |
| *Naja multifasciata* | 0 | 1 | 0 | 1 |
| *Naja naja* | 0 | 1 | 0 | 1 |
| *Naja nigricollis* | 0 | 1 | 0 | 1 |
| *Naja nivea* | 0 | 1 | 0 | 1 |
| *Naja nubiae* | 1 | 0 | 0 | 0 |
| *Naja pallida* | 0 | 1 | 0 | 1 |
| *Naja siamensis* | 0 | 1 | 0 | 1 |
| *Naja sumatrana* | 0 | 1 | 0 | 1 |
| *Nangura spinosa* | 0 | 1 | 0 | 1 |
| *Nannoscincus garrulus* | 0 | 1 | 0 | 1 |
| *Nannoscincus gracilis* | 0 | 1 | 0 | 1 |
| *Nannoscincus greeri* | 0 | 1 | 0 | 1 |
| *Nannoscincus hanchisteus* | 0 | 1 | 0 | 1 |
| *Nannoscincus humectus* | 0 | 1 | 0 | 1 |
| *Nannoscincus mariei* | 0 | 1 | 0 | 1 |
| *Nannoscincus slevini* | 0 | 1 | 0 | 1 |
| *Narudasia festiva* | 0 | 1 | 0 | 1 |
| *Natriciteres olivacea* | 0 | 1 | 0 | 1 |
| *Natrix maura* | 1 | 0 | 0 | 0 |
| *Natrix natrix* | 1 | 0 | 0 | 0 |
| *Natrix tessellata* | 1 | 0 | 0 | 0 |
| *Naultinus elegans* | 0 | 0 | 1 | 0 |
| *Naultinus gemmeus* | 0 | 0 | 1 | 0 |
| *Naultinus grayii* | 0 | 0 | 1 | 0 |
| *Naultinus manukanus* | 0 | 0 | 1 | 0 |
| *Naultinus poecilochlorus* | 0 | 0 | 1 | 0 |
| *Naultinus rudis* | 0 | 0 | 1 | 0 |
| *Naultinus stellatus* | 0 | 0 | 1 | 0 |
| *Naultinus tuberculatus* | 0 | 0 | 1 | 0 |
| *Nephrurus amyae* | 0 | 1 | 0 | 1 |
| *Nephrurus asper* | 0 | 1 | 0 | 1 |
| *Nephrurus deleani* | 0 | 1 | 0 | 1 |
| *Nephrurus laevissimus* | 0 | 1 | 0 | 1 |
| *Nephrurus levis* | 0 | 1 | 0 | 1 |
| *Nephrurus sheai* | 0 | 1 | 0 | 1 |
| *Nephrurus stellatus* | 0 | 1 | 0 | 1 |
| *Nephrurus vertebralis* | 0 | 1 | 0 | 1 |
| *Nephrurus wheeleri* | 0 | 1 | 0 | 1 |
| *Nerodia cyclopion* | 1 | 0 | 0 | 0 |
| *Nerodia erythrogaster* | 1 | 0 | 0 | 0 |
| *Nerodia fasciata* | 1 | 0 | 0 | 0 |
| *Nerodia floridana* | 1 | 0 | 0 | 0 |
| *Nerodia harteri* | 1 | 0 | 0 | 0 |
| *Nerodia rhombifer* | 1 | 1 | 0 | 0 |
| *Nerodia sipedon* | 1 | 0 | 0 | 0 |
| *Nerodia taxispilota* | 1 | 0 | 0 | 0 |
| *Neusticurus bicarinatus* | 0 | 1 | 0 | 1 |
| *Neusticurus rudis* | 0 | 1 | 0 | 1 |
| *Ninia atrata* | 0 | 1 | 0 | 1 |
| *Niveoscincus greeni* | 0 | 1 | 0 | 1 |
| *Niveoscincus metallicus* | 0 | 1 | 0 | 1 |
| *Niveoscincus ocellatus* | 0 | 1 | 0 | 1 |
| *Niveoscincus pretiosus* | 0 | 1 | 0 | 1 |
| *Notechis scutatus* | 0 | 1 | 0 | 1 |
| *Nothobachia ablephara* | 0 | 1 | 0 | 1 |
| *Nothopsis rugosus* | 0 | 1 | 0 | 1 |
| *Notoscincus ornatus* | 0 | 1 | 0 | 1 |
| *Nucras lalandii* | 0 | 1 | 0 | 1 |
| *Nucras tessellata* | 0 | 1 | 0 | 1 |
| *Oedodera marmorata* | 0 | 1 | 0 | 1 |
| *Oedura castelnaui* | 0 | 1 | 0 | 1 |
| *Oedura coggeri* | 0 | 1 | 0 | 1 |
| *Oedura filicipoda* | 0 | 1 | 0 | 1 |
| *Oedura gemmata* | 0 | 1 | 0 | 1 |
| *Oedura gracilis* | 0 | 1 | 0 | 1 |
| *Oedura lesueurii* | 0 | 1 | 0 | 1 |
| *Oedura marmorata* | 0 | 1 | 0 | 1 |
| *Oedura monilis* | 0 | 1 | 0 | 1 |
| *Oedura obscura* | 0 | 1 | 0 | 1 |
| *Oedura reticulata* | 0 | 1 | 0 | 1 |
| *Oedura rhombifer* | 0 | 1 | 0 | 1 |
| *Oedura robusta* | 0 | 1 | 0 | 1 |
| *Oedura tryoni* | 0 | 1 | 0 | 1 |
| *Oligodon arnensis* | 0 | 1 | 0 | 1 |
| *Oligodon barroni* | 0 | 1 | 0 | 1 |
| *Oligodon chinensis* | 0 | 1 | 0 | 1 |
| *Oligodon cinereus* | 0 | 1 | 0 | 1 |
| *Oligodon cruentatus* | 0 | 1 | 0 | 1 |
| *Oligodon cyclurus* | 0 | 1 | 0 | 1 |
| *Oligodon formosanus* | 0 | 1 | 0 | 1 |
| *Oligodon maculatus* | 0 | 1 | 0 | 1 |
| *Oligodon ocellatus* | 0 | 1 | 0 | 1 |
| *Oligodon octolineatus* | 0 | 1 | 0 | 1 |
| *Oligodon planiceps* | 0 | 1 | 0 | 1 |
| *Oligodon splendidus* | 0 | 1 | 0 | 1 |
| *Oligodon sublineatus* | 0 | 1 | 0 | 1 |
| *Oligodon taeniatus* | 0 | 1 | 0 | 1 |
| *Oligodon taeniolatus* | 1 | 0 | 0 | 0 |
| *Oligodon theobaldi* | 0 | 1 | 0 | 1 |
| *Oligodon torquatus* | 0 | 1 | 0 | 1 |
| *Oligosoma acrinasum* | 0 | 0 | 1 | 0 |
| *Oligosoma chloronoton* | 0 | 0 | 1 | 0 |
| *Oligosoma fallai* | 0 | 0 | 1 | 0 |
| *Oligosoma grande* | 0 | 0 | 1 | 0 |
| *Oligosoma homalonotum* | 0 | 0 | 1 | 0 |
| *Oligosoma inconspicuum* | 0 | 0 | 1 | 0 |
| *Oligosoma infrapunctatum* | 0 | 0 | 1 | 0 |
| *Oligosoma lineoocellatum* | 0 | 0 | 1 | 0 |
| *Oligosoma longipes* | 0 | 0 | 1 | 0 |
| *Oligosoma maccanni* | 0 | 0 | 1 | 0 |
| *Oligosoma microlepis* | 0 | 0 | 1 | 0 |
| *Oligosoma moco* | 0 | 0 | 1 | 0 |
| *Oligosoma nigriplantare* | 0 | 0 | 1 | 0 |
| *Oligosoma notosaurus* | 0 | 0 | 1 | 0 |
| *Oligosoma otagense* | 0 | 0 | 1 | 0 |
| *Oligosoma pikitanga* | 0 | 0 | 1 | 0 |
| *Oligosoma smithi* | 0 | 0 | 1 | 0 |
| *Oligosoma stenotis* | 0 | 0 | 1 | 0 |
| *Oligosoma striatum* | 0 | 0 | 1 | 0 |
| *Oligosoma suteri* | 0 | 0 | 1 | 0 |
| *Oligosoma taumakae* | 0 | 0 | 1 | 0 |
| *Oligosoma waimatense* | 0 | 0 | 1 | 0 |
| *Oligosoma zelandicum* | 0 | 0 | 1 | 0 |
| *Omanosaura cyanura* | 0 | 1 | 0 | 1 |
| *Omanosaura jayakari* | 0 | 1 | 0 | 1 |
| *Opheodrys aestivus* | 1 | 0 | 0 | 0 |
| *Opheodrys vernalis* | 1 | 0 | 0 | 0 |
| *Ophidiocephalus taeniatus* | 0 | 1 | 0 | 1 |
| *Ophiodes striatus* | 0 | 1 | 0 | 1 |
| *Ophiomorus latastii* | 1 | 0 | 0 | 0 |
| *Ophiomorus punctatissimus* | 1 | 0 | 0 | 0 |
| *Ophiophagus hannah* | 0 | 1 | 0 | 1 |
| *Ophioscincus ophioscincus* | 0 | 1 | 0 | 1 |
| *Ophioscincus truncatus* | 0 | 1 | 0 | 1 |
| *Ophisaurus attenuatus* | 1 | 0 | 0 | 0 |
| *Ophisaurus gracilis* | 0 | 1 | 0 | 1 |
| *Ophisaurus harti* | 0 | 1 | 0 | 1 |
| *Ophisaurus koellikeri* | 1 | 0 | 0 | 0 |
| *Ophisaurus ventralis* | 1 | 0 | 0 | 0 |
| *Ophisops elegans* | 1 | 0 | 0 | 0 |
| *Ophisops occidentalis* | 1 | 0 | 0 | 0 |
| *Ophryacus melanurus* | 0 | 1 | 0 | 1 |
| *Ophryacus undulatus* | 0 | 1 | 0 | 1 |
| *Opisthotropis cheni* | 0 | 1 | 0 | 1 |
| *Opisthotropis guangxiensis* | 0 | 1 | 0 | 1 |
| *Opisthotropis lateralis* | 0 | 1 | 0 | 1 |
| *Opisthotropis latouchii* | 0 | 1 | 0 | 1 |
| *Oplurus cuvieri* | 0 | 1 | 0 | 1 |
| *Oplurus cyclurus* | 0 | 1 | 0 | 1 |
| *Oplurus fierinensis* | 0 | 1 | 0 | 1 |
| *Oplurus grandidieri* | 0 | 1 | 0 | 1 |
| *Oplurus quadrimaculatus* | 0 | 1 | 0 | 1 |
| *Oplurus saxicola* | 0 | 1 | 0 | 1 |
| *Oreocryptophis porphyracea* | 0 | 1 | 0 | 1 |
| *Orraya occultus* | 0 | 1 | 0 | 1 |
| *Orthriophis cantoris* | 0 | 1 | 0 | 1 |
| *Orthriophis hodgsoni* | 1 | 0 | 0 | 0 |
| *Orthriophis moellendorffi* | 0 | 1 | 0 | 1 |
| *Orthriophis taeniurus* | 0 | 1 | 0 | 1 |
| *Otocryptis wiegmanni* | 0 | 1 | 0 | 1 |
| *Ovophis monticola* | 0 | 1 | 0 | 1 |
| *Ovophis okinavensis* | 0 | 1 | 0 | 1 |
| *Ovophis tonkinensis* | 0 | 1 | 0 | 1 |
| *Ovophis zayuensis* | 0 | 1 | 0 | 1 |
| *Oxybelis aeneus* | 0 | 1 | 0 | 1 |
| *Oxybelis fulgidus* | 0 | 1 | 0 | 1 |
| *Oxyrhabdium leporinum* | 0 | 1 | 0 | 1 |
| *Oxyrhopus clathratus* | 0 | 1 | 0 | 1 |
| *Oxyrhopus formosus* | 0 | 1 | 0 | 1 |
| *Oxyrhopus guibei* | 0 | 1 | 0 | 1 |
| *Oxyrhopus melanogenys* | 0 | 1 | 0 | 1 |
| *Oxyrhopus petola* | 0 | 1 | 0 | 1 |
| *Oxyrhopus rhombifer* | 0 | 1 | 0 | 1 |
| *Oxyrhopus trigeminus* | 0 | 1 | 0 | 1 |
| *Oxyuranus microlepidotus* | 0 | 1 | 0 | 1 |
| *Oxyuranus scutellatus* | 0 | 1 | 0 | 1 |
| *Pachydactylus affinis* | 0 | 1 | 0 | 1 |
| *Pachydactylus austeni* | 0 | 1 | 0 | 1 |
| *Pachydactylus barnardi* | 0 | 1 | 0 | 1 |
| *Pachydactylus bicolor* | 0 | 1 | 0 | 1 |
| *Pachydactylus capensis* | 0 | 1 | 0 | 1 |
| *Pachydactylus caraculicus* | 0 | 1 | 0 | 1 |
| *Pachydactylus carinatus* | 0 | 1 | 0 | 1 |
| *Pachydactylus fasciatus* | 0 | 1 | 0 | 1 |
| *Pachydactylus formosus* | 0 | 1 | 0 | 1 |
| *Pachydactylus gaiasensis* | 0 | 1 | 0 | 1 |
| *Pachydactylus geitje* | 0 | 1 | 0 | 1 |
| *Pachydactylus griffini* | 0 | 1 | 0 | 1 |
| *Pachydactylus haackei* | 0 | 1 | 0 | 1 |
| *Pachydactylus kladaroderma* | 0 | 1 | 0 | 1 |
| *Pachydactylus labialis* | 0 | 1 | 0 | 1 |
| *Pachydactylus laevigatus* | 0 | 1 | 0 | 1 |
| *Pachydactylus maculatus* | 0 | 1 | 0 | 1 |
| *Pachydactylus mariquensis* | 0 | 1 | 0 | 1 |
| *Pachydactylus mclachlani* | 0 | 1 | 0 | 1 |
| *Pachydactylus monicae* | 0 | 1 | 0 | 1 |
| *Pachydactylus montanus* | 0 | 1 | 0 | 1 |
| *Pachydactylus namaquensis* | 0 | 1 | 0 | 1 |
| *Pachydactylus oculatus* | 0 | 1 | 0 | 1 |
| *Pachydactylus oreophilus* | 0 | 1 | 0 | 1 |
| *Pachydactylus oshaughnessyi* | 0 | 1 | 0 | 1 |
| *Pachydactylus parascutatus* | 0 | 1 | 0 | 1 |
| *Pachydactylus punctatus* | 0 | 1 | 0 | 1 |
| *Pachydactylus purcelli* | 0 | 1 | 0 | 1 |
| *Pachydactylus rangei* | 0 | 1 | 0 | 1 |
| *Pachydactylus reconditus* | 0 | 1 | 0 | 1 |
| *Pachydactylus robertsi* | 0 | 1 | 0 | 1 |
| *Pachydactylus rugosus* | 0 | 1 | 0 | 1 |
| *Pachydactylus sansteynae* | 0 | 1 | 0 | 1 |
| *Pachydactylus scherzi* | 0 | 1 | 0 | 1 |
| *Pachydactylus scutatus* | 0 | 1 | 0 | 1 |
| *Pachydactylus serval* | 0 | 1 | 0 | 1 |
| *Pachydactylus tigrinus* | 0 | 1 | 0 | 1 |
| *Pachydactylus tsodiloensis* | 0 | 1 | 0 | 1 |
| *Pachydactylus vansoni* | 0 | 1 | 0 | 1 |
| *Pachydactylus vanzyli* | 0 | 1 | 0 | 1 |
| *Pachydactylus waterbergensis* | 0 | 1 | 0 | 1 |
| *Pachydactylus weberi* | 0 | 1 | 0 | 1 |
| *Pamelaescincus gardineri* | 0 | 1 | 0 | 1 |
| *Panaspis breviceps* | 0 | 1 | 0 | 1 |
| *Panaspis togoensis* | 0 | 1 | 0 | 1 |
| *Pantherophis alleghaniensis* | 1 | 0 | 0 | 0 |
| *Pantherophis bairdi* | 1 | 0 | 0 | 0 |
| *Pantherophis emoryi* | 1 | 0 | 0 | 0 |
| *Pantherophis guttatus* | 1 | 0 | 0 | 0 |
| *Pantherophis obsoletus* | 1 | 0 | 0 | 0 |
| *Pantherophis slowinskii* | 1 | 0 | 0 | 0 |
| *Pantherophis spiloides* | 1 | 0 | 0 | 0 |
| *Pantherophis vulpinus* | 1 | 0 | 0 | 0 |
| *Papuascincus stanleyanus* | 0 | 1 | 0 | 1 |
| *Paracontias brocchii* | 0 | 1 | 0 | 1 |
| *Paracontias hildebrandti* | 0 | 1 | 0 | 1 |
| *Paracontias holomelas* | 0 | 1 | 0 | 1 |
| *Paracontias manify* | 0 | 1 | 0 | 1 |
| *Paracontias rothschildi* | 0 | 1 | 0 | 1 |
| *Paradelma orientalis* | 0 | 1 | 0 | 1 |
| *Paragehyra gabriellae* | 0 | 1 | 0 | 1 |
| *Parahydrophis mertoni* | 0 | 1 | 0 | 1 |
| *Pareas boulengeri* | 1 | 0 | 0 | 0 |
| *Pareas carinatus* | 0 | 1 | 0 | 1 |
| *Pareas formosensis* | 0 | 1 | 0 | 1 |
| *Pareas hamptoni* | 0 | 1 | 0 | 1 |
| *Pareas macularius* | 0 | 1 | 0 | 1 |
| *Pareas margaritophorus* | 0 | 1 | 0 | 1 |
| *Pareas monticola* | 0 | 1 | 0 | 1 |
| *Pareas nuchalis* | 0 | 1 | 0 | 1 |
| *Parias flavomaculatus* | 0 | 1 | 0 | 1 |
| *Parias hageni* | 0 | 1 | 0 | 1 |
| *Parias malcolmi* | 0 | 1 | 0 | 1 |
| *Parias schultzei* | 0 | 1 | 0 | 1 |
| *Parias sumatranus* | 0 | 1 | 0 | 1 |
| *Paroedura androyensis* | 0 | 1 | 0 | 1 |
| *Paroedura bastardi* | 0 | 1 | 0 | 1 |
| *Paroedura gracilis* | 0 | 1 | 0 | 1 |
| *Paroedura homalorhina* | 0 | 1 | 0 | 1 |
| *Paroedura karstophila* | 0 | 1 | 0 | 1 |
| *Paroedura lohatsara* | 0 | 1 | 0 | 1 |
| *Paroedura masobe* | 0 | 1 | 0 | 1 |
| *Paroedura oviceps* | 0 | 1 | 0 | 1 |
| *Paroedura picta* | 0 | 1 | 0 | 1 |
| *Paroedura sanctijohannis* | 0 | 1 | 0 | 1 |
| *Paroedura stumpffi* | 0 | 1 | 0 | 1 |
| *Paroedura tanjaka* | 0 | 1 | 0 | 1 |
| *Paroedura vazimba* | 0 | 1 | 0 | 1 |
| *Parvilacerta fraasii* | 1 | 0 | 0 | 0 |
| *Parvilacerta parva* | 1 | 0 | 0 | 0 |
| *Parvoscincus sisoni* | 0 | 1 | 0 | 1 |
| *Pedioplanis breviceps* | 0 | 1 | 0 | 1 |
| *Pedioplanis burchelli* | 0 | 1 | 0 | 1 |
| *Pedioplanis gaerdesi* | 0 | 1 | 0 | 1 |
| *Pedioplanis husabensis* | 0 | 1 | 0 | 1 |
| *Pedioplanis inornata* | 0 | 1 | 0 | 1 |
| *Pedioplanis laticeps* | 0 | 1 | 0 | 1 |
| *Pedioplanis lineoocellata* | 0 | 1 | 0 | 1 |
| *Pedioplanis namaquensis* | 0 | 1 | 0 | 1 |
| *Pedioplanis rubens* | 0 | 1 | 0 | 1 |
| *Pedioplanis undata* | 0 | 1 | 0 | 1 |
| *Pelamis platura* | 0 | 1 | 0 | 1 |
| *Perochirus ateles* | 0 | 1 | 0 | 1 |
| *Petracola ventrimaculatus* | 0 | 1 | 0 | 1 |
| *Petrosaurus mearnsi* | 1 | 0 | 0 | 0 |
| *Petrosaurus repens* | 1 | 0 | 0 | 0 |
| *Petrosaurus thalassinus* | 0 | 1 | 0 | 1 |
| *Phalotris lativittatus* | 0 | 1 | 0 | 1 |
| *Phalotris lemniscatus* | 0 | 1 | 0 | 1 |
| *Phalotris mertensi* | 0 | 1 | 0 | 1 |
| *Phalotris nasutus* | 0 | 1 | 0 | 1 |
| *Phelsuma abbotti* | 0 | 1 | 0 | 1 |
| *Phelsuma andamanense* | 0 | 1 | 0 | 1 |
| *Phelsuma antanosy* | 0 | 1 | 0 | 1 |
| *Phelsuma astriata* | 0 | 1 | 0 | 1 |
| *Phelsuma barbouri* | 0 | 1 | 0 | 1 |
| *Phelsuma berghofi* | 0 | 1 | 0 | 1 |
| *Phelsuma borbonica* | 0 | 1 | 0 | 1 |
| *Phelsuma breviceps* | 0 | 1 | 0 | 1 |
| *Phelsuma cepediana* | 0 | 1 | 0 | 1 |
| *Phelsuma comorensis* | 0 | 1 | 0 | 1 |
| *Phelsuma dubia* | 0 | 1 | 0 | 1 |
| *Phelsuma edwardnewtoni* | 0 | 1 | 0 | 1 |
| *Phelsuma flavigularis* | 0 | 1 | 0 | 1 |
| *Phelsuma gigas* | 0 | 1 | 0 | 1 |
| *Phelsuma guentheri* | 0 | 1 | 0 | 1 |
| *Phelsuma guimbeaui* | 0 | 1 | 0 | 1 |
| *Phelsuma guttata* | 0 | 1 | 0 | 1 |
| *Phelsuma hielscheri* | 0 | 1 | 0 | 1 |
| *Phelsuma inexpectata* | 0 | 1 | 0 | 1 |
| *Phelsuma kely* | 0 | 1 | 0 | 1 |
| *Phelsuma klemmeri* | 0 | 1 | 0 | 1 |
| *Phelsuma laticauda* | 0 | 1 | 0 | 1 |
| *Phelsuma lineata* | 0 | 1 | 0 | 1 |
| *Phelsuma madagascariensis* | 0 | 1 | 0 | 1 |
| *Phelsuma malamakibo* | 0 | 1 | 0 | 1 |
| *Phelsuma modesta* | 0 | 1 | 0 | 1 |
| *Phelsuma mutabilis* | 0 | 1 | 0 | 1 |
| *Phelsuma nigristriata* | 0 | 1 | 0 | 1 |
| *Phelsuma ocellata* | 0 | 1 | 0 | 1 |
| *Phelsuma ornata* | 0 | 1 | 0 | 1 |
| *Phelsuma parkeri* | 0 | 1 | 0 | 1 |
| *Phelsuma pronki* | 0 | 1 | 0 | 1 |
| *Phelsuma pusilla* | 0 | 1 | 0 | 1 |
| *Phelsuma quadriocellata* | 0 | 1 | 0 | 1 |
| *Phelsuma ravenala* | 0 | 1 | 0 | 1 |
| *Phelsuma robertmertensi* | 0 | 1 | 0 | 1 |
| *Phelsuma seippi* | 0 | 1 | 0 | 1 |
| *Phelsuma serraticauda* | 0 | 1 | 0 | 1 |
| *Phelsuma standingi* | 0 | 1 | 0 | 1 |
| *Phelsuma sundbergi* | 0 | 1 | 0 | 1 |
| *Phelsuma vanheygeni* | 0 | 1 | 0 | 1 |
| *Philochortus spinalis* | 0 | 1 | 0 | 1 |
| *Philodryas aestivus* | 0 | 1 | 0 | 1 |
| *Philodryas baroni* | 0 | 1 | 0 | 1 |
| *Philodryas mattogrossensis* | 0 | 1 | 0 | 1 |
| *Philodryas nattereri* | 0 | 1 | 0 | 1 |
| *Philodryas olfersii* | 0 | 1 | 0 | 1 |
| *Philodryas patagoniensis* | 0 | 1 | 0 | 1 |
| *Philodryas psammophidea* | 0 | 1 | 0 | 1 |
| *Philodryas viridissima* | 0 | 1 | 0 | 1 |
| *Philothamnus angolensis* | 0 | 1 | 0 | 1 |
| *Philothamnus carinatus* | 0 | 1 | 0 | 1 |
| *Philothamnus girardi* | 0 | 1 | 0 | 1 |
| *Philothamnus heterodermus* | 0 | 1 | 0 | 1 |
| *Philothamnus hoplogaster* | 0 | 1 | 0 | 1 |
| *Philothamnus natalensis* | 0 | 1 | 0 | 1 |
| *Philothamnus nitidus* | 0 | 1 | 0 | 1 |
| *Philothamnus semivariegatus* | 0 | 1 | 0 | 1 |
| *Philothamnus thomensis* | 0 | 1 | 0 | 1 |
| *Phimophis guerini* | 0 | 1 | 0 | 1 |
| *Phimophis iglesiasi* | 0 | 1 | 0 | 1 |
| *Phoboscincus garnieri* | 0 | 1 | 0 | 1 |
| *Phoenicolacerta cyanisparsa* | 1 | 0 | 0 | 0 |
| *Phoenicolacerta kulzeri* | 1 | 0 | 0 | 0 |
| *Phoenicolacerta laevis* | 1 | 0 | 0 | 0 |
| *Pholidobolus macbrydei* | 0 | 1 | 0 | 1 |
| *Pholidobolus montium* | 0 | 1 | 0 | 1 |
| *Phoxophrys nigrilabris* | 0 | 1 | 0 | 1 |
| *Phrynocephalus albolineatus* | 1 | 0 | 0 | 0 |
| *Phrynocephalus axillaris* | 1 | 0 | 0 | 0 |
| *Phrynocephalus forsythii* | 1 | 0 | 0 | 0 |
| *Phrynocephalus guttatus* | 1 | 0 | 0 | 0 |
| *Phrynocephalus helioscopus* | 1 | 0 | 0 | 0 |
| *Phrynocephalus interscapularis* | 1 | 0 | 0 | 0 |
| *Phrynocephalus lidskii* | 1 | 0 | 0 | 0 |
| *Phrynocephalus melanurus* | 1 | 0 | 0 | 0 |
| *Phrynocephalus mystaceus* | 1 | 0 | 0 | 0 |
| *Phrynocephalus przewalskii* | 1 | 0 | 0 | 0 |
| *Phrynocephalus putjatai* | 1 | 0 | 0 | 0 |
| *Phrynocephalus raddei* | 1 | 0 | 0 | 0 |
| *Phrynocephalus scutellatus* | 1 | 0 | 0 | 0 |
| *Phrynocephalus theobaldi* | 1 | 0 | 0 | 0 |
| *Phrynocephalus versicolor* | 1 | 0 | 0 | 0 |
| *Phrynocephalus vlangalii* | 1 | 0 | 0 | 0 |
| *Phrynosoma asio* | 0 | 1 | 0 | 1 |
| *Phrynosoma blainvillii* | 1 | 0 | 0 | 0 |
| *Phrynosoma braconnieri* | 0 | 1 | 0 | 1 |
| *Phrynosoma cerroense* | 1 | 0 | 0 | 0 |
| *Phrynosoma cornutum* | 1 | 0 | 0 | 0 |
| *Phrynosoma coronatum* | 1 | 0 | 0 | 0 |
| *Phrynosoma ditmarsi* | 1 | 0 | 0 | 0 |
| *Phrynosoma douglassii* | 1 | 0 | 0 | 0 |
| *Phrynosoma hernandesi* | 1 | 0 | 0 | 0 |
| *Phrynosoma mcallii* | 1 | 0 | 0 | 0 |
| *Phrynosoma modestum* | 1 | 0 | 0 | 0 |
| *Phrynosoma orbiculare* | 1 | 0 | 0 | 0 |
| *Phrynosoma platyrhinos* | 1 | 0 | 0 | 0 |
| *Phrynosoma solare* | 1 | 0 | 0 | 0 |
| *Phrynosoma taurus* | 0 | 1 | 0 | 1 |
| *Phrynosoma wigginsi* | 1 | 0 | 0 | 0 |
| *Phyllodactylus bordai* | 0 | 1 | 0 | 1 |
| *Phyllodactylus bugastrolepis* | 1 | 0 | 0 | 0 |
| *Phyllodactylus davisi* | 0 | 1 | 0 | 1 |
| *Phyllodactylus delcampoi* | 0 | 1 | 0 | 1 |
| *Phyllodactylus duellmani* | 0 | 1 | 0 | 1 |
| *Phyllodactylus homolepidurus* | 0 | 1 | 0 | 1 |
| *Phyllodactylus lanei* | 0 | 1 | 0 | 1 |
| *Phyllodactylus nocticolus* | 1 | 0 | 0 | 0 |
| *Phyllodactylus paucituberculatus* | 0 | 1 | 0 | 1 |
| *Phyllodactylus reissii* | 0 | 1 | 0 | 1 |
| *Phyllodactylus tuberculosus* | 0 | 1 | 0 | 1 |
| *Phyllodactylus unctus* | 0 | 1 | 0 | 1 |
| *Phyllodactylus wirshingi* | 0 | 1 | 0 | 1 |
| *Phyllodactylus xanti* | 0 | 1 | 0 | 1 |
| *Phyllopezus maranjonensis* | 0 | 1 | 0 | 1 |
| *Phyllopezus periosus* | 0 | 1 | 0 | 1 |
| *Phyllopezus pollicaris* | 0 | 1 | 0 | 1 |
| *Phyllorhynchus decurtatus* | 1 | 0 | 0 | 0 |
| *Phyllurus amnicola* | 0 | 1 | 0 | 1 |
| *Phyllurus kabikabi* | 0 | 1 | 0 | 1 |
| *Phyllurus platurus* | 0 | 1 | 0 | 1 |
| *Phymaturus antofagastensis* | 0 | 0 | 1 | 0 |
| *Phymaturus dorsimaculatus* | 0 | 0 | 1 | 0 |
| *Phymaturus indistinctus* | 0 | 0 | 1 | 0 |
| *Phymaturus mallimaccii* | 0 | 0 | 1 | 0 |
| *Phymaturus palluma* | 0 | 0 | 1 | 0 |
| *Phymaturus patagonicus* | 0 | 0 | 1 | 0 |
| *Phymaturus punae* | 0 | 0 | 1 | 0 |
| *Phymaturus somuncurensis* | 0 | 0 | 1 | 0 |
| *Physignathus cocincinus* | 0 | 1 | 0 | 1 |
| *Physignathus lesueurii* | 0 | 1 | 0 | 1 |
| *Pituophis catenifer* | 1 | 0 | 0 | 0 |
| *Pituophis deppei* | 0 | 1 | 0 | 1 |
| *Pituophis lineaticollis* | 0 | 1 | 0 | 1 |
| *Pituophis melanoleucus* | 1 | 0 | 0 | 0 |
| *Pituophis ruthveni* | 1 | 0 | 0 | 0 |
| *Pituophis vertebralis* | 1 | 0 | 0 | 0 |
| *Placosoma cordylinum* | 0 | 1 | 0 | 1 |
| *Placosoma glabellum* | 0 | 1 | 0 | 1 |
| *Plagiopholis styani* | 1 | 0 | 0 | 0 |
| *Platyceps collaris* | 1 | 0 | 0 | 0 |
| *Platyceps florulentus* | 1 | 1 | 0 | 0 |
| *Platyceps karelini* | 1 | 0 | 0 | 0 |
| *Platyceps najadum* | 1 | 0 | 0 | 0 |
| *Platyceps rhodorachis* | 1 | 1 | 0 | 0 |
| *Platyceps rogersi* | 1 | 0 | 0 | 0 |
| *Platyceps ventromaculatus* | 1 | 0 | 0 | 0 |
| *Platysaurus broadleyi* | 0 | 1 | 0 | 1 |
| *Platysaurus capensis* | 0 | 1 | 0 | 1 |
| *Platysaurus intermedius* | 0 | 1 | 0 | 1 |
| *Platysaurus minor* | 0 | 1 | 0 | 1 |
| *Platysaurus mitchelli* | 0 | 1 | 0 | 1 |
| *Platysaurus monotropis* | 0 | 1 | 0 | 1 |
| *Platysaurus pungweensis* | 0 | 1 | 0 | 1 |
| *Plestiodon anthracinus* | 1 | 0 | 0 | 0 |
| *Plestiodon barbouri* | 0 | 1 | 0 | 1 |
| *Plestiodon brevirostris* | 0 | 1 | 0 | 1 |
| *Plestiodon callicephalus* | 1 | 0 | 0 | 0 |
| *Plestiodon capito* | 1 | 0 | 0 | 0 |
| *Plestiodon chinensis* | 0 | 1 | 0 | 1 |
| *Plestiodon copei* | 0 | 1 | 0 | 1 |
| *Plestiodon dugesii* | 0 | 1 | 0 | 1 |
| *Plestiodon egregius* | 1 | 0 | 0 | 0 |
| *Plestiodon elegans* | 0 | 1 | 0 | 1 |
| *Plestiodon fasciatus* | 1 | 0 | 0 | 0 |
| *Plestiodon gilberti* | 1 | 0 | 0 | 0 |
| *Plestiodon inexpectatus* | 1 | 0 | 0 | 0 |
| *Plestiodon japonicus* | 1 | 0 | 0 | 0 |
| *Plestiodon kishinouyei* | 0 | 1 | 0 | 1 |
| *Plestiodon lagunensis* | 1 | 0 | 0 | 0 |
| *Plestiodon laticeps* | 1 | 0 | 0 | 0 |
| *Plestiodon latiscutatus* | 1 | 0 | 0 | 0 |
| *Plestiodon longirostris* | 1 | 0 | 0 | 0 |
| *Plestiodon lynxe* | 0 | 1 | 0 | 1 |
| *Plestiodon marginatus* | 0 | 1 | 0 | 1 |
| *Plestiodon multivirgatus* | 1 | 0 | 0 | 0 |
| *Plestiodon obsoletus* | 1 | 0 | 0 | 0 |
| *Plestiodon ochoterenae* | 0 | 1 | 0 | 1 |
| *Plestiodon parviauriculatus* | 0 | 1 | 0 | 1 |
| *Plestiodon parvulus* | 0 | 1 | 0 | 1 |
| *Plestiodon quadrilineatus* | 0 | 1 | 0 | 1 |
| *Plestiodon reynoldsi* | 1 | 0 | 0 | 0 |
| *Plestiodon septentrionalis* | 1 | 0 | 0 | 0 |
| *Plestiodon skiltonianus* | 1 | 0 | 0 | 0 |
| *Plestiodon stimpsonii* | 0 | 1 | 0 | 1 |
| *Plestiodon sumichrasti* | 0 | 1 | 0 | 1 |
| *Plestiodon tamdaoensis* | 0 | 1 | 0 | 1 |
| *Plestiodon tetragrammus* | 1 | 0 | 0 | 0 |
| *Plestiodon tunganus* | 0 | 1 | 0 | 1 |
| *Pletholax gracilis* | 0 | 1 | 0 | 1 |
| *Plica lumaria* | 0 | 1 | 0 | 1 |
| *Plica plica* | 0 | 1 | 0 | 1 |
| *Plica umbra* | 0 | 1 | 0 | 1 |
| *Podarcis atrata* | 1 | 0 | 0 | 0 |
| *Podarcis bocagei* | 1 | 0 | 0 | 0 |
| *Podarcis carbonelli* | 1 | 0 | 0 | 0 |
| *Podarcis erhardii* | 1 | 0 | 0 | 0 |
| *Podarcis filfolensis* | 1 | 0 | 0 | 0 |
| *Podarcis gaigeae* | 1 | 0 | 0 | 0 |
| *Podarcis hispanicus* | 1 | 0 | 0 | 0 |
| *Podarcis lilfordi* | 1 | 0 | 0 | 0 |
| *Podarcis melisellensis* | 1 | 0 | 0 | 0 |
| *Podarcis milensis* | 1 | 0 | 0 | 0 |
| *Podarcis muralis* | 1 | 0 | 0 | 0 |
| *Podarcis peloponnesiacus* | 1 | 0 | 0 | 0 |
| *Podarcis pityusensis* | 1 | 0 | 0 | 0 |
| *Podarcis raffonei* | 1 | 0 | 0 | 0 |
| *Podarcis siculus* | 1 | 0 | 0 | 0 |
| *Podarcis tauricus* | 1 | 0 | 0 | 0 |
| *Podarcis tiliguerta* | 1 | 0 | 0 | 0 |
| *Podarcis vaucheri* | 1 | 0 | 0 | 0 |
| *Pogona barbata* | 0 | 1 | 0 | 1 |
| *Pogona henrylawsoni* | 0 | 1 | 0 | 1 |
| *Pogona minima* | 0 | 1 | 0 | 1 |
| *Pogona minor* | 0 | 1 | 0 | 1 |
| *Pogona nullarbor* | 0 | 1 | 0 | 1 |
| *Pogona vitticeps* | 0 | 1 | 0 | 1 |
| *Polemon acanthias* | 0 | 1 | 0 | 1 |
| *Polemon collaris* | 0 | 1 | 0 | 1 |
| *Polemon notatus* | 0 | 1 | 0 | 1 |
| *Polychrus acutirostris* | 0 | 1 | 0 | 1 |
| *Polychrus femoralis* | 0 | 1 | 0 | 1 |
| *Polychrus gutturosus* | 0 | 1 | 0 | 1 |
| *Polychrus marmoratus* | 0 | 1 | 0 | 1 |
| *Polyodontognathus caerulescens* | 0 | 1 | 0 | 1 |
| *Popeia popeiorum* | 0 | 1 | 0 | 1 |
| *Poromera fordii* | 0 | 1 | 0 | 1 |
| *Porthidium dunni* | 0 | 1 | 0 | 1 |
| *Porthidium lansbergii* | 0 | 1 | 0 | 1 |
| *Porthidium nasutum* | 0 | 1 | 0 | 1 |
| *Porthidium ophryomegas* | 0 | 1 | 0 | 1 |
| *Porthidium porrasi* | 0 | 1 | 0 | 1 |
| *Porthidium yucatanicum* | 0 | 1 | 0 | 1 |
| *Potamites ecpleopus* | 0 | 1 | 0 | 1 |
| *Potamites juruazensis* | 0 | 1 | 0 | 1 |
| *Prasinohaema virens* | 0 | 1 | 0 | 1 |
| *Pristidactylus scapulatus* | 0 | 0 | 1 | 0 |
| *Pristidactylus torquatus* | 0 | 0 | 1 | 0 |
| *Pristurus abdelkuri* | 0 | 1 | 0 | 1 |
| *Pristurus carteri* | 0 | 1 | 0 | 1 |
| *Pristurus celerrimus* | 0 | 1 | 0 | 1 |
| *Pristurus crucifer* | 0 | 1 | 0 | 1 |
| *Pristurus flavipunctatus* | 1 | 1 | 0 | 0 |
| *Pristurus guichardi* | 0 | 1 | 0 | 1 |
| *Pristurus insignis* | 0 | 1 | 0 | 1 |
| *Pristurus minimus* | 0 | 1 | 0 | 1 |
| *Pristurus rupestris* | 1 | 1 | 0 | 0 |
| *Pristurus sokotranus* | 0 | 1 | 0 | 1 |
| *Pristurus somalicus* | 0 | 1 | 0 | 1 |
| *Proablepharus reginae* | 0 | 1 | 0 | 1 |
| *Proatheris superciliaris* | 0 | 1 | 0 | 1 |
| *Procellosaurinus erythrocercus* | 0 | 1 | 0 | 1 |
| *Procellosaurinus tetradactylus* | 0 | 1 | 0 | 1 |
| *Proctoporus bolivianus* | 0 | 1 | 0 | 1 |
| *Proctoporus guentheri* | 0 | 1 | 0 | 1 |
| *Proctoporus pachyurus* | 0 | 1 | 0 | 1 |
| *Proctoporus subsolanus* | 0 | 1 | 0 | 1 |
| *Proctoporus sucullucu* | 0 | 1 | 0 | 1 |
| *Proctoporus unsaacae* | 0 | 1 | 0 | 1 |
| *Proscelotes eggeli* | 0 | 1 | 0 | 1 |
| *Prosymna greigerti* | 0 | 1 | 0 | 1 |
| *Prosymna janii* | 0 | 1 | 0 | 1 |
| *Prosymna meleagris* | 0 | 1 | 0 | 1 |
| *Prosymna ruspolii* | 0 | 1 | 0 | 1 |
| *Prosymna visseri* | 0 | 1 | 0 | 1 |
| *Protobothrops cornutus* | 0 | 1 | 0 | 1 |
| *Protobothrops elegans* | 0 | 1 | 0 | 1 |
| *Protobothrops flavoviridis* | 0 | 1 | 0 | 1 |
| *Protobothrops jerdonii* | 1 | 0 | 0 | 0 |
| *Protobothrops kaulbacki* | 0 | 1 | 0 | 1 |
| *Protobothrops mucrosquamatus* | 1 | 1 | 0 | 0 |
| *Protobothrops tokarensis* | 0 | 1 | 0 | 1 |
| *Protobothrops xiangchengensis* | 0 | 1 | 0 | 1 |
| *Psammodromus algirus* | 1 | 0 | 0 | 0 |
| *Psammodromus blanci* | 1 | 0 | 0 | 0 |
| *Psammodromus hispanicus* | 1 | 0 | 0 | 0 |
| *Psammodynastes pictus* | 0 | 1 | 0 | 1 |
| *Psammodynastes pulverulentus* | 0 | 1 | 0 | 1 |
| *Psammophis angolensis* | 0 | 1 | 0 | 1 |
| *Psammophis biseriatus* | 0 | 1 | 0 | 1 |
| *Psammophis condanarus* | 0 | 1 | 0 | 1 |
| *Psammophis crucifer* | 0 | 1 | 0 | 1 |
| *Psammophis jallae* | 0 | 1 | 0 | 1 |
| *Psammophis leightoni* | 0 | 1 | 0 | 1 |
| *Psammophis leopardinus* | 0 | 1 | 0 | 1 |
| *Psammophis lineatus* | 0 | 1 | 0 | 1 |
| *Psammophis lineolatus* | 1 | 0 | 0 | 0 |
| *Psammophis mossambicus* | 0 | 1 | 0 | 1 |
| *Psammophis notostictus* | 0 | 1 | 0 | 1 |
| *Psammophis orientalis* | 0 | 1 | 0 | 1 |
| *Psammophis phillipsi* | 0 | 1 | 0 | 1 |
| *Psammophis praeornatus* | 0 | 1 | 0 | 1 |
| *Psammophis punctulatus* | 0 | 1 | 0 | 1 |
| *Psammophis rukwae* | 0 | 1 | 0 | 1 |
| *Psammophis schokari* | 1 | 0 | 0 | 0 |
| *Psammophis sibilans* | 0 | 1 | 0 | 1 |
| *Psammophis subtaeniatus* | 0 | 1 | 0 | 1 |
| *Psammophis sudanensis* | 0 | 1 | 0 | 1 |
| *Psammophis tanganicus* | 0 | 1 | 0 | 1 |
| *Psammophis trigrammus* | 0 | 1 | 0 | 1 |
| *Psammophylax acutus* | 0 | 1 | 0 | 1 |
| *Psammophylax rhombeatus* | 0 | 1 | 0 | 1 |
| *Psammophylax tritaeniatus* | 0 | 1 | 0 | 1 |
| *Psammophylax variabilis* | 0 | 1 | 0 | 1 |
| *Pseudablabes agassizii* | 0 | 1 | 0 | 1 |
| *Pseudaspis cana* | 0 | 1 | 0 | 1 |
| *Pseudechis australis* | 0 | 1 | 0 | 1 |
| *Pseudechis butleri* | 0 | 1 | 0 | 1 |
| *Pseudechis colletti* | 0 | 1 | 0 | 1 |
| *Pseudechis guttatus* | 0 | 1 | 0 | 1 |
| *Pseudechis papuanus* | 0 | 1 | 0 | 1 |
| *Pseudechis porphyriacus* | 0 | 1 | 0 | 1 |
| *Pseudelaphe flavirufa* | 0 | 1 | 0 | 1 |
| *Pseudemoia entrecasteauxii* | 0 | 1 | 0 | 1 |
| *Pseudemoia pagenstecheri* | 0 | 1 | 0 | 1 |
| *Pseuderemias smithii* | 0 | 1 | 0 | 1 |
| *Pseudoacontias menamainty* | 0 | 1 | 0 | 1 |
| *Pseudoboa coronata* | 0 | 1 | 0 | 1 |
| *Pseudoboa neuwiedii* | 0 | 1 | 0 | 1 |
| *Pseudoboa nigra* | 0 | 1 | 0 | 1 |
| *Pseudoboodon lemniscatus* | 0 | 1 | 0 | 1 |
| *Pseudocalotes brevipes* | 0 | 1 | 0 | 1 |
| *Pseudocalotes flavigula* | 0 | 1 | 0 | 1 |
| *Pseudocerastes fieldi* | 1 | 0 | 0 | 0 |
| *Pseudocerastes persicus* | 1 | 0 | 0 | 0 |
| *Pseudocyclophis persicus* | 1 | 0 | 0 | 0 |
| *Pseudoeryx plicatilis* | 0 | 1 | 0 | 1 |
| *Pseudoficimia frontalis* | 0 | 1 | 0 | 1 |
| *Pseudogekko compressicorpus* | 0 | 1 | 0 | 1 |
| *Pseudogekko smaragdinus* | 0 | 1 | 0 | 1 |
| *Pseudogonatodes guianensis* | 0 | 1 | 0 | 1 |
| *Pseudogonatodes lunulatus* | 0 | 1 | 0 | 1 |
| *Pseudogonatodes manessi* | 0 | 1 | 0 | 1 |
| *Pseudoleptodeira latifasciata* | 0 | 1 | 0 | 1 |
| *Pseudoleptodeira uribei* | 0 | 1 | 0 | 1 |
| *Pseudonaja modesta* | 0 | 1 | 0 | 1 |
| *Pseudonaja textilis* | 0 | 1 | 0 | 1 |
| *Pseudopus apodus* | 1 | 0 | 0 | 0 |
| *Pseudorabdion oxycephalum* | 0 | 1 | 0 | 1 |
| *Pseudothecadactylus lindneri* | 0 | 1 | 0 | 1 |
| *Pseudotomodon trigonatus* | 0 | 0 | 1 | 0 |
| *Pseudotrapelus sinaitus* | 1 | 0 | 0 | 0 |
| *Pseudotyphlops philippinus* | 0 | 1 | 0 | 1 |
| *Pseudoxenodon bambusicola* | 0 | 1 | 0 | 1 |
| *Pseudoxenodon karlschmidti* | 0 | 1 | 0 | 1 |
| *Pseudoxenodon macrops* | 0 | 1 | 0 | 1 |
| *Pseudoxyrhopus ambreensis* | 0 | 1 | 0 | 1 |
| *Pseustes sulphureus* | 0 | 1 | 0 | 1 |
| *Psilophthalmus paeminosus* | 0 | 1 | 0 | 1 |
| *Psomophis genimaculatus* | 0 | 1 | 0 | 1 |
| *Psomophis joberti* | 0 | 1 | 0 | 1 |
| *Psomophis obtusus* | 0 | 1 | 0 | 1 |
| *Ptenopus carpi* | 0 | 1 | 0 | 1 |
| *Ptyas korros* | 0 | 1 | 0 | 1 |
| *Ptyas mucosa* | 1 | 0 | 0 | 0 |
| *Ptychoglossus brevifrontalis* | 0 | 1 | 0 | 1 |
| *Ptychophis flavovirgatus* | 0 | 1 | 0 | 1 |
| *Ptychozoon kuhli* | 0 | 1 | 0 | 1 |
| *Ptychozoon lionotum* | 0 | 1 | 0 | 1 |
| *Ptychozoon rhacophorus* | 0 | 1 | 0 | 1 |
| *Ptyctolaemus collicristatus* | 0 | 1 | 0 | 1 |
| *Ptyctolaemus gularis* | 0 | 1 | 0 | 1 |
| *Ptyodactylus guttatus* | 1 | 0 | 0 | 0 |
| *Ptyodactylus hasselquistii* | 1 | 0 | 0 | 0 |
| *Ptyodactylus oudrii* | 1 | 0 | 0 | 0 |
| *Ptyodactylus ragazzii* | 1 | 0 | 0 | 0 |
| *Pygomeles braconnieri* | 0 | 1 | 0 | 1 |
| *Pygopus lepidopodus* | 0 | 1 | 0 | 1 |
| *Pygopus nigriceps* | 0 | 1 | 0 | 1 |
| *Python brongersmai* | 0 | 1 | 0 | 1 |
| *Python curtus* | 0 | 1 | 0 | 1 |
| *Python molurus* | 0 | 1 | 0 | 1 |
| *Python regius* | 0 | 1 | 0 | 1 |
| *Python sebae* | 0 | 1 | 0 | 1 |
| *Pythonodipsas carinata* | 0 | 1 | 0 | 1 |
| *Quedenfeldtia moerens* | 1 | 0 | 0 | 0 |
| *Quedenfeldtia trachyblepharus* | 1 | 0 | 0 | 0 |
| *Rachidelus brazili* | 0 | 1 | 0 | 1 |
| *Ramphotyphlops acuticauda* | 0 | 1 | 0 | 1 |
| *Ramphotyphlops albiceps* | 0 | 1 | 0 | 1 |
| *Ramphotyphlops bicolor* | 0 | 1 | 0 | 1 |
| *Ramphotyphlops braminus* | 0 | 1 | 0 | 1 |
| *Ramphotyphlops lineatus* | 0 | 1 | 0 | 1 |
| *Ramphotyphlops polygrammicus* | 0 | 1 | 0 | 1 |
| *Rankinia diemensis* | 0 | 1 | 0 | 1 |
| *Regina alleni* | 1 | 0 | 0 | 0 |
| *Regina grahami* | 1 | 0 | 0 | 0 |
| *Regina rigida* | 1 | 0 | 0 | 0 |
| *Regina septemvittata* | 1 | 0 | 0 | 0 |
| *Rhabdophis nuchalis* | 0 | 1 | 0 | 1 |
| *Rhabdophis subminiatus* | 0 | 1 | 0 | 1 |
| *Rhabdophis tigrinus* | 1 | 0 | 0 | 0 |
| *Rhachisaurus brachylepis* | 0 | 1 | 0 | 1 |
| *Rhacodactylus auriculatus* | 0 | 1 | 0 | 1 |
| *Rhacodactylus chahoua* | 0 | 1 | 0 | 1 |
| *Rhacodactylus ciliatus* | 0 | 1 | 0 | 1 |
| *Rhacodactylus leachianus* | 0 | 1 | 0 | 1 |
| *Rhacodactylus sarasinorum* | 0 | 1 | 0 | 1 |
| *Rhacodactylus trachyrhynchus* | 0 | 1 | 0 | 1 |
| *Rhadinaea flavilata* | 1 | 0 | 0 | 0 |
| *Rhadinaea fulvivittis* | 0 | 1 | 0 | 1 |
| *Rhadinophis frenatum* | 0 | 1 | 0 | 1 |
| *Rhadinophis prasina* | 0 | 1 | 0 | 1 |
| *Rhamphiophis oxyrhynchus* | 0 | 1 | 0 | 1 |
| *Rhamphiophis rubropunctatus* | 0 | 1 | 0 | 1 |
| *Rhampholeon acuminatus* | 0 | 1 | 0 | 1 |
| *Rhampholeon beraduccii* | 0 | 1 | 0 | 1 |
| *Rhampholeon boulengeri* | 0 | 1 | 0 | 1 |
| *Rhampholeon chapmanorum* | 0 | 1 | 0 | 1 |
| *Rhampholeon marshalli* | 0 | 1 | 0 | 1 |
| *Rhampholeon moyeri* | 0 | 1 | 0 | 1 |
| *Rhampholeon nchisiensis* | 0 | 1 | 0 | 1 |
| *Rhampholeon platyceps* | 0 | 1 | 0 | 1 |
| *Rhampholeon spectrum* | 0 | 1 | 0 | 1 |
| *Rhampholeon spinosus* | 0 | 1 | 0 | 1 |
| *Rhampholeon temporalis* | 0 | 1 | 0 | 1 |
| *Rhampholeon uluguruensis* | 0 | 1 | 0 | 1 |
| *Rhampholeon viridis* | 0 | 1 | 0 | 1 |
| *Rhinechis scalaris* | 1 | 0 | 0 | 0 |
| *Rhineura floridana* | 1 | 0 | 0 | 0 |
| *Rhinobothryum lentiginosum* | 0 | 1 | 0 | 1 |
| *Rhinocheilus lecontei* | 1 | 0 | 0 | 0 |
| *Rhinophis blythii* | 0 | 1 | 0 | 1 |
| *Rhinophis dorsimaculatus* | 0 | 1 | 0 | 1 |
| *Rhinophis drummondhayi* | 0 | 1 | 0 | 1 |
| *Rhinophis homolepis* | 0 | 1 | 0 | 1 |
| *Rhinophis oxyrhynchus* | 0 | 1 | 0 | 1 |
| *Rhinophis philippinus* | 0 | 1 | 0 | 1 |
| *Rhinophis travancoricus* | 0 | 1 | 0 | 1 |
| *Rhinoplocephalus bicolor* | 0 | 1 | 0 | 1 |
| *Rhinoplocephalus nigrescens* | 0 | 1 | 0 | 1 |
| *Rhinotyphlops feae* | 0 | 1 | 0 | 1 |
| *Rhinotyphlops lalandei* | 0 | 1 | 0 | 1 |
| *Rhinotyphlops newtoni* | 0 | 1 | 0 | 1 |
| *Rhinotyphlops schlegelii* | 0 | 1 | 0 | 1 |
| *Rhoptropus afer* | 0 | 1 | 0 | 1 |
| *Rhoptropus barnardi* | 0 | 1 | 0 | 1 |
| *Rhoptropus biporosus* | 0 | 1 | 0 | 1 |
| *Rhoptropus boultoni* | 0 | 1 | 0 | 1 |
| *Rhoptropus bradfieldi* | 0 | 1 | 0 | 1 |
| *Rhynchoedura ornata* | 0 | 1 | 0 | 1 |
| *Rhynchophis boulengeri* | 0 | 1 | 0 | 1 |
| *Riama colomaromani* | 0 | 1 | 0 | 1 |
| *Riama simoterus* | 0 | 1 | 0 | 1 |
| *Riama unicolor* | 0 | 1 | 0 | 1 |
| *Rieppeleon brachyurus* | 0 | 1 | 0 | 1 |
| *Rieppeleon brevicaudatus* | 0 | 1 | 0 | 1 |
| *Rieppeleon kerstenii* | 0 | 1 | 0 | 1 |
| *Ristella rurkii* | 0 | 1 | 0 | 1 |
| *Saiphos equalis* | 0 | 1 | 0 | 1 |
| *Salea horsfieldii* | 0 | 1 | 0 | 1 |
| *Salea kakhienensis* | 0 | 1 | 0 | 1 |
| *Saltuarius cornutus* | 0 | 1 | 0 | 1 |
| *Saltuarius kateae* | 0 | 1 | 0 | 1 |
| *Saltuarius moritzi* | 0 | 1 | 0 | 1 |
| *Saltuarius salebrosus* | 0 | 1 | 0 | 1 |
| *Saltuarius swaini* | 0 | 1 | 0 | 1 |
| *Saltuarius wyberba* | 0 | 1 | 0 | 1 |
| *Salvadora mexicana* | 0 | 1 | 0 | 1 |
| *Sanzinia madagascariensis* | 0 | 1 | 0 | 1 |
| *Saproscincus basiliscus* | 0 | 1 | 0 | 1 |
| *Saproscincus challengeri* | 0 | 1 | 0 | 1 |
| *Saproscincus czechurai* | 0 | 1 | 0 | 1 |
| *Saproscincus hannahae* | 0 | 1 | 0 | 1 |
| *Saproscincus lewisi* | 0 | 1 | 0 | 1 |
| *Saproscincus mustelinus* | 0 | 1 | 0 | 1 |
| *Saproscincus oriarius* | 0 | 1 | 0 | 1 |
| *Saproscincus rosei* | 0 | 1 | 0 | 1 |
| *Saproscincus spectabilis* | 0 | 1 | 0 | 1 |
| *Saproscincus tetradactylus* | 0 | 1 | 0 | 1 |
| *Saurodactylus fasciatus* | 1 | 0 | 0 | 0 |
| *Saurodactylus mauritanicus* | 1 | 0 | 0 | 0 |
| *Sauromalus ater* | 1 | 0 | 0 | 0 |
| *Sauromalus hispidus* | 1 | 0 | 0 | 0 |
| *Sauromalus klauberi* | 1 | 0 | 0 | 0 |
| *Sauromalus varius* | 1 | 0 | 0 | 0 |
| *Scaphiodontophis annulatus* | 0 | 1 | 0 | 1 |
| *Scaphiophis albopunctatus* | 0 | 1 | 0 | 1 |
| *Scelarcis perspicillata* | 1 | 0 | 0 | 0 |
| *Sceloporus adleri* | 0 | 1 | 0 | 1 |
| *Sceloporus aeneus* | 0 | 1 | 0 | 1 |
| *Sceloporus angustus* | 1 | 0 | 0 | 0 |
| *Sceloporus arenicolus* | 1 | 0 | 0 | 0 |
| *Sceloporus bicanthalis* | 0 | 1 | 0 | 1 |
| *Sceloporus bulleri* | 0 | 1 | 0 | 1 |
| *Sceloporus carinatus* | 0 | 1 | 0 | 1 |
| *Sceloporus cautus* | 1 | 0 | 0 | 0 |
| *Sceloporus chaneyi* | 0 | 1 | 0 | 1 |
| *Sceloporus chrysostictus* | 0 | 1 | 0 | 1 |
| *Sceloporus clarkii* | 1 | 0 | 0 | 0 |
| *Sceloporus consobrinus* | 1 | 0 | 0 | 0 |
| *Sceloporus couchii* | 1 | 0 | 0 | 0 |
| *Sceloporus cozumelae* | 0 | 1 | 0 | 1 |
| *Sceloporus cryptus* | 0 | 1 | 0 | 1 |
| *Sceloporus cyanogenys* | 1 | 0 | 0 | 0 |
| *Sceloporus dugesii* | 0 | 1 | 0 | 1 |
| *Sceloporus edwardtaylori* | 0 | 1 | 0 | 1 |
| *Sceloporus formosus* | 0 | 1 | 0 | 1 |
| *Sceloporus gadoviae* | 0 | 1 | 0 | 1 |
| *Sceloporus goldmani* | 0 | 1 | 0 | 1 |
| *Sceloporus graciosus* | 1 | 0 | 0 | 0 |
| *Sceloporus grammicus* | 1 | 0 | 0 | 0 |
| *Sceloporus grandaevus* | 0 | 1 | 0 | 1 |
| *Sceloporus heterolepis* | 0 | 1 | 0 | 1 |
| *Sceloporus horridus* | 0 | 1 | 0 | 1 |
| *Sceloporus hunsakeri* | 0 | 1 | 0 | 1 |
| *Sceloporus insignis* | 0 | 1 | 0 | 1 |
| *Sceloporus jalapae* | 0 | 1 | 0 | 1 |
| *Sceloporus jarrovii* | 1 | 0 | 0 | 0 |
| *Sceloporus licki* | 0 | 1 | 0 | 1 |
| *Sceloporus lineatulus* | 1 | 0 | 0 | 0 |
| *Sceloporus lundelli* | 0 | 1 | 0 | 1 |
| *Sceloporus macdougalli* | 0 | 1 | 0 | 1 |
| *Sceloporus maculosus* | 1 | 0 | 0 | 0 |
| *Sceloporus magister* | 1 | 0 | 0 | 0 |
| *Sceloporus malachiticus* | 0 | 1 | 0 | 1 |
| *Sceloporus megalepidurus* | 0 | 1 | 0 | 1 |
| *Sceloporus melanorhinus* | 0 | 1 | 0 | 1 |
| *Sceloporus merriami* | 1 | 0 | 0 | 0 |
| *Sceloporus minor* | 0 | 1 | 0 | 1 |
| *Sceloporus mucronatus* | 0 | 1 | 0 | 1 |
| *Sceloporus nelsoni* | 0 | 1 | 0 | 1 |
| *Sceloporus occidentalis* | 1 | 0 | 0 | 0 |
| *Sceloporus ochoterenae* | 0 | 1 | 0 | 1 |
| *Sceloporus olivaceus* | 1 | 0 | 0 | 0 |
| *Sceloporus orcutti* | 1 | 0 | 0 | 0 |
| *Sceloporus ornatus* | 1 | 0 | 0 | 0 |
| *Sceloporus palaciosi* | 0 | 1 | 0 | 1 |
| *Sceloporus parvus* | 0 | 1 | 0 | 1 |
| *Sceloporus poinsettii* | 1 | 0 | 0 | 0 |
| *Sceloporus pyrocephalus* | 0 | 1 | 0 | 1 |
| *Sceloporus samcolemani* | 0 | 1 | 0 | 1 |
| *Sceloporus scalaris* | 0 | 1 | 0 | 1 |
| *Sceloporus serrifer* | 1 | 1 | 0 | 0 |
| *Sceloporus siniferus* | 0 | 1 | 0 | 1 |
| *Sceloporus slevini* | 1 | 0 | 0 | 0 |
| *Sceloporus smaragdinus* | 0 | 1 | 0 | 1 |
| *Sceloporus smithi* | 0 | 1 | 0 | 1 |
| *Sceloporus spinosus* | 0 | 1 | 0 | 1 |
| *Sceloporus squamosus* | 0 | 1 | 0 | 1 |
| *Sceloporus stejnegeri* | 0 | 1 | 0 | 1 |
| *Sceloporus subpictus* | 0 | 1 | 0 | 1 |
| *Sceloporus taeniocnemis* | 0 | 1 | 0 | 1 |
| *Sceloporus teapensis* | 0 | 1 | 0 | 1 |
| *Sceloporus torquatus* | 0 | 1 | 0 | 1 |
| *Sceloporus undulatus* | 1 | 0 | 0 | 0 |
| *Sceloporus utiformis* | 0 | 1 | 0 | 1 |
| *Sceloporus vandenburgianus* | 1 | 0 | 0 | 0 |
| *Sceloporus variabilis* | 0 | 1 | 0 | 1 |
| *Sceloporus virgatus* | 1 | 0 | 0 | 0 |
| *Sceloporus woodi* | 1 | 0 | 0 | 0 |
| *Sceloporus zosteromus* | 1 | 0 | 0 | 0 |
| *Scelotes anguineus* | 0 | 1 | 0 | 1 |
| *Scelotes arenicolus* | 0 | 1 | 0 | 1 |
| *Scelotes bipes* | 0 | 1 | 0 | 1 |
| *Scelotes caffer* | 0 | 1 | 0 | 1 |
| *Scelotes gronovii* | 0 | 1 | 0 | 1 |
| *Scelotes kasneri* | 0 | 1 | 0 | 1 |
| *Scelotes mirus* | 0 | 1 | 0 | 1 |
| *Scelotes montispectus* | 0 | 1 | 0 | 1 |
| *Scelotes sexlineatus* | 0 | 1 | 0 | 1 |
| *Scincella gemmingeri* | 0 | 1 | 0 | 1 |
| *Scincella lateralis* | 1 | 0 | 0 | 0 |
| *Scincella reevesii* | 0 | 1 | 0 | 1 |
| *Scincopus fasciatus* | 1 | 0 | 0 | 0 |
| *Scincus mitranus* | 1 | 0 | 0 | 0 |
| *Scincus scincus* | 1 | 1 | 0 | 0 |
| *Seminatrix pygaea* | 1 | 0 | 0 | 0 |
| *Senticolis triaspis* | 0 | 1 | 0 | 1 |
| *Sepsina angolensis* | 0 | 1 | 0 | 1 |
| *Shinisaurus crocodilurus* | 0 | 1 | 0 | 1 |
| *Sibon nebulatus* | 0 | 1 | 0 | 1 |
| *Sibynomorphus mikanii* | 0 | 1 | 0 | 1 |
| *Sibynomorphus neuwiedi* | 0 | 1 | 0 | 1 |
| *Sibynomorphus turgidus* | 0 | 1 | 0 | 1 |
| *Sibynomorphus ventrimaculatus* | 0 | 1 | 0 | 1 |
| *Sibynophis bistrigatus* | 0 | 1 | 0 | 1 |
| *Sibynophis chinensis* | 0 | 1 | 0 | 1 |
| *Sibynophis collaris* | 0 | 1 | 0 | 1 |
| *Sibynophis subpunctatus* | 0 | 1 | 0 | 1 |
| *Sibynophis triangularis* | 0 | 1 | 0 | 1 |
| *Sigaloseps deplanchei* | 0 | 1 | 0 | 1 |
| *Sigaloseps ruficauda* | 0 | 1 | 0 | 1 |
| *Simiscincus aurantiacus* | 0 | 1 | 0 | 1 |
| *Simoselaps anomalus* | 0 | 1 | 0 | 1 |
| *Simoselaps bertholdi* | 0 | 1 | 0 | 1 |
| *Simoselaps calonotus* | 0 | 1 | 0 | 1 |
| *Simoselaps semifasciatus* | 0 | 1 | 0 | 1 |
| *Sinomicrurus japonicus* | 0 | 1 | 0 | 1 |
| *Sinomicrurus kelloggi* | 0 | 1 | 0 | 1 |
| *Sinomicrurus macclellandi* | 0 | 1 | 0 | 1 |
| *Sinonatrix aequifasciata* | 0 | 1 | 0 | 1 |
| *Sinonatrix annularis* | 0 | 1 | 0 | 1 |
| *Sinonatrix percarinata* | 0 | 1 | 0 | 1 |
| *Siphlophis cervinus* | 0 | 1 | 0 | 1 |
| *Siphlophis compressus* | 0 | 1 | 0 | 1 |
| *Siphlophis longicaudatus* | 0 | 1 | 0 | 1 |
| *Siphlophis pulcher* | 0 | 1 | 0 | 1 |
| *Sistrurus catenatus* | 1 | 0 | 0 | 0 |
| *Sistrurus miliarius* | 1 | 0 | 0 | 0 |
| *Sitana ponticeriana* | 0 | 1 | 0 | 1 |
| *Sonora semiannulata* | 1 | 0 | 0 | 0 |
| *Sordellina punctata* | 0 | 1 | 0 | 1 |
| *Spalerosophis diadema* | 1 | 0 | 0 | 0 |
| *Spalerosophis microlepis* | 1 | 0 | 0 | 0 |
| *Sphaerodactylus altavelensis* | 0 | 1 | 0 | 1 |
| *Sphaerodactylus argus* | 0 | 1 | 0 | 1 |
| *Sphaerodactylus armstrongi* | 0 | 1 | 0 | 1 |
| *Sphaerodactylus cinereus* | 0 | 1 | 0 | 1 |
| *Sphaerodactylus copei* | 0 | 1 | 0 | 1 |
| *Sphaerodactylus cricoderus* | 0 | 1 | 0 | 1 |
| *Sphaerodactylus cryphius* | 0 | 1 | 0 | 1 |
| *Sphaerodactylus darlingtoni* | 0 | 1 | 0 | 1 |
| *Sphaerodactylus elegans* | 0 | 1 | 0 | 1 |
| *Sphaerodactylus elegantulus* | 0 | 1 | 0 | 1 |
| *Sphaerodactylus fantasticus* | 0 | 1 | 0 | 1 |
| *Sphaerodactylus gaigeae* | 0 | 1 | 0 | 1 |
| *Sphaerodactylus glaucus* | 0 | 1 | 0 | 1 |
| *Sphaerodactylus goniorhynchus* | 0 | 1 | 0 | 1 |
| *Sphaerodactylus intermedius* | 0 | 1 | 0 | 1 |
| *Sphaerodactylus kirbyi* | 0 | 1 | 0 | 1 |
| *Sphaerodactylus klauberi* | 0 | 1 | 0 | 1 |
| *Sphaerodactylus leucaster* | 0 | 1 | 0 | 1 |
| *Sphaerodactylus macrolepis* | 0 | 1 | 0 | 1 |
| *Sphaerodactylus microlepis* | 0 | 1 | 0 | 1 |
| *Sphaerodactylus molei* | 0 | 1 | 0 | 1 |
| *Sphaerodactylus nicholsi* | 0 | 1 | 0 | 1 |
| *Sphaerodactylus nigropunctatus* | 0 | 1 | 0 | 1 |
| *Sphaerodactylus notatus* | 0 | 1 | 0 | 1 |
| *Sphaerodactylus ocoae* | 0 | 1 | 0 | 1 |
| *Sphaerodactylus oliveri* | 0 | 1 | 0 | 1 |
| *Sphaerodactylus parvus* | 0 | 1 | 0 | 1 |
| *Sphaerodactylus ramsdeni* | 0 | 1 | 0 | 1 |
| *Sphaerodactylus richardi* | 0 | 1 | 0 | 1 |
| *Sphaerodactylus roosevelti* | 0 | 1 | 0 | 1 |
| *Sphaerodactylus sabanus* | 0 | 1 | 0 | 1 |
| *Sphaerodactylus schwartzi* | 0 | 1 | 0 | 1 |
| *Sphaerodactylus semasiops* | 0 | 1 | 0 | 1 |
| *Sphaerodactylus shrevei* | 0 | 1 | 0 | 1 |
| *Sphaerodactylus sputator* | 0 | 1 | 0 | 1 |
| *Sphaerodactylus thompsoni* | 0 | 1 | 0 | 1 |
| *Sphaerodactylus torrei* | 0 | 1 | 0 | 1 |
| *Sphaerodactylus townsendi* | 0 | 1 | 0 | 1 |
| *Sphaerodactylus vincenti* | 0 | 1 | 0 | 1 |
| *Sphenodon punctatus* | 0 | 0 | 1 | 0 |
| *Sphenomorphus abdictus* | 0 | 1 | 0 | 1 |
| *Sphenomorphus acutus* | 0 | 1 | 0 | 1 |
| *Sphenomorphus aesculeticola* | 0 | 1 | 0 | 1 |
| *Sphenomorphus arborens* | 0 | 1 | 0 | 1 |
| *Sphenomorphus assatus* | 0 | 1 | 0 | 1 |
| *Sphenomorphus atrigularis* | 0 | 1 | 0 | 1 |
| *Sphenomorphus beyeri* | 0 | 1 | 0 | 1 |
| *Sphenomorphus buenloicus* | 0 | 1 | 0 | 1 |
| *Sphenomorphus cherriei* | 0 | 1 | 0 | 1 |
| *Sphenomorphus concinnatus* | 0 | 1 | 0 | 1 |
| *Sphenomorphus coxi* | 0 | 1 | 0 | 1 |
| *Sphenomorphus cranei* | 0 | 1 | 0 | 1 |
| *Sphenomorphus cumingi* | 0 | 1 | 0 | 1 |
| *Sphenomorphus cyanolaemus* | 0 | 1 | 0 | 1 |
| *Sphenomorphus decipiens* | 0 | 1 | 0 | 1 |
| *Sphenomorphus diwata* | 0 | 1 | 0 | 1 |
| *Sphenomorphus fasciatus* | 0 | 1 | 0 | 1 |
| *Sphenomorphus hallieri* | 0 | 1 | 0 | 1 |
| *Sphenomorphus indicus* | 0 | 1 | 0 | 1 |
| *Sphenomorphus jagori* | 0 | 1 | 0 | 1 |
| *Sphenomorphus jobiensis* | 0 | 1 | 0 | 1 |
| *Sphenomorphus kitangladensis* | 0 | 1 | 0 | 1 |
| *Sphenomorphus laterimaculatus* | 0 | 1 | 0 | 1 |
| *Sphenomorphus lawtoni* | 0 | 1 | 0 | 1 |
| *Sphenomorphus leptofasciatus* | 0 | 1 | 0 | 1 |
| *Sphenomorphus leucospilos* | 0 | 1 | 0 | 1 |
| *Sphenomorphus llanosi* | 0 | 1 | 0 | 1 |
| *Sphenomorphus luzonense* | 0 | 1 | 0 | 1 |
| *Sphenomorphus maculatus* | 0 | 1 | 0 | 1 |
| *Sphenomorphus maindroni* | 0 | 1 | 0 | 1 |
| *Sphenomorphus melanopogon* | 0 | 1 | 0 | 1 |
| *Sphenomorphus mindanensis* | 0 | 1 | 0 | 1 |
| *Sphenomorphus muelleri* | 0 | 1 | 0 | 1 |
| *Sphenomorphus multisquamatus* | 0 | 1 | 0 | 1 |
| *Sphenomorphus parvus* | 0 | 1 | 0 | 1 |
| *Sphenomorphus praesignis* | 0 | 1 | 0 | 1 |
| *Sphenomorphus sabanus* | 0 | 1 | 0 | 1 |
| *Sphenomorphus scutatus* | 0 | 1 | 0 | 1 |
| *Sphenomorphus simus* | 0 | 1 | 0 | 1 |
| *Sphenomorphus solomonis* | 0 | 1 | 0 | 1 |
| *Sphenomorphus steerei* | 0 | 1 | 0 | 1 |
| *Sphenomorphus stellatus* | 0 | 1 | 0 | 1 |
| *Sphenomorphus tagapayo* | 0 | 1 | 0 | 1 |
| *Sphenomorphus variegatus* | 0 | 1 | 0 | 1 |
| *Sphenomorphus victoria* | 0 | 1 | 0 | 1 |
| *Sphenomorphus wrighti* | 0 | 1 | 0 | 1 |
| *Spilotes pullatus* | 0 | 1 | 0 | 1 |
| *Stenocercus angel* | 0 | 1 | 0 | 1 |
| *Stenocercus angulifer* | 0 | 1 | 0 | 1 |
| *Stenocercus apurimacus* | 0 | 1 | 0 | 1 |
| *Stenocercus azureus* | 0 | 1 | 0 | 1 |
| *Stenocercus boettgeri* | 0 | 1 | 0 | 1 |
| *Stenocercus caducus* | 0 | 1 | 0 | 1 |
| *Stenocercus chota* | 0 | 1 | 0 | 1 |
| *Stenocercus chrysopygus* | 0 | 1 | 0 | 1 |
| *Stenocercus crassicaudatus* | 0 | 1 | 0 | 1 |
| *Stenocercus cupreus* | 0 | 1 | 0 | 1 |
| *Stenocercus doellojuradoi* | 0 | 0 | 1 | 0 |
| *Stenocercus empetrus* | 0 | 1 | 0 | 1 |
| *Stenocercus eunetopsis* | 0 | 1 | 0 | 1 |
| *Stenocercus festae* | 0 | 1 | 0 | 1 |
| *Stenocercus formosus* | 0 | 1 | 0 | 1 |
| *Stenocercus guentheri* | 0 | 1 | 0 | 1 |
| *Stenocercus humeralis* | 0 | 1 | 0 | 1 |
| *Stenocercus imitator* | 0 | 1 | 0 | 1 |
| *Stenocercus iridescens* | 0 | 1 | 0 | 1 |
| *Stenocercus latebrosus* | 0 | 1 | 0 | 1 |
| *Stenocercus limitaris* | 0 | 1 | 0 | 1 |
| *Stenocercus marmoratus* | 0 | 1 | 0 | 1 |
| *Stenocercus melanopygus* | 0 | 1 | 0 | 1 |
| *Stenocercus ochoai* | 0 | 1 | 0 | 1 |
| *Stenocercus orientalis* | 0 | 1 | 0 | 1 |
| *Stenocercus ornatissimus* | 0 | 1 | 0 | 1 |
| *Stenocercus ornatus* | 0 | 1 | 0 | 1 |
| *Stenocercus percultus* | 0 | 1 | 0 | 1 |
| *Stenocercus puyango* | 0 | 1 | 0 | 1 |
| *Stenocercus rhodomelas* | 0 | 1 | 0 | 1 |
| *Stenocercus roseiventris* | 0 | 1 | 0 | 1 |
| *Stenocercus scapularis* | 0 | 1 | 0 | 1 |
| *Stenocercus stigmosus* | 0 | 1 | 0 | 1 |
| *Stenocercus torquatus* | 0 | 1 | 0 | 1 |
| *Stenocercus varius* | 0 | 1 | 0 | 1 |
| *Stenodactylus arabicus* | 1 | 0 | 0 | 0 |
| *Stenodactylus doriae* | 1 | 0 | 0 | 0 |
| *Stenodactylus khobarensis* | 1 | 0 | 0 | 0 |
| *Stenodactylus leptocosymbotus* | 1 | 0 | 0 | 0 |
| *Stenodactylus petrii* | 1 | 0 | 0 | 0 |
| *Stenodactylus sthenodactylus* | 1 | 0 | 0 | 0 |
| *Stenodactylus yemenensis* | 0 | 1 | 0 | 1 |
| *Stenolepis ridleyi* | 0 | 1 | 0 | 1 |
| *Stenophis betsileanus* | 0 | 1 | 0 | 1 |
| *Stenophis citrinus* | 0 | 1 | 0 | 1 |
| *Stenophis granuliceps* | 0 | 1 | 0 | 1 |
| *Stenophis inopinae* | 0 | 1 | 0 | 1 |
| *Stenophis inornatus* | 0 | 1 | 0 | 1 |
| *Stenophis pseudogranuliceps* | 0 | 1 | 0 | 1 |
| *Stenorrhina freminvillei* | 0 | 1 | 0 | 1 |
| *Stoliczkaia borneensis* | 0 | 1 | 0 | 1 |
| *Storeria dekayi* | 1 | 1 | 0 | 0 |
| *Storeria occipitomaculata* | 1 | 0 | 0 | 0 |
| *Strobilurus torquatus* | 0 | 1 | 0 | 1 |
| *Strophurus assimilis* | 0 | 1 | 0 | 1 |
| *Strophurus ciliaris* | 0 | 1 | 0 | 1 |
| *Strophurus elderi* | 0 | 1 | 0 | 1 |
| *Strophurus intermedius* | 0 | 1 | 0 | 1 |
| *Strophurus jeanae* | 0 | 1 | 0 | 1 |
| *Strophurus krisalys* | 0 | 1 | 0 | 1 |
| *Strophurus mcmillani* | 0 | 1 | 0 | 1 |
| *Strophurus rankini* | 0 | 1 | 0 | 1 |
| *Strophurus spinigerus* | 0 | 1 | 0 | 1 |
| *Strophurus strophurus* | 0 | 1 | 0 | 1 |
| *Strophurus taeniatus* | 0 | 1 | 0 | 1 |
| *Strophurus taenicauda* | 0 | 1 | 0 | 1 |
| *Strophurus wellingtonae* | 0 | 1 | 0 | 1 |
| *Strophurus williamsi* | 0 | 1 | 0 | 1 |
| *Suta fasciata* | 0 | 1 | 0 | 1 |
| *Suta monachus* | 0 | 1 | 0 | 1 |
| *Suta spectabilis* | 0 | 1 | 0 | 1 |
| *Suta suta* | 0 | 1 | 0 | 1 |
| *Sympholis lippiens* | 0 | 1 | 0 | 1 |
| *Tachymenis peruviana* | 0 | 1 | 0 | 1 |
| *Taeniophallus affinis* | 0 | 1 | 0 | 1 |
| *Taeniophallus brevirostris* | 0 | 1 | 0 | 1 |
| *Taeniophallus nicagus* | 0 | 1 | 0 | 1 |
| *Takydromus amurensis* | 1 | 0 | 0 | 0 |
| *Takydromus dorsalis* | 0 | 1 | 0 | 1 |
| *Takydromus formosanus* | 0 | 1 | 0 | 1 |
| *Takydromus hsuehshanensis* | 0 | 1 | 0 | 1 |
| *Takydromus intermedius* | 1 | 1 | 0 | 0 |
| *Takydromus kuehnei* | 0 | 1 | 0 | 1 |
| *Takydromus sauteri* | 0 | 1 | 0 | 1 |
| *Takydromus septentrionalis* | 1 | 1 | 0 | 0 |
| *Takydromus sexlineatus* | 0 | 1 | 0 | 1 |
| *Takydromus smaragdinus* | 0 | 1 | 0 | 1 |
| *Takydromus stejnegeri* | 0 | 1 | 0 | 1 |
| *Takydromus sylvaticus* | 0 | 1 | 0 | 1 |
| *Takydromus tachydromoides* | 1 | 0 | 0 | 0 |
| *Takydromus toyamai* | 0 | 1 | 0 | 1 |
| *Takydromus wolteri* | 1 | 1 | 0 | 0 |
| *Tantalophis discolor* | 0 | 1 | 0 | 1 |
| *Tantilla melanocephala* | 0 | 1 | 0 | 1 |
| *Tarentola americana* | 0 | 1 | 0 | 1 |
| *Tarentola angustimentalis* | 1 | 0 | 0 | 0 |
| *Tarentola annularis* | 1 | 0 | 0 | 0 |
| *Tarentola boehmei* | 1 | 0 | 0 | 0 |
| *Tarentola boettgeri* | 1 | 0 | 0 | 0 |
| *Tarentola caboverdianus* | 0 | 1 | 0 | 1 |
| *Tarentola darwini* | 0 | 1 | 0 | 1 |
| *Tarentola delalandii* | 1 | 0 | 0 | 0 |
| *Tarentola deserti* | 1 | 0 | 0 | 0 |
| *Tarentola ephippiata* | 1 | 0 | 0 | 0 |
| *Tarentola gigas* | 0 | 1 | 0 | 1 |
| *Tarentola gomerensis* | 1 | 0 | 0 | 0 |
| *Tarentola mauritanica* | 1 | 0 | 0 | 0 |
| *Tarentola mindiae* | 1 | 0 | 0 | 0 |
| *Tarentola neglecta* | 1 | 0 | 0 | 0 |
| *Tarentola rudis* | 0 | 1 | 0 | 1 |
| *Teira dugesii* | 1 | 0 | 0 | 0 |
| *Teius teyou* | 0 | 1 | 0 | 1 |
| *Telescopus fallax* | 1 | 0 | 0 | 0 |
| *Teratolepis fasciata* | 0 | 1 | 0 | 1 |
| *Teratoscincus microlepis* | 1 | 0 | 0 | 0 |
| *Teratoscincus przewalskii* | 1 | 0 | 0 | 0 |
| *Teratoscincus roborowskii* | 1 | 0 | 0 | 0 |
| *Teratoscincus scincus* | 1 | 0 | 0 | 0 |
| *Tetradactylus africanus* | 0 | 1 | 0 | 1 |
| *Tetradactylus seps* | 0 | 1 | 0 | 1 |
| *Tetradactylus tetradactylus* | 0 | 1 | 0 | 1 |
| *Thamnodynastes hypoconia* | 0 | 1 | 0 | 1 |
| *Thamnodynastes lanei* | 0 | 1 | 0 | 1 |
| *Thamnodynastes pallidus* | 0 | 1 | 0 | 1 |
| *Thamnodynastes rutilus* | 0 | 1 | 0 | 1 |
| *Thamnodynastes strigatus* | 0 | 1 | 0 | 1 |
| *Thamnophis atratus* | 1 | 0 | 0 | 0 |
| *Thamnophis brachystoma* | 1 | 0 | 0 | 0 |
| *Thamnophis butleri* | 1 | 0 | 0 | 0 |
| *Thamnophis chrysocephalus* | 0 | 1 | 0 | 1 |
| *Thamnophis couchii* | 1 | 0 | 0 | 0 |
| *Thamnophis cyrtopsis* | 1 | 1 | 0 | 0 |
| *Thamnophis elegans* | 1 | 0 | 0 | 0 |
| *Thamnophis eques* | 1 | 0 | 0 | 0 |
| *Thamnophis exsul* | 0 | 1 | 0 | 1 |
| *Thamnophis fulvus* | 0 | 1 | 0 | 1 |
| *Thamnophis gigas* | 1 | 0 | 0 | 0 |
| *Thamnophis godmani* | 0 | 1 | 0 | 1 |
| *Thamnophis hammondii* | 1 | 0 | 0 | 0 |
| *Thamnophis marcianus* | 0 | 1 | 0 | 1 |
| *Thamnophis melanogaster* | 0 | 1 | 0 | 1 |
| *Thamnophis mendax* | 0 | 1 | 0 | 1 |
| *Thamnophis ordinoides* | 1 | 0 | 0 | 0 |
| *Thamnophis proximus* | 1 | 1 | 0 | 0 |
| *Thamnophis radix* | 1 | 0 | 0 | 0 |
| *Thamnophis rufipunctatus* | 1 | 0 | 0 | 0 |
| *Thamnophis sauritus* | 1 | 0 | 0 | 0 |
| *Thamnophis scaliger* | 0 | 1 | 0 | 1 |
| *Thamnophis sirtalis* | 1 | 0 | 0 | 0 |
| *Thamnophis sumichrasti* | 0 | 1 | 0 | 1 |
| *Thamnophis valida* | 0 | 1 | 0 | 1 |
| *Thecadactylus rapicauda* | 0 | 1 | 0 | 1 |
| *Thecadactylus solimoensis* | 0 | 1 | 0 | 1 |
| *Thelotornis capensis* | 0 | 1 | 0 | 1 |
| *Thermophis baileyi* | 1 | 0 | 0 | 0 |
| *Thermophis zhaoermii* | 1 | 0 | 0 | 0 |
| *Thrasops jacksonii* | 0 | 1 | 0 | 1 |
| *Tiliqua adelaidensis* | 0 | 1 | 0 | 1 |
| *Tiliqua gigas* | 0 | 1 | 0 | 1 |
| *Tiliqua nigrolutea* | 0 | 1 | 0 | 1 |
| *Tiliqua occipitalis* | 0 | 1 | 0 | 1 |
| *Tiliqua rugosa* | 0 | 1 | 0 | 1 |
| *Tiliqua scincoides* | 0 | 1 | 0 | 1 |
| *Timon lepidus* | 1 | 0 | 0 | 0 |
| *Timon pater* | 1 | 0 | 0 | 0 |
| *Timon princeps* | 1 | 0 | 0 | 0 |
| *Timon tangitanus* | 1 | 0 | 0 | 0 |
| *Tomodon dorsatus* | 0 | 1 | 0 | 1 |
| *Toxicocalamus loriae* | 0 | 1 | 0 | 1 |
| *Toxicocalamus preussi* | 0 | 1 | 0 | 1 |
| *Tracheloptychus madagascariensis* | 0 | 1 | 0 | 1 |
| *Tracheloptychus petersi* | 0 | 1 | 0 | 1 |
| *Trachischium monticola* | 0 | 1 | 0 | 1 |
| *Trachyboa boulengeri* | 0 | 1 | 0 | 1 |
| *Trachyboa gularis* | 0 | 1 | 0 | 1 |
| *Trachylepis acutilabris* | 0 | 1 | 0 | 1 |
| *Trachylepis affinis* | 0 | 1 | 0 | 1 |
| *Trachylepis atlantica* | 0 | 1 | 0 | 1 |
| *Trachylepis aurata* | 1 | 0 | 0 | 0 |
| *Trachylepis aureopunctata* | 0 | 1 | 0 | 1 |
| *Trachylepis boettgeri* | 0 | 1 | 0 | 1 |
| *Trachylepis brevicollis* | 0 | 1 | 0 | 1 |
| *Trachylepis capensis* | 0 | 1 | 0 | 1 |
| *Trachylepis dumasi* | 0 | 1 | 0 | 1 |
| *Trachylepis elegans* | 0 | 1 | 0 | 1 |
| *Trachylepis gravenhorstii* | 0 | 1 | 0 | 1 |
| *Trachylepis hoeschi* | 0 | 1 | 0 | 1 |
| *Trachylepis homalocephala* | 0 | 1 | 0 | 1 |
| *Trachylepis maculilabris* | 0 | 1 | 0 | 1 |
| *Trachylepis madagascariensis* | 0 | 1 | 0 | 1 |
| *Trachylepis margaritifera* | 0 | 1 | 0 | 1 |
| *Trachylepis occidentalis* | 0 | 1 | 0 | 1 |
| *Trachylepis perrotetii* | 0 | 1 | 0 | 1 |
| *Trachylepis quinquetaeniata* | 0 | 1 | 0 | 1 |
| *Trachylepis seychellensis* | 0 | 1 | 0 | 1 |
| *Trachylepis socotrana* | 0 | 1 | 0 | 1 |
| *Trachylepis spilogaster* | 0 | 1 | 0 | 1 |
| *Trachylepis striata* | 0 | 1 | 0 | 1 |
| *Trachylepis sulcata* | 0 | 1 | 0 | 1 |
| *Trachylepis varia* | 0 | 1 | 0 | 1 |
| *Trachylepis variegata* | 0 | 1 | 0 | 1 |
| *Trachylepis vato* | 0 | 1 | 0 | 1 |
| *Trachylepis vittata* | 1 | 0 | 0 | 0 |
| *Trachylepis wrightii* | 0 | 1 | 0 | 1 |
| *Trapelus agilis* | 1 | 0 | 0 | 0 |
| *Trapelus flavimaculatus* | 1 | 0 | 0 | 0 |
| *Trapelus mutabilis* | 1 | 0 | 0 | 0 |
| *Trapelus pallidus* | 1 | 0 | 0 | 0 |
| *Trapelus ruderatus* | 1 | 0 | 0 | 0 |
| *Trapelus sanguinolentus* | 1 | 0 | 0 | 0 |
| *Trapelus savignii* | 1 | 0 | 0 | 0 |
| *Tretanorhinus nigroluteus* | 0 | 1 | 0 | 1 |
| *Tretanorhinus variabilis* | 0 | 1 | 0 | 1 |
| *Tretioscincus agilis* | 0 | 1 | 0 | 1 |
| *Tretioscincus oriximinensis* | 0 | 1 | 0 | 1 |
| *Tribolonotus blanchardi* | 0 | 1 | 0 | 1 |
| *Tribolonotus brongersmai* | 0 | 1 | 0 | 1 |
| *Tribolonotus gracilis* | 0 | 1 | 0 | 1 |
| *Tribolonotus novaeguineae* | 0 | 1 | 0 | 1 |
| *Tribolonotus ponceleti* | 0 | 1 | 0 | 1 |
| *Tribolonotus pseudoponceleti* | 0 | 1 | 0 | 1 |
| *Tribolonotus schmidti* | 0 | 1 | 0 | 1 |
| *Triceratolepidophis sieversorum* | 0 | 1 | 0 | 1 |
| *Trimeresurus borneensis* | 0 | 1 | 0 | 1 |
| *Trimeresurus gracilis* | 0 | 1 | 0 | 1 |
| *Trimeresurus gramineus* | 0 | 1 | 0 | 1 |
| *Trimeresurus malabaricus* | 0 | 1 | 0 | 1 |
| *Trimeresurus puniceus* | 0 | 1 | 0 | 1 |
| *Trimeresurus trigonocephalus* | 0 | 1 | 0 | 1 |
| *Trimetopon gracile* | 0 | 1 | 0 | 1 |
| *Trimorphodon biscutatus* | 1 | 1 | 0 | 0 |
| *Trogonophis wiegmanni* | 1 | 0 | 0 | 0 |
| *Tropidechis carinatus* | 0 | 1 | 0 | 1 |
| *Tropidoclonion lineatum* | 1 | 0 | 0 | 0 |
| *Tropidodipsas sartorii* | 0 | 1 | 0 | 1 |
| *Tropidodryas serra* | 0 | 1 | 0 | 1 |
| *Tropidodryas striaticeps* | 0 | 1 | 0 | 1 |
| *Tropidolaemus wagleri* | 0 | 1 | 0 | 1 |
| *Tropidophis feicki* | 0 | 1 | 0 | 1 |
| *Tropidophis greenwayi* | 0 | 1 | 0 | 1 |
| *Tropidophis haetianus* | 0 | 1 | 0 | 1 |
| *Tropidophis melanurus* | 0 | 1 | 0 | 1 |
| *Tropidophis pardalis* | 0 | 1 | 0 | 1 |
| *Tropidophis wrighti* | 0 | 1 | 0 | 1 |
| *Tropidophorus baconi* | 0 | 1 | 0 | 1 |
| *Tropidophorus baviensis* | 0 | 1 | 0 | 1 |
| *Tropidophorus beccarii* | 0 | 1 | 0 | 1 |
| *Tropidophorus berdmorei* | 0 | 1 | 0 | 1 |
| *Tropidophorus brookei* | 0 | 1 | 0 | 1 |
| *Tropidophorus cocincinensis* | 0 | 1 | 0 | 1 |
| *Tropidophorus grayi* | 0 | 1 | 0 | 1 |
| *Tropidophorus hainanus* | 0 | 1 | 0 | 1 |
| *Tropidophorus latiscutatus* | 0 | 1 | 0 | 1 |
| *Tropidophorus matsuii* | 0 | 1 | 0 | 1 |
| *Tropidophorus microlepis* | 0 | 1 | 0 | 1 |
| *Tropidophorus misaminius* | 0 | 1 | 0 | 1 |
| *Tropidophorus murphyi* | 0 | 1 | 0 | 1 |
| *Tropidophorus noggei* | 0 | 1 | 0 | 1 |
| *Tropidophorus partelloi* | 0 | 1 | 0 | 1 |
| *Tropidophorus robinsoni* | 0 | 1 | 0 | 1 |
| *Tropidophorus sinicus* | 0 | 1 | 0 | 1 |
| *Tropidophorus thai* | 0 | 1 | 0 | 1 |
| *Tropidosaura gularis* | 0 | 1 | 0 | 1 |
| *Tropidoscincus aubrianus* | 0 | 1 | 0 | 1 |
| *Tropidoscincus boreus* | 0 | 1 | 0 | 1 |
| *Tropidoscincus variabilis* | 0 | 1 | 0 | 1 |
| *Tropidurus bogerti* | 0 | 1 | 0 | 1 |
| *Tropidurus callathelys* | 0 | 1 | 0 | 1 |
| *Tropidurus cocorobensis* | 0 | 1 | 0 | 1 |
| *Tropidurus erythrocephalus* | 0 | 1 | 0 | 1 |
| *Tropidurus etheridgei* | 0 | 1 | 0 | 1 |
| *Tropidurus hispidus* | 0 | 1 | 0 | 1 |
| *Tropidurus hygomi* | 0 | 1 | 0 | 1 |
| *Tropidurus insulanus* | 0 | 1 | 0 | 1 |
| *Tropidurus itambere* | 0 | 1 | 0 | 1 |
| *Tropidurus montanus* | 0 | 1 | 0 | 1 |
| *Tropidurus mucujensis* | 0 | 1 | 0 | 1 |
| *Tropidurus oreadicus* | 0 | 1 | 0 | 1 |
| *Tropidurus psammonastes* | 0 | 1 | 0 | 1 |
| *Tropidurus spinulosus* | 0 | 1 | 0 | 1 |
| *Tropidurus torquatus* | 0 | 1 | 0 | 1 |
| *Tropiocolotes helenae* | 1 | 0 | 0 | 0 |
| *Tropiocolotes tripolitanus* | 1 | 0 | 0 | 0 |
| *Tupinambis duseni* | 0 | 1 | 0 | 1 |
| *Tupinambis longilineus* | 0 | 1 | 0 | 1 |
| *Tupinambis merianae* | 0 | 1 | 0 | 1 |
| *Tupinambis quadrilineatus* | 0 | 1 | 0 | 1 |
| *Tupinambis rufescens* | 0 | 1 | 0 | 1 |
| *Tupinambis teguixin* | 0 | 1 | 0 | 1 |
| *Tympanocryptis cephalus* | 0 | 1 | 0 | 1 |
| *Tympanocryptis intima* | 0 | 1 | 0 | 1 |
| *Tympanocryptis lineata* | 0 | 1 | 0 | 1 |
| *Tympanocryptis pinguicolla* | 0 | 1 | 0 | 1 |
| *Tympanocryptis tetraporophora* | 0 | 1 | 0 | 1 |
| *Tympanocryptis uniformis* | 0 | 1 | 0 | 1 |
| *Typhlacontias brevipes* | 0 | 1 | 0 | 1 |
| *Typhlacontias punctatissimus* | 0 | 1 | 0 | 1 |
| *Typhlophis squamosus* | 0 | 1 | 0 | 1 |
| *Typhlops agoralionis* | 0 | 1 | 0 | 1 |
| *Typhlops anchaurus* | 0 | 1 | 0 | 1 |
| *Typhlops angolensis* | 0 | 1 | 0 | 1 |
| *Typhlops anousius* | 0 | 1 | 0 | 1 |
| *Typhlops arator* | 0 | 1 | 0 | 1 |
| *Typhlops arenarius* | 0 | 1 | 0 | 1 |
| *Typhlops bibronii* | 0 | 1 | 0 | 1 |
| *Typhlops biminiensis* | 0 | 1 | 0 | 1 |
| *Typhlops brongersmianus* | 0 | 1 | 0 | 1 |
| *Typhlops capitulatus* | 0 | 1 | 0 | 1 |
| *Typhlops catapontus* | 0 | 1 | 0 | 1 |
| *Typhlops caymanensis* | 0 | 1 | 0 | 1 |
| *Typhlops congestus* | 0 | 1 | 0 | 1 |
| *Typhlops contorhinus* | 0 | 1 | 0 | 1 |
| *Typhlops dominicanus* | 0 | 1 | 0 | 1 |
| *Typhlops elegans* | 0 | 1 | 0 | 1 |
| *Typhlops eperopeus* | 0 | 1 | 0 | 1 |
| *Typhlops fornasinii* | 0 | 1 | 0 | 1 |
| *Typhlops granti* | 0 | 1 | 0 | 1 |
| *Typhlops hectus* | 0 | 1 | 0 | 1 |
| *Typhlops hedraeus* | 0 | 1 | 0 | 1 |
| *Typhlops hypomethes* | 0 | 1 | 0 | 1 |
| *Typhlops jamaicensis* | 0 | 1 | 0 | 1 |
| *Typhlops lineolatus* | 0 | 1 | 0 | 1 |
| *Typhlops lumbricalis* | 0 | 1 | 0 | 1 |
| *Typhlops luzonensis* | 0 | 1 | 0 | 1 |
| *Typhlops mirus* | 0 | 1 | 0 | 1 |
| *Typhlops monastus* | 0 | 1 | 0 | 1 |
| *Typhlops notorachius* | 0 | 1 | 0 | 1 |
| *Typhlops pammeces* | 0 | 1 | 0 | 1 |
| *Typhlops platycephalus* | 0 | 1 | 0 | 1 |
| *Typhlops punctatus* | 0 | 1 | 0 | 1 |
| *Typhlops pushpakumara* | 0 | 1 | 0 | 1 |
| *Typhlops pusillus* | 0 | 1 | 0 | 1 |
| *Typhlops reticulatus* | 0 | 1 | 0 | 1 |
| *Typhlops richardi* | 0 | 1 | 0 | 1 |
| *Typhlops rostellatus* | 0 | 1 | 0 | 1 |
| *Typhlops ruber* | 0 | 1 | 0 | 1 |
| *Typhlops schwartzi* | 0 | 1 | 0 | 1 |
| *Typhlops sulcatus* | 0 | 1 | 0 | 1 |
| *Typhlops sylleptor* | 0 | 1 | 0 | 1 |
| *Typhlops syntherus* | 0 | 1 | 0 | 1 |
| *Typhlops titanops* | 0 | 1 | 0 | 1 |
| *Typhlops vermicularis* | 1 | 0 | 0 | 0 |
| *Typhlosaurus braini* | 0 | 1 | 0 | 1 |
| *Typhlosaurus caecus* | 0 | 1 | 0 | 1 |
| *Typhlosaurus gariepensis* | 0 | 1 | 0 | 1 |
| *Typhlosaurus lineatus* | 0 | 1 | 0 | 1 |
| *Typhlosaurus lomiae* | 0 | 1 | 0 | 1 |
| *Typhlosaurus meyeri* | 0 | 1 | 0 | 1 |
| *Typhlosaurus vermis* | 0 | 1 | 0 | 1 |
| *Uma exsul* | 1 | 0 | 0 | 0 |
| *Uma inornata* | 1 | 0 | 0 | 0 |
| *Uma notata* | 1 | 0 | 0 | 0 |
| *Uma paraphygas* | 1 | 0 | 0 | 0 |
| *Uma scoparia* | 1 | 0 | 0 | 0 |
| *Umbrivaga pygmaea* | 0 | 1 | 0 | 1 |
| *Underwoodisaurus milii* | 0 | 1 | 0 | 1 |
| *Underwoodisaurus sphyrurus* | 0 | 1 | 0 | 1 |
| *Ungaliophis continentalis* | 0 | 1 | 0 | 1 |
| *Uracentron flaviceps* | 0 | 1 | 0 | 1 |
| *Uranoscodon superciliosus* | 0 | 1 | 0 | 1 |
| *Urocotyledon inexpectata* | 0 | 1 | 0 | 1 |
| *Uromacer catesbyi* | 0 | 1 | 0 | 1 |
| *Uromacer frenatus* | 0 | 1 | 0 | 1 |
| *Uromacer oxyrhynchus* | 0 | 1 | 0 | 1 |
| *Uromastyx acanthinura* | 1 | 1 | 0 | 0 |
| *Uromastyx aegyptia* | 1 | 0 | 0 | 0 |
| *Uromastyx asmussi* | 1 | 0 | 0 | 0 |
| *Uromastyx benti* | 0 | 1 | 0 | 1 |
| *Uromastyx dispar* | 0 | 1 | 0 | 1 |
| *Uromastyx geyri* | 0 | 1 | 0 | 1 |
| *Uromastyx hardwickii* | 1 | 0 | 0 | 0 |
| *Uromastyx leptieni* | 1 | 0 | 0 | 0 |
| *Uromastyx loricata* | 1 | 0 | 0 | 0 |
| *Uromastyx macfadyeni* | 0 | 1 | 0 | 1 |
| *Uromastyx ocellata* | 0 | 1 | 0 | 1 |
| *Uromastyx ornata* | 1 | 0 | 0 | 0 |
| *Uromastyx princeps* | 0 | 1 | 0 | 1 |
| *Uromastyx thomasi* | 1 | 0 | 0 | 0 |
| *Uromastyx yemenensis* | 0 | 1 | 0 | 1 |
| *Uropeltis ceylanicus* | 0 | 1 | 0 | 1 |
| *Uropeltis liura* | 0 | 1 | 0 | 1 |
| *Uropeltis melanogaster* | 0 | 1 | 0 | 1 |
| *Uropeltis phillipsi* | 0 | 1 | 0 | 1 |
| *Uroplatus alluaudi* | 0 | 1 | 0 | 1 |
| *Uroplatus ebenaui* | 0 | 1 | 0 | 1 |
| *Uroplatus fimbriatus* | 0 | 1 | 0 | 1 |
| *Uroplatus giganteus* | 0 | 1 | 0 | 1 |
| *Uroplatus guentheri* | 0 | 1 | 0 | 1 |
| *Uroplatus henkeli* | 0 | 1 | 0 | 1 |
| *Uroplatus lineatus* | 0 | 1 | 0 | 1 |
| *Uroplatus malahelo* | 0 | 1 | 0 | 1 |
| *Uroplatus malama* | 0 | 1 | 0 | 1 |
| *Uroplatus phantasticus* | 0 | 1 | 0 | 1 |
| *Uroplatus pietschmanni* | 0 | 1 | 0 | 1 |
| *Uroplatus sikorae* | 0 | 1 | 0 | 1 |
| *Urosaurus auriculatus* | 0 | 1 | 0 | 1 |
| *Urosaurus bicarinatus* | 0 | 1 | 0 | 1 |
| *Urosaurus clarionensis* | 0 | 1 | 0 | 1 |
| *Urosaurus gadovi* | 0 | 1 | 0 | 1 |
| *Urosaurus graciosus* | 1 | 0 | 0 | 0 |
| *Urosaurus lahtelai* | 1 | 0 | 0 | 0 |
| *Urosaurus nigricaudus* | 1 | 0 | 0 | 0 |
| *Urosaurus ornatus* | 1 | 0 | 0 | 0 |
| *Urostrophus gallardoi* | 0 | 1 | 0 | 1 |
| *Urostrophus vautieri* | 0 | 1 | 0 | 1 |
| *Uta palmeri* | 1 | 0 | 0 | 0 |
| *Uta squamata* | 1 | 0 | 0 | 0 |
| *Uta stansburiana* | 1 | 0 | 0 | 0 |
| *Uta stejnegeri* | 1 | 0 | 0 | 0 |
| *Vanzosaura rubricauda* | 0 | 1 | 0 | 1 |
| *Varanus acanthurus* | 0 | 1 | 0 | 1 |
| *Varanus albigularis* | 0 | 1 | 0 | 1 |
| *Varanus baritji* | 0 | 1 | 0 | 1 |
| *Varanus beccarii* | 0 | 1 | 0 | 1 |
| *Varanus bengalensis* | 0 | 1 | 0 | 1 |
| *Varanus boehmei* | 0 | 1 | 0 | 1 |
| *Varanus brevicauda* | 0 | 1 | 0 | 1 |
| *Varanus bushi* | 0 | 1 | 0 | 1 |
| *Varanus caerulivirens* | 0 | 1 | 0 | 1 |
| *Varanus caudolineatus* | 0 | 1 | 0 | 1 |
| *Varanus cerambonensis* | 0 | 1 | 0 | 1 |
| *Varanus doreanus* | 0 | 1 | 0 | 1 |
| *Varanus dumerilii* | 0 | 1 | 0 | 1 |
| *Varanus eremius* | 0 | 1 | 0 | 1 |
| *Varanus exanthematicus* | 0 | 1 | 0 | 1 |
| *Varanus finschi* | 0 | 1 | 0 | 1 |
| *Varanus flavescens* | 0 | 1 | 0 | 1 |
| *Varanus giganteus* | 0 | 1 | 0 | 1 |
| *Varanus gilleni* | 0 | 1 | 0 | 1 |
| *Varanus glauerti* | 0 | 1 | 0 | 1 |
| *Varanus glebopalma* | 0 | 1 | 0 | 1 |
| *Varanus gouldii* | 0 | 1 | 0 | 1 |
| *Varanus griseus* | 1 | 0 | 0 | 0 |
| *Varanus indicus* | 0 | 1 | 0 | 1 |
| *Varanus jobiensis* | 0 | 1 | 0 | 1 |
| *Varanus keithhornei* | 0 | 1 | 0 | 1 |
| *Varanus kingorum* | 0 | 1 | 0 | 1 |
| *Varanus komodoensis* | 0 | 1 | 0 | 1 |
| *Varanus macraei* | 0 | 1 | 0 | 1 |
| *Varanus marmoratus* | 0 | 1 | 0 | 1 |
| *Varanus melinus* | 0 | 1 | 0 | 1 |
| *Varanus mertensi* | 0 | 1 | 0 | 1 |
| *Varanus mitchelli* | 0 | 1 | 0 | 1 |
| *Varanus niloticus* | 0 | 1 | 0 | 1 |
| *Varanus olivaceus* | 0 | 1 | 0 | 1 |
| *Varanus panoptes* | 0 | 1 | 0 | 1 |
| *Varanus pilbarensis* | 0 | 1 | 0 | 1 |
| *Varanus prasinus* | 0 | 1 | 0 | 1 |
| *Varanus primordius* | 0 | 1 | 0 | 1 |
| *Varanus rainerguentheri* | 0 | 1 | 0 | 1 |
| *Varanus rosenbergi* | 0 | 1 | 0 | 1 |
| *Varanus rudicollis* | 0 | 1 | 0 | 1 |
| *Varanus salvadorii* | 0 | 1 | 0 | 1 |
| *Varanus salvator* | 0 | 1 | 0 | 1 |
| *Varanus scalaris* | 0 | 1 | 0 | 1 |
| *Varanus semiremex* | 0 | 1 | 0 | 1 |
| *Varanus spenceri* | 0 | 1 | 0 | 1 |
| *Varanus storri* | 0 | 1 | 0 | 1 |
| *Varanus timorensis* | 0 | 1 | 0 | 1 |
| *Varanus tristis* | 0 | 1 | 0 | 1 |
| *Varanus varius* | 0 | 1 | 0 | 1 |
| *Varanus yemenensis* | 0 | 1 | 0 | 1 |
| *Varanus yuwonoi* | 0 | 1 | 0 | 1 |
| *Vermicella intermedia* | 0 | 1 | 0 | 1 |
| *Vipera albizona* | 1 | 0 | 0 | 0 |
| *Vipera ammodytes* | 1 | 0 | 0 | 0 |
| *Vipera aspis* | 1 | 0 | 0 | 0 |
| *Vipera barani* | 1 | 0 | 0 | 0 |
| *Vipera berus* | 1 | 0 | 0 | 0 |
| *Vipera bornmuelleri* | 1 | 0 | 0 | 0 |
| *Vipera dinniki* | 1 | 0 | 0 | 0 |
| *Vipera eriwanensis* | 1 | 0 | 0 | 0 |
| *Vipera kaznakovi* | 1 | 0 | 0 | 0 |
| *Vipera latastei* | 1 | 0 | 0 | 0 |
| *Vipera lotievi* | 1 | 0 | 0 | 0 |
| *Vipera nikolskii* | 1 | 0 | 0 | 0 |
| *Vipera palaestinae* | 1 | 0 | 0 | 0 |
| *Vipera raddei* | 1 | 0 | 0 | 0 |
| *Vipera renardi* | 1 | 0 | 0 | 0 |
| *Vipera seoanei* | 1 | 0 | 0 | 0 |
| *Vipera ursinii* | 1 | 0 | 0 | 0 |
| *Vipera wagneri* | 1 | 0 | 0 | 0 |
| *Vipera xanthina* | 1 | 0 | 0 | 0 |
| *Virginia striatula* | 1 | 0 | 0 | 0 |
| *Viridovipera gumprechti* | 0 | 1 | 0 | 1 |
| *Viridovipera medoensis* | 0 | 1 | 0 | 1 |
| *Viridovipera stejnegeri* | 0 | 1 | 0 | 1 |
| *Viridovipera vogeli* | 0 | 1 | 0 | 1 |
| *Viridovipera yunnanensis* | 0 | 1 | 0 | 1 |
| *Voeltzkowia fierinensis* | 0 | 1 | 0 | 1 |
| *Voeltzkowia lineata* | 0 | 1 | 0 | 1 |
| *Voeltzkowia rubrocaudata* | 0 | 1 | 0 | 1 |
| *Waglerophis merremi* | 0 | 1 | 0 | 1 |
| *Walterinnesia aegyptia* | 1 | 0 | 0 | 0 |
| *Xantusia arizonae* | 1 | 0 | 0 | 0 |
| *Xantusia bezyi* | 1 | 0 | 0 | 0 |
| *Xantusia bolsonae* | 1 | 0 | 0 | 0 |
| *Xantusia gracilis* | 1 | 0 | 0 | 0 |
| *Xantusia henshawi* | 1 | 0 | 0 | 0 |
| *Xantusia riversiana* | 1 | 0 | 0 | 0 |
| *Xantusia sanchezi* | 1 | 0 | 0 | 0 |
| *Xantusia vigilis* | 1 | 0 | 0 | 0 |
| *Xantusia wigginsi* | 1 | 0 | 0 | 0 |
| *Xenagama taylori* | 0 | 1 | 0 | 1 |
| *Xenocalamus transvaalensis* | 0 | 1 | 0 | 1 |
| *Xenochrophis asperrimus* | 0 | 1 | 0 | 1 |
| *Xenochrophis flavipunctatus* | 0 | 1 | 0 | 1 |
| *Xenochrophis piscator* | 0 | 1 | 0 | 1 |
| *Xenochrophis punctulatus* | 0 | 1 | 0 | 1 |
| *Xenochrophis vittatus* | 0 | 1 | 0 | 1 |
| *Xenodermus javanicus* | 0 | 1 | 0 | 1 |
| *Xenodon guentheri* | 0 | 1 | 0 | 1 |
| *Xenodon neuwiedii* | 0 | 1 | 0 | 1 |
| *Xenodon severus* | 0 | 1 | 0 | 1 |
| *Xenodon werneri* | 0 | 1 | 0 | 1 |
| *Xenopeltis unicolor* | 0 | 1 | 0 | 1 |
| *Xenophidion schaeferi* | 0 | 1 | 0 | 1 |
| *Xenopholis scalaris* | 0 | 1 | 0 | 1 |
| *Xenopholis undulatus* | 0 | 1 | 0 | 1 |
| *Xenosaurus grandis* | 0 | 1 | 0 | 1 |
| *Xenosaurus platyceps* | 0 | 1 | 0 | 1 |
| *Xenotyphlops grandidieri* | 0 | 1 | 0 | 1 |
| *Xenoxybelis argenteus* | 0 | 1 | 0 | 1 |
| *Xenoxybelis boulengeri* | 0 | 1 | 0 | 1 |
| *Zamenis hohenackeri* | 1 | 0 | 0 | 0 |
| *Zamenis lineata* | 1 | 0 | 0 | 0 |
| *Zamenis longissimus* | 1 | 0 | 0 | 0 |
| *Zamenis persica* | 1 | 0 | 0 | 0 |
| *Zamenis situla* | 1 | 0 | 0 | 0 |
| *Zhaoermia mangshanensis* | 0 | 1 | 0 | 1 |
| *Zonosaurus aeneus* | 0 | 1 | 0 | 1 |
| *Zonosaurus anelanelany* | 0 | 1 | 0 | 1 |
| *Zonosaurus bemaraha* | 0 | 1 | 0 | 1 |
| *Zonosaurus boettgeri* | 0 | 1 | 0 | 1 |
| *Zonosaurus brygooi* | 0 | 1 | 0 | 1 |
| *Zonosaurus haraldmeieri* | 0 | 1 | 0 | 1 |
| *Zonosaurus karsteni* | 0 | 1 | 0 | 1 |
| *Zonosaurus laticaudatus* | 0 | 1 | 0 | 1 |
| *Zonosaurus madagascariensis* | 0 | 1 | 0 | 1 |
| *Zonosaurus ornatus* | 0 | 1 | 0 | 1 |
| *Zonosaurus quadrilineatus* | 0 | 1 | 0 | 1 |
| *Zonosaurus rufipes* | 0 | 1 | 0 | 1 |
| *Zonosaurus subunicolor* | 0 | 1 | 0 | 1 |
| *Zonosaurus trilineatus* | 0 | 1 | 0 | 1 |
| *Zonosaurus tsingy* | 0 | 1 | 0 | 1 |
| *Zootoca vivipara* | 1 | 0 | 0 | 0 |
