## Supplementary Table 3 for "Ancient tropical extinctions contributed to the latitudinal diversity gradient"

**Table S3.** Geographic distribution of extant crocodile species. For each species, we coded its geographic range as defined by 3 discrete areas. *Abbreviations for the 3 areas: North Temp. and South Temp. = Northern (Southern) temperate regions.*

| Species | North Temp. | Tropical | South Temp. |
| --- | --- | --- | --- |
| *Alligator mississippiensis* | 1 | 0 | 0 |
| *Alligator sinensis* | 1 | 0 | 0 |
| *Caiman crocodilus* | 0 | 1 | 0 |
| *Caiman latirostris* | 0 | 1 | 0 |
| *Caiman yacare* | 0 | 1 | 0 |
| *Crocodylus acutus* | 0 | 1 | 0 |
| *Crocodylus intermedius* | 0 | 1 | 0 |
| *Crocodylus johnsoni* | 0 | 1 | 0 |
| *Crocodylus mindorensis* | 0 | 1 | 0 |
| *Crocodylus moreletii* | 0 | 1 | 0 |
| *Crocodylus niloticus* | 0 | 1 | 0 |
| *Crocodylus niloticus* | 0 | 1 | 0 |
| *Crocodylus novaeguineae* | 0 | 1 | 0 |
| *Crocodylus palustris* | 0 | 1 | 0 |
| *Crocodylus porosus* | 0 | 1 | 0 |
| *Crocodylus rhombifer* | 0 | 1 | 0 |
| *Crocodylus siamensis* | 0 | 1 | 0 |
| *Gavialis gangeticus* | 0 | 1 | 0 |
| *Mecistops cataphractus* | 0 | 1 | 0 |
| *Melanosuchus niger* | 0 | 1 | 0 |
| *Osteolaemus* *tetraspis* | 0 | 1 | 0 |
| *Osteolaemus* *tetraspis* | 0 | 1 | 0 |
| *Paleosuchus palpebrosus* | 0 | 1 | 0 |
| *Paleosuchus trigonatus* | 0 | 1 | 0 |
| *Tomistoma* *schlegelii* | 0 | 1 | 0 |
