## Supplementary Table 7 for "Ancient tropical extinctions contributed to the latitudinal diversity gradient"

**Table S7.** **Fossil constraints used for the biogeographic analyses of turtles.** Based on fossil occurrences assigned to a given clade and to the estimated age of the clade, we found 23 nodes to which fossil constraints could be applied on the turtle phylogeny (see *Methods* for more details on the procedure). Abbreviations for fossil ranges: a = Holarctic, b =Equatorial and subtropical regions, and c = southern regions of the Southern Hemisphere. MRCA = most recent common ancestor, and Ma = million years ago.

| **Node number on tree (Fig. 5b)** | **MRCA** | **Node age**  **(Ma)** | **Fossil range** | **Fossil ages**  **(periods, epochs)** | **Number of fossil occurrences** |
| --- | --- | --- | --- | --- | --- |
| 1 | Testudines | 209 | a + b + c | Late Triassic-Early Jurassic | 8 |
| 2 | Trionychidae | 123.29 | a | Early Cretaceous | 19 |
| 3 | Dermatemydiiae + Kinosternidae | 96.75 | a | Early Cretaceous | 9 |
| 4 | Chelidae | 85 | c | Early Cretaceous | 7 |
| 5 | Carettochelyidae | 150.5 | a | Late Jurassic-Early Cretaceous | 7 |
| 6 | Podocnemididae | 103.71 | b | Late Cretaceous | 14 |
| 7 | Pelomedusidae | 141.43 | a + b | Early-Late Cretaceous | 17 |
| 8 | Cryptodira | 162.24 | a | Middle-Late Jurassic | 37 |
| 9 | *Chelodina* | 57.1 | a | Late Cretaceous | 3 |
| 10 | Cheloniidae | 68.48 | a | Late Cretaceous | 29 |
| 11 | Playusternidae + Chelydridae | 72.8 | a | Late Cretaceous | 32 + 119 |
| 12 | Emydidae | 38.79 | a | Eocene | 73 |
| 13 | Testudinoidea | 77.27 | a | Late Cretaceous | 17 |
| 14 | Testudinidae | 51.54 | a | Eocene | 69 |
| 15 | *Manoura* + *Gopherus* | 39.98 | a | Eocene | 1 |
| 16 | *Testudo* | 29.37 | a | Eocene-Oligocene | 5 |
| 17 | *Geochelone* | 28.62 | a | late Eocene-Oligocene | 9 |
| 18 | Geoemydidae | 57.47 | a | Eocene | 124 |
| 19 | *Rhinoclemmys* | 26.71 | a + b | late Eocene-Oligocene | 4 |
| 20 | *Geoemyda* | 18.44 | a + b | Oligocene-Miocene | 5 |
| 21 | *Podocnemis* | 47.67 | a + b | Eocene | 11 |
| 22 | Trionychinae | 76.62 | a | Late Cretaceous | 192 |
| 23 | Pleurodira | 193.94 | a + b | Late Jurassic | 7 |
