## Supplementary Table 8 for "Ancient tropical extinctions contributed to the latitudinal diversity gradient"

**Table S8.** **Fossil constraints used for the biogeographic analyses of Squamata**. Based on fossil occurrences assigned to a given clade and to the estimated age of the clade, we found 30 nodes to which fossil constraints could be applied on the squamates phylogeny (see *Methods* for more details on the procedure). Abbreviations for fossil ranges: a = Holarctic, b =Equatorial and subtropical regions, and c = southern regions of the Southern Hemisphere. MRCA = most recent common ancestor, and Ma = million years ago.

| **Node number on tree (Appendix 1)** | **MRCA** | **Node age**  **(Ma)** | **Fossil range** | **Fossil ages**  **(periods, epochs)** | **Number of fossil occurrences** |
| --- | --- | --- | --- | --- | --- |
| 4163 | Lepidosauria | 228 | a + b + c | Middle-Late Triassic | 180 |
| 4164 | Squamata | 174.1 | a + b | Early-Middle Jurassic | 32 |
| 4173 | Gekkota | 86.5 | a | Late Cretaceous | 6 |
| 7643 | Serpentes: Elapidae | 42 | a | Eocene (Ypresian) | 1 |
| 7234 | Serpentes: Loxocemidae | 44.6 | a | Eocene (Lutetian-Priabonian) | 6 |
| 7183 | Serpentes: Boidae | 63.3 | a + b | Paleocene (Danian, Selandian) | 12 |
| 7172 | Serpentes: Tropidophiidae | 33 | a + b | Paleocene | 48 |
| 7063 | Serpentes | 131 | a | Late Jurassic-Early Cretaceous | 5 |
| 7170 | Alethinophidia | 90.3 | a + b + c | Late Cretaceous | 47 |
| 6851 | Iguania: Corytophanidae | 42.4 | a | Eocene (Ypresian) | 1 |
| 6703 | Iguania: Polychrotidae | 44.14 | a + b | Eocene (Ypresian-Priabonian) | 4 |
| 6702 | Iguania: Hoplocercidae stem | 81.2 | a | Late Cretaceous (Campanian) | 3 |
| 6540 | Iguania: Iguanidae | 66 | a + b + c | Late Cretaceous–Paleocene | 13 |
| 6238 | Iguania: Agamidae | 110 | a | Late Cretaceous (Cenomanian) | 2 |
| 6097 | Iguania: Chamaeleonidae | 66 | a | Late Cretaceous | 1 |
| 6043 | Platynota: Varanidae | 32 | a | Paleogene | 4 |
| 6008 | Diploglossa: Anguidae | 53.7 | a | Paleocene, Ypresian | 248 |
| 5999 | Platynota: Helodermatidae | 20.2 | a | Oligocene | 4 |
| 5997 | Diploglossa: Xenosauridae | 16 | a | Miocene | 1 |
| 4835 | Scincomorpha: Xantusiidae | 103 | a | Late Cretaceous (Albian) | 1 |
| 4888 | Scincomorpha: Cordylidae | 67 | a | Late Cretaceous (Maastrichtian) | 3 |
| 4929 | Scincomorpha: Scincidae | 94 | a | Late Cretaceous | 5 |
| 8090 | Serpentes: Thamnophis | 8 | a | Miocene | 18 |
| 7682 | Serpentes: Naja | 20 | a + c | Miocene | 21 |
| 7650 | Serpentes: Micrurus | 21.7 | a | Miocene | 9 |
| 7327 | Serpentes: Bitis | 23 | c | Miocene | 1 |
| 5612 | Lacertoidea | 154 | a + b | Late Jurassic-Early Cretaceous | 11 |
| 5757 | Amphisbaena | 120 | a | Early Cretaceous | 1 |
| 5808 | Lacertidae | 84.4 | a | Early Cretaceous | 1 |
| 5614 | Teiidae | 82.5 | a | Late Cretaceous | 27 |
