## Supplementary Table 9 for "Ancient tropical extinctions contributed to the latitudinal diversity gradient"

**Table S9.** **Fossil constraints used for the biogeographic analyses of Crocodiles**. Based on fossil occurrences assigned to a given clade and to the estimated age of the clade, we found 8 nodes to which fossil constraints could be applied on the crocodilian phylogeny (see *Methods* for more details on the procedure). Abbreviations for fossil ranges: a = Holarctic, b =Equatorial and subtropical regions, and c = southern regions of the Southern Hemisphere. MRCA = most recent common ancestor, and Ma = million years ago.

| Node number on tree (Fig. S15) | MRCA | Node age  (Ma) | Fossil range | Fossil ages  (periods, epochs) | Number of fossil occurrences |
| --- | --- | --- | --- | --- | --- |
| 24 | Crocodilia | 87.14 | a + b + c | Late Cretaceous | 694 |
| 7 | Alligatoridae | 65.5 | a + c | Paleocene | 43 |
| 21 | Crocodylidae | 22.10 | a + b + c | Miocene | 128 |
| 22 | Gavialidae | 23.23 | b | Miocene | 22 |
| 23 | Crocodylidae + Gavialidae | 54.53 | a | Paleocene-Ypresian | 56 |
| 6 | *Alligator* | 47.25 | a | Eocene | 15 |
| 2 | *Caiman* | 8.3 | b | Miocene | 26 |
| 18 | *Crocodylus* | 11 | a + b + c | Miocene | 60 |
