## Supplementary Table 10 for "Ancient tropical extinctions contributed to the latitudinal diversity gradient"

**Table S10.** **Estimated number of dispersals events through time** from the Holarctic into the equator and out of the equatorial zone for Testudines, Squamata and Crocodiles under the *unconstrained* (Unc.), based on present evidence only, and the fossil-based *hard* (HFC) and *soft fossil constraint* (SFC) biogeographic models. To calculate the number of dispersal events through time, we divided each phylogeny in 25 Ma intervals, and calculated the number of branches in which range expansion was inferred that cross a particular time interval. Note that in DEC, extinction and dispersal are modelled as anagenetic processes that occur across the branches of the tree, with an equal probability to occur across all the branch length (Ree and Smith, 2008). To account for this uncertainty in the position of an event, when dispersal was estimated in a branch crossing two or more continuous intervals, we counted this event for all time intervals concerned.

| Organism | Dispersal direction | Model | 200 | 175 | 150 | 125 | 100 | 75 | 50 | 25 | 0 | Total |
| --- | --- | --- | --- | --- | --- | --- | --- | --- | --- | --- | --- | --- |
| Testudines | Into the Equator | Unc. | 0 | 0 | 0 | 0 | 0 | 0 | 0 | 0 | 3 | **3** |
|  |  | SFC | 1 | 3 | 3 | 2 | 2 | 4 | 3 | 0 | 3 | **21** |
|  |  | HFC | 0 | 0 | 0 | 1 | 2 | 4 | 10 | 9 | 10 | **36** |
|  | Out of the Equator | Unc. | 0 | 0 | 0 | 0 | 0 | 2 | 3 | 4 | 5 | **14** |
|  |  | SFC | 0 | 0 | 0 | 0 | 0 | 0 | 0 | 3 | 5 | **8** |
|  |  | HFC | 0 | 0 | 0 | 0 | 0 | 1 | 1 | 1 | 6 | **9** |
| Squamates | Into the Equator | Unc. | - | - | 0 | 0 | 0 | 0 | 2 | 8 | 30 | **40** |
|  |  | SFC | - | - | 2 | 5 | 9 | 5 | 11 | 20 | 40 | **92** |
|  |  | HFC | - | - | 0 | 2 | 6 | 20 | 23 | 31 | 42 | **124** |
|  | Out of the Equator | Unc. | - | - | 0 | 0 | 3 | 8 | 21 | 44 | 78 | **154** |
|  |  | SFC | - | - | 0 | 1 | 1 | 5 | 24 | 38 | 71 | **140** |
|  |  | HFC | - | - | 0 | 0 | 0 | 3 | 19 | 41 | 85 | **148** |
| Crocodiles | Into the Equator | Unc. | - | - | - | - | - | 0 | 0 | 0 | 0 | **0** |
|  |  | SFC | - | - | - | - | - | 0 | 3 | 3 | 2 | **8** |
|  |  | HFC | - | - | - | - | - | 0 | 2 | 2 | 2 | **6** |
|  | Out of the Equator | Unc. | - | - | - | - | - | 0 | 0 | 0 | 0 | **0** |
|  |  | SFC | - | - | - | - | - | 0 | 0 | 0 | 0 | **0** |
|  |  | HFC | - | - | - | - | - | 0 | 0 | 0 | 0 | **0** |
