## Supplementary Table 14 for "Ancient tropical extinctions contributed to the latitudinal diversity gradient"

**Table S14**. Uncertainty of the biogeographic reconstruction of Crocodilia inferred from Lagrange (DEC) under three different models (unconstrained, Unc.; soft fossil constraint, SFC; and hard fossil constraint model, HFC; see methods) on the 10 most basal nodes of the tree. Relative probabilities for the best and the second most supported ancestral scenario are provided. Ancestral scenarios are indicated with letters that correspond with our three operational areas: Holarctic (a), equator (b), southern temperate regions (c). In the ancestral scenarios, the symbol “|” separates the ranges inherited by each descendant lineage. Node numbers correspond with the tree supplied on Appendix 1. Results for all nodes on the tree are provided on Appendix 1.

| node | model | Best scenario | Relative prob. | Second best | Relative prob. |
| --- | --- | --- | --- | --- | --- |
| 24 | Unc. | a_b\|b | 0.982 | - | - |
|  | SFC | a\|a | 0.381 | a\|a_b | 0.235 |
|  | HFC | a_b_c\|a | 0.630 | b_c\|a | 0.104 |
| 7 | Unc. | b\|a | 0.970 | - | - |
|  | SFC | a\|a | 0.617 | a_b\|a | 0.173 |
|  | HFC | c\|a | 0.490 | a_c\|a | 0.352 |
| 23 | Unc. | b\|b | 0.999 | - | - |
|  | SFC | a\|a | 0.360 | a\|a_b | 0.142 |
|  | HFC | a\|a | 1 | - | - |
| 5 | Unc. | b\|b | 0.997 | - | - |
|  | SFC | a_b\|b | 0.709 | b\|b | 0.229 |
|  | HFC | b\|b | 0.830 | - | - |
| 6 | Unc. | a\|a | 0.994 | - | - |
|  | SFC | a\|a | 0.973 | - | - |
|  | HFC | a\|a | 1 | - | - |
| 21 | Unc. | b\|b | 0.999 | - | - |
|  | SFC | a_b\|b | 0.951 | - | - |
|  | HFC | a_b_c\|b | 0.898 | - | - |
| 22 | Unc. | b\|b | 0.999 | - | - |
|  | SFC | a_b\|b | 0.469 | - | - |
|  | HFC | b\|b | 1 | - | - |
| 3 | Unc. | b\|b | 0.999 | - | - |
|  | SFC | b\|a_b | 0.819 | b\|b | 0.162 |
|  | HFC | b\|b | 0.982 | - | - |
| 4 | Unc. | b\|b | 0.999 | - | - |
|  | SFC | b\|b | 0.994 | - | - |
|  | HFC | b\|b | 0.979 | - | - |
| 18 | Unc. | b\|b | 0.999 | - | - |
|  | SFC | a_b\|b | 0.613 | b\|a_b | 0.380 |
|  | HFC | a_b_c\|b | 0.716 | b\|a_b_c | 0.265 |
