## Supplementary Table 15 for "Ancient tropical extinctions contributed to the latitudinal diversity gradient"

**Table S15.** Number of fossil occurrences (occ.) per genera in Equatorial, Tropical, Temperate and Holarctic datasets for each group (occurrences for the Southern Hemisphere temperate regions not shown).

|  | **Total**  **(occ. / genera)** | **Equatorial**  **(occ. / genera)** | **Holarctic**  **(occ. / genera)** | **Tropical**  **(occ. / genera)** | **Temperate**  **(occ. / genera)** |
| --- | --- | --- | --- | --- | --- |
| **Testudines** | 4083/422 = 9.7 | 429/123 = 3.5 | 3568/320 = 11.2 | 2996/360 = 8.3 | 993/85 = 11.7 |
| **Squamata** | 4798/638 = 7.5 | 307/106 = 2.9 | 4289/525 = 8.2 | 2428/470 = 5.2 | 2168/174 = 12.5 |
| **Crocodilia** | 1596/121 = 13.2 | 509/53 = 9.6 | 990/70 = 14.1 | 1267/105 = 12.1 | 237/16 = 14.8 |
